# Distinct mutational processes shape selection of MHC class I and class II mutations across primary and metastatic tumors

**DOI:** 10.1101/2023.01.22.523447

**Authors:** Michael B. Mumphrey, Noshad Hosseini, Abhijit Parolia, Jie Geng, Weiping Zou, Malini Raghavan, Arul Chinnaiyan, Marcin Cieslik

**Author notes:** Co-senior authors (Arul Chinnaiyan, Marcin Cieslik). Correspondence: Marcin Cieslik.

## Abstract

Disruption of antigen presentation via loss of MHC expression is a strategy whereby cancer cells escape immune surveillance and develop resistance to immunotherapy. We developed the personalized genomics algorithm Hapster and accurately called somatic mutations within the MHC genes of 10,001 primary and 2,199 metastatic tumors, creating a catalog of 1663 nonsynonymous mutations that provide key insights into MHC mutagenesis. We found that MHC-I genes are among the most frequently mutated genes in both primary and metastatic tumors, while MHC-II mutations are more restricted. Recurrent deleterious mutations are found within haplotype and cancer-type specific hotspots associated with distinct mutational processes. Functional classification of MHC residues revealed significant positive selection for mutations disruptive to the B2M, peptide, and T-cell binding interfaces, as well as MHC chaperones. At the cohort level, all cancers with positive selection for MHC mutations are responsive to immune checkpoint inhibitors, underscoring the translational relevance of our findings.

## 1. Introduction

The immune system is capable of identifying and eliminating cancer cells via CD8^+^ T cell mediated cytotoxicity ^1^. This process has been described as the cancer immunity cycle ^2^ and when functioning provides a defense against malignant cells. To avoid destruction, successful cancers often evolve strategies to disrupt the cancer immunity cycle, such as overexpression of the immunosuppressive PD-L1 ^3^ or reduced expression of key immune system proteins such as the MHC class I molecules ^4^. Research into how tumors escape T cell surveillance has led to a breakthrough in immunotherapies that restore cancer immunity in select patients ^5–10^. However, even the most promising immunotherapies still only provide a clinical benefit in a minority of cases ^11^. Understanding the molecular and genetic mechanisms that lead to primary and acquired resistance to T cell based immunotherapies is critical for the continued improvement of patient outcomes.

T cells are able to identify malignant cells via the presence of antigenic mutant peptides known as neoantigens ^1^. In order to detect these neoantigens, T cells require the peptides to be presented at the cell surface by the MHC. The MHC class I proteins present neoantigens to cytotoxic CD8^+^ T cells ^1^, and as a result are directly involved in the destruction of malignant cells. Consistent with this role as a tumor suppressor, correlations between decreased MHC class I expression and poor outcomes for cancer patients have been repeatedly observed across multiple cancer types ^12–16^. The MHC class II proteins are less well studied in cancers, but evidence suggests that they also play an important role in tumor suppression ^17–19^. The MHC class II proteins are responsible for presenting neoantigens to CD4^+^ T cells which have both regulatory and effector functions and have been shown to play a role in tumor immunity ^20^. Classically, MHC class II expression was thought to be restricted to professional antigen presenting cells (APCs). However, their expression can be induced in most cell types ^21^, including cancer cells ^22–24^, and MHC class II-restricted neoantigen vaccines have been shown to produce APCs from cancer cells ^25^. In this context, the MHC class II expressing cancer cells promote an anti-tumor response suggesting that loss of MHC class II function may also promote tumor survival. However, this process is not well understood, and may only be relevant to a subset of cancers.

Given the central role of MHC class I and class II proteins in antigen presentation, identifying the molecular and genetic mechanisms that lead to loss of the MHC is key to understanding primary and acquired resistance to T cell based immunotherapies. However, genetic studies of the MHC are hindered by the extreme polymorphism of the MHC genes ^26^. We have addressed this issue and created Hapster, a generalized algorithm that constructs personalized reference sequences to allow for high quality alignment and mutation calling within polymorphic genes. The high sensitivity and specificity of the identified mutations enabled us to study the mutational and evolutionary processes driving pervasive MHC mutational loss in primary and metastatic tumors.

## 2. Results

### 2.1 Hapster allows for more sensitive and specific somatic mutation calls in MHC genes

Hapster is a complete mutation calling pipeline that performs three fundamental functions: (1) selection of personalized reference haplotypes, (2) pruning of contaminant and misaligned reads, and (3) detection ^27,28^ of variants and filtering of false positives that may have been called due to remaining misalignments or sample contamination (**Figure 1A**). For the first function, while in principle any of the many existing HLA haplotypers ^29–35^ could be used to identify HLA haplotype sequences, in practice existing haplotypers often report HLA types for which only the sequence for exons 2 and 3 are known ^36^, which precludes somatic mutation calling as we aim to identify mutations in all exons and introns (**Supplementary figure S1A**). Further, some leading haplotypers such as OptiType ^29^ cannot type the MHC class II genes. We therefore developed a generalized haplotyping algorithm to guarantee the return of full length genomic sequences for MHC class I and II genes. For the second and third functions, we developed MHC specific strategies for pruning of contaminant and misaligned reads originating from other MHC genes and pseudogenes, as well as for the identification of false positive somatic mutation calls. A detailed overview of the Hapster pipeline is included (**Supplementary note S1, S2**).

**Figure 1.**
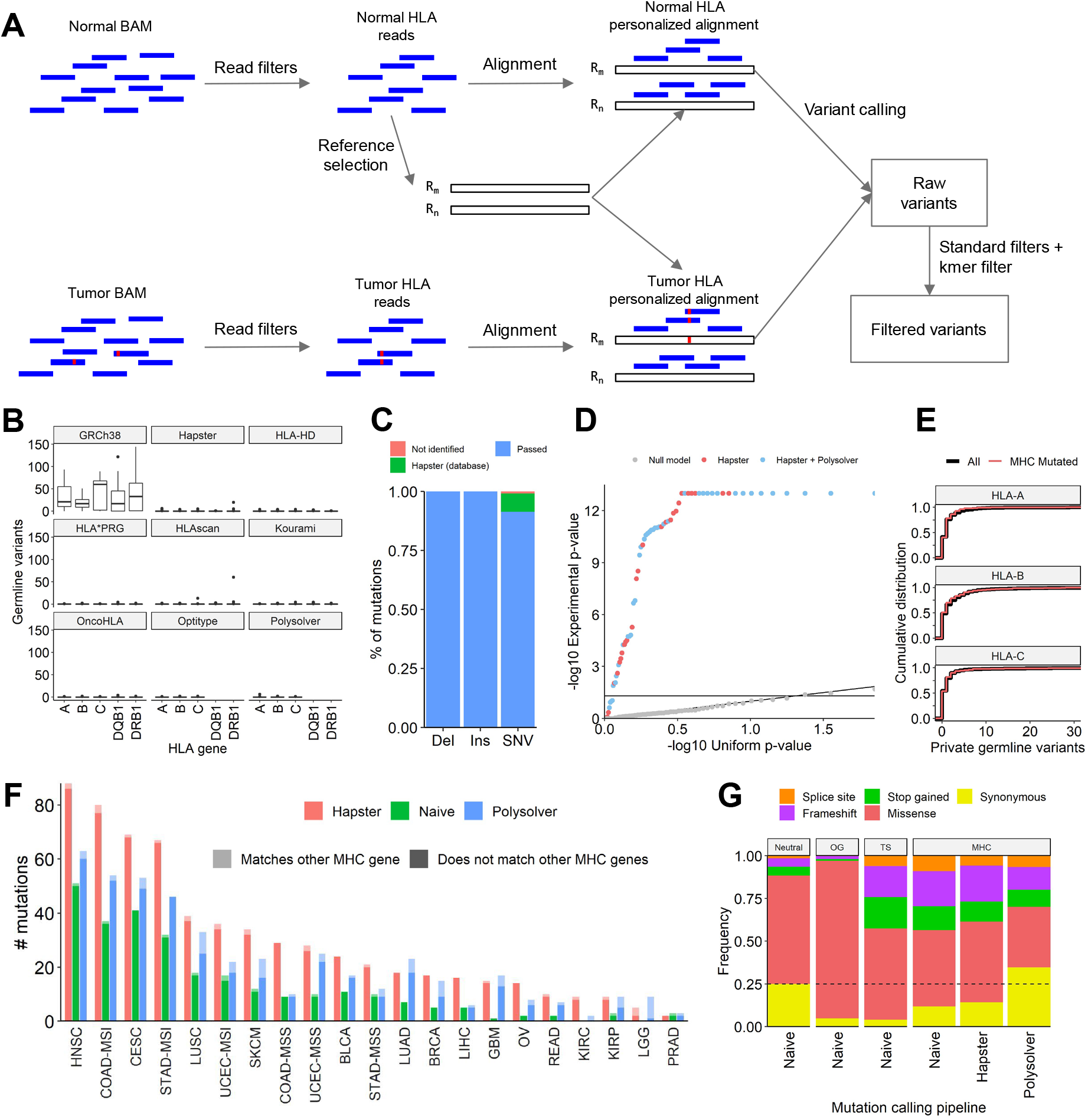
Hapster algorithm and validation. **(A)** Simplified overview of the Hapster algorithm. For each gene, Hapster infers optimal reference sequences R_m_, R_n_ from normal sequencing data, realigns to these personalized references, and calls mutations. For a complete description see **supplementary note S2**. **(B)** Germline variants identified from 69 WES samples from the 1000 Genomes project relative to either the standard reference GRCh38, or to dynamically selected references using 8 different haplotypers. A perfect reference sequence should produce 0 apparent germline variants. **(C)** Fraction of simulated true-positive deletions (Del), insertions (Ins), and SNVs that were either identified and ‘Passed’ all filters, were erroneously filtered by Hapster, or were not identified by the variant caller. **(D)** Probability of read support for observed or randomly generated variants being caused by sequencing error according to a Beta-binomial model. **(E)** Distribution of the number of private germline variants observed per tumor in either all cases or restricted to those cases with MHC mutations. **(F)** Comparison of non-synonymous mutation calls for the MHC class I genes between the naive GDC pipeline, Polysolver, and Hapster across various cancer types from TCGA. Lightly-shaded bars represent possible false positives. **(G)** Comparison of mutational consequences for variants called by alternative approaches in the MHC genes vs oncogenes, tumor suppressors, and neutral genes in TCGA. Oncogenes (OG): KRAS, PIK3CA, IDH1, CTNNB1, FOXA1, BRAF, AKT1, EGFR. Tumor suppressors (TS): TP53, RB1, PTEN, APC, BRCA2, VHL. Neutral genes: All others.

To benchmark the haplotype inference portion of the Hapster pipeline we used a set of 69 WES samples from the 1000 genomes project that have previously reported MHC-I and MHC-II haplotype calls both via sanger sequencing ^37^ and seven *in silico* prediction methods ^34^. MHC haplotyping algorithms are generally ‘digit-optimizing’, in that they attempt to maximize the number of correct digits in the names of the inferred alleles, which due to HLA nomenclature has the effect of emphasizing protein similarity over DNA similarity. However, for the purposes of mutation calling, it is most critical that the reference haplotype minimizes mismatches between an individual’s germline DNA sequence and the chosen haplotypes. We therefore compared Hapster to other haplotyping methods by calling germline variants in WES sequencing data relative to each haplotyper’s inferred haplotype sequence for that individual. We consider the sequencing reads to be the ground truth, and a perfectly identified reference sequence would lead to no germline variants being identified in these reads. We see that relative to the fixed sequences of the standard reference Grch38, there are a median of 17-38 germline mutations observed per gene. All tested haplotypers improve upon this, with each having a median of 0 and a mean of <0.5 observed germline mutation per gene (**Figure 1B**), which is on par with the ~1 variant per kilobase rate observed relative to the standard reference in other non-polymorphic regions ^38^. When Hapster was applied to a larger population of 10,001 normal tissue samples from TCGA, we again found a median of only 0-3 germline variants per allele (**Supplementary table S1**).

To assess mutation calling sensitivity, we first simulated 150 synthetic MHC haplotypes with a random mutation. Of the 150 simulated mutations, 149 were successfully identified **(Figure 1C)**. Following filtering, 9 calls were removed by Hapster’s filters due to the variant randomly matching a known germline polymorphism within the HLA database, giving an overall sensitivity of 93% (140/150). To assess specificity, we called somatic mutations in all 450 samples from the HNSC TCGA cohort with tumor and normal labels swapped, such that no somatic variants should be identified. In 9 cases, an apparent somatic variant was identified that passed all filters. Assuming all 9 calls are false positives gives a specificity of 98% (441/450).

To further assess mutation calling accuracy using an orthogonal sequencing technology, we called somatic variants in the HNSC cohort for validation using RNA-seq. While established RNA-seq validation methods would be ideal, they rely on alignment of reads to a reference in order to identify mutations, which would be inappropriate in validating Hapster. We therefore developed a fully orthogonal alignment-free kmer based approach to determine the read support for each variant in the RNA (**Methods**), avoiding potential reference selection or alignment biases. Of the 75 variants with high enough coverage in the RNA to undergo validation, 70 variants (93%) had either read support significantly exceeding (p < 0.05) the null model of sequencing error (65, 86%) (**Figure 1D**) or were truncating variants with evidence of nonsense mediated decay (5, 7%) **(Supplementary figure S1B**). This leaves only 5 variants (7%) without RNA evidence, a reasonable proportion given the limitations of our statistical model, variable tumor cellularity, loss-of-heterozygosity (LOH), and ubiquitous transcriptional silencing of the MHC locus in tumors^39^. For a second orthogonal validation, we performed Sanger sequencing on tumors from MI-ONCOSEQ ^40^ with sufficient DNA or tissue samples. All 14 candidate variants called by Hapster were clearly detected in the Sanger chromatograms from tumor specimens, while being absent in traces from patient-matched normal tissues **(Supplementary figure S1C**). Addressing the possibility of germline variants being miscalled as somatic due to poor reference selections, we found no evidence of enrichment of somatic mutations in cases with higher numbers of private germline variants **(Figure 1E)**.

Finally, we applied Hapster to a larger set of 7,746 samples from TCGA that have previously reported mutations called by both the NCI Genomic Data Commons (GDC) standard reference based pipeline and the Polysolver personalized pipeline^35,41^. We found that when calling mutations in the MHC class I genes, Hapster detected over twice as many non-synonymous mutations as the GDC pipeline, and 36% more than Polysolver (**Supplementary table S8, Figure 1F, Supplementary figure S1D**). To assess the quality of the mutation calls we first performed an exhaustive search for potential alternative explanations for each variant called by each pipeline. We reasoned that given an accurate haplotype inference, the most likely cause of false positives should be misalignment of sequencing reads originating from other homologous MHC genes or pseudogenes. We found that only 6% of Hapster’s calls matched known sequences in any other MHC gene, a rate significantly lower than that of both the naive GDC pipeline (11%, Fisher’s exact test p < .01) and Polysolver (42%, Fisher’s exact test p < 1e-16) (**Supplementary table S9, Figure 1F, Supplementary figure S1E**). Of note are the highly recurrent synonymous variants p.T214T and p.A269A that are identified as somatic mutations by Polysolver **(Supplementary figure S1F)**. These mutations are unlikely to be under extreme positive selection, but have sequences exactly matching non-classical MHC class I genes *i.e*. are likely due to alignment errors from *HLA-E*, *HLA-F*, or HLA pseudogenes.

Many methods to identify positive selection of mutations within a gene rely on the detection of deviations from a neutral nonsynonymous to synonymous (dN/dS) ratio, and any pipeline biases in functional consequences can confound these analyses. We therefore additionally compared the distribution of functional consequences of HLA mutations called by each of the approaches. For both Hapster and the naive pipeline, synonymous mutation calls were underrepresented when compared to neutral genes, consistent with what would be expected for a potential driver gene **(Figure 1G)**. In contrast, we found that Polysolver had a surprising over-representation of synonymous calls, many of which can likely be attributed to misaligned reads originating from non-classical MHC class 1 genes **(Supplementary table S9, Supplementary figure S1F**).

### 2.2 Pan-cancer compendium of MHC class I and MHC class II mutations

To comprehensively characterize MHC class I and MHC class II mutation rates in human cancer we analyzed 10,001 primary tumors across 35 cancer types from TCGA (**Supplementary table S2**) and 2,199 metastatic and refractory tumors across 24 cancer-types within MI-ONCOSEQ ^40^ (**Supplementary table S3**), for a total compendium of 2069 mutations (**Figure 2A, Supplementary table S4**). Previously only MHC class I mutations in primary tumors have been reported ^35^. Microsatellite unstable (MSI) tumors are immunologically distinct due to their significantly higher neoantigen burden ^42^ and we have therefore separated them from their microsatellite stable (MSS) counterparts within the colon (COAD), stomach (STAD), and endometrial (UCEC) TCGA cohorts. The mutations were in general distributed uniformly across the gene body, but occasionally concentrated within prominent hotspots (**Figure 2A, Supplementary figure S2A**). We found that for the MHC class I HLA-A and HLA-B contained significantly more mutations than HLA-C, and for the MHC class II HLA-DRA contained significantly more mutations than all other MHC class II genes except for HLA-DQB1 (**Figure 2B**). In primary tumors, we noted substantial variation in both mutational frequency and their predicted consequences across tumor types and the MHC gene classes (**Figure 2C, Supplementary data S1**). We found nonsynonymous MHC class I and MHC class II mutations in 10.5% of primary tumors (ranging from 2.7% to 72.5% across cancer types) (**Supplementary figure S2B**), with 5.6% (range 0.2% to 62.3%) of patients harboring an MHC class I and 5.7% (range 1.1% to 21.7%) an MHC class II somatic variant. Consistent with previous reports that MSI tumors should be under strong pressure to acquire loss of MHC function ^43,44^, the COAD-MSI, STAD-MSI, and UCEC-MSI cohorts make up 3 of the top 4 cohorts for MHC class I mutations (**Figure 2D, Figure 2B)** with the majority being loss of function (LOF) frameshifts or stop gains. MHC class II mutations were also most prevalent in cancers with high mutation burden including MSI tumors and melanoma (**Figure 2E**). However, LOF mutations in the top-mutated cohorts were less frequent and the variation in mutation-rates across cancer types was lower compared to MHC class I (**Figure 2E, Supplementary figure S2B, Supplementary data S1**).

**Figure 2.**
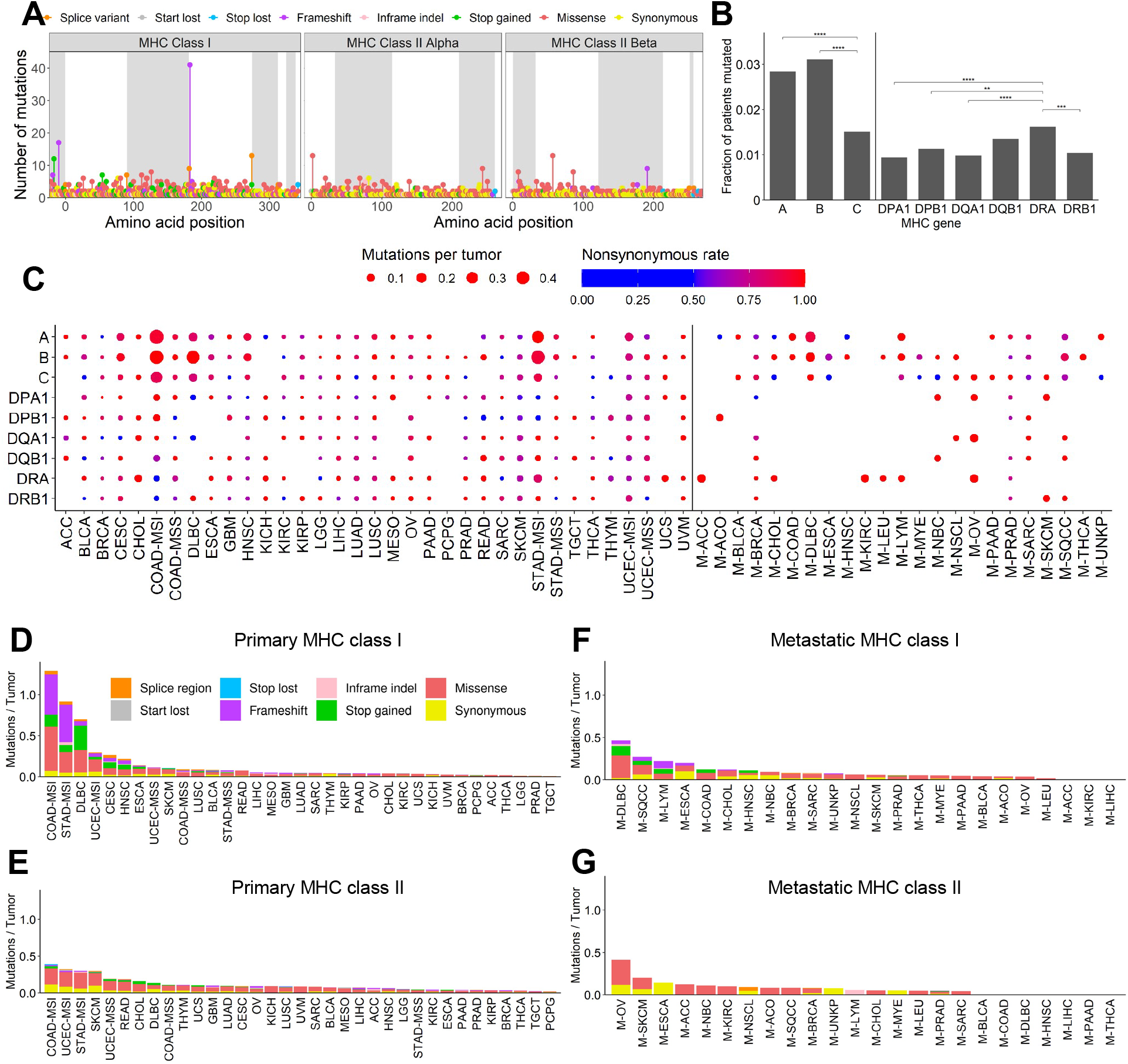
Compendium of MHC class I and class II mutations in primary and metastatic tumors. **(A)** Distribution of all observed mutations in both primary and metastatic cancers across the coding region of the MHC genes. **(B)** Significant differences in the prevalence of nonsynonymous mutations and indels of individual MHC class I and MHC class II genes. **(C)** Cohort specific mutation rates for MHC class I and MHC class II genes across all primary and metastatic cancers. Values are scaled to the number of individuals within each cohort. Colors represent the fraction of cancers with nonsynonymous/indel mutations. At neutrality, the expected nonsynonymous rate should be approximately 0.75. **(D,E,F,G)** Cohort summaries of coding region mutations in MHC class I **(D,F)** and MHC class II **(E,G)** genes in primary **(D,E)** and metastatic **(F,G)** cancers. Values are scaled by the number of individuals within each cohort.

### 2.3 Prevalence of MHC class I and MHC class II mutations in primary vs metastatic tumors

The prevalence of MHC mutations in metastatic tumors is unknown, a critical gap in knowledge considering the predominant use of immunotherapies in this setting and immunological differences between the primary and metastatic tumor micro-environment (TME) ^40,45–48^. Overall, we observed nonsynonymous MHC class I and MHC class II mutations in 7.6% (range 3.3%-20.0%) of metastatic patients, with substantial variation in mutational frequency and functional consequences between cancer types (**Figure 2F-G, Supplementary figure S2C, Supplementary data S1**). To directly compare mutation rates between primary and metastatic cancers we created a set of pairings to match TCGA cohorts to MI-ONCOSEQ cohorts (**Supplementary table S5**). For 14/17 pairings (82%) there were no significant changes in primary vs metastatic MHC class I or MHC class II mutation rates, however select cohorts did show a significant difference (**Supplementary figure S2D**). For prostate and breast cancers we observed a significant increase in MHC class I mutations in metastatic cancers compared to primary (prostate: F(1, 909) = 9.35, p = 0.002; breast: F(1, 1140) = 12.8, p = 0.0004). Additionally, in breast cancers we see a slight increase in MHC class II mutations in metastatic cancers (F(1, 1033) = 5.81, p = 0.016). In ovarian cancers we see elevated MHC class II mutations (**Figure 2G**) and a nearly 3-fold increase in metastatic cancers compared to primary (F(1, 260) = 3.86, p = 0.05, **Supplementary figure S2D**). While both primary and refractory DLBC cohorts showed a high MHC class I mutation rate relative to other cancer types, there was a trend towards decreased MHC class 1 mutations in refractory cases, although this did not reach significance (F(1, 82) = 2.94, p = 0.09).

### 2.4 Impact of tumor mutation burden on MHC class I and MHC class II mutation frequency

To investigate the association between tumor mutational burden (TMB) and MHC mutations we compared the local TMB within the MHC genes to the genome-wide TMB for each cancer cohort. As global TMB increases, neoantigen burden increases, and we would expect increased selective pressures for LOF MHC mutations. This trend is evident for MHC class I mutations in primary cancers where increasing global TMB is strongly associated with increasing local TMB (p < 1e-11, Spearman’s ρ = 0.83, **Supplementary figure S2E**) with some high TMB tumors reaching a striking local TMB of >100 mutations/Mb. However, this association is weaker for the MHC class II genes (p < 0.001, Spearman’s ρ = 0.54) with lower TMB cancer types frequently observed having a local MHC class II TMB on par with high TMB cancer types. While the associations are generally weaker in metastatic and refractory cancers, we nonetheless see the same trend with local TMB showing a much stronger association with global TMB in the MHC class I genes (p = 0.05, Spearman’s ρ = 0.40) than in the MHC class II genes (p = 0.57, Spearman’s ρ = 0.12) (**Supplementary figure S2F**). We originally hypothesized that somatic loss of MHC class II should mirror that of MHC class I given that both have been shown to promote anti-tumor immune responses. However, there was no association at the cohort level between MHC class I mutations and MHC class II mutations after controlling for TMB **(Supplementary figure S2G-I)**. Strikingly, while MHC class I mutations appeared to be most prevalent in cancer types with high TMB, MHC class II mutations were frequently increased in low TMB cancers with few MHC class I mutations. Importantly, predicted deleterious MHC class I mutations are significantly enriched in cancer types with approved immune-checkpoint inhibitors (OR: 2.29, p < 1e-16) (**Supplementary table S6**).

Overall these data provide, to our knowledge, the first comprehensive look at MHC class I and MHC class II mutations pan-cancer, across both primary and metastatic tumors. We find that somatic mutations of HLA-A and HLA-B are most common while HLA-C and MHC class II genes are less frequently mutated and are likely only relevant in specific cancer types. In addition, TMB, and therefore neoantigen load ^49^, is significantly associated with MHC class I mutations to a greater extent than with MHC class II mutations. While some significant differences in MHC mutation rate between primary and metastatic tumors are noted, the majority of MHC mutations in metastatic tumors is expected to be already present in the primary tumor.

### 2.5 Positive selection of non-synonymous MHC somatic mutations

Given the high proportion of deleterious mutations in cancer types with the highest frequency of MHC mutations, we asked whether there was significant evidence for positive selection of functional mutations within the MHC genes. We applied CBaSE ^50^, a tool that estimates the gene-specific strength of positive or negative selection for functional mutations, to each primary and metastatic cohort from TCGA and MI-ONCOSEQ, respectively. HLA genes and haplotypes are codominant and each allele presents a largely unique set of neoantigens ^51^. In addition, specific T cell responses are often immunodominant and mounted against only a few of the presented neoantigens. Mutation of a single HLA allele may therefore result in the complete inability to present an immunodominant neoantigen. Accordingly, in the following analyses we treat all MHC class I genes (and separately, all MHC class II genes) as one functional unit, analogous to multiple genes of a protein complex ^52^, taking into account the increased genomic length of this combined set of genes. In primary cancers, CBaSE identified 6 cohorts (COAD-MSI, STAD-MSI, DLBC, CESC, HNSC, LUSC) with statistically significant evidence for positive selection of non-synonymous variants in the MHC class I genes, and 3 cohorts (CHOL, KICH, UVM) for the MHC class II genes (**Figure 3A**). By this measure, the MHC class I are tied for 7th and the MHC class II are tied for the 17th most recurrent driver genes pan-cancer as determined by applying CBaSE to all protein-coding genes across primary cancers. A similar trend was identified in metastatic and refractory cancers with the MHC class I genes being mutated in two cohorts (M-DLBC, M-LYM) making them tied for 6th most recurrent pan-cancer driver gene by number of cohorts significantly mutated (**Figure 3B**). As an alternative measure of positive selection, we used Fisher’s method to create a combined score Φ_pos_ for the strength of selection across all cohorts (**Figures 3C-D, Supplementary figure S3)**. We found that in both primary and metastatic cancers the MHC class I genes scored in the top 0.1% of all protein-coding genes according to this metric of positive selection **(Figures 3C-D)**, and in primary cancers the MHC class II genes scored in the top 0.5% (**Figure 3C**). Due to the exclusion of MHC class II genes from the sequencing panel in a subset of MI-ONCOSEQ samples, we were not statistically powered to investigate selection of MHC class II genes in metastatic cohorts.

**Figure 3.**
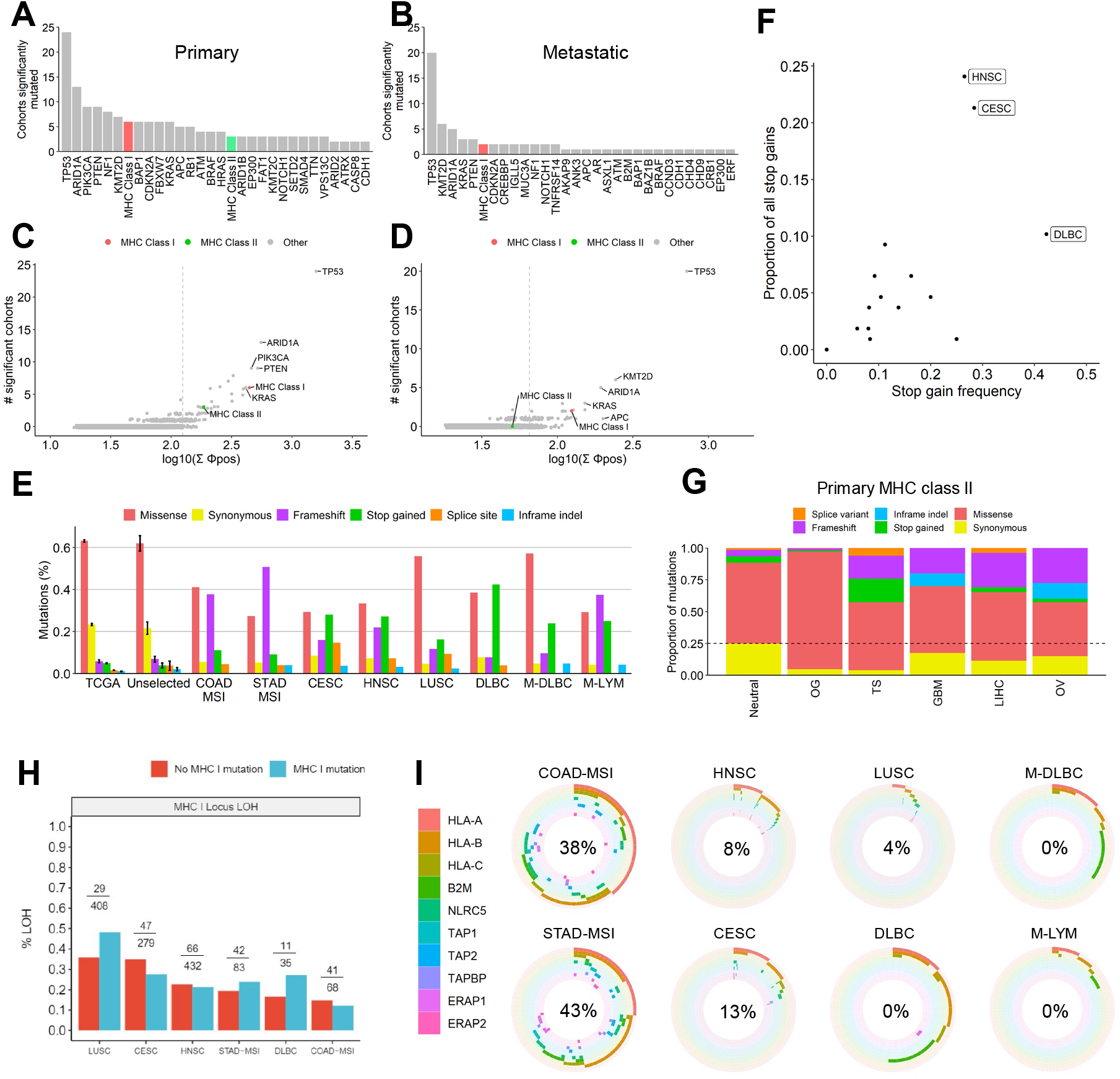
Evidence for strong positive selection and deleteriousness of MHC somatic mutations. **(A, B)** Top 30 genes showing evidence of positive selection in primary **(A)** or metastatic **(B)** cancers by CBaSE by number of cohorts with significant evidence. **(C, D)** Comparison of the number of cohorts significantly mutated vs pan-cancer metastatic Φ_pos_ for protein-coding genes in primary **(C)** or metastatic **(D)** cancers as measured by CBaSE. Vertical dashed lines show the cutoff for the top 0.5% of genes by Φ_pos_. **(E)** Proportion of functional consequences observed in various groups: “TCGA” 2,600,654 pan-cancer mutations from TCGA an approx. neutral model; “Unselected” - MHC class I mutations from all primary and metastatic cohorts showing no evidence of positive selection; others - MHC class I mutations from cohorts showing evidence of positive selection (n = 21-96 mutations within positively selected cohorts). “TCGA” and “Unselected” are average frequencies across cohorts, with error bars showing SEM. **(F)** Cohort specific MHC stop gain frequency compared to overall proportion of stop gains contributed by each primary cohort. **(G)** Functional consequences of MHC class II mutations in select primary cohorts. For comparison, mutational consequence distribution of known oncogenes (OG: KRAS, PIK3CA, IDH1, CTNNB1, FOXA1, BRAF, AKT1, EGFR) and tumor suppressors (TS: TP53, RB1, PTEN, APC, BRCA2, VHL) are shown. **(H)** Association of HLA mutations and LOH, numbers indicate portion of mutated tumors (mutated | total) **(I)** Sample level co-occurrence of mutations in either the MHC class I or APM genes within positively selected cohorts. Percentage values show percent of mutated samples containing a hit in both the MHC class I and APM, with lower percentages suggesting mutual exclusivity.

### 2.6 Functional consequences of MHC class I and MHC class II mutations

To better understand the imprint of positive selection of non-synonymous mutations in MHC genes, we characterized their functional consequences and compared their distributions in cohorts with and without evidence of positive selection. We constructed an approximately neutral model by looking at the distribution of functional consequences across 2.6 million mutations called from the entirety of the TCGA, the overwhelming majority of which are known to be passengers ^53^ (**Figure 3E**, "TCGA”). MHC class I mutations within cohorts showing no evidence of positive selection showed a consequence distribution nearly identical to that of the neutral model (**Figure 3E**, "Unselected”), supporting the notion that mutations observed in these cohorts are primarily passengers. However, in the six primary and 2 refractory cancer types that did show positive selection, there was a clear deviation from the neutral model (p < 1e-3 - 1e-16). Consistent with the MHC’s role as a tumor suppressor, this deviation was caused by an increase in truncating mutations which accounted for more than 40% of mutations in most cohorts, as compared to the expected neutral rate of ~12%. The B-cell lymphoma (DLBC), cervical (CESC), and head and neck (HNSC) cohorts all have a high proportion of stop gains (46%, 32%, and 28%, respectively) within the MHC class I genes, accounting for 56% (60/108) of all observed stop gains despite only comprising 8% (792/10,001) of TCGA patients (**Figure 3F**). Notably, frameshift mutations in MHC class II were rare even in MSI tumors, but unexpectedly common in some MSS tumors including GBM, OV, and LIHC. These cohorts were also depleted of synonymous mutations (**Figure 3G**).

Similar to the DLBC cohort from TCGA, the refractory M-DLBC cohort showed both a high mutation rate and a strong bias towards truncating mutations in the MHC class I genes (35%). Other non-DLBC refractory lymphomas (M-LYM) had a lower overall MHC class I mutation rate, but still had a large bias towards truncating mutations (65%) (**Figure 3E, Supplementary figure S4A**). The lymphomas alone account for 52% of stop gains observed across all MI-ONCOSEQ cohorts (11/21) despite containing only 7% of patients. The HNSC, CESC, and LUSC cohorts in TCGA are all types of squamous cell carcinomas which correspond to a single cohort M-SQCC within MI-ONCOSEQ. Similar to what was observed across the primary squamous cancers, the pan-squamous M-SQCC cohort showed an overall elevated mutation rate and a high rate of LOF mutations when considering frameshifts, stop gains, and splice region variants (35%, **Supplementary figure S4B**). Metastatic MSI tumors are underrepresented in MI-ONCOSEQ preventing any comparison to primary MSI. Altogether, these data reveal striking differences in mutation frequency and deleteriousness not only across cancer types but also between MHC class I and class II genes.

### 2.7 Patterns of mutual exclusivity and independence of MHC mutations

We sought to determine whether mutations in the MHC are independent of other mutational drivers, and concomitant allelic losses. We focused on the cohorts with evidence of positive selection (**Figure 3E**), and examined if MHC class I mutational status was associated with HLA loss-of-heterozygosity (LOH) (**Supplementary figure S4C, Methods**). As predicted from the high heterozygosity and co-dominance of HLA alleles, and immunodominance of T cell responses (see above), we did not find statistically significant evidence for bi-allelic inactivation (mutation compounded by LOH) of MHC class I in primary tumors (**Figure 3H**). Since the ability to present antigens can be restricted by mutations of other genes that make up the antigen processing machinery (APM), we next looked at the relationships between deleterious mutations in the MHC class I genes and the APM ^54^ (**Figure 3I**). Other genes linked to the MHC class 1 have been identified as cancer driver genes (e.g. *B2M ^55^*), and it has been shown that driver genes that fall within the same pathway frequently show mutual exclusivity. This effect is most clear in the lymphomas where there is significant mutual exclusivity between MHC mutations and the APM (p < 0.01) with no observed tumors having hits in both gene sets. However, there is no mutual exclusivity in the squamous cell carcinomas (p = 0.99) with the LUSC, HNSC, and CESC cohorts having 4%, 8%, and 13%, respectively, of mutated tumors with simultaneous hits in the MHC genes and the APM. MSI tumors had even higher overlap with 38% of COAD-MSI and 43% of STAD-MSI tumors containing simultaneous hits, which does not support mutual exclusivity (p = 0.81). These data suggest that allelic loss of HLA does not significantly reduce (or increase) the pressure to select for additional mutations limiting antigen presentation in solid tumors, but appears dominant in lymphomas. However, it is also possible that low mutual exclusivity is the result of high tumor heterogeneity, with multiple subclones having independent loss of APM function that only appear to co-occur due to bulk sequencing. Finally, we looked at potential co-mutation of the MHC class I genes with known driver genes (**Supplementary figure S4D**). The strongest observed co-occurrence was found in the HNSC cohort with deleterious mutations of *CASP8*, a gene that plays a key role in an alternative pathway for destruction of malignant cells by the immune system ^56^. This observation has been confirmed in an independent cohort of primary HNSCC (**Supplementary figure S4E**). Also observed in the HNSC cohort was co-mutation with HRAS, a member of the RAS family of oncogenes that has been shown to be associated with increased immune activity within head and neck cancers ^57^ **(Supplementary figure S4D**.

### 2.8 Mutational processes shape cancer type specific MHC mutational patterns

We next sought to determine which mutational processes may contribute to the generation of the non-synonymous mutations within cohorts showing evidence of positive selection. For MSI cancers, mismatch repair deficiency (MMRd) is the primary mutational process leading to a large number of indels within microsatellites ^58^. We have already observed a high rate of frameshift indels within MSI tumors (**Figure 3E**) and notable hotspots (**Figure 2A, Supplementary figure S2A**), and upon further investigation the majority of these frameshifts (57/63, 90%) are due to single base pair insertions or deletions at homopolymer microsatellites (**Figure 4A**) which occur at a rate much higher than observed in MSS cancers (p < 1e-16) (**Figure 4B**). We also observed that MSI-associated indels were preferentially in longer homopolymers, while MSS-associated indels showed no relationship with homopolymer length (p < 0.001) (**Figure 4C**), which is consistent with the mutational signature for MMRd ^59^. For the lymphoma and squamous cell carcinoma cohorts, we observe a striking number of stop gain mutations, including multiple recurrent (n > 2) hotspot positions (**Figure 4D**). Interestingly, 100% of the stop gains that we observed in these cohorts are caused by C>T or C>A mutations. This is consistent with the well characterized process of cytosine deamination which is frequently observed in lymphomas due to activation-induced cytosine deaminase (AID) ^60^ and in squamous cell carcinomas due to the closely related APOBEC family of enzymes ^61^. Both AID and APOBEC have distinct sequence preferences for their deaminase activity that should be visible in the sequence motifs surrounding each mutation. In the lymphomas we find that 13/22 (59%) of the observed stop gains match the canonical AID motif WR<u>C</u> (W = A/T, R = A/G) (**Figure 4D-F**). Similarly, across the squamous cell carcinoma cohorts we find that 49/62 (79%) of observed stop gains match either the APOBEC3A/B/H/F motif T<u>C</u> or the APOBEC3G motif CC<u>C</u> (**Figure 4D,G,H**). Further analysis showed that mutational signatures SBS2 and SBS13, which have been reported to be associated with APOBEC activity ^62^, are significantly more active in squamous cell carcinomas with observed stop gain mutations (**Figure 4I**).

**Figure 4.**
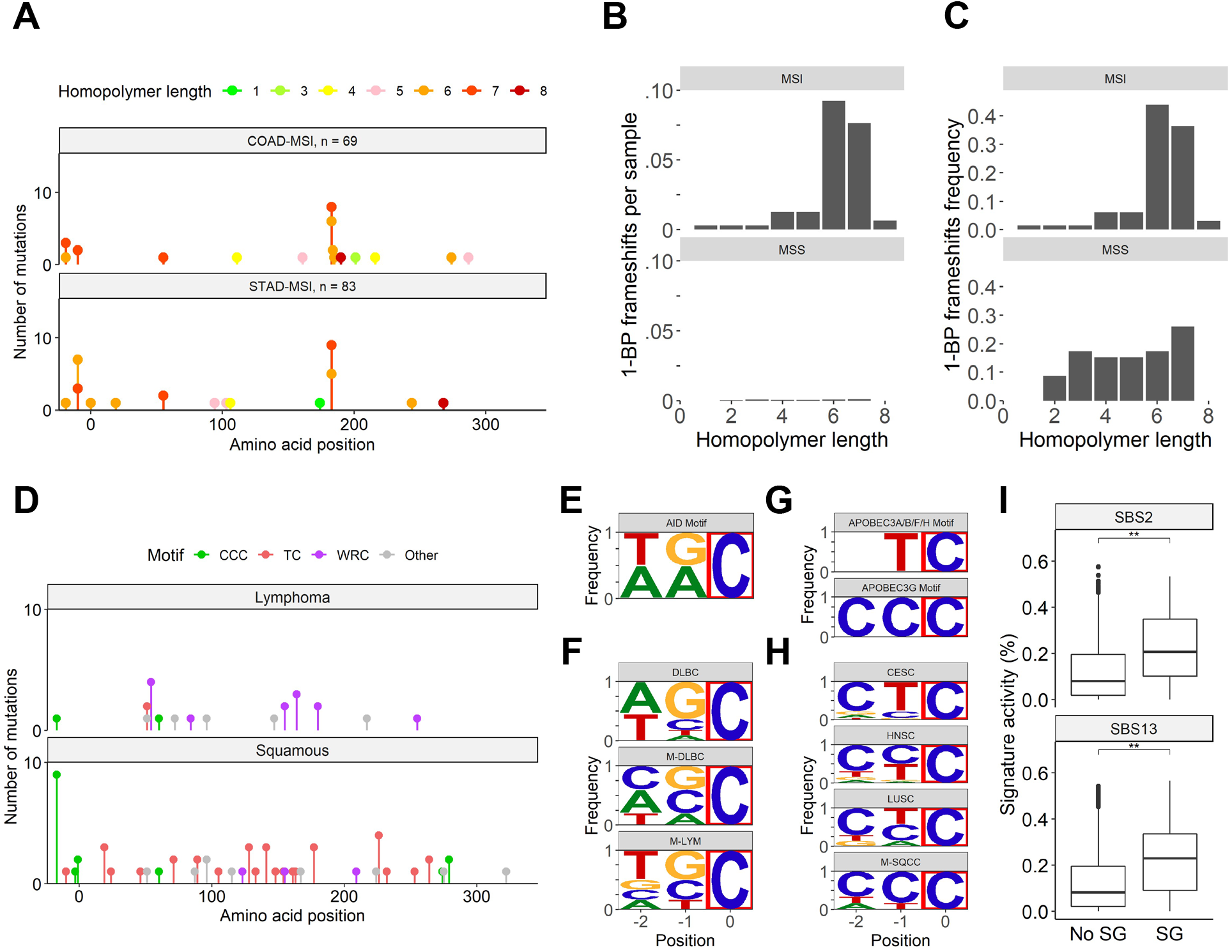
Distinct mutational processes shape cancer-type specific MHC mutational patterns. **(A)** Position of observed frameshift mutations within the MHC class I genes of COAD-MSI and STAD-MSI tumors. Colors show length of homopolymer microsatellites at each observed frameshift. **(B)** Total number of 1-BP frameshifts observed per tumor at homopolymers of varying length in MSI and MSS tumors. **(C)** Frequency of 1-BP frameshifts observed at homopolymers of varying length in MSI and MSS tumors. **(D)** Position of observed stop gain mutations within the MHC class I genes of lymphomas and squamous cell carcinomas. Colors show motif of mutated position. **(E)** Canonical motif for AID **(F)** DNA motifs for stop gain mutations observed in lymphoma cohorts. Mutated base marked with red box**. (G)** Canonical motifs for APOBEC proteins. **(H)** DNA motifs for stop gain mutations observed in squamous cell carcinoma cohorts. Mutated base marked with red box. **(I)**% of mutations within Squamous cell carcinomas with (SG) or without (No SG) observed stop gains that can be attributed to signatures SBS2 and SBS13, which have been associated with APOBEC activity. **: p < 0.01

Altogether, these observations strongly suggest that truncating mutations within the MHC genes originate due to specific mutational processes active within select cancer types. The active mutational processes are responsible not only for producing highly immunogenic tumors that are under pressure to select truncating mutations within the MHC class I genes, but are also directly responsible for creating the majority of the LOF mutations in the first place. Haplotypes harboring homopolymer repeats and AID/APOBEC templates are therefore potentially more susceptible to this immune-escape mechanism.

### 2.9 Missense mutations are enriched in specific MHC functional domains

While frameshift and stop gain mutations are easy to classify as LOF, missense mutations are more difficult to interpret as they can be LOF, neutral, gain of function, or even neomorphic ^63,64^. We hypothesized that deleterious missense mutations within positively selected cohorts should accumulate predominantly within the functional domains that provide the most immune escape potential for a tumor. To detect this enrichment we constructed two null models (**Figure 5A, Methods**), which we compared to the observed mutations across MHC functional domains which we established through systematic expert-knowledge and crystal structure guided annotation of individual amino acids within the MHC class I proteins (**Figure 5B, Supplementary figure S5A-C, Supplementary table S7**). The first ‘simulated’ null model was based on a large number of HLA mutations generated randomly taking into account HLA sequence trinucleotide contexts and observed mutational signature activities within each positively selected cohort **(Methods)**. The second ‘observed’ null was based on 251 actual mutations called in cohorts that showed no evidence of positive selection. We first compared the observed null dN/dS ratios to the simulated null ratios across all functional domains (**Figure 5C**). The log fold change of the observed *vs* simulated local dN/dS ratios were normally distributed (Shapiro-Wilk p = 0.99) with a mean not significantly different than 0 (p = 0.38), showing appropriately that there are no significant differences between the two null models. In contrast, in cohorts showing evidence of positive selection, the local dN/dS fold change was also normally distributed (Shapiro-Wilk p = 0.98, p = .433) but with a mean significantly above 0 (p = 0.02, p = 0.006) when compared to both the simulated null and the observed null, respectively (**Figure 5C**). This shows a strong general trend towards excess nonsynonymous mutations across all annotation regions.

**Figure 5.**
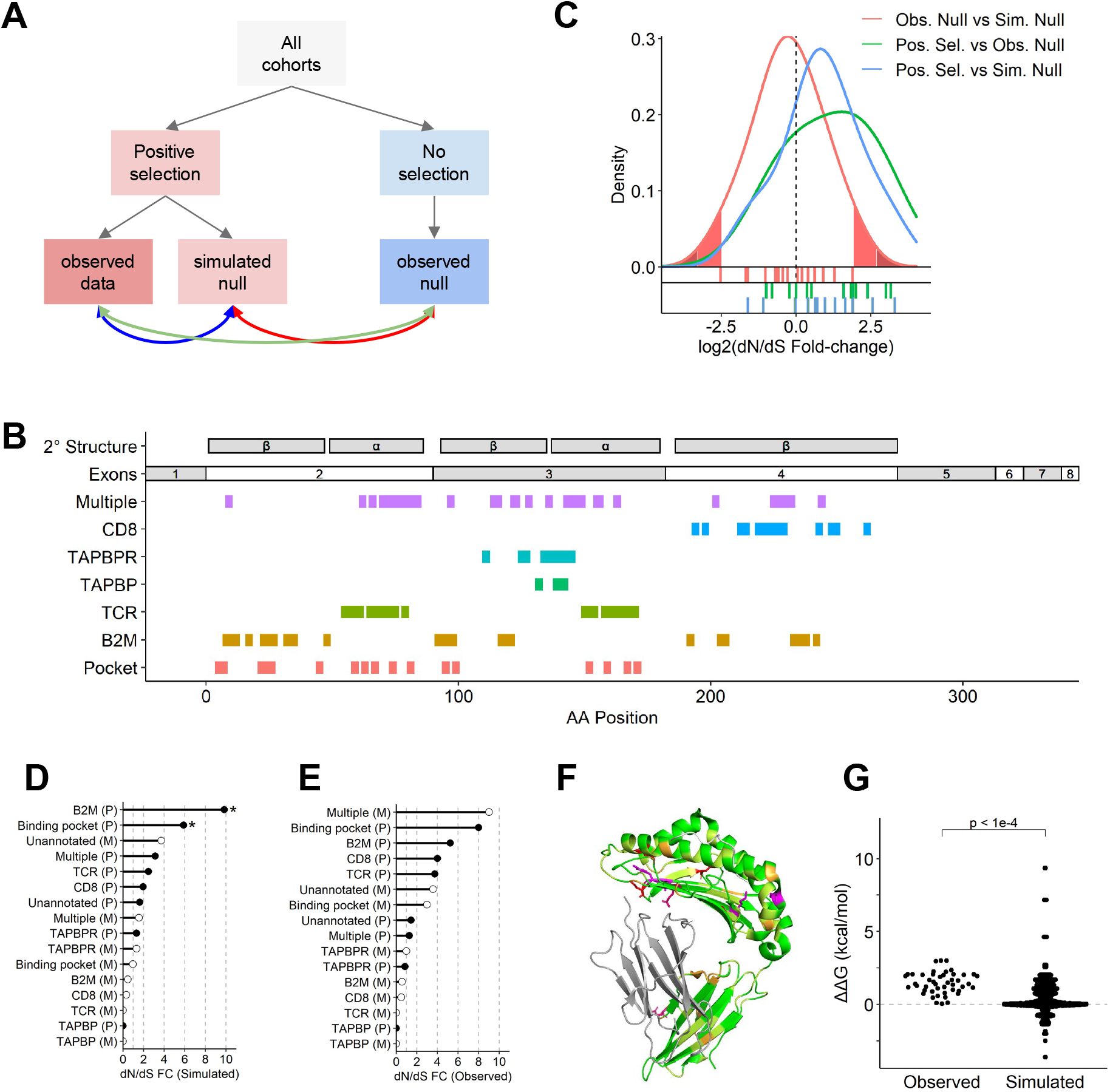
MHC class I missense mutations are enriched within specific protein functional domains. **(A)** Schematic representation of the constructed null models and their comparisons (arrows). **(B)** Schematic overview of HLA proteins showing secondary structure, exon boundaries, and amino acid interactions with various binding partners. **(C)** Distribution of observed vs simulated dN/dS ratio fold-change for amino acids that are predicted to interact with various MHC interacting partners. Observed dN/dS ratios are compared to dN/dS ratios from simulations taking into account the mutational signature activities within each of the cohorts showing evidence of positive selection. Rug plots show individual data points. Filled regions show tails of empirical null distribution. (light-red: top and bottom 5%; dark-red: top and bottom 1%) **(D-E)** dN/dS ratio fold-changes vs simulated / theoretical **(D)** and observed / empirical **(E)** null models for individual annotation regions within cohorts showing evidence of positive selection (P: primary cancers; M: Metastatic/refractory cancers). Stars denote observations above the 95th percentile based on the observed null distribution. **(F)** Structure of the MHC/B2M complex showing 3D clustering of recurrently mutated amino acids. Positions within the MHC protein are colored based on the number of observed mutations (0: green, 1: yellow, 2: orange, 3: pink, 4: magenta, 5: red). Positions mutated 3 or more times are shown with side chains visible. B2M and bound peptide in grey. **(G)** Change in binding energy due to mutations in amino acids at the HLA:B2M interface as predicted by the software SSIPe.

Analysis of missense mutations in the context of functional domains acts not only as a signal for positive selection, but also helps in the interpretation of the mutations’ likely functional consequences. Therefore, using our amino acid annotations, we examined which functional domains had the highest enrichment of non-synonymous mutations in both primary and metastatic tumors (**Figure 5D-E**). Compared to both the null models, 5 of the 7 annotated functional domains showed a two-fold or higher enrichment of non-synonymous mutations. Some differences were noted between primary and metastatic tumors. Specifically, based on the ‘simulated’ null model, multi-functional residues are under a stronger positive selection in metastatic compared to primary tumors (**Figure 5D-E**), while on the other hand residues involved specifically in the B2M, TCR and CD8 binding interfaces show strong enrichment only in the primary tumors (**Figure 5D-E**).

### 2.10 Mutations at the B2M interface are predicted to disrupt MHC-B2M complex formation

In primary tumors, particularly striking are the B2M interacting and binding pocket residues displaying dN/dS ratios of 5 to 10 fold with respect to both null models **(Figure 5B,D,E)**. This suggests that within cohorts showing evidence of positive selection, missense mutations may be LOF by disrupting the ability of the mature MHC proteins to interact with either B2M or their cognate neoantigen peptides. To examine this, we overlaid all missense mutations from the positively selected cohorts on the crystal-structure of the MHC class 1 - B2M complex^65^ (**Figure 5F**), which revealed a clustering of recurrently mutated positions in 3D space at both the interface between the MHC and B2M proteins and at the anchor points of the peptide binding pocket (**Figure 5F**), strongly suggesting that mutations disrupt this interface leading to loss of MHC function. This is consistent with previous studies identifying B2M itself as a driver gene in all cancer types implicated here ^14,50,66^. To determine whether missense mutations at the MHC class 1 - B2M interface are potentially deleterious we used SSIPe ^67^ to predict the change in binding energy resulting from each observed mutation in comparison to that of our previously ‘simulated’ null mutations (**Figure 5G**). Observed mutations had a significantly higher predicted ΔΔG than appropriately simulated mutations (median ΔΔG 1.37 vs 0.15, p < 1e-4, **Supplementary figure S5D**). Additionally, 42% of observed mutations had a ΔΔG > 1.5 kcal, the threshold suggested by SSIPe as evidence for significant disruption of a protein-protein interface. Overall, this suggests that the observed somatic mutations in residues at the MHC-B2M interface are more disruptive than expected by chance. Interestingly, even though *B2M* itself was enriched for loss of function mutations **(Supplementary figure S5E,F)**, there were no observed missense mutations in *B2M* in residues at the MHC class 1 interface.

Altogether, these findings demonstrate that observed missense MHC class I mutations are strongly enriched at residues enabling MHC antigen binding complex formation, and have a likely deleterious function.

## 3. Discussion

Despite overwhelming evidence from IHC studies demonstrating frequent loss of MHC expression pan-cancer ^39^, there is still much that is unknown about the molecular mechanisms that drive or underlie it. To address this problem, we have developed Hapster, a novel algorithm to detect germline and somatic mutations in the MHC locus with high sensitivity and specificity. Using Hapster we identified the cancer types most affected by somatic mutation of the MHC and the mutational processes that promote and generate these mutations. We quantified positive selection and patterns of mutual exclusivity and independence of MHC mutations. We also characterized the deleterious consequences of truncating and missense mutations at the level of functional residues and protein domains.

We have identified six cancer types (colon and stomach adenocarcinomas with microsatellite instability; head and neck, cervical, and lung squamous cell carcinomas; lymphomas) that are significantly enriched for somatic non-synonymous mutations of the MHC class I. Notably, all of these cancers display above-average levels of tumor-immune infiltration ^68^. At the pan-cancer level MHC mutant tumors are significantly more likely to have approved immunotherapies (p < 2.2e-16, OR: 2.29) (**Supplementary table S6**). A logical interpretation of these results is that at the cohort-level immunologically ‘hot’ tumors tend to both respond to ICI and mutate the MHC as an immune-escape mechanism. However, individual patients with impaired MHC function may be partially or completely invisible to T cells ^39^, making them less likely to respond to ICI. Since somatic loss of one or more MHC genes is essentially irreversible ^69^ it may preclude T cell based immunotherapies as viable treatments of MHC-mutated tumors. Within the above 6 cancer types, nearly 10% of patients harbor a functional mutation within the MHC class I genes, highlighting the necessity of further studies of MHC function in the context of both primary and acquired resistance to ICI.

We provide a first look at MHC mutations in metastatic and refractory cancers obtained using personalized genomics. Metastatic cancers typically have a higher TMB, and therefore neoantigen load, and may be expected to be more immunogenic than primary tumors. However, metastases may also originate from less immunogenic sub-clones in the first place, and seed locations that are immunosuppressive such as bone marrow ^70^ or liver ^71^, making them less visible targets for the immune system. IHC studies have also observed both MHC^+^ primary cancers that produce MHC^−^ metastases, and MHC^−^ primary tumors that produce metastases that regain MHC function ^72^. It is therefore unclear how pan-cancer mutational patterns in the MHC locus should compare to primary tumors. We show here that, broadly, MHC mutations in metastatic cancers mirror that of primary tumors. In refractory lymphomas there is a downward trend in MHC class I mutations when compared to primary, however this trend did not reach significance and evidence for positive selection still remains very high. Similarly, in the metastatic squamous cell carcinomas we see that there is no longer evidence for positive selection of MHC class I mutations as was observed in the primary cases. However, in contrast, we note a significant increase in MHC class I mutations in metastatic breast and prostate cancers, as well as MHC class II mutations in metastatic breast and ovarian cancers. Where statistically significant differences are observed, we note that effect sizes are relatively small, so the results should be interpreted with caution.

We also provide novel insights into MHC class II mutations pan-cancer. While MHC class II function is not strictly required for identification of neoantigens by CD8^+^ T cells, it has been shown to regulate anti-tumor T cell responses ^20^ and can act as a therapeutic target for CD4^+^ T cell based cancer vaccines ^25,73^. While select cohorts (CHOL, KICH, UVM) did show evidence for positive selection of functional MHC class II mutations, there were overall fewer mutations in the MHC class II genes than in the MHC class I. This is expected as MHC class II expression is not constitutive and must first be induced before a tumor can be under pressure to select for a LOF mutation, which is in contrast to the MHC class I where selective pressures are present from the onset of tumor formation. What was not expected, however, was the lack of a relationship between cancer types that lose MHC class I and those that lose MHC class II function given that both sets of proteins act to present neo-antigens to T cells to promote an anti-tumor response. We observed no overlap in cohorts showing evidence of positive selection for LOF mutations in the MHC class I and MHC class II genes, with MHC class I loss being more prevalent in high TMB cancer types and MHC class II loss being more prevalent in low TMB cancer types. It will take future studies to determine if this is driven by differences in MHC class II induction in different tissue types, differences in APC or CD4^+^ T cell infiltration in the TME of different cancers, or if this is an actual relationship between low TMB tumors and MHC class II loss.

We have investigated the role of TMB overall as well as specific mutational processes in MHC mutagenesis. We identified three mutational signatures associated with mismatch repair deficiency, APOBEC activity, and AID activity as major drivers of MHC mutations in specific cancer types. We posit that for any given cancer under pressure to lose MHC function, it will be lost via “the path of least resistance” given the processes active in each different cancer type. In MSI cancers, squamous cell carcinomas, and lymphomas, the path of least resistance may be specific to the mutational processes active in these cancers. In contrast, there is melanoma which is known to have high TMB and to respond to immunotherapy, but is not observed to have many MHC mutations. However, transcriptional downregulation of MHC class I, as well as *B2M* mutations, are known acquired resistance mechanisms in this cancer type. Other cancers, such as prostate adenocarcinoma, are shown in IHC studies to have extremely high rates of loss of MHC expression ^39^, yet show no apparent somatic mutations in the MHC genes.

Altogether, our study demonstrates the high prevalence, positive selection, and deleterious nature of MHC mutations, and suggests that immune-escape through MHC mutagenesis is a common and early step in the progression of several common cancer types. While this study primarily focused on immunotherapy naive tumors, future studies looking at tumors post-immunotherapy may reveal MHC mutations to be drivers of acquired immunotherapy resistance. In this work we have focused on MHC mutations, however, in other cancer types, MHC function may be more easily lost via structural loss of the MHC locus, LOH ^16^, transcriptional repression, or even post-translational inactivation. Given Hapster’s ability to create personalized haplotype references, we aim in future studies to apply Hapster to other sequence based analytical methods to identify patterns of loss at the RNA and protein level. This will provide a deeper understanding of the diversity of mechanisms that can lead to loss of MHC expression, and as a result immune evasion and immunotherapy resistance.

## Methods

### Data availability

WES data for the 29 HapMap samples with previously reported HLA haplotypes ^30^ were obtained from the NCBI Sequence Read Archive (https://www.ncbi.nlm.nih.gov/sra). WES and RNA-seq data for all TCGA samples (**Supplementary table S2**) were obtained from the NIH Genomic Data Commons portal (https://portal.gdc.cancer.gov). WES and targeted sequencing data from MI-ONCOSEQ were obtained from ^40,74^, with raw sequencing data supporting the HLA mutation calls made available through Zenodo (ID). HLA reference sequences were downloaded from the IMGT/HLA public database release (https://www.ebi.ac.uk/ipd/imgt/hla). HLA mutation calls are provided in the supplement (**Supplementary tables S8-S9**). Crystal structures used to identify HLA residues involved in protein:protein interactions were obtained from the RCSB protein data bank (https://www.rcsb.org).

### Code availability

Source code for the Hapster tool is available online (https://github.com/mctp/hapster).

### Reference Selection Validation

Quantitative reference selection validation was performed using a ground truth set of 69 WES samples from the 1000 Genomes project with previously reported HLA haplotype calls ^34^. For each sample and each method, an HLA reference was constructed by taking the genomic sequence in the IMGT/HLA database (https://www.ebi.ac.uk/ipd/imgt/hla) corresponding to the called HLA type. For a comparison to GRCh38, the standard reference haplotype (A*03:01:01:01, B*07:02:01:01, C*07:02:01:01, DQB1*06:02:01:01, DRB1*15:03:01:01) was used for all samples. For samples where the reported haplotype call did not refer to a single unique sequence within the IMGT/HLA database, the alphanumerically first sequence was used. Reads from each sample were aligned to each reference using BWA-mem ^27^, and germline variants were called using GATK’s HaplotypeCaller ^75^.

### Alignment and mutation calling

HLA extracted reads were aligned to Hapster inferred reference sequences using BWA-mem ^27^. Mutation calling was performed using GATK’s Mutect2 ^28^. Mutations were filtered based on the following criteria: 1) Must pass the GATK filters *FilterMutectCalls* and *FilterByOrientationBias*, 2) The alternate base must not have been observed in the same position in other alleles of the same gene, 3) Read support in the tumor must be at least 3 reads, or at least 20% VAF, 4) Variant must have no more than 1 read support in the normal after a kmer search. For the kmer filter, all 25-mers covering the variant position are used to search for any matching reads in the normal in an alignment-free manner.

### Simulation validation

150 synthetic haplotypes were constructed by taking two random allelic sequences from the IMGT/HLA database (https://www.ebi.ac.uk/ipd/imgt/hla) for each MHC class 1 and class 2 gene (HLA-A, -B, -C, -DPA1, -DPB1, -DQA1, -DQB1, -DRA, -DRB1). For each synthetic haplotype, a mutation was inserted randomly using the following logic: 1) One allele from the haplotype was selected uniformly randomly 2) One position within the coding region of the allele was selected uniformly randomly 3) A mutational consequence was chosen with a 75% chance of creating an SNV, a 12.5% chance of creating a deletion, and a 12.5% chance of creating an insertion 4) For SNVs a random alternate base was chosen uniformly evenly, for deletions a deletion of length 1-5 nucleotides was chosen uniformly randomly, and for insertions an insertion of length 1-5 nucleotides was chosen uniformly randomly, with the insertion being uniformly random nucleotides. From each haplotype, inserts of length 125-300 were simulated from both the normal and mutated reference sequences for each sample using the BBmap ^76^ function randomreads.sh. Inserts were considered captured if they had significant overlap with MI-ONCOSEQ WES capture probes. Paired reads of length 125 were taken from the ends of each captured insert. Mutations were then called using the Hapster pipeline and were filtered using Mutect2 and Hapster filters.

### RNA validation of somatic mutations

All variants from the TCGA HNSC cohort with sufficiently high coverage in the RNA to detect low allelic fraction variants (>1000 reads at the variant position) were selected, resulting in a set of 75 variants. MHC class II variants were not selected for RNA validation as the expression of MHC class II genes is expected to be dominated by immune cells and not cancer cells. For each somatic mutation, a set of all 25-mers containing either the called variant or the germline sequence were created. In the case that other variants were called within 25 bases of the primary variant, kmers were produced with both the primary variant alone and with other variants in combination to account for possible phasing. Kmers that contained the variant within 8 bases of the edge of the kmer were discarded to avoid confounding results when deletions or insertions in small repetitive regions lead to variant kmers that are identical to the germline sequence. All reads within the RNA seq data for a sample were searched for germline or variant supporting kmers. Somatic mutations were considered validated if the number of variant supporting reads could not be explained by sequencing errors in the germline reads. We modeled sequencing errors as a Beta-Binomial distribution with the probability of error as a Beta distribution determined experimentally from RNA-seq data. An experimental null sample was created by selecting random bases at random positions within sequenced samples and evaluating the support for the random variant based on the null distribution. Variants were considered validated if the probability of observing equal or more variant supporting reads was less than 5% given our distribution.

### Sanger sequencing validation of somatic mutations

14 somatic mutations called within samples from the MI-ONCOSEQ project that had tissue samples available were chosen to be validated via Sanger sequencing. For each mutation, Primer3 was used to create custom PCR probes for the specific HLA allele that the mutation was called within. Genomic DNA from the tumor was amplified using PCR and amplicons isolated using gel electrophoresis. Following Sanger sequencing, point mutations were considered validated if a clear peak containing the called variant was observed, and indels were considered validated if a clear peak-offset corresponding to the number of inserted or deleted bases was observed.

### Tumor mutational burden calculations

Global mutation burden was calculated using mutations obtained from MAF files provided by the Broad Institute’s GDAC Firehose (https://gdac.broadinstitute.org/). Mutations observed within the capture kit region for each WES sample was divided by the total area covered by the capture kit. Capture kit information for each sample was obtained from official TCGA sample level metadata. Local mutation burden for each MHC gene was reported as the number of coding region mutations divided by the total exon length of each gene.

### Positive selection of somatic mutations

Positive selection was evaluated by CBaSE, a tool that provides gene-specific measures of the strength of positive or negative selection for functional mutations based on the distribution of synonymous and non-synonymous variants after accounting for sequence contexts and cohort-specific mutational signature activities. CBaSE was run on each TCGA and MI-ONCOSEQ cohort separately. For MHC genes, CBaSE was run on mutations called by Hapster. For all other genes, CBaSE was run on mutations reported via MAF files obtained from the Broad Institute’s GDAC Firehose (https://gdac.broadinstitute.org/). To calculate the pan-cancer metastatistic ϕ_pos_, the cohort-level ϕ_pos_ reported by CBaSE was summed across all cohorts.

### Mutual exclusivity and co-mutation analyses

Both mutual exclusivity and co-mutation analyses were performed at the cohort level. Mutual exclusivity of MHC class I and APM mutations were evaluated using CoMEt. Exact tests for mutual exclusivity were calculated by comparing any mutation with the MHC class I genes to any mutation within an APM gene. Co-mutation analyses were performed using a SNP-seq kernel association test as implemented in the R package SKAT (https://cran.r-project.org/web/packages/SKAT/). Only driver genes listed in the COSMIC Tier 1 Cancer Gene Census (https://cancer.sanger.ac.uk/census) were considered. Within each cohort, only genes mutated in more than 2% of patients were analyzed.

### AID/APOBEC mutational signature activity

Mutational signature activity within specific TCGA samples was obtained from the ICGC Pan Cancer Analysis Mutational Signatures Working Group (https://doi.org/10.7303/syn11726601). For each tumor within the CESC, HNSC, LUSC, and DLBC cohorts, the relative activity of AID/APOBEC associated mutational signatures SBS2 and SBS13 were identified. Cases were split into those either containing or not containing stop gain mutations within the MHC genes, and differences in SBS2/SBS13 activity were tested using t-tests.

### Mutation simulations

Mutations were simulated for each significantly mutated cohort by taking into account trinucleotide mutational signatures that are active in each cancer type. Mutational signature activity was obtained from the ICGC Pan Cancer Analysis Mutational Signatures Working Group (https://doi.org/10.7303/syn11726601). 10,000 mutations for each cohort were simulated as follows: 1) Each base along the length of HLA-A*01:01:01:01 was weighted based on its trinucleotide context, and the probability of that trinucleotide being mutated given known mutational signature activity. 2) A random position was picked based on the weighted probabilities of each base being mutated. 3) A random alternate base was picked for the selected position, with each potential alternate base weighted based on trinucleotide context and mutational signature activity.

### MHC amino acid annotations

Protein interaction analysis was performed using annotations derived from structures of the relevant MHC class I complexes (PDB IDs provided below). The distance cutoff of the contacts was ≤4 Å. Structures used for MHC class I : peptide interaction and the interaction between MHC class I heavy chain and B2M were 4NQV, 6IEX, 3MGO, 3KPL, 3RL2, 3DX7, 4F7M, 1E28, 4HX1, 2RFX, 5EO0, 5IM7, 1XR8, 5W6A, 1JGE, 5VGE, 3LKR and 6JTP. Structure used for MHC class I : CD8 interaction was 3DMM. Structures used for MHC class I : TCR interaction were 5WKF, 4G8G, 5WKH, 4G9F, 5NQK, 5XOT, 6EQA, 4PRP, 3VXM, 6BJ2, 3W0W, 3MV7, 6AVF, 4JRX, 6AVG, 4JRY, 4QRP, 4PRI, 1MI5, 3DXA, 3FFC, 3KPR, 3SJV, 3KPS and 4QRP. Structures used for MHC class I : TAPBPR interaction were 5OPI and 5WER. Structure used for MHC class I : TAPBP interaction was 6ENY.

### PPI binding energy predictions

Predicted changes to the MHC-B2M interface binding energy were generated using SSIPe. All predictions were performed using the 3D structure at https://www.rcsb.org/structure/4U6X. While the MHC class I proteins are highly polymorphic, the residues that make up the B2M interface are highly conserved, allowing us to predict binding energy changes using a single 3D structure regardless of which MHC protein the mutation was called in.

### Statistics

For comparisons of levenshtein distance from HapMap samples to Grch38 or Hapster references, t-tests were performed. For comparison of MHC mutation rates between cancers approved or not for immune checkpoint inhibitors, and comparison of MSI frameshifts vs MSS frameshifts, Fisher’s exact test was used. For comparisons between functional consequence distributions of MHC mutations in significantly mutated cohorts to that of the neutral model, the Chi-Squared test was used as sample sizes were too large for Fisher’s exact test. For comparisons between primary and metastatic MHC mutation rates, ANOVA was performed with mutational burden as a covariate. For associations between local and global TMB, data distributions did not meet the assumptions for linear regression, so Spearman’s rank correlation was used. For local dN/dS ratios within various MHC annotation regions, the Shapiro-Wilks normality test was performed to show that samples were normally distributed, and t-tests were performed to determine if samples had means significantly different than 0.

### MHC class I LOH Calls

To call the Loss of heterozygosity (LOH) for each gene per sample, we used Hapster generated BAMs, containing the HLA reads mapped to the inferred haplotypes references. We used BAM files to calculate the read coverage for each allele of each HLA gene. Using those calculated coverages we were able to estimate the B-Allele frequency (BAF) and Coverage Log2-ratios (LR) values for each gene, both in tumor samples and their matching normal sample. To call a gene with LOH, a gene must have met 3 conditions: (I) tumor BAF value is deviated from 0.5 by more than 0.1 to indicate a possible LOH, (II) normal BAF value is not deviated from 0.5 by more than 0.1 to ensure the detected deviation in tumor sample is not due to any biases in mapping or reference detection and finally, (III) the LR value is less than 0.01 to ensure the deviation in tumor samples is not due to a amplification, but rather a possible copy-neutral/deleterious LOH. We considered these conditions for all the MHC-I genes separately per sample.

## Data availability

WES and RNA-seq data for all TCGA samples were obtained from the NIH Genomic Data Commons portal (https://portal.gdc.cancer.gov). MAF files containing TCGA WES mutation calls were obtained from the Broad Institute’s GDAC Firehose (https://gdac.broadinstitute.org/). Mutational signature activity within specific TCGA samples was obtained from the ICGC Pan Cancer Analysis Mutational Signatures Working Group (https://doi.org/10.7303/syn11726601). WES and targeted sequencing data from MI-ONCOSEQ were obtained from (Cobain et al., 2021; Robinson et al., 2017), with raw sequencing data supporting the HLA mutation calls made available through Zenodo (ID). HLA reference sequences were downloaded from the IMGT/HLA public database release (https://www.ebi.ac.uk/ipd/imgt/hla). HLA mutation calls are provided in the supplement (**Supplementary tables S8-S9**). Crystal structures used to identify HLA residues involved in protein:protein interactions were obtained from the RCSB protein data bank (https://www.rcsb.org).

