## Supplemental figures S1-S5 for "Distinct mutational processes shape selection of MHC class I and class II mutations across primary and metastatic tumors"

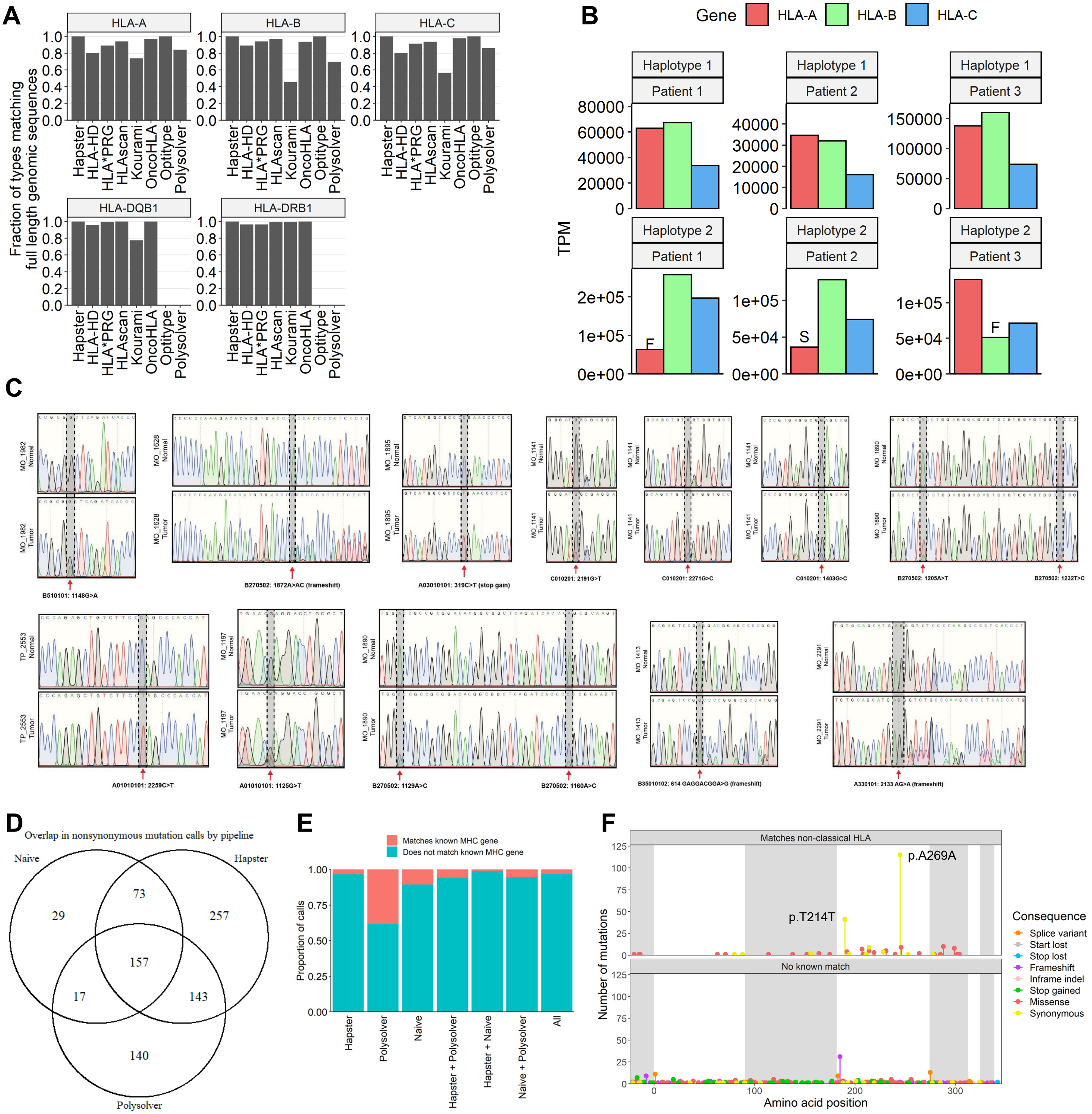

**Supplementary figure S1 - Validation of hapster**

**(A)** Fraction of HLA types from 69 1000 Genomes samples reported by 8 different HLA haplotypers that correspond to a full length genomic sequence found in the IMGT/HLA database. HLA haplotypers often report types that only have known sequence for the polymorphic binding pocket, and full length genomic sequences are unavailable. **(B)** Reduced transcript levels observed in alleles with truncating stop-gains or frameshifts, potentially caused by nonsense mediated decay. In each case, haplotype 1 shows expected RNA expression with HLA-A and HLA-B expressed at a higher level than HLA-C. Haplotype 2 shows allele specific expression losses within alleles containing a truncating frameshift (F) or stop gain (S) mutation. **(C)** Sanger traces for 14 mutations, representing single-nucleotide variants, multi-nucleotide variants, and frameshift variants, called in MI-ONCOSEQ samples. **(D)** Venn-diagram showing overlap of non-synonymous mutation calls for the MHC class I genes between the naive GDC pipeline, Polysolver, and Hapster. **(E)** Proportion of nonsynonymous mutation calls that match known DNA sequences from homologous MHC genes and pseudogenes and could provide an alternative explanation as misaligned reads rather than actual somatic variants. Bars with '+' represent intersections. **(F)** Positions of mutations called by the Polysolver pipeline, separated by whether or not the variant matches a known sequence in another non-classical MHC class I gene. Annotated recurrent synonymous variants are suspected false positives.

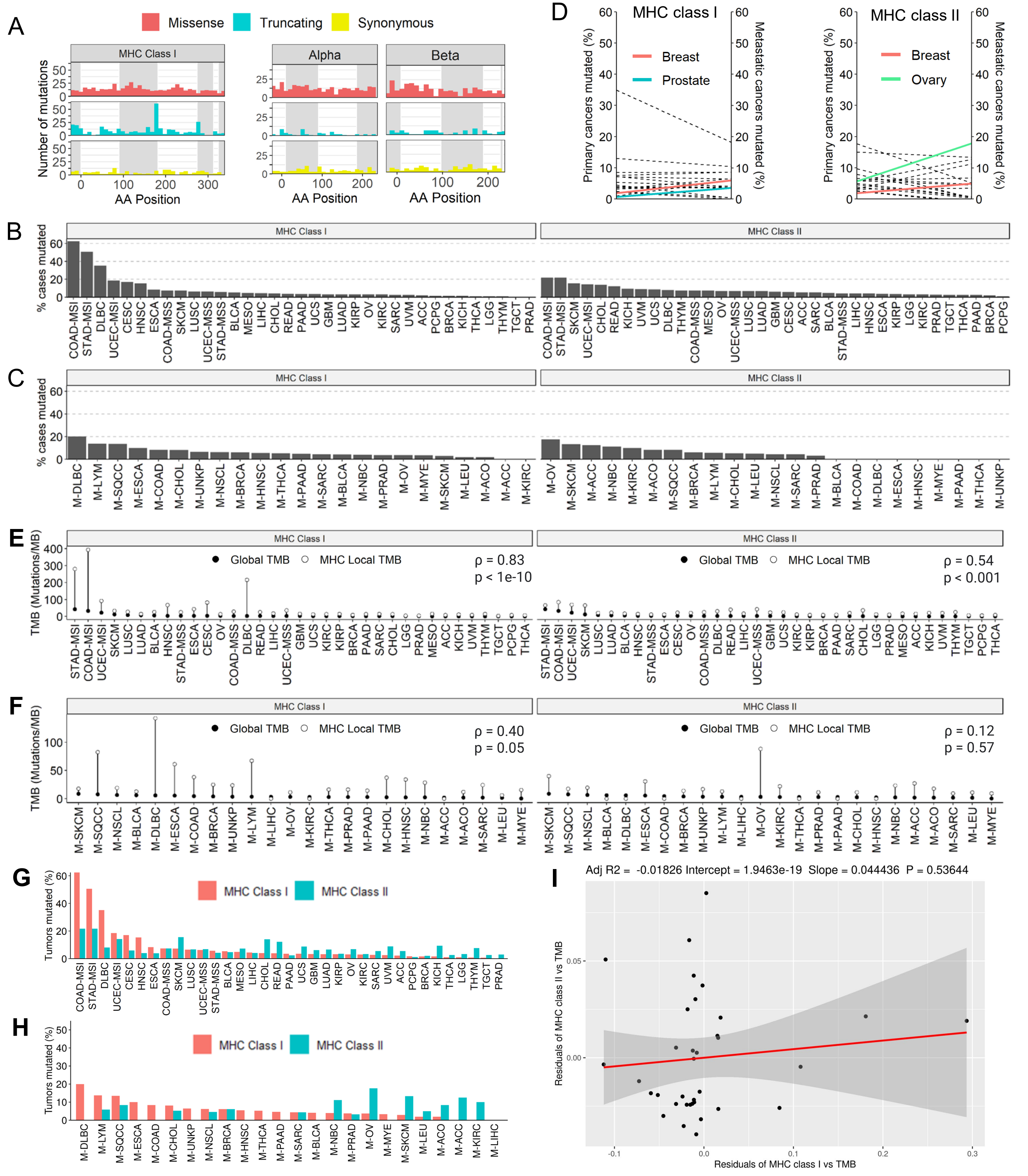


**Supplementary figure S2 - MHC class 1 and class 2 mutation distributions and associations, related to figure 2**

**(A)** Distribution of MHC mutations by mutation consequence, MHC class, and peptide chain. Alternating grey/white backgrounds show exon boundaries. **(B, C)** Fraction of primary cancers from TCGA **(B)** and metastatic cancers from MI-OncoSeq **(C)** harboring mutations in any MHC class I and any MHC class II gene. **(D)** Change in percent of tumors harboring MHC class I or MHC class II mutations between primary and metastatic cancers. Values are marginal means after adjusting for tumor mutation burden. Cohorts with significantly different numbers of mutated primary and metastatic cases are colored. Sample sizes are slightly reduced from full cohort sizes due to missing WES mutation calls with which to calculate the global tumor mutation burden covariate. Breast: primary n = 854, metastatic n = 289; Prostate: primary n = 434, metastatic n = 478; Ovary: primary n = 246, metastatic n = 17. **(E, F)** Comparison between global TMB and local TMB within MHC class I and MHC class II genes in primary **(E)** and metastatic **(F)** cancers. Average global TMB is calculated based on non-synonymous mutations in all protein-coding genes, average MHC local TMB is the number of mutations in MHC class I or class II genes divided by their length. **(G, H)** Proportion of tumors with mutations in MHC class I or MHC class II in **(G)** primary and **(H)** metastatic tumors. **(I)** Residual correlation between the numbers of MHC class I and class II mutations in primary tumors after adjusting for tumor mutational burden. Points correspond to each of the 35 cancer types from TCGA.

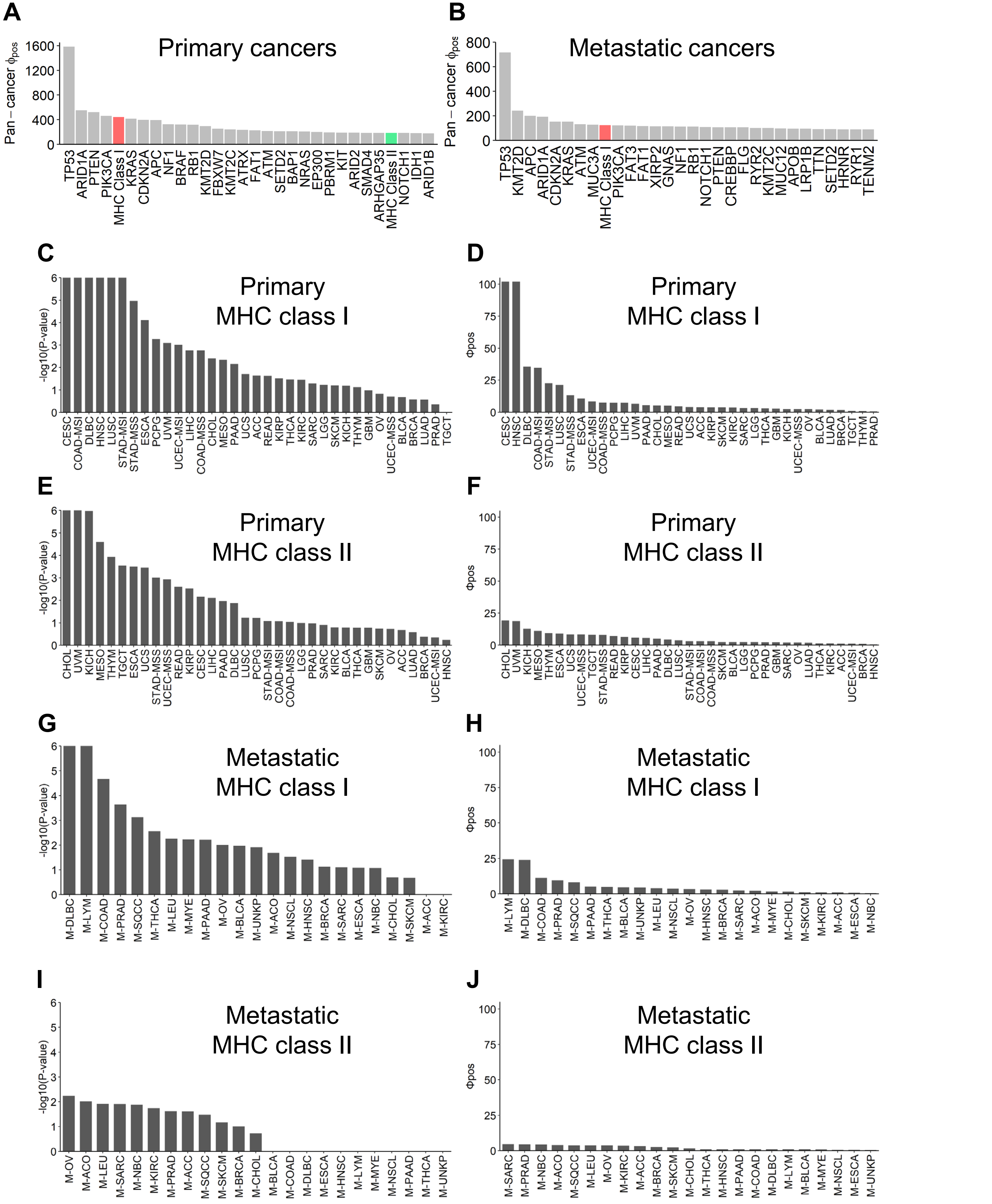


**Supplementary figure S3 - Positive selection in primary and metastatic cancers, related to figure 3**
**(A, B)** Top-30 positively selected genes by summed metastatistic $\phi_{pos}$ for evidence of positive selection across all primary **(A)** and metastatic/refractory **(B)** cohorts pan-cancer. **(C-J)** Cohort-level evidence for MHC class I and II positive selection as reported by CBaSE. P-values **(C, E, G, I)** and metastatistic $\Phi_{pos}$ **(D, F, H, J)** are shown. P-values reported by CBaSE are capped at 1e-6. Ties between cohorts reaching this level of significance can be broken using the metastatistic $\Phi_{pos}$.

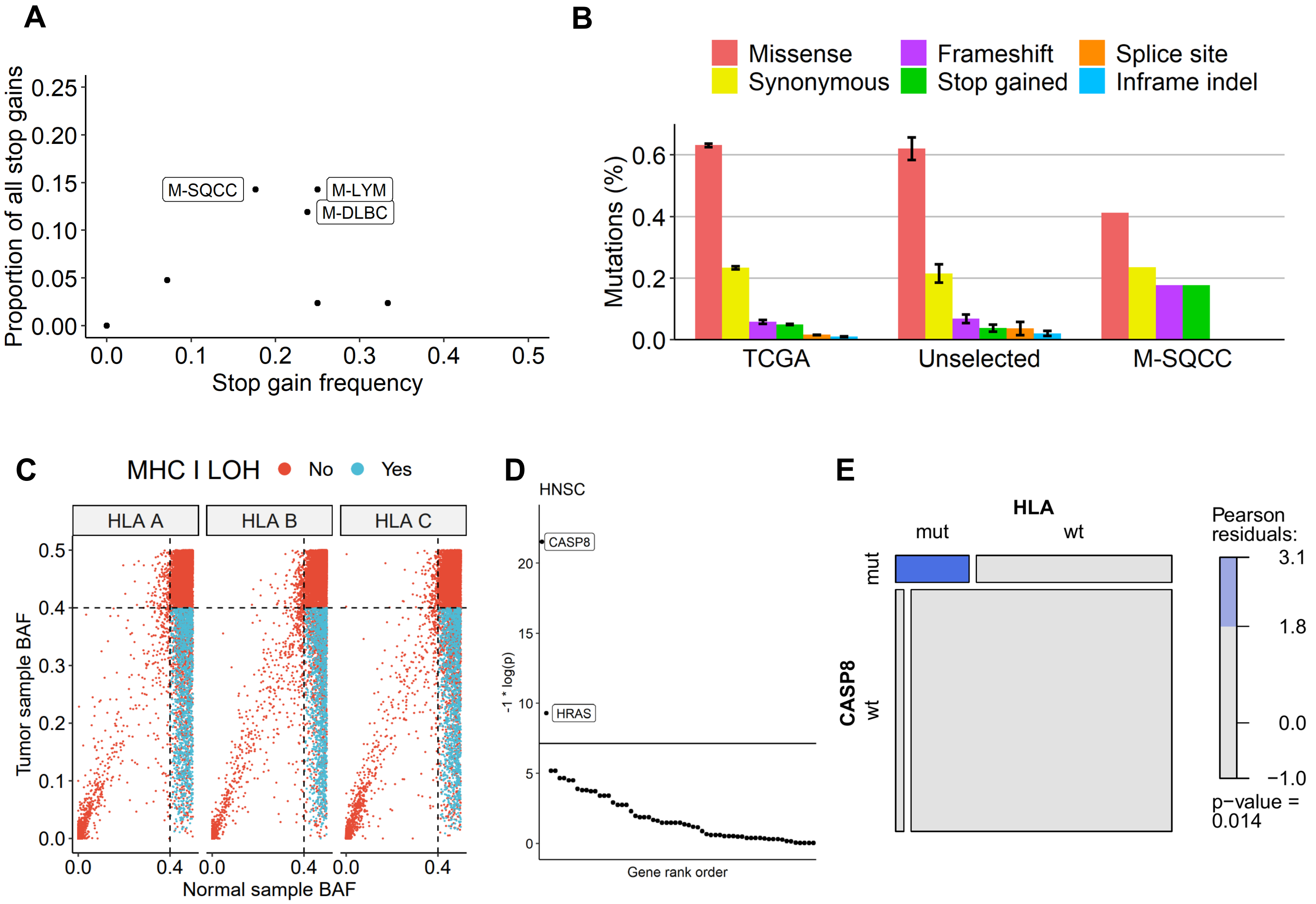


**Supplementary figure S4 - Related to figure 3**

**(A)** Cohort specific MHC stop gain frequency compared to overall proportion of stop gains contributed by each MI-ONCOSEQ cohort. Enriched cohorts are labeled. **(B)** Proportion of functional consequences observed in various groups: "TCGA" 2,600,654 pan-cancer mutations from TCGA, an approx. neutral model; "Unselected" - MHC class I mutations from all primary and metastatic cohorts showing no evidence of positive selection; "M-SQCC" - MHC class I mutations from the M-SQCC cohort. "TCGA" and "Unselected" are average frequencies across cohorts, with error bars showing SEM. **(C)** BAF of normal vs tumor samples used to establish thresholds for LOH calls. Normal samples show variability with BAF ranging from 0.4 to 0.5. Samples with normal BAF lower than 0.4 were rejected for LOH calls. Tumor samples were only accepted as having LOH if the BAF in the tumor was lower than 0.4. Finally, cases with normal BAF > 0.4 and tumor BAF < 0.4 were still rejected if the LR value was greater than 0.1, which could imply amplification rather than LOH. **(D)** Co-mutation analysis for functional mutations in all cancer gene census tier 1 genes vs MHC class I mutations in the HNSC cohort. Horizontal lines show significance cutoff after Bonferroni correction. Significantly co-mutated genes are labeled. **(E)** Pearson residual plot showing the enrichment of tumors with both CASP8 and MHC class I mutations in an independent cohort of 109 HNSCC tumors.

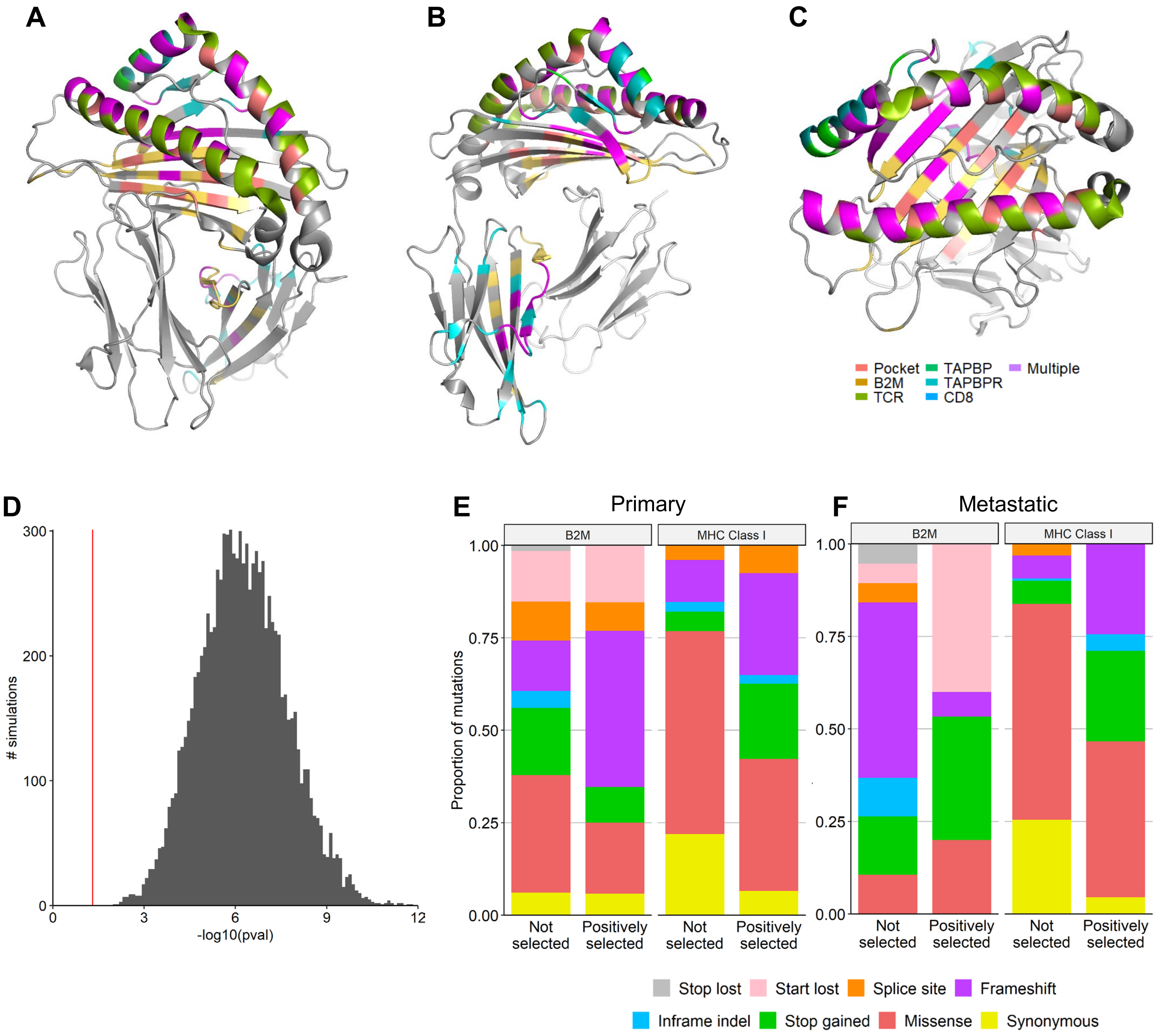


**Supplementary figure S5 - Mutations at the MHC1:B2M interface, related to figure 5**
**(A-C)** Multiple views of the MHC class I crystal structure. Residues are colored based on interactions with MHC associated proteins. **(D)** P-values (t-test) comparing observed mutations at the β2m interface to a set of random samples of the same size from a pool of simulated mutations. 10,000 random samples were taken and tested, all of which showed a significant difference (red line, $p < 0.05$) *i.e.* all random sets of mutations had significantly smaller ΔΔG compared to the observed ones. **(E, F)** B2M shows a high rate of loss of function mutations in both **(E)** primary and **(F)** metastatic/refractory cancers that do or do not show evidence of positive selection for MHC class I mutations. Mutational consequence distributions of mutations within the MHC class I proteins are provided for comparison.
