## Supplemental notes S1-S2 for "Distinct mutational processes shape selection of MHC class I and class II mutations across primary and metastatic tumors"

**Supplementary note S1. Hapster dynamic reference selection summary, related to figure 1**

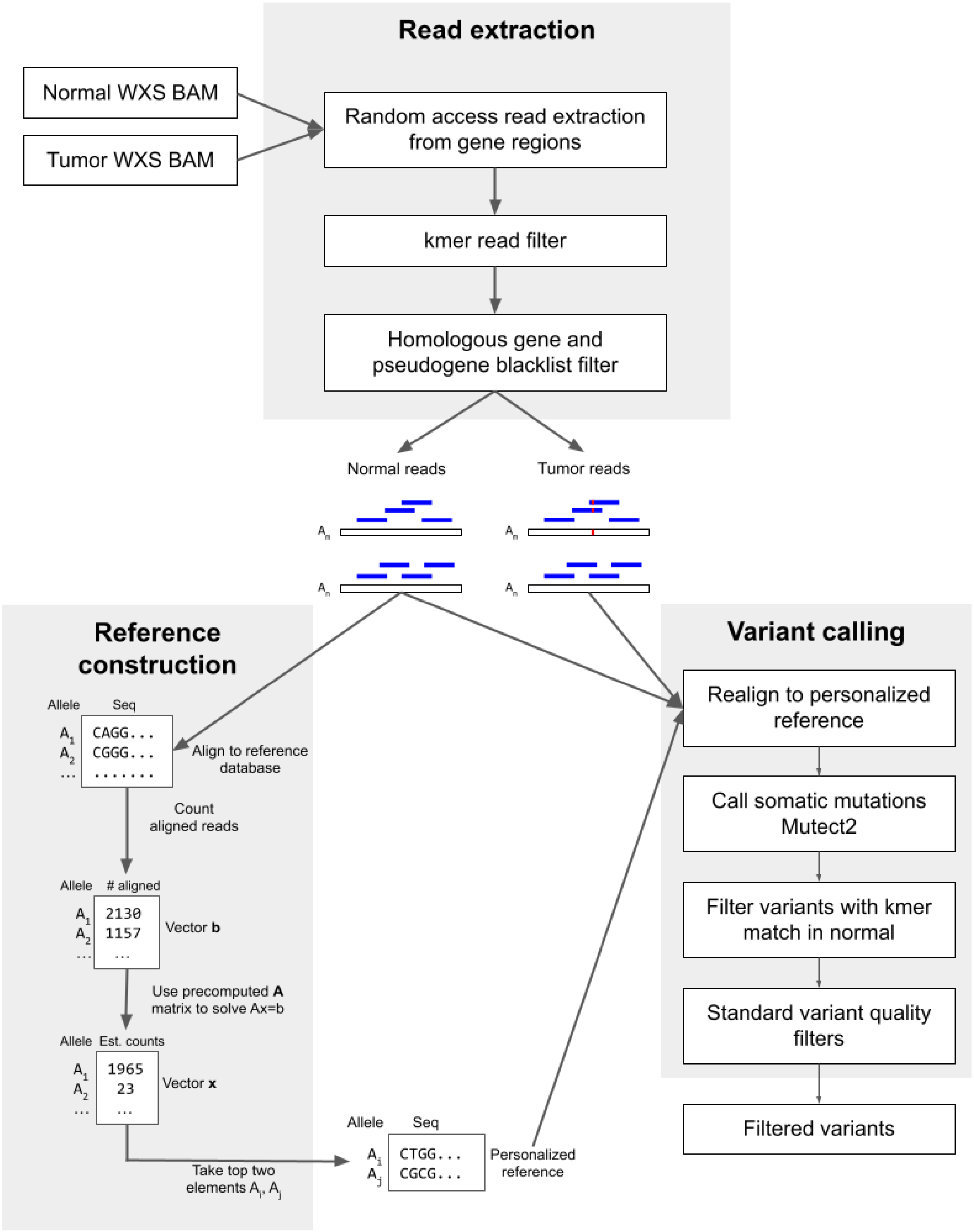


**Schematic overview of Hapster's mutation calling pipeline.**

The current paradigm for mutation calling involves the alignment of DNA sequencing reads to a standard reference genome, followed by identification of variants relative to that reference. This approach fails for the MHC genes where high sequence divergence from the standard reference, as well as the presence of homologous pseudogenes, frequently causes reads to fail to align appropriately.

To address this problem, we have developed Hapster. Hapster models alignment of sequencing reads to alternative haplotypes as a linear system that can be solved to identify the closest underlying haplotypes **(Figure 1A)**, a method which generalizes to other polymorphic genes. We also use a curated blacklist of homologous genes and pseudogenes to remove reads that erroneously cross-map to our genes of interest. Given appropriate haplotype references and filtered reads, Hapster builds upon state-of-the-art aligners and mutation callers (BWA-MEM[1] and Mutect2[2]) to provide accurate mutation calls. As a final step, we use an alignment-free kmer search to flag variants that may have been called due to remaining erroneous alignments or sample contamination.

The Hapster reference selection algorithm, in brief, is as follows. First, reads are simulated from each sequence within a database of known MHC alleles. Reads are then aligned simultaneously to all other known sequences. An all vs all matrix **A** is constructed where each entry describes the percentage of reads simulated from one allele that align well to another allele. Following sequencing, MHC reads are extracted and simultaneously aligned to all alleles within the MHC sequence database. Observed counts of reads that align to each allele are put into a vector **b**. The system **Ax = b** is solved for **x**, where **A** describes alternate-sequence aware alignment, **x** describes the number of reads originating from each known allele, and **b** describes the number of reads that are aligned to each known allele. After solving, vector **x** should contain all values of ~0 except for the one or two (depending on whether the individual is homozygous or heterozygous at the locus) non-zero entries representing alleles that were actually present in the germline and therefore generated sequencing reads. These two non-zero entries are taken as the most likely diploid haplotype for the individual, and their reference sequences are used for downstream alignment and mutation calling. Personalized references provide improved alignments, and allow for recovery of missed mutation calls relative to the standard reference approach **(See IGV view below)**.

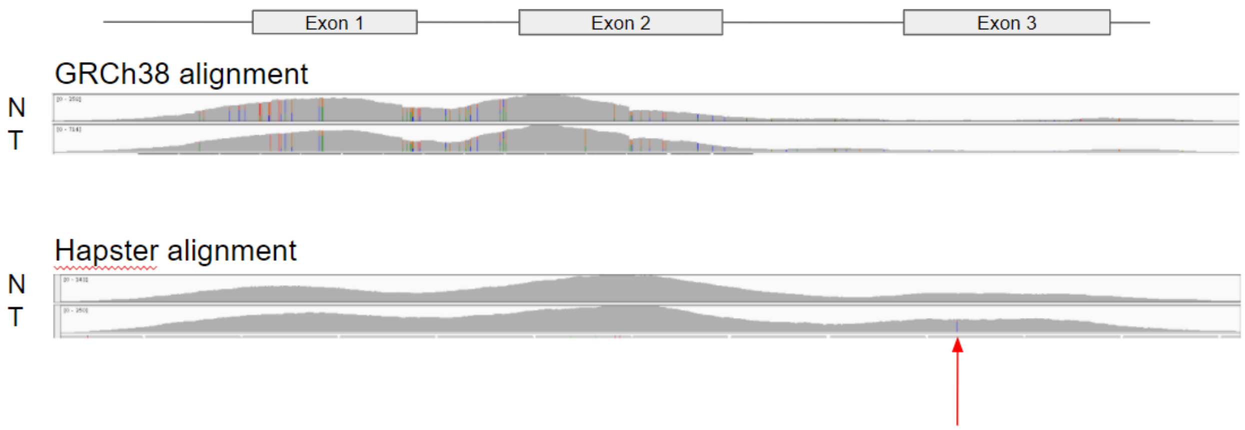


**Recovery of somatic mutations missed by naive pipelines**
IGV view of sequencing reads from paired normal and tumor whole exome sequencing data aligning to the first 3 exons of HLA-A using the standard reference GRCh38 or Hapster. Arrow shows a somatic variant missed by the standard reference approach due to a failure to align reads from exon 3 to the correct location. N: Normal, T: Tumor.

**Supplementary note S2. Complete Hapster pipeline description, related to figure 1**

*Alt-aware reference construction*

Hapster leverages BWA mem's alt-aware alignment mode for its haplotyping, and as such needs a genomic reference that contains alternate sequences for the genes of interest. To construct an alt-aware reference for the HLA class I and class II genes we first retrieved sequences from the IMGT/HLA database [3] to create a set of alternate alleles for each gene. The sequences for each alternate allele were appended to the primary assembly of Grch38 as independent contigs, and an alt index file was created by performing long read alignment of each sequence to Grch38 using minimap2[4].

*HLA read kmer extraction*

To efficiently extract true HLA reads, we created a 4 step procedure: (1) Random access retrieval of reads from all HLA regions in Grch38, all unaligned reads, and additional locations throughout Grch38 where we have found HLA reads mapping erroneously. (2) Passing reads retrieved by random access through a kmer filter, keeping any read that contains any 30-mer found in our set of alternate HLA allele sequences (3) Alt-aware alignment of kmer extracted reads to the previously created reference containing our set of alternate HLA alleles, keeping only reads that have at least one alignment to an HLA contig (4) Alignment of remaining reads to a reference containing both the sequences for our genes of interest (whitelist), as well as the sequences for any homologous genes/pseudogenes that are not of interest (blacklist), and keeping only reads that preferentially align to the whitelist.

*HLA haplotype inference*

For each HLA gene, extracted reads are aligned using BWA-MEM's alt-aware alignment mode to the constructed reference containing our set of alternate HLA sequences. The number of read pairs aligning to each allele with a total NM score of less than or equal to 1 is counted and put into vector **b**. Using vector **b** and the precomputed probability matrix **A** (construction of **A** matrix described below), the denoised read vector **x** is calculated by solving the equation **Ax=b**. To reduce problems caused by highly correlated alleles, a dynamic stepwise variable selection process is used to eliminate variables from **A** as follows: (1) identify alleles with pairwise correlations above a specific threshold (parameter tuning for this threshold described below) (2) For each highly correlated pair of alleles, find the magnitude difference between the alleles within vector **x** (3) Remove the most negative allele within the pair with the highest magnitude difference from both matrix **A** and vector **b** (4) Solve for **x** using the pared **A** and **b** (5) Repeat until there are no more pairs of alleles with correlations above the predetermined correlation cutoff. Once all highly correlated alleles are removed, the two highest value alleles in vector **x** are assigned to the individual's haplotype. The top two alleles are always chosen due to our observation that during alignment of reads to our inferred haplotype, a homozygous individual's reads will preferentially align to the single allele that is from their true haplotype. The presence of an extra allele's sequence in the reference file in the homozygous case therefore does not affect alignment or mutation calling of reads aligning to the true allele.

*Construction of probability matrix A*

Insert size metrics for a given sequencing protocol are calculated using the Picard tools command CollectInsertSizeMetrics. Using the read length, mean insert size, and standard deviation for the protocol, a random set of read pairs are simulated from the genomic sequence of each allele within a given HLA gene as follows: (1) Inserts of mean insert size ± 2 standard deviations are simulated using BBMap's randomreads.sh (2) To simulate exome capture, read pairs derived from each insert are only kept if they have significant overlap with a provided set of capture probe sequences. The simulated captured reads are aligned in an alt-aware manner to our reference containing all alternate HLA alleles and are then processed and counted to create a vector **b** as described in the methods for haplotype inference. The vector **b** is then divided by the total number of read pairs that were simulated to obtain a vector that reflects the probability that any randomly selected read derived from the simulated allele would align to each other allele when using alt-aware alignment. This probability vector is calculated for each allele, and the probability matrix **A** is constructed using each of these probability vectors as its columns. The simulation process leads to a slightly non-symmetrical matrix, so the matrix was made to be symmetric by taking the mean of each paired off-diagonal term. The final matrix **A** is an all-vs-all matrix where each element represents the probability of reads generated from one allele aligning to another allele.

*Parameter tuning*

Parameter tuning must be performed to find the optimal correlation cutoff values for stepwise variable selection for every new sequencing experiment **(See cutoff heatmap below)**. Parameter tuning with HapMap samples was performed to find the optimal correlation cutoffs to minimize Levenshtein distances across all genes. For many experiments this will not be possible as ground truth haplotypes will be unavailable, making parameter tuning with Levenshtein distance impossible. However, we reasoned that every mismatch that contributes to the true Levenshtein distance could be recognizable as a germline mutation relative to the inferred reference sequences. To test this, we realigned sequences from our validation set to Hapster's inferred haplotype sequences and called germline mutations using HaplotypeCaller. We found strong correlations between the true Levenshtein distance and the number of germline mutations called **(See correlation plots below)**, suggesting that parameter tuning can be performed using the number of germline mutations called as a proxy for Levenshtein distance. For parameter tuning for TCGA and MI-ONCOSEQ samples, all correlation cutoff values between 0.70-0.99 were tested to find the optimal threshold per gene that minimizes observed germline variants.

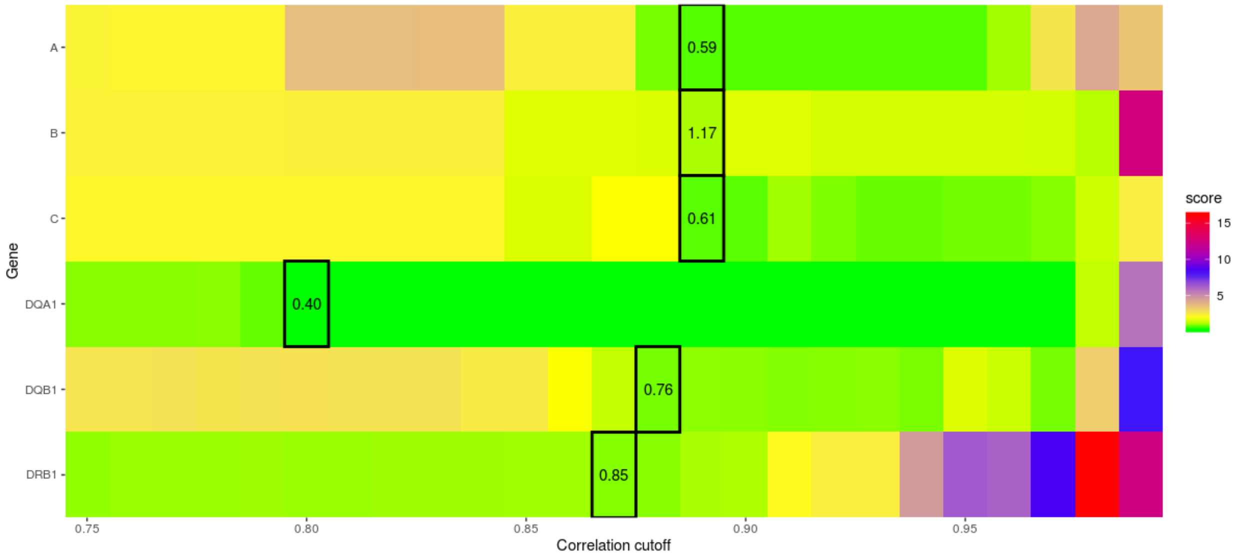


**Correlation cutoff parameter tuning**
Example of the search space for parameter tuning for optimal correlation cutoff values in the stepwise variable selection step. For each gene, correlation cutoff values from 0.75 to 0.99 are tested. Optimal values minimize either Levenshtein distance of germline variants, depending on what information is available. Selected parameters given this data set are highlighted. Values inside boxes denote the average Levenshtein distance of Hapster inferred references to ground truth sequences for 29 samples.

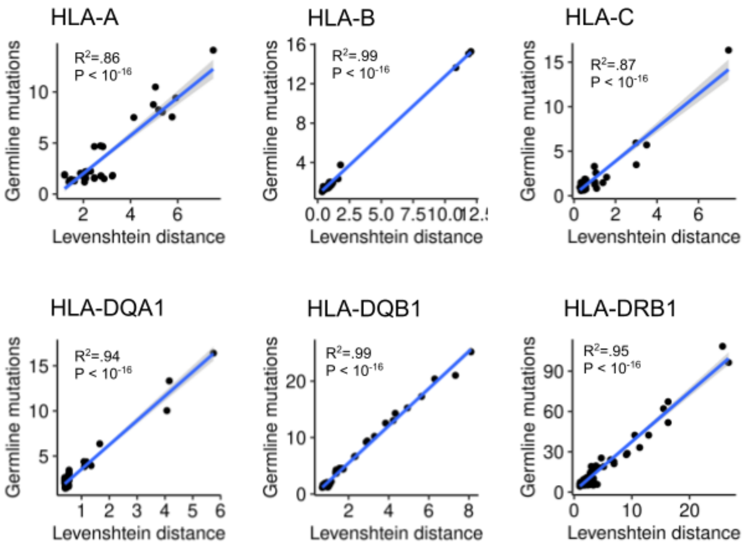


**Private germline variants as a proxy for Levenshtein distance**
Correlation plots showing high concordance between private germline variants and Levenshtein distance as a quantitative measure for similarity of a reference sequence to the true underlying genomic sequence.

*Mutation calling and filtering*

For each gene, the extracted read set is re-aligned with BWA-mem[1] to a new genomic reference file dynamically constructed to only contain the genomic sequence of each allele in the inferred HLA haplotype. In this new genomic reference file, each HLA allele is present as an individual contig. Mutations are called in the re-aligned BAM using the GATK4 function Mutect2[2], and variants are filtered using the function FilterMutectCalls. The GATK4 filter FilterByOrientationBias was applied to remove 8-Oxog artifacts that were observed within TCGA cohorts. We additionally removed any low coverage variants that had 100% orientation bias even if there were not enough reads to fail the bias test. We next constructed a custom kmer filter that searches for the presence of any reads supporting a called variant in an alignment free manner, removing any variants that find kmer support in the normal. 25-mers covering the variant were used, excluding those kmers where the variant is within 8 bases from the edge of the kmer. As a final filter, we removed any somatic variants that matched a known polymorphism for an alternative allele of the same gene within the HLA database. Known polymorphisms were identified by looking at a variant position within the global multiple sequence alignment of all alleles of a given gene. Only single nucleotide variants were identified this way.

**Supplementary notes references**


1. Li, H. (2013). Aligning sequence reads, clone sequences and assembly contigs with BWA-MEM. arXiv [q-bio.GN].

2. Benjamin, D., Sato, T., Cibulskis, K., Getz, G., Stewart, C., and Lichtenstein, L. (2019). Calling Somatic SNVs and Indels with Mutect2. bioRxiv, 861054. 10.1101/861054.

3. Robinson, J., Halliwell, J.A., Hayhurst, J.D., Flicek, P., Parham, P., and Marsh, S.G.E. (2015). The IPD and IMGT/HLA database: allele variant databases. Nucleic Acids Res. *43*, D423–D431.

4. Li, H. (2018). Minimap2: pairwise alignment for nucleotide sequences. Bioinformatics *34*, 3094–3100.
