## Supplemental data for "Distinct mutational processes shape selection of MHC class I and class II mutations across primary and metastatic tumors"

### Supplementary data S1, related to figure 2

This file contains visualizations of the position and mutational consequence of all individual variants called by Hapster within primary cancers from all 35 TCGA cohorts and all metastatic/refractory cancers from 24 MI-ONCOSEQ cohorts. Mutations are shown for the MHC class 1 genes (HLA-A, -B, -C), MHC class 2 alpha genes (HLA-DPA1, -DQA1, -DRA), and MHC class 2 beta genes (HLA-DPB1, -DQB1, -DRB1). Alternating grey/white backgrounds denote exon boundaries.

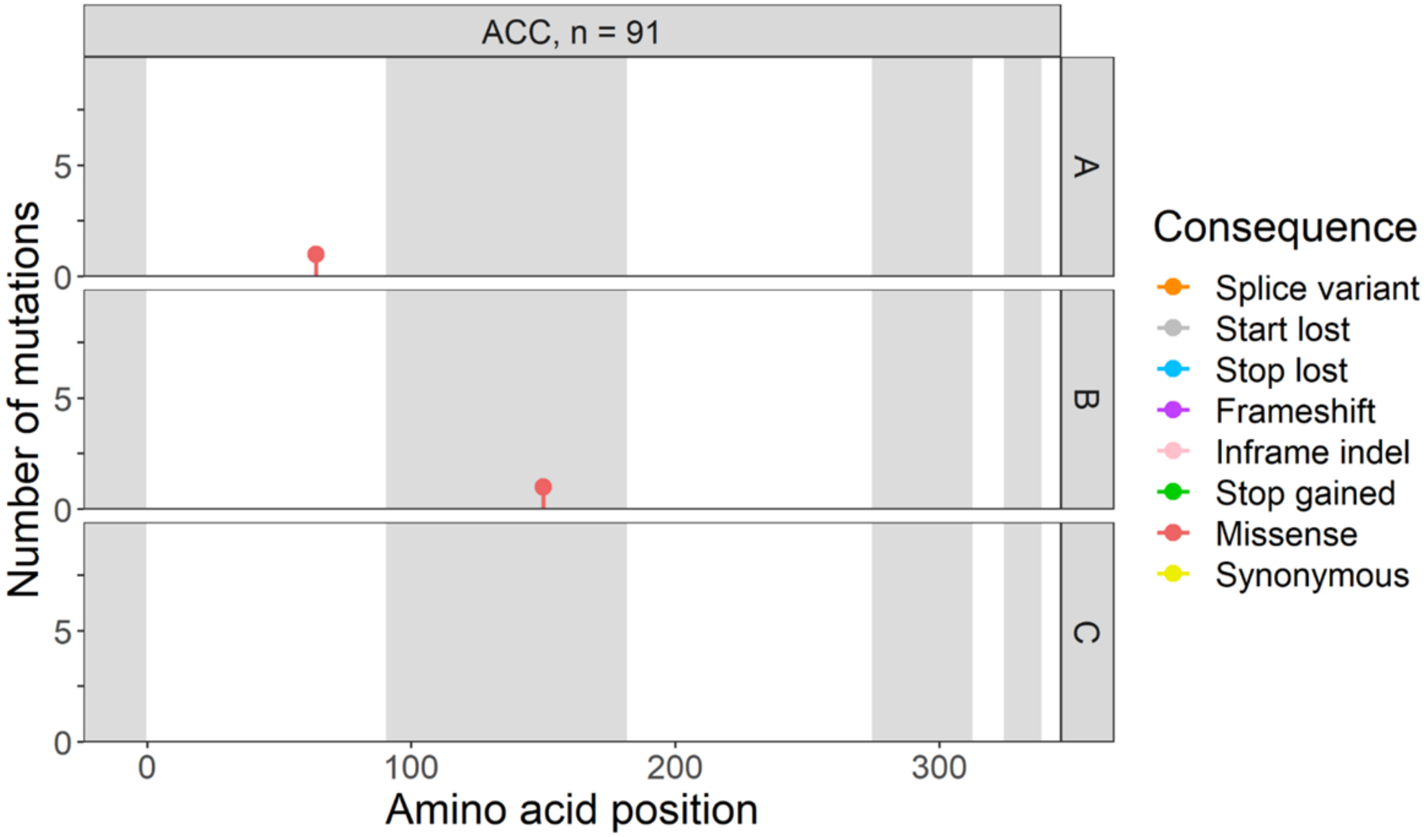

ACC, n = 91
A
B
C
Number of mutations
Amino acid position
Consequence
Splice variant
Start lost
Stop lost
Frameshift
Inframe indel
Stop gained
Missense
Synonymous


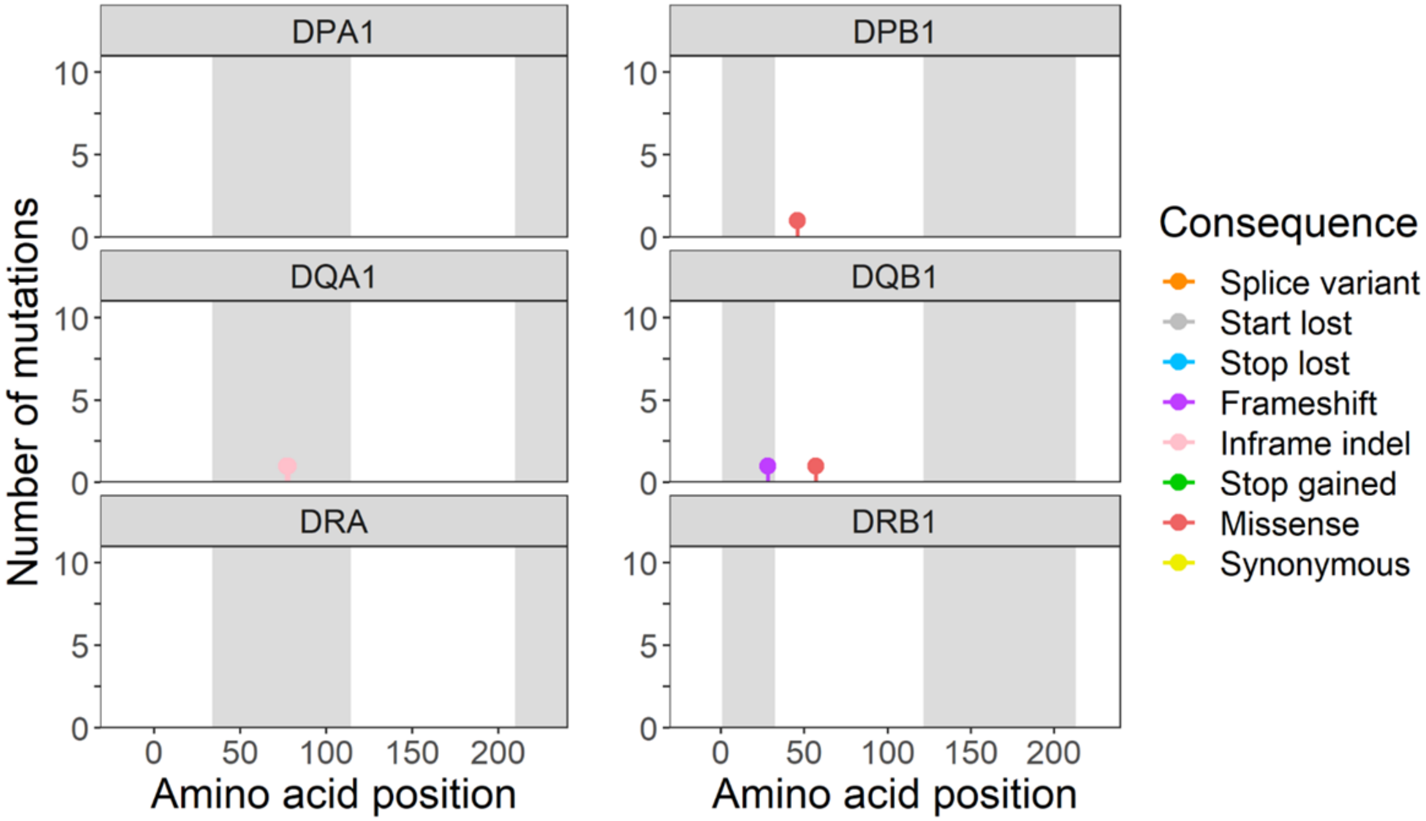

DPA1
DPB1
DQA1
DQB1
DRA
DRB1
Number of mutations
Amino acid position
Amino acid position
Consequence
Splice variant
Start lost
Stop lost
Frameshift
Inframe indel
Stop gained
Missense
Synonymous

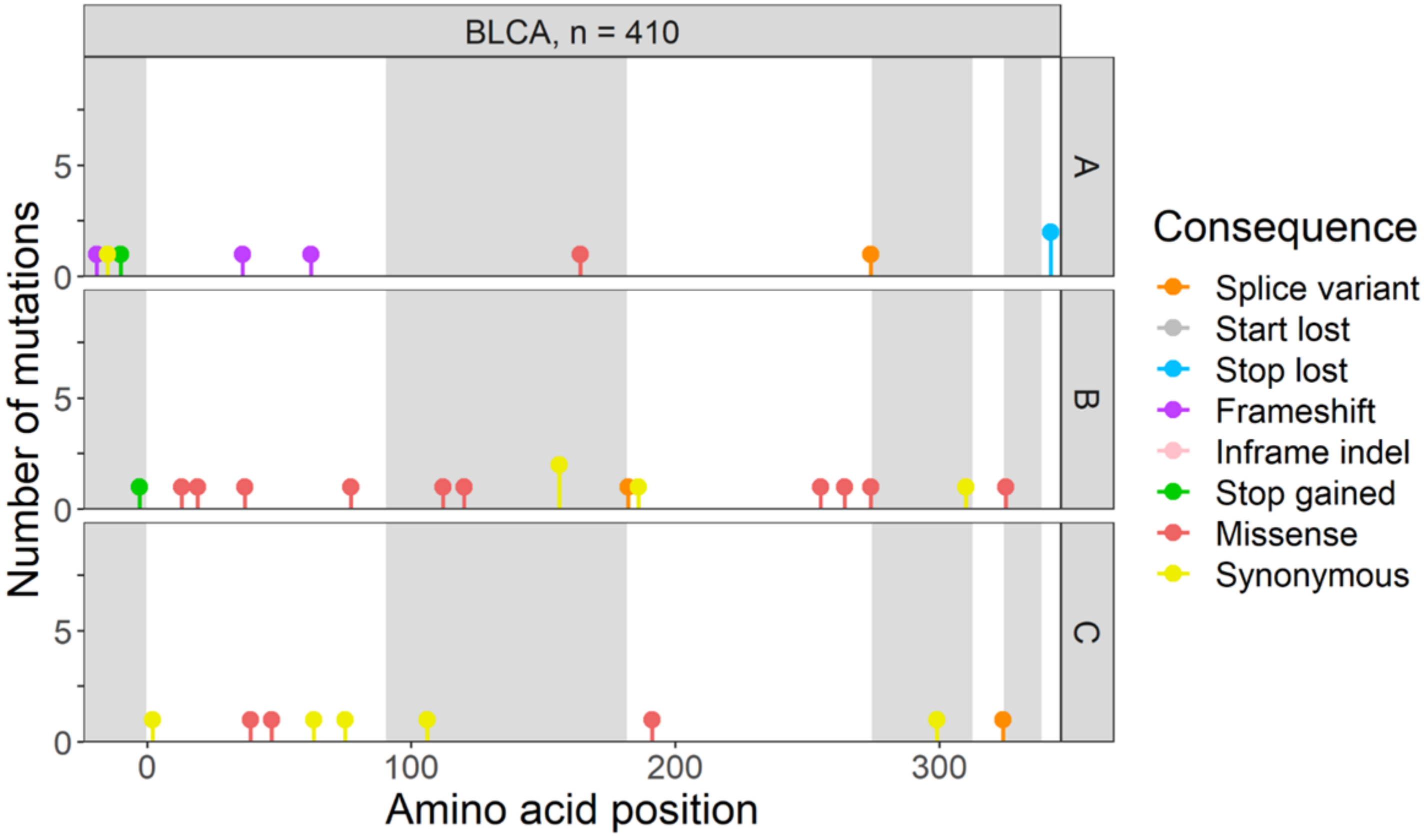

BLCA, n = 410
A
B
C
Number of mutations
Amino acid position
Consequence
Splice variant
Start lost
Stop lost
Frameshift
Inframe indel
Stop gained
Missense
Synonymous


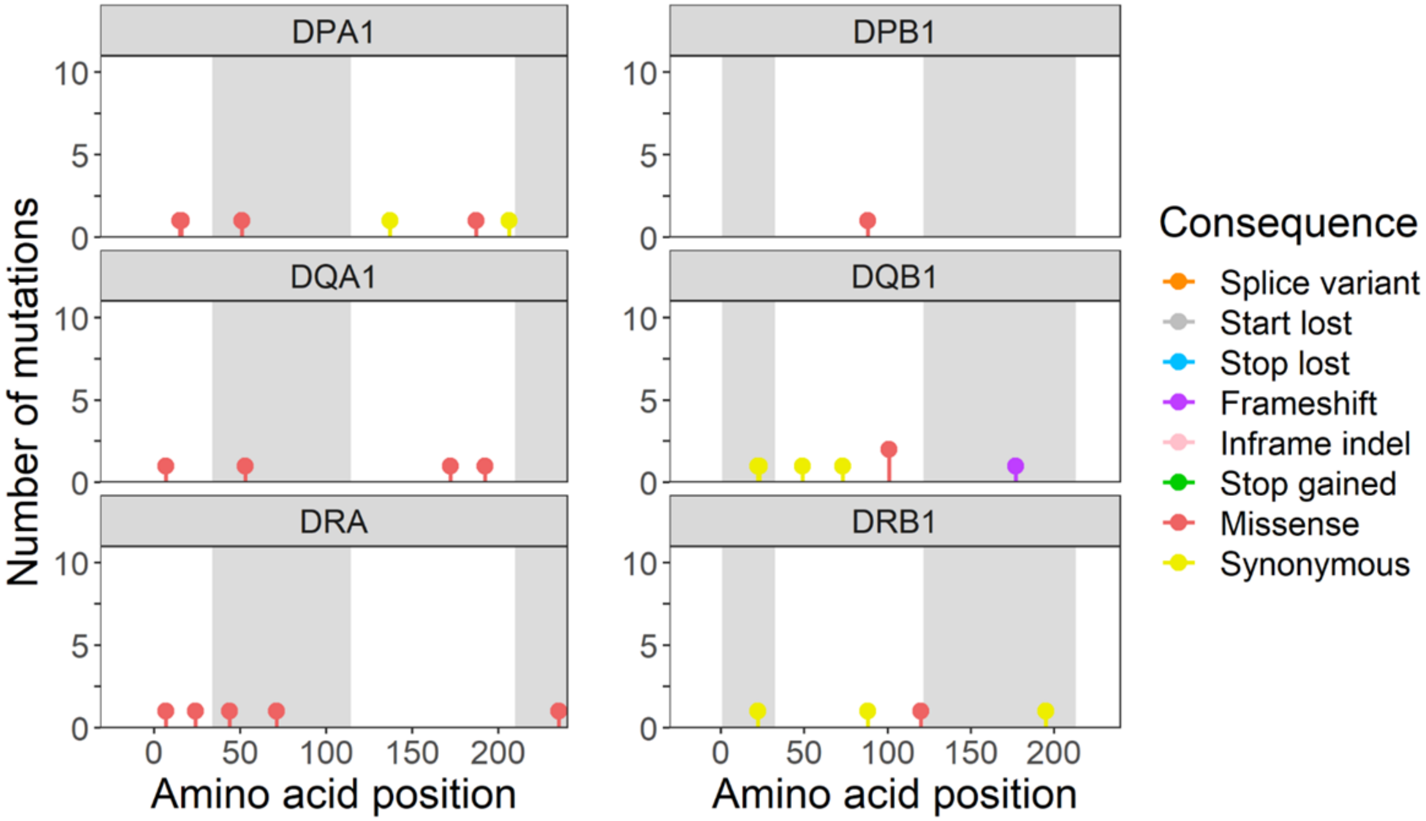

DPA1
DPB1
DQA1
DQB1
DRA
DRB1
Number of mutations
Amino acid position
Amino acid position
Consequence
Splice variant
Start lost
Stop lost
Frameshift
Inframe indel
Stop gained
Missense
Synonymous

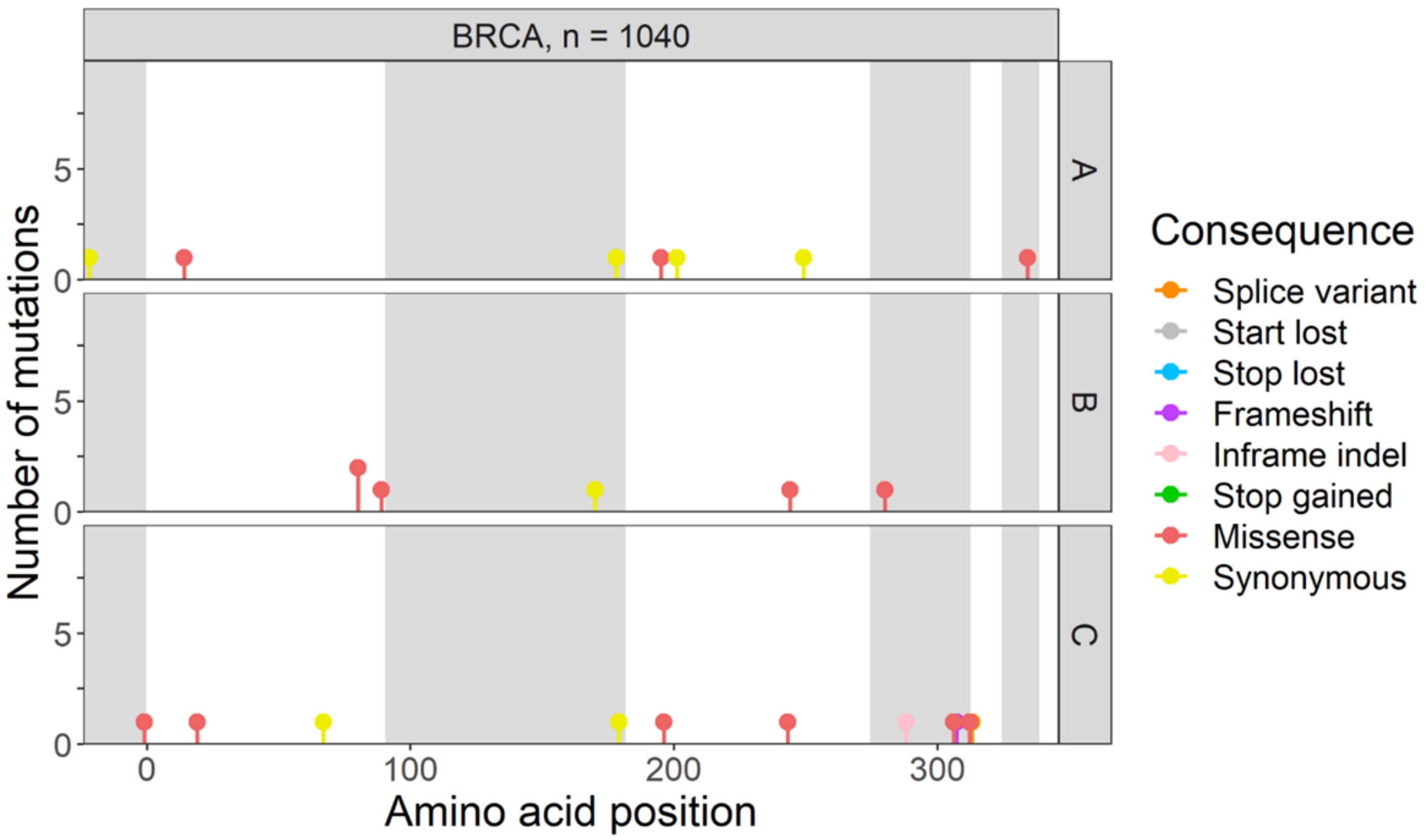

BRCA, n = 1040
A
B
C
Number of mutations
Amino acid position
Consequence
Splice variant
Start lost
Stop lost
Frameshift
Inframe indel
Stop gained
Missense
Synonymous


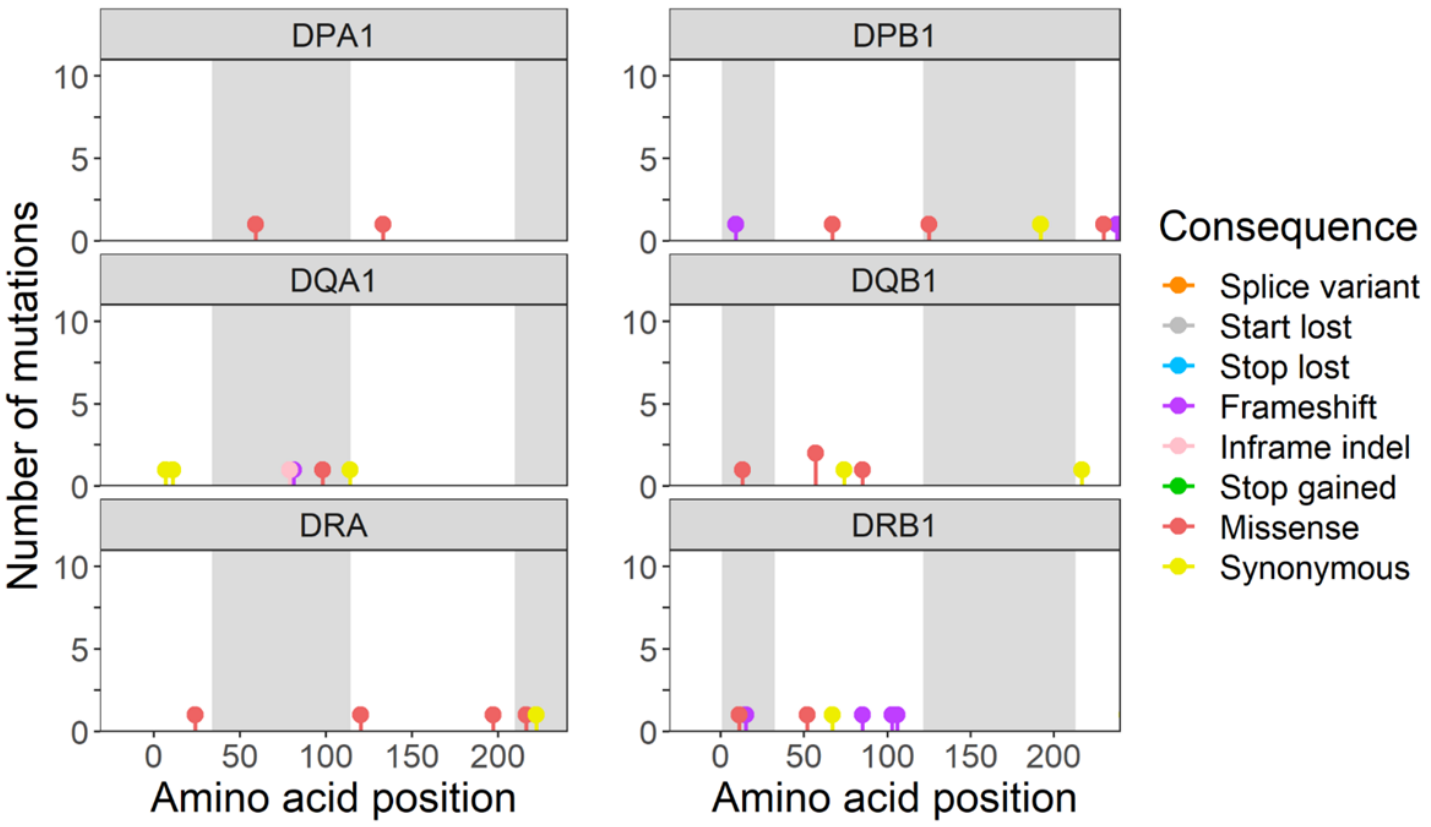

DPA1
DPB1
DQA1
DQB1
DRA
DRB1
Number of mutations
Amino acid position
Amino acid position
Consequence
Splice variant
Start lost
Stop lost
Frameshift
Inframe indel
Stop gained
Missense
Synonymous

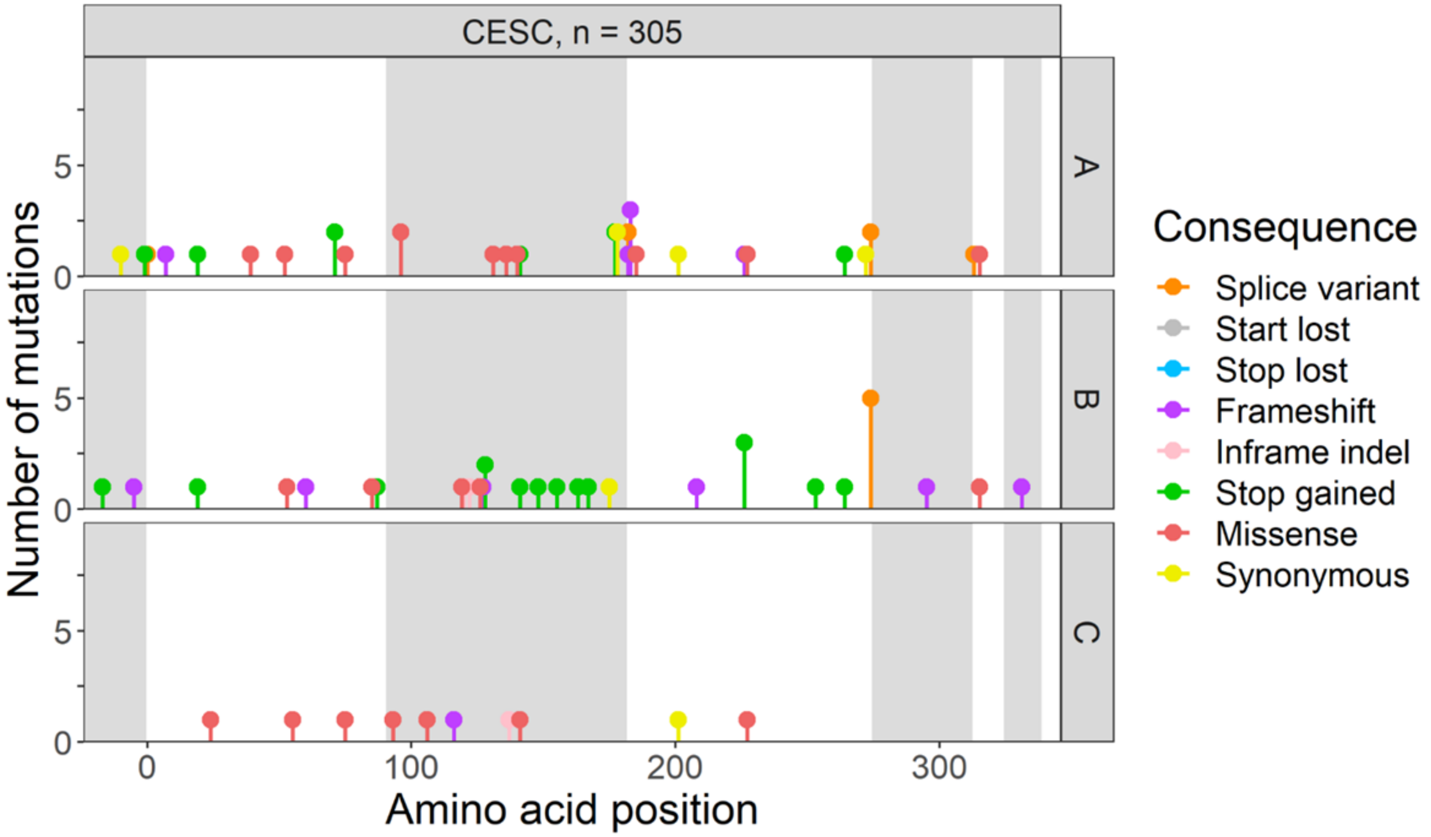

CESC, n = 305
A
B
C
Number of mutations
Amino acid position
Consequence
Splice variant
Start lost
Stop lost
Frameshift
Inframe indel
Stop gained
Missense
Synonymous


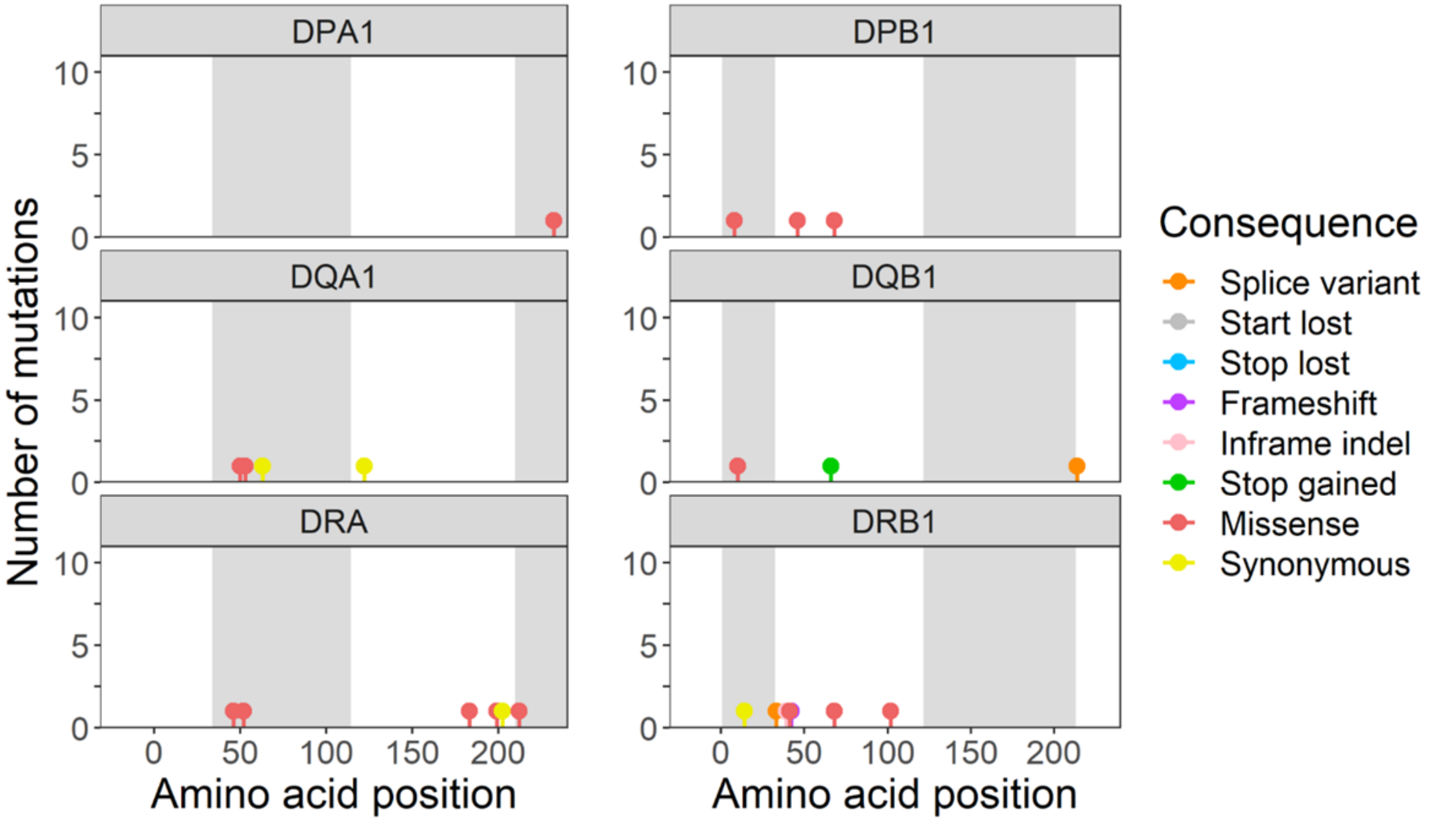

DPA1
DPB1
DQA1
DQB1
DRA
DRB1
Number of mutations
Amino acid position
Amino acid position
Consequence
Splice variant
Start lost
Stop lost
Frameshift
Inframe indel
Stop gained
Missense
Synonymous

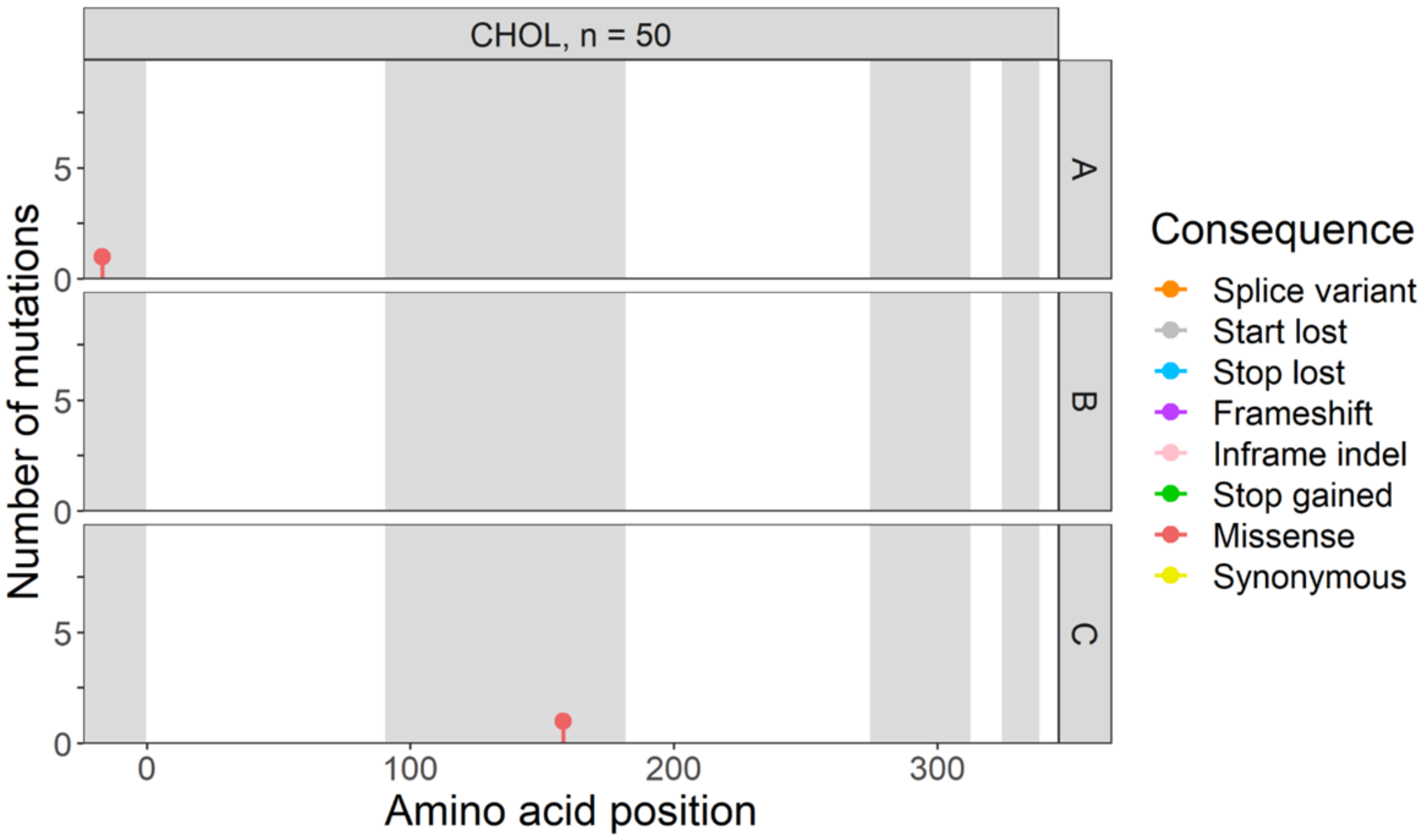
CHOL, n = 50
A
B
C
Number of mutations
0
5
0
100
200
300
Amino acid position
Consequence
Splice variant
Start lost
Stop lost
Frameshift
Inframe indel
Stop gained
Missense
Synonymous

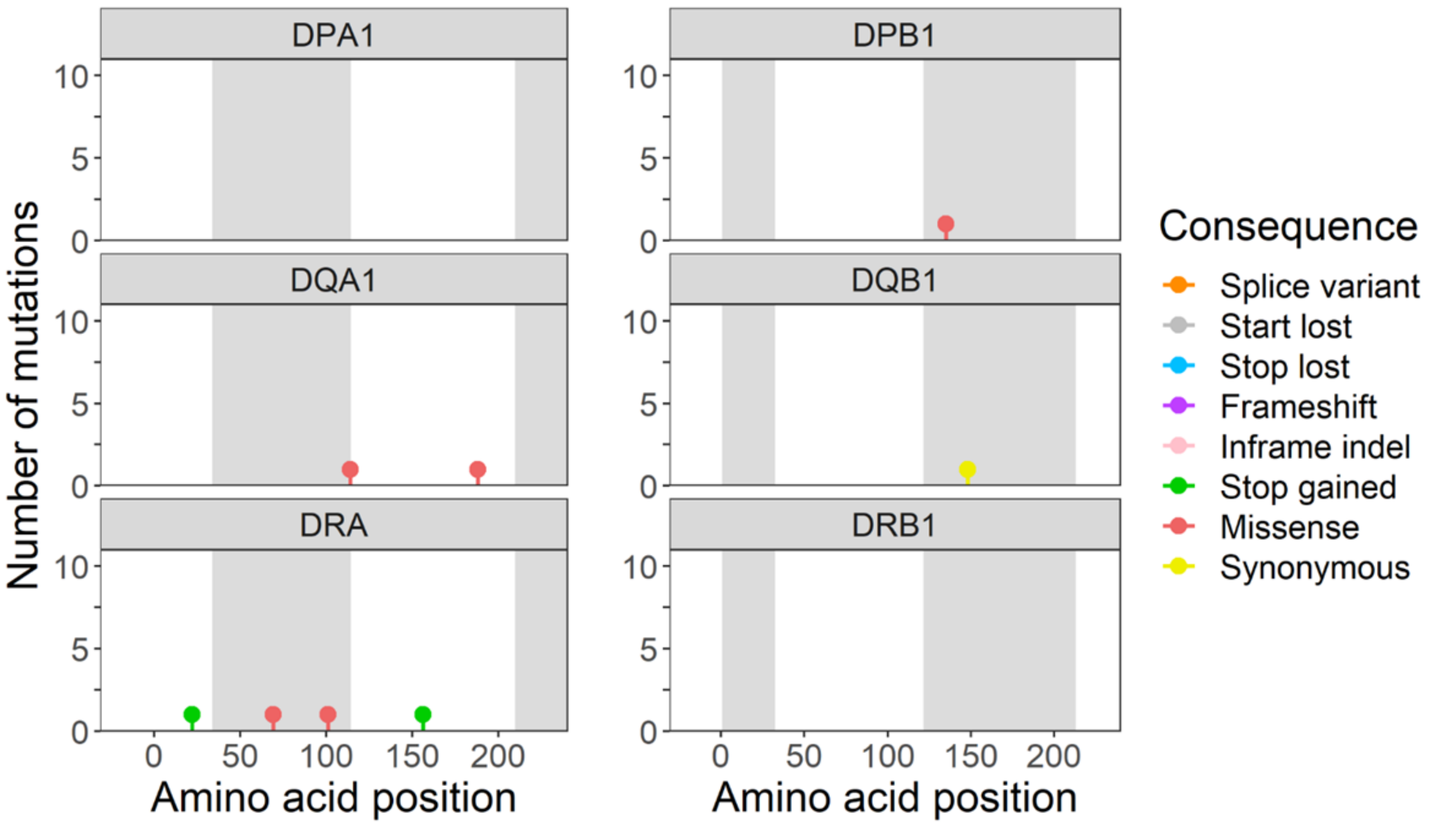
DPA1
DPB1
DQA1
DQB1
DRA
DRB1
Number of mutations
0
5
10
0
50
100
150
200
Amino acid position
Amino acid position
Consequence
Splice variant
Start lost
Stop lost
Frameshift
Inframe indel
Stop gained
Missense
Synonymous

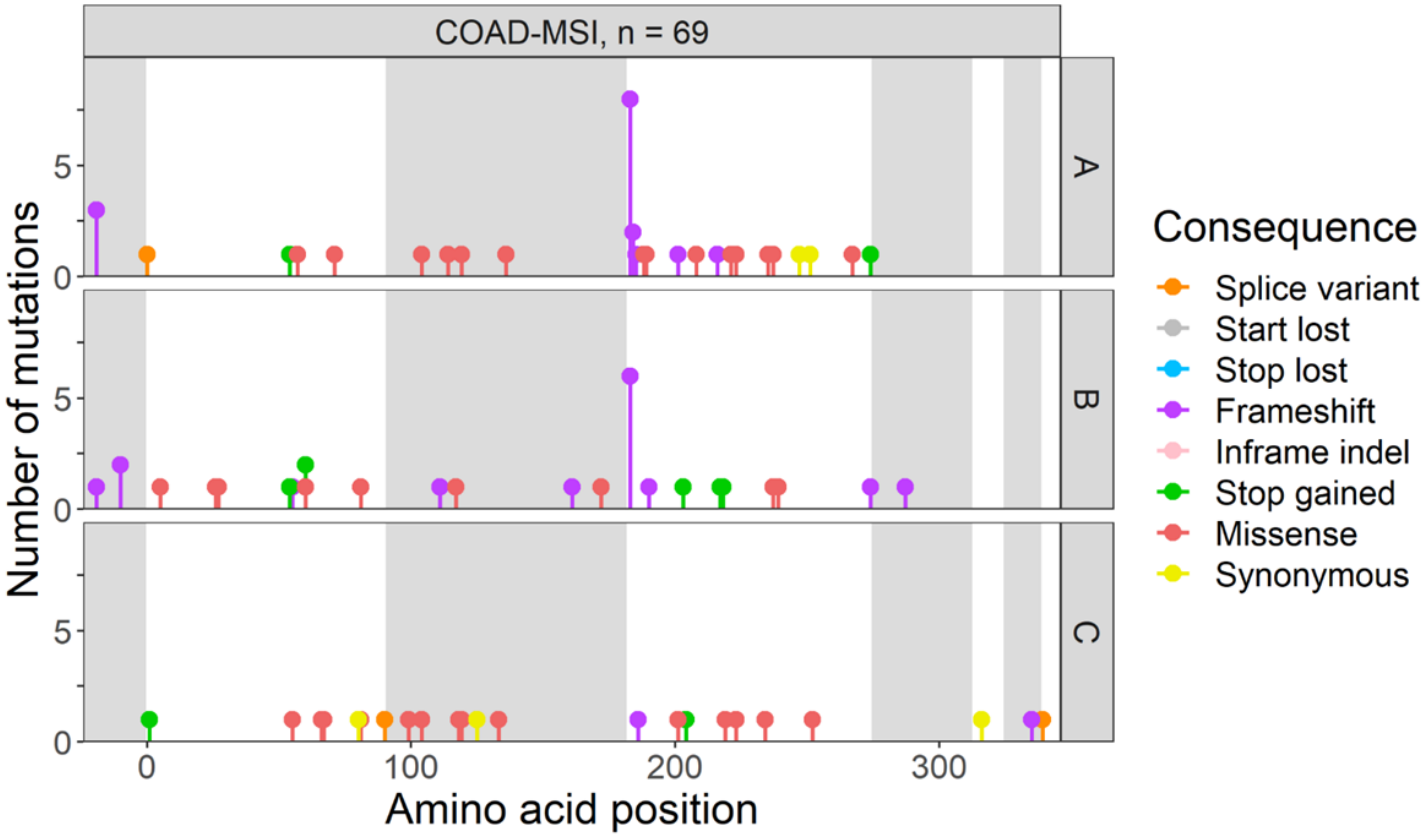
COAD-MSI, n = 69
A
B
C
Number of mutations
Amino acid position
Consequence
Splice variant
Start lost
Stop lost
Frameshift
Inframe indel
Stop gained
Missense
Synonymous

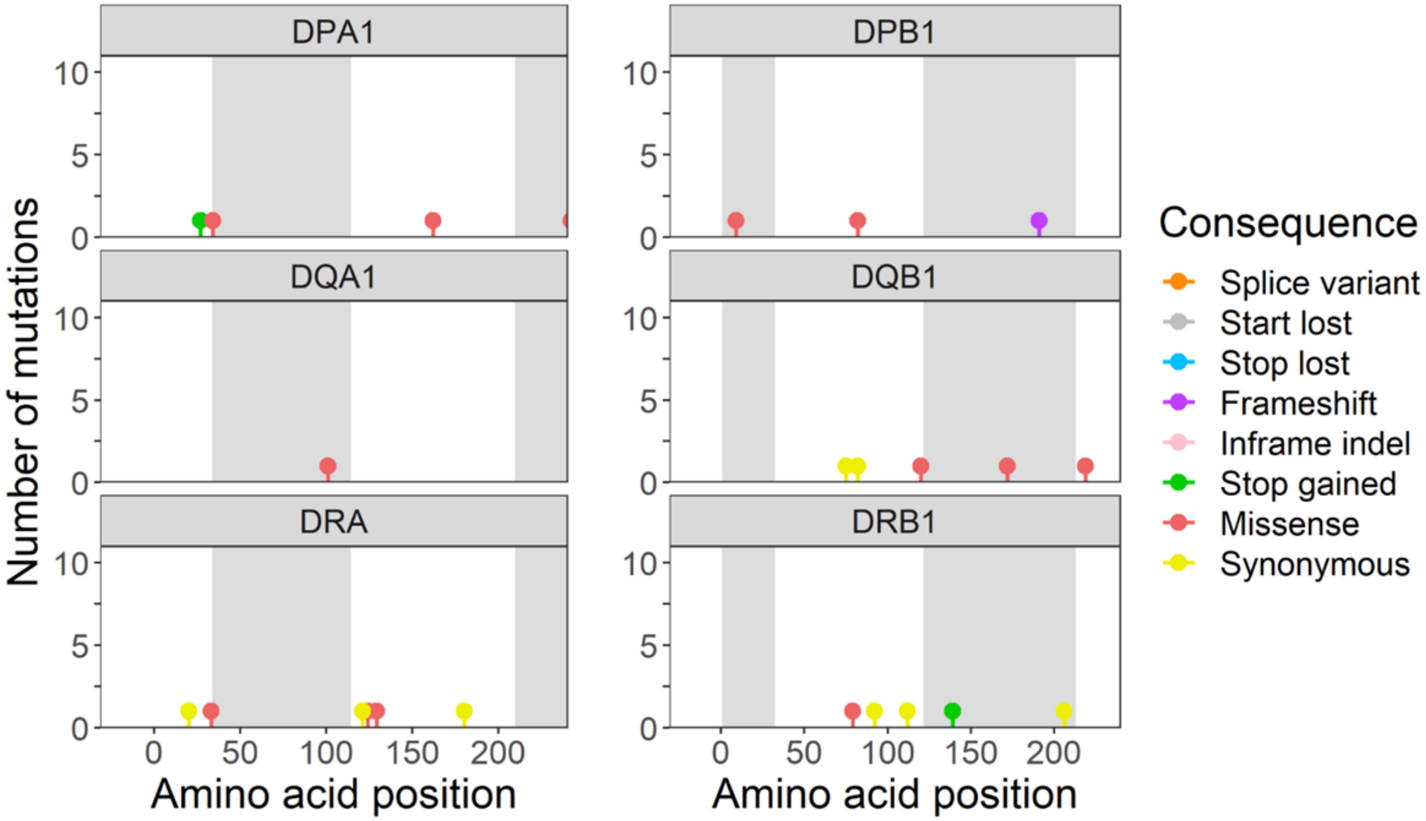
DPA1
DPB1
DQA1
DQB1
DRA
DRB1
Number of mutations
Amino acid position
Amino acid position
Consequence
Splice variant
Start lost
Stop lost
Frameshift
Inframe indel
Stop gained
Missense
Synonymous

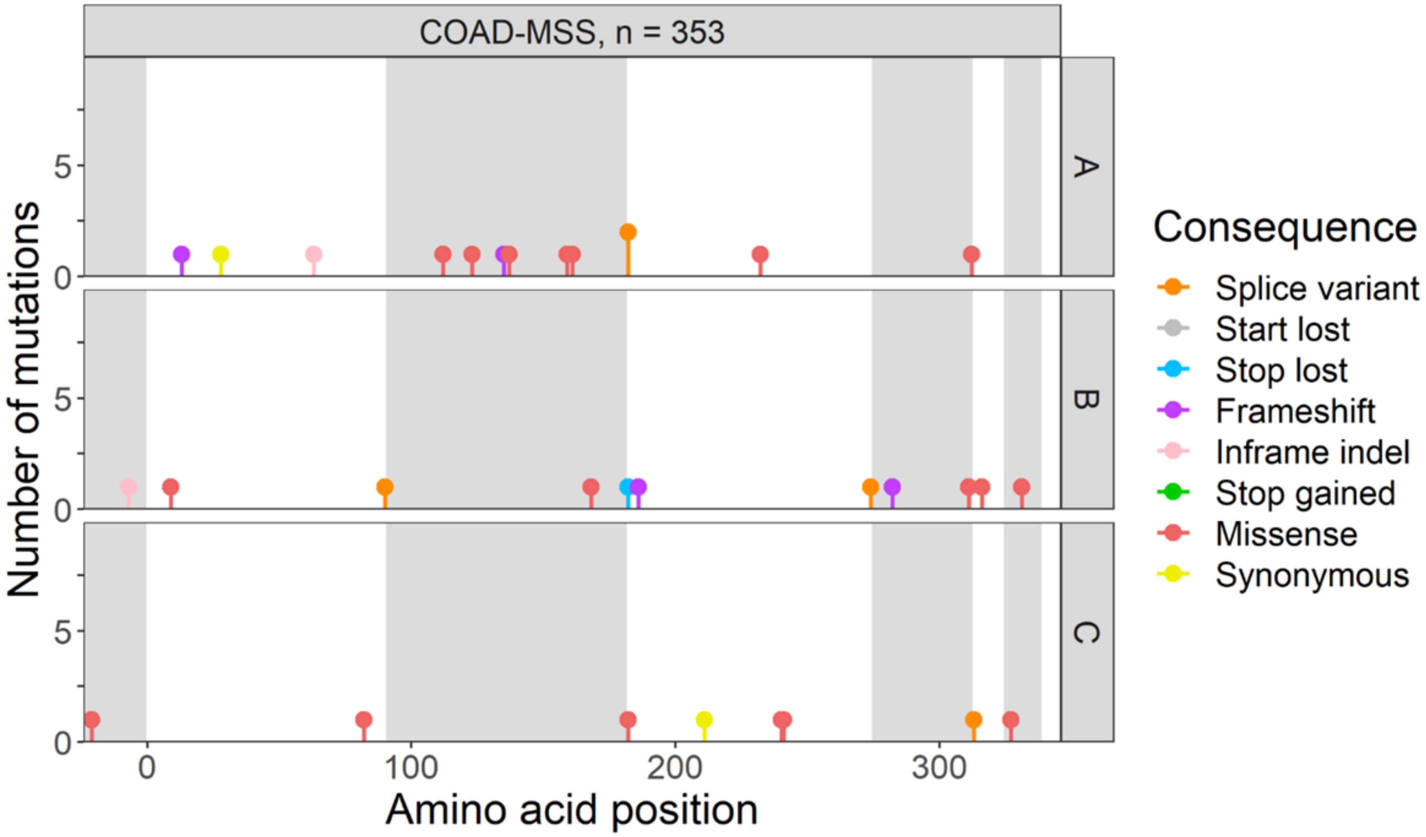
COAD-MSS, n = 353
A
B
C
Number of mutations
Amino acid position
Consequence
Splice variant
Start lost
Stop lost
Frameshift
Inframe indel
Stop gained
Missense
Synonymous

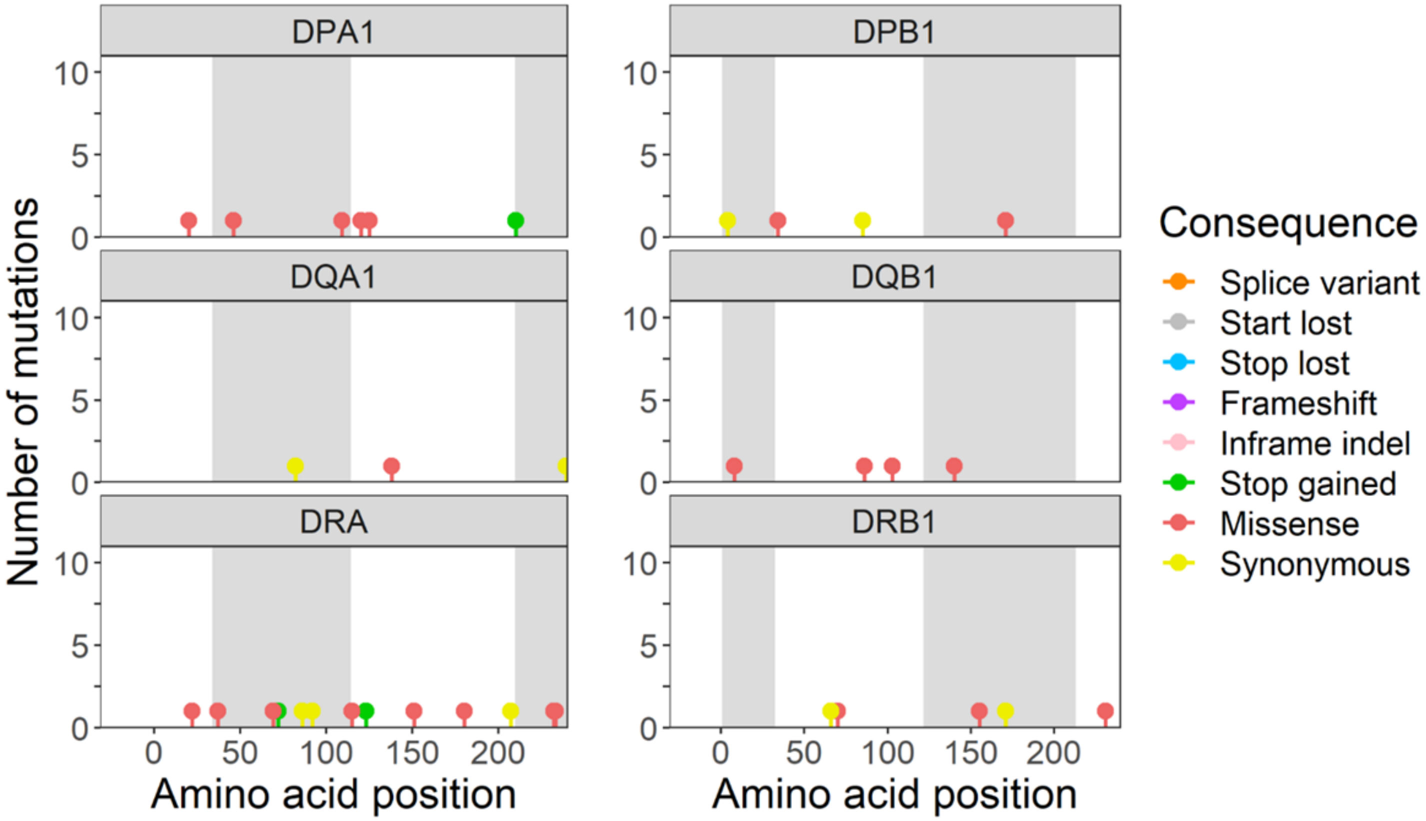
DPA1
DPB1
DQA1
DQB1
DRA
DRB1
Number of mutations
Amino acid position
Amino acid position
Consequence
Splice variant
Start lost
Stop lost
Frameshift
Inframe indel
Stop gained
Missense
Synonymous

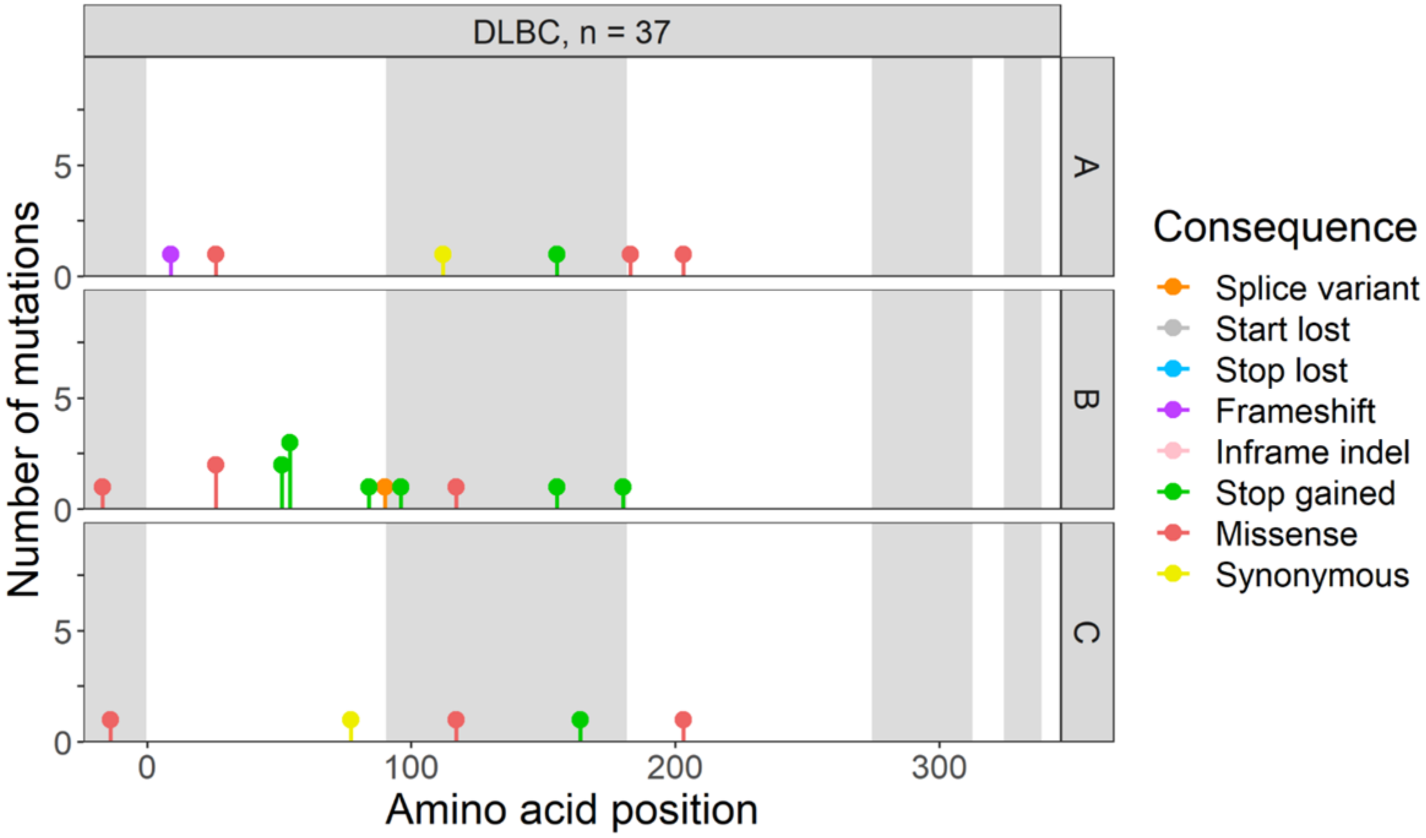
DLBC, n = 37
A
B
C
Number of mutations
Amino acid position
Consequence
Splice variant
Start lost
Stop lost
Frameshift
Inframe indel
Stop gained
Missense
Synonymous

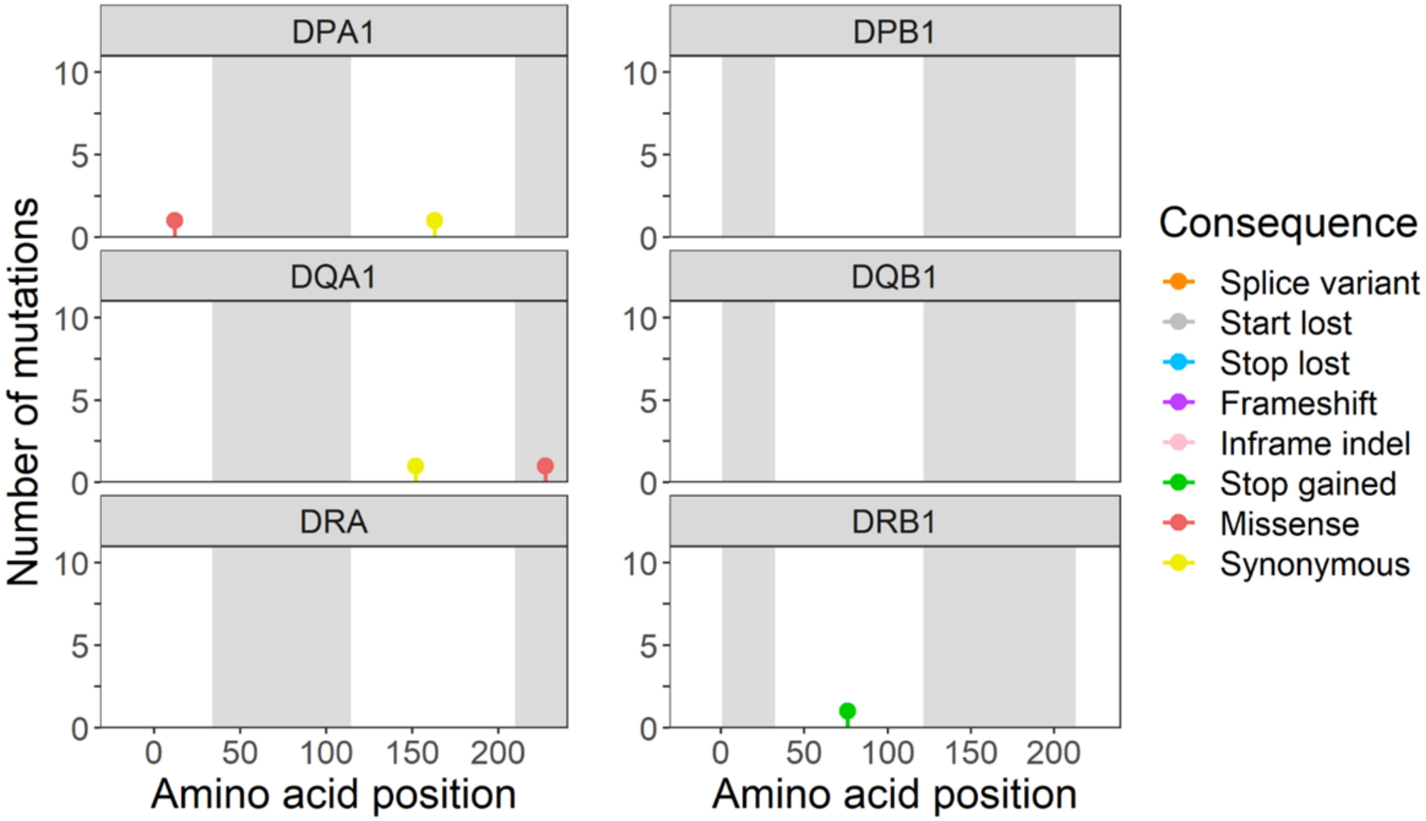
DPA1
DPB1
DQA1
DQB1
DRA
DRB1
Number of mutations
Amino acid position
Consequence
Splice variant
Start lost
Stop lost
Frameshift
Inframe indel
Stop gained
Missense
Synonymous

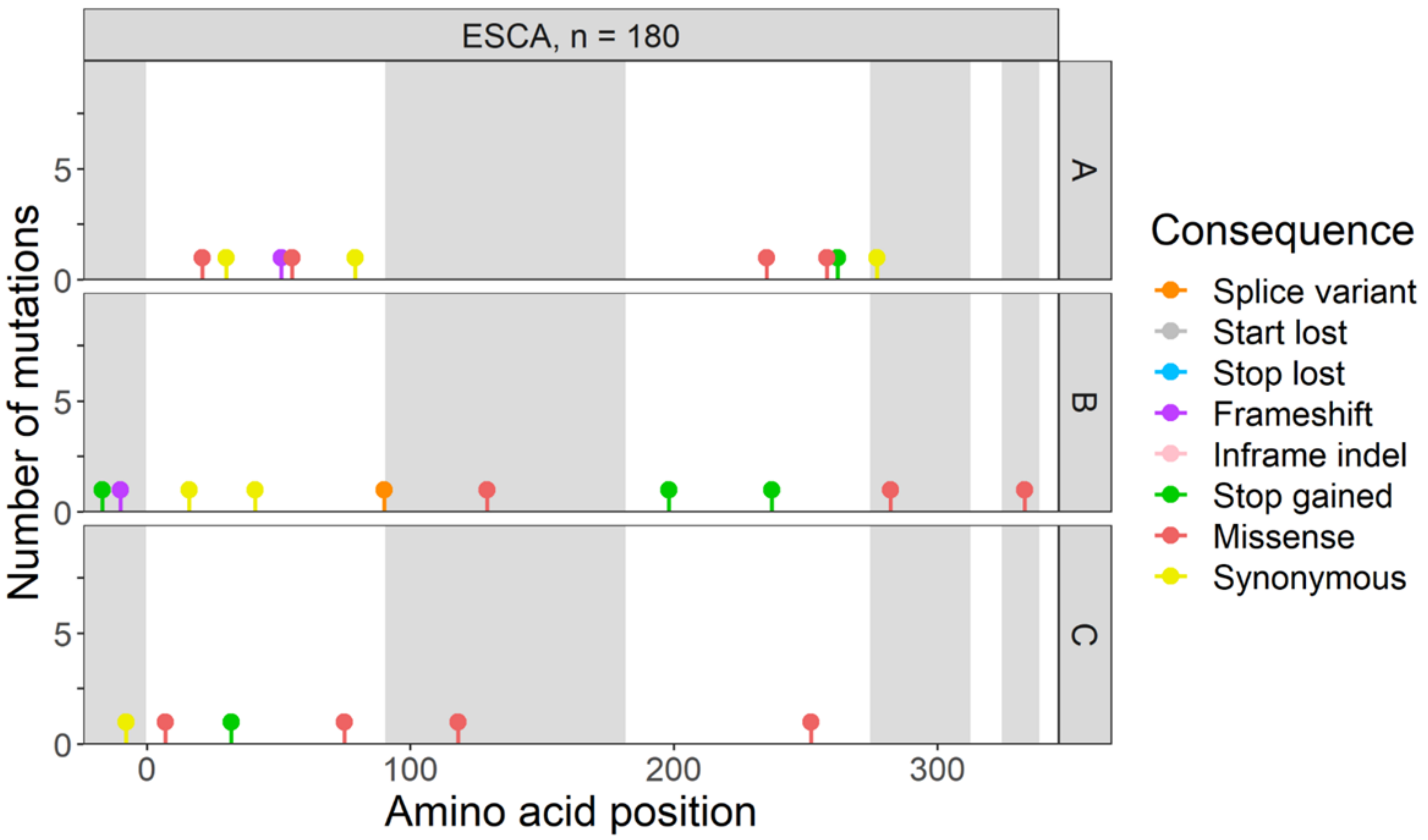
ESCA, n = 180
A
B
C
Number of mutations
0
5
0
100
200
300
Amino acid position
Consequence
Splice variant
Start lost
Stop lost
Frameshift
Inframe indel
Stop gained
Missense
Synonymous

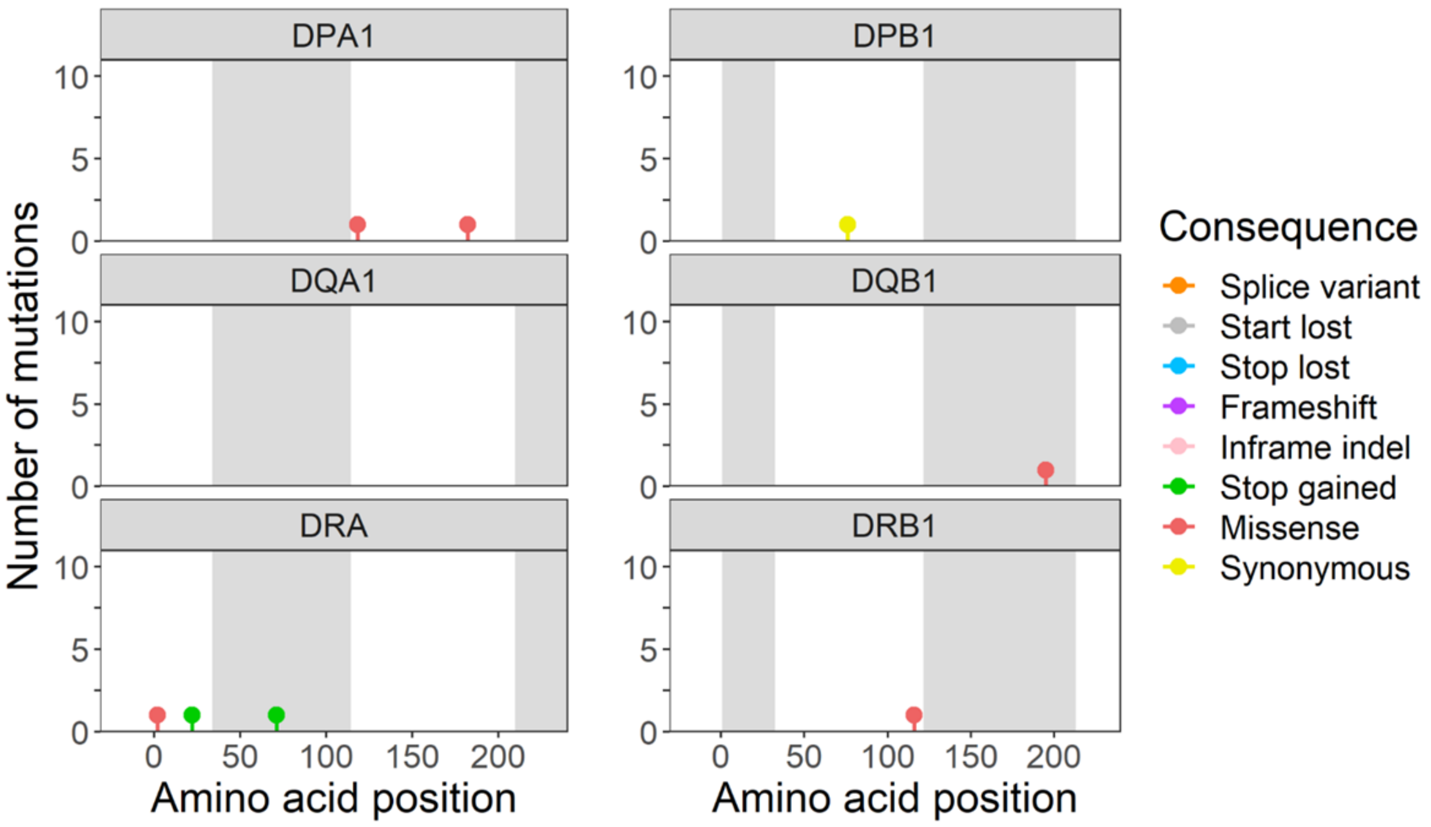
DPA1
DPB1
DQA1
DQB1
DRA
DRB1
Number of mutations
0
5
10
0
50
100
150
200
Amino acid position
Amino acid position
Consequence
Splice variant
Start lost
Stop lost
Frameshift
Inframe indel
Stop gained
Missense
Synonymous

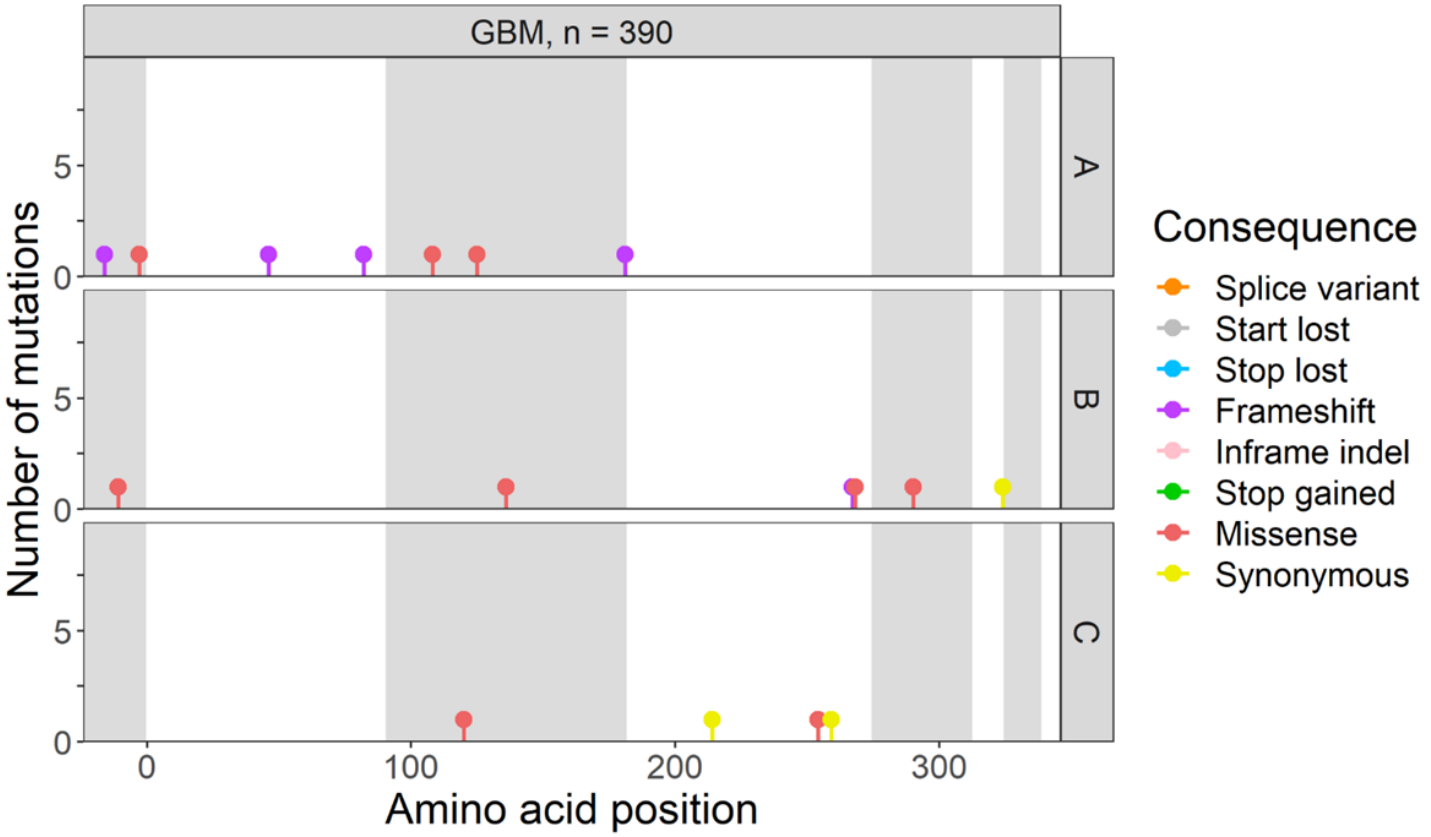
GBM, n = 390
A
B
C
Number of mutations
Amino acid position
Consequence
Splice variant
Start lost
Stop lost
Frameshift
Inframe indel
Stop gained
Missense
Synonymous

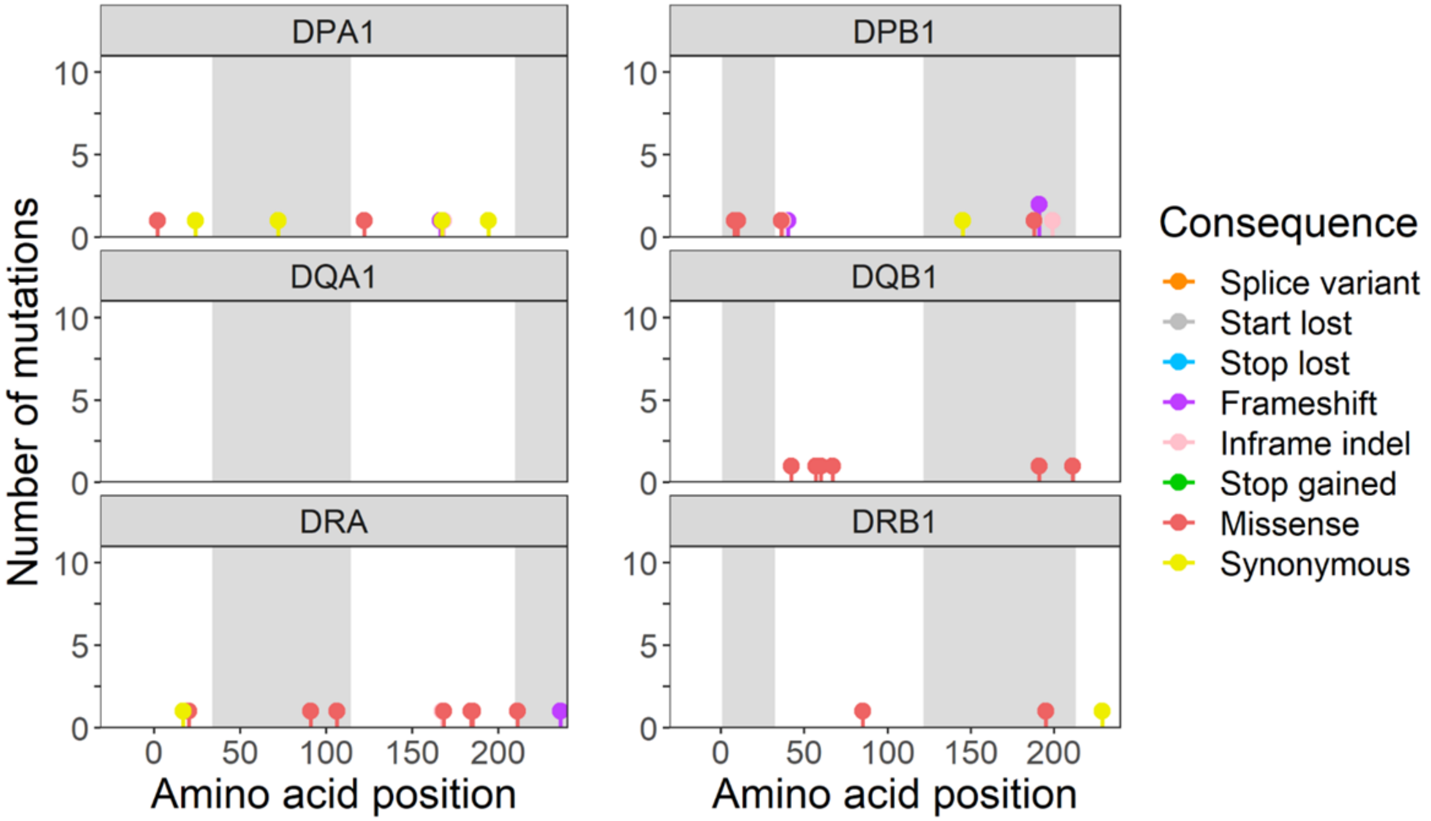
DPA1
DPB1
DQA1
DQB1
DRA
DRB1
Number of mutations
Amino acid position
Amino acid position
Consequence
Splice variant
Start lost
Stop lost
Frameshift
Inframe indel
Stop gained
Missense
Synonymous

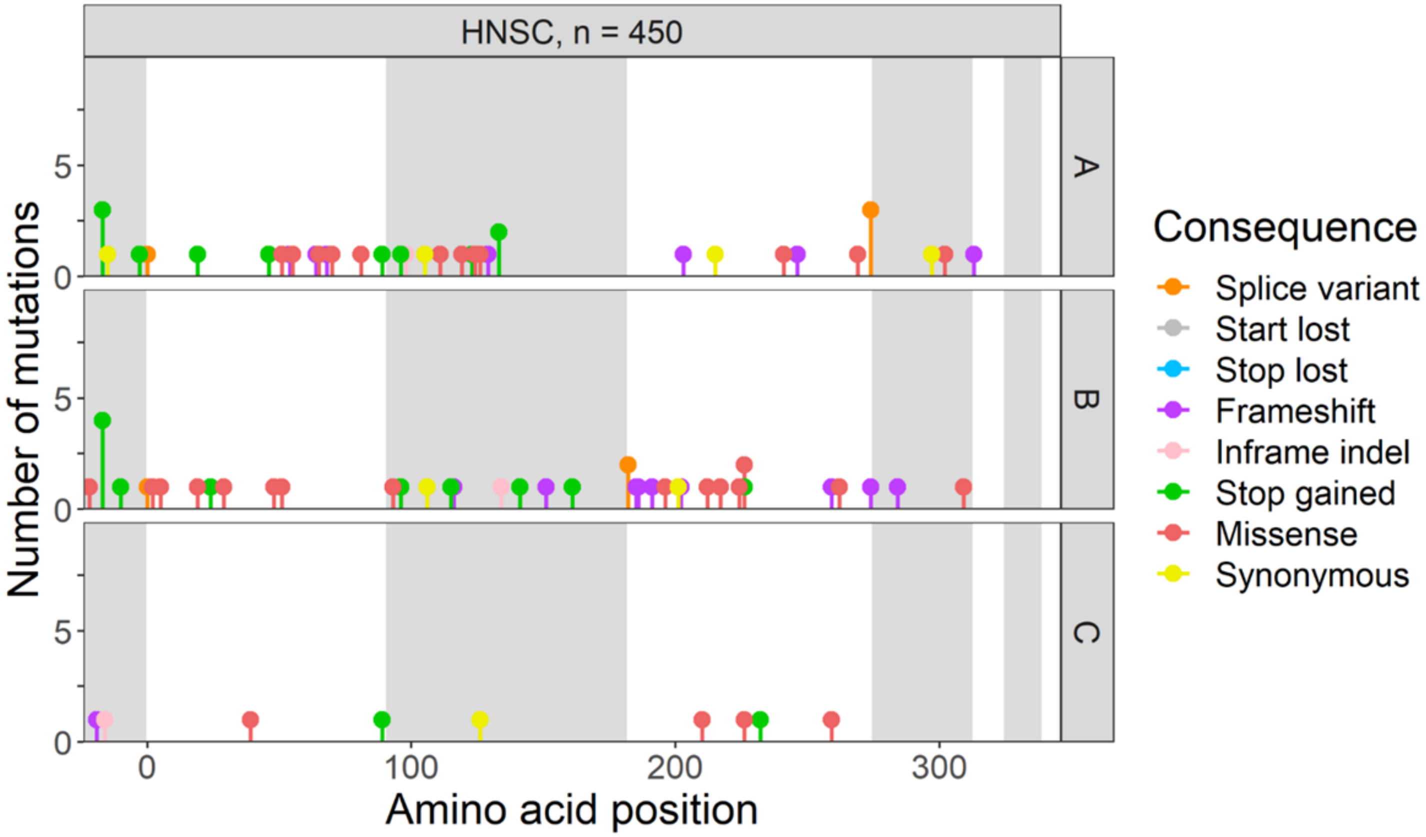

HNSC, n = 450
A
B
C
Number of mutations
Amino acid position
Consequence
Splice variant
Start lost
Stop lost
Frameshift
Inframe indel
Stop gained
Missense
Synonymous


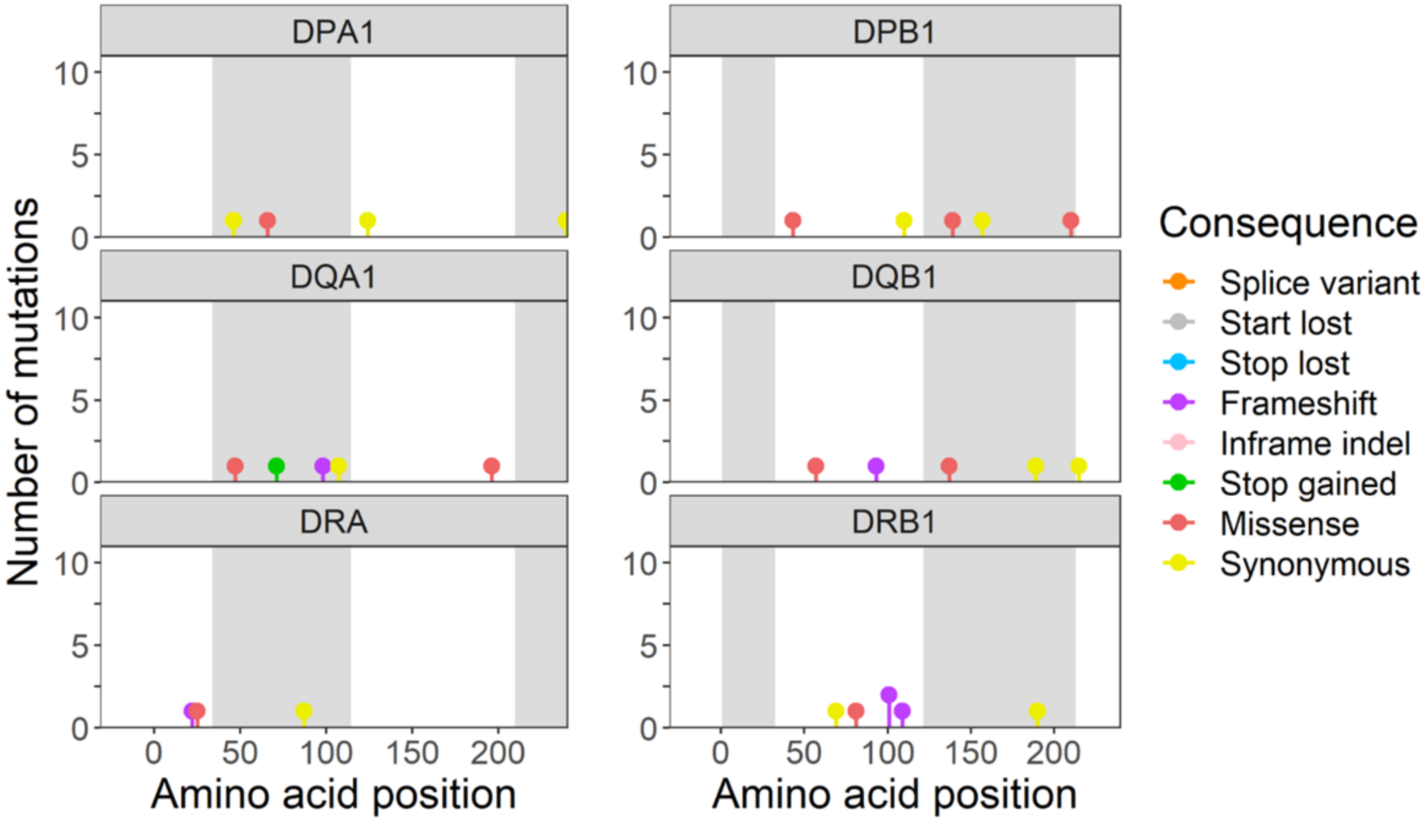

DPA1
DPB1
DQA1
DQB1
DRA
DRB1
Number of mutations
Amino acid position
Amino acid position
Consequence
Splice variant
Start lost
Stop lost
Frameshift
Inframe indel
Stop gained
Missense
Synonymous

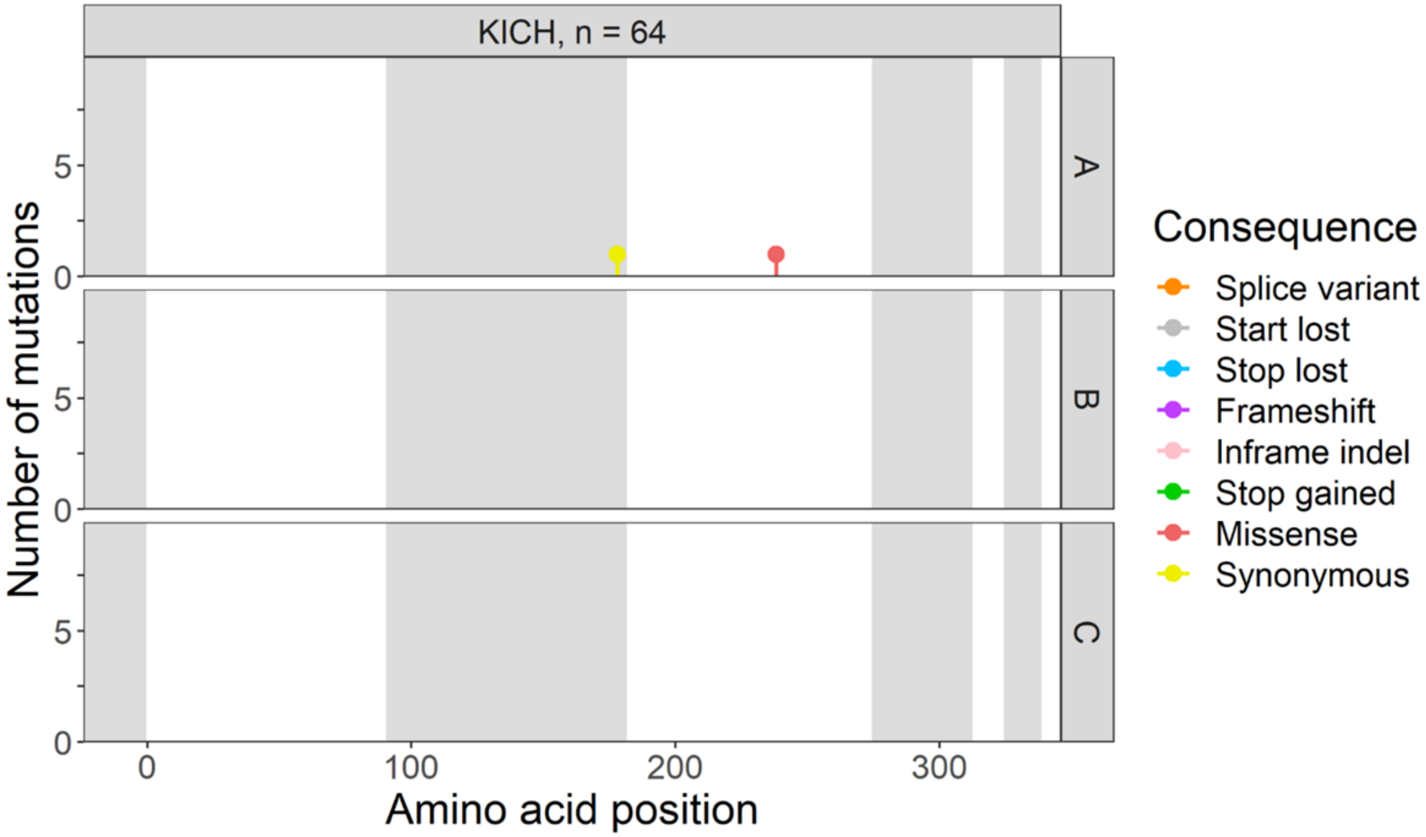
KICH, n = 64
A
B
C
Number of mutations
Amino acid position
Consequence
Splice variant
Start lost
Stop lost
Frameshift
Inframe indel
Stop gained
Missense
Synonymous

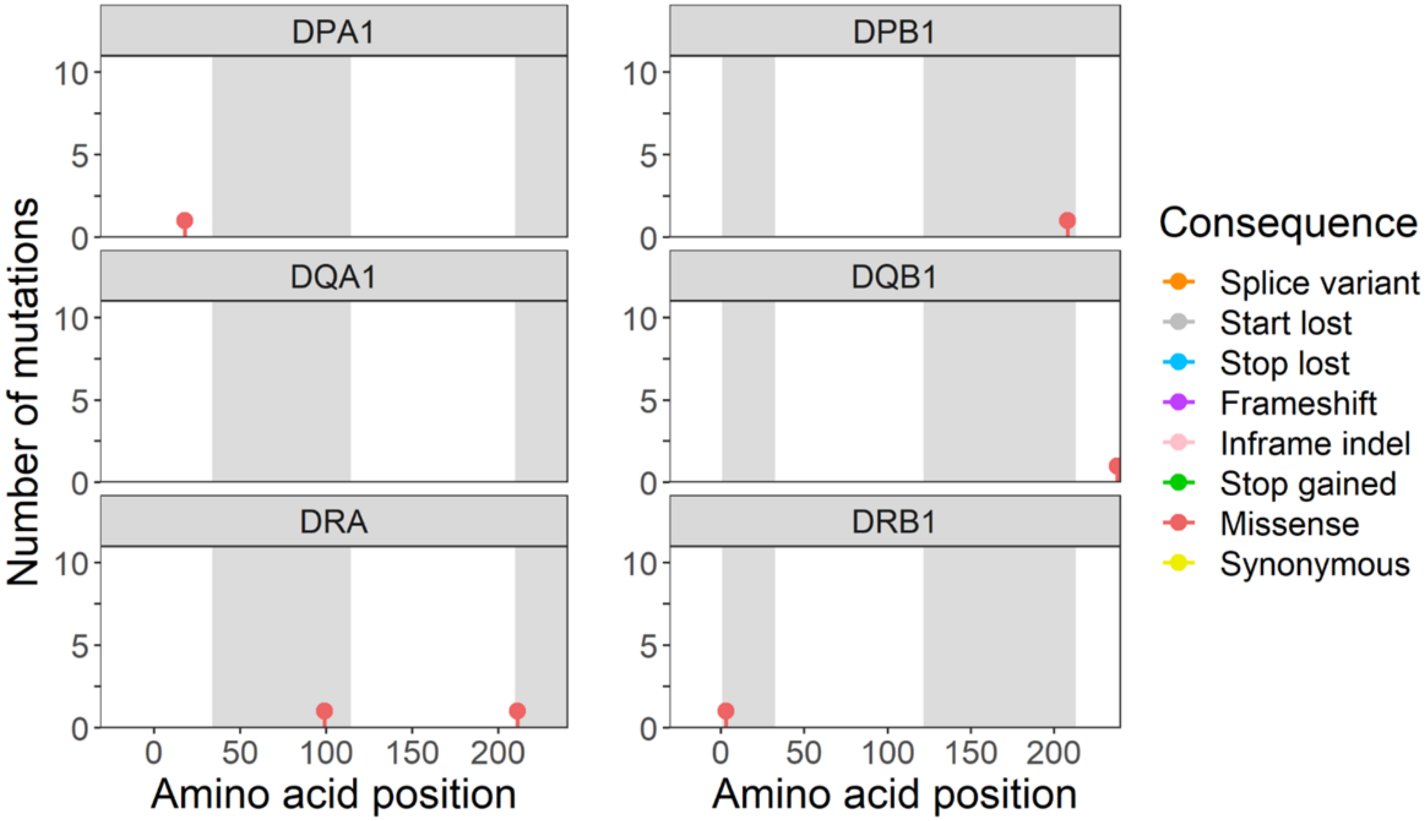
DPA1
DPB1
DQA1
DQB1
DRA
DRB1
Number of mutations
Amino acid position
Amino acid position
Consequence
Splice variant
Start lost
Stop lost
Frameshift
Inframe indel
Stop gained
Missense
Synonymous

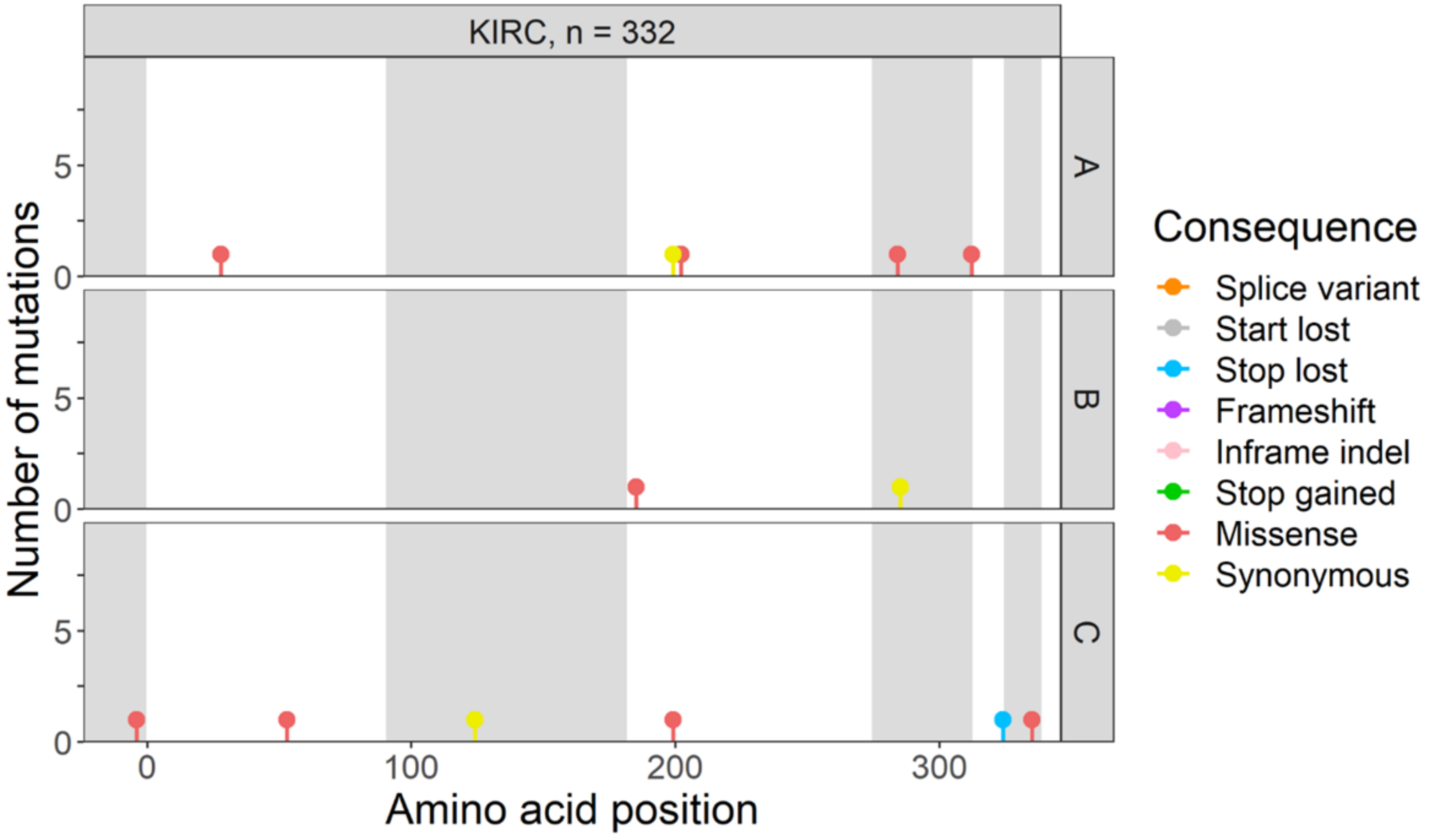

KIRC, n = 332
A
B
C
Number of mutations
Amino acid position
Consequence
Splice variant
Start lost
Stop lost
Frameshift
Inframe indel
Stop gained
Missense
Synonymous


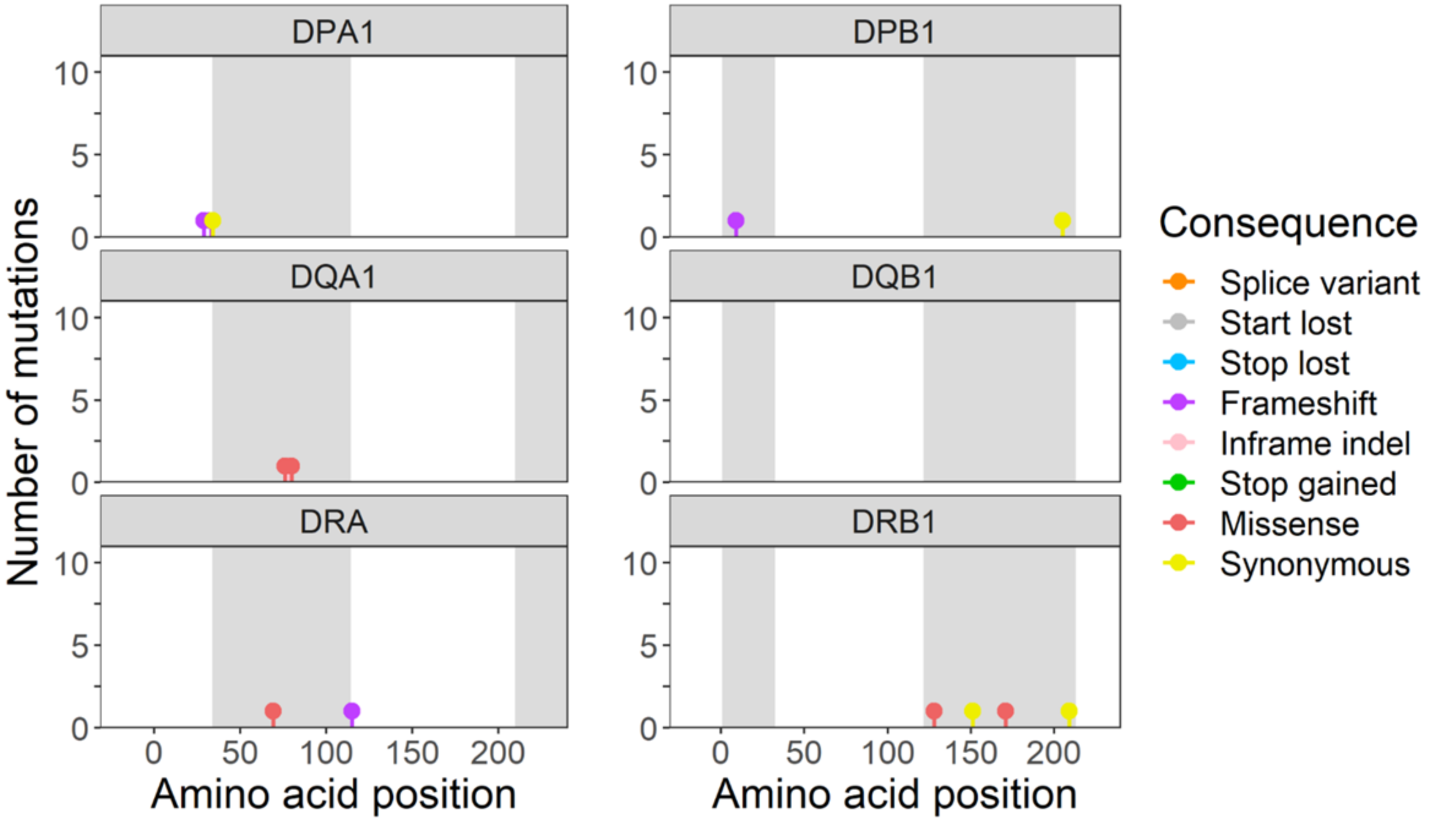

DPA1
DPB1
DQA1
DQB1
DRA
DRB1
Number of mutations
Amino acid position
Amino acid position
Consequence
Splice variant
Start lost
Stop lost
Frameshift
Inframe indel
Stop gained
Missense
Synonymous

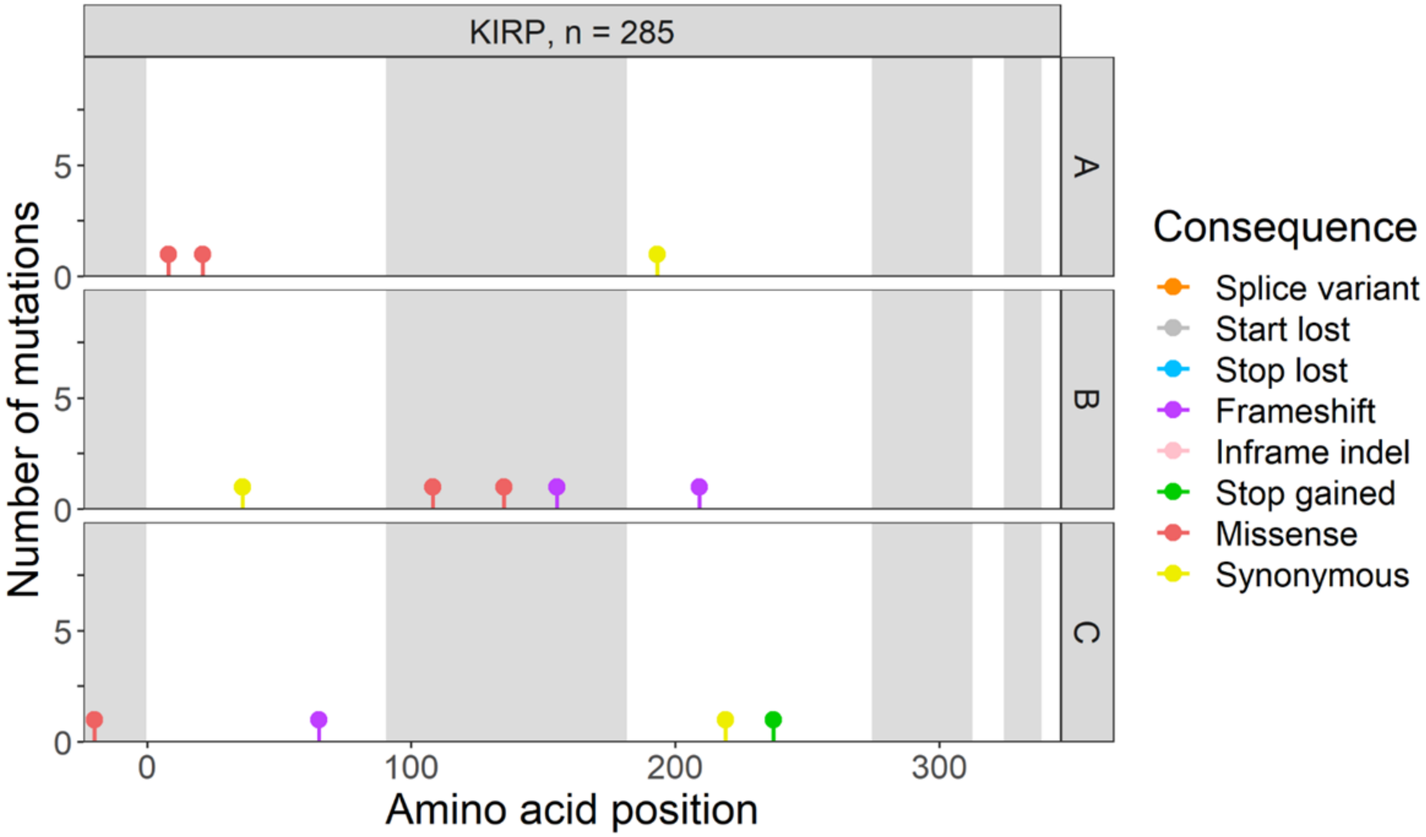

KIRP, n = 285
A
B
C
Number of mutations
Amino acid position
Consequence
Splice variant
Start lost
Stop lost
Frameshift
Inframe indel
Stop gained
Missense
Synonymous


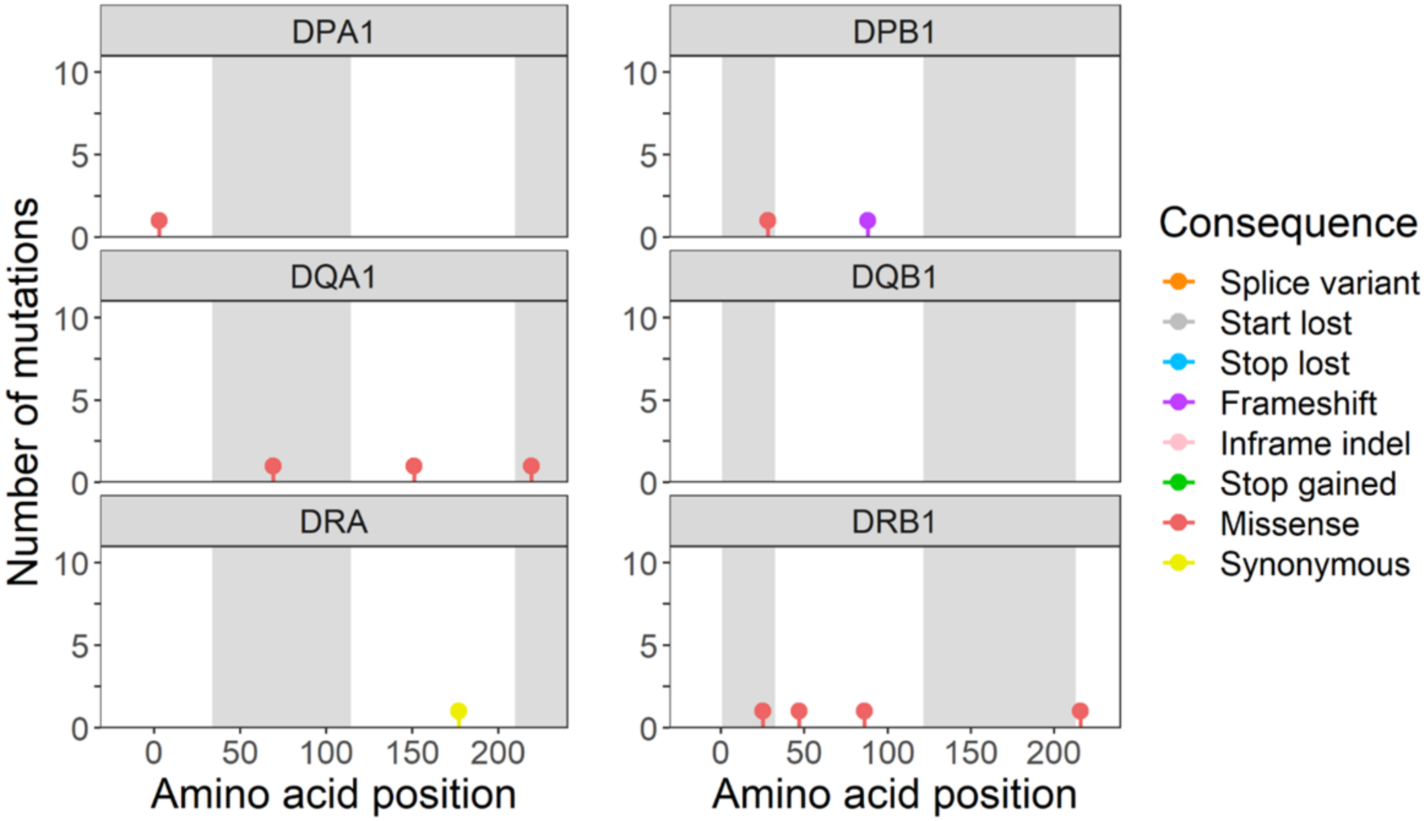

DPA1
DPB1
DQA1
DQB1
DRA
DRB1
Number of mutations
Amino acid position
Amino acid position
Consequence
Splice variant
Start lost
Stop lost
Frameshift
Inframe indel
Stop gained
Missense
Synonymous

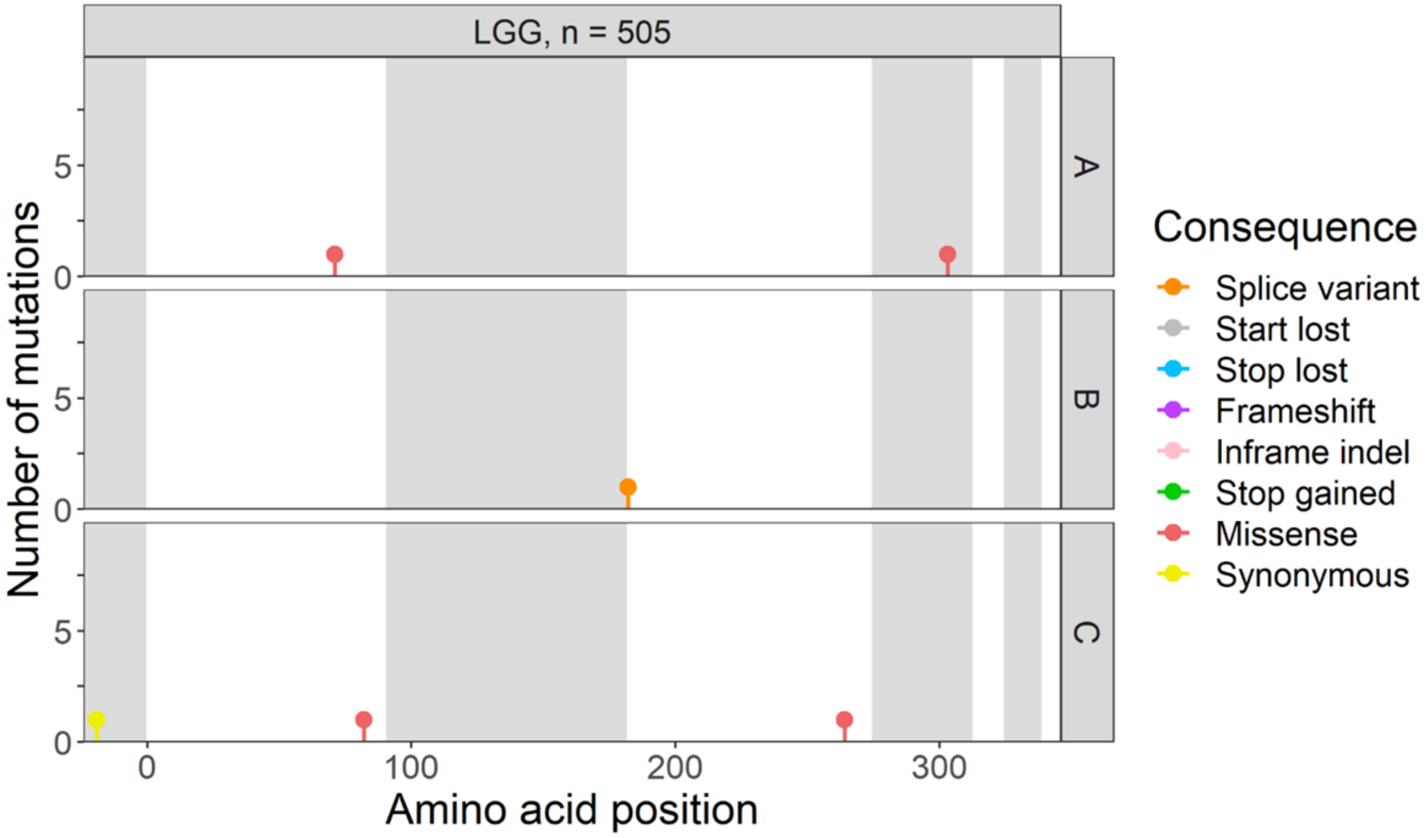

LGG, n = 505
A
B
C
Number of mutations
Amino acid position
Consequence
Splice variant
Start lost
Stop lost
Frameshift
Inframe indel
Stop gained
Missense
Synonymous


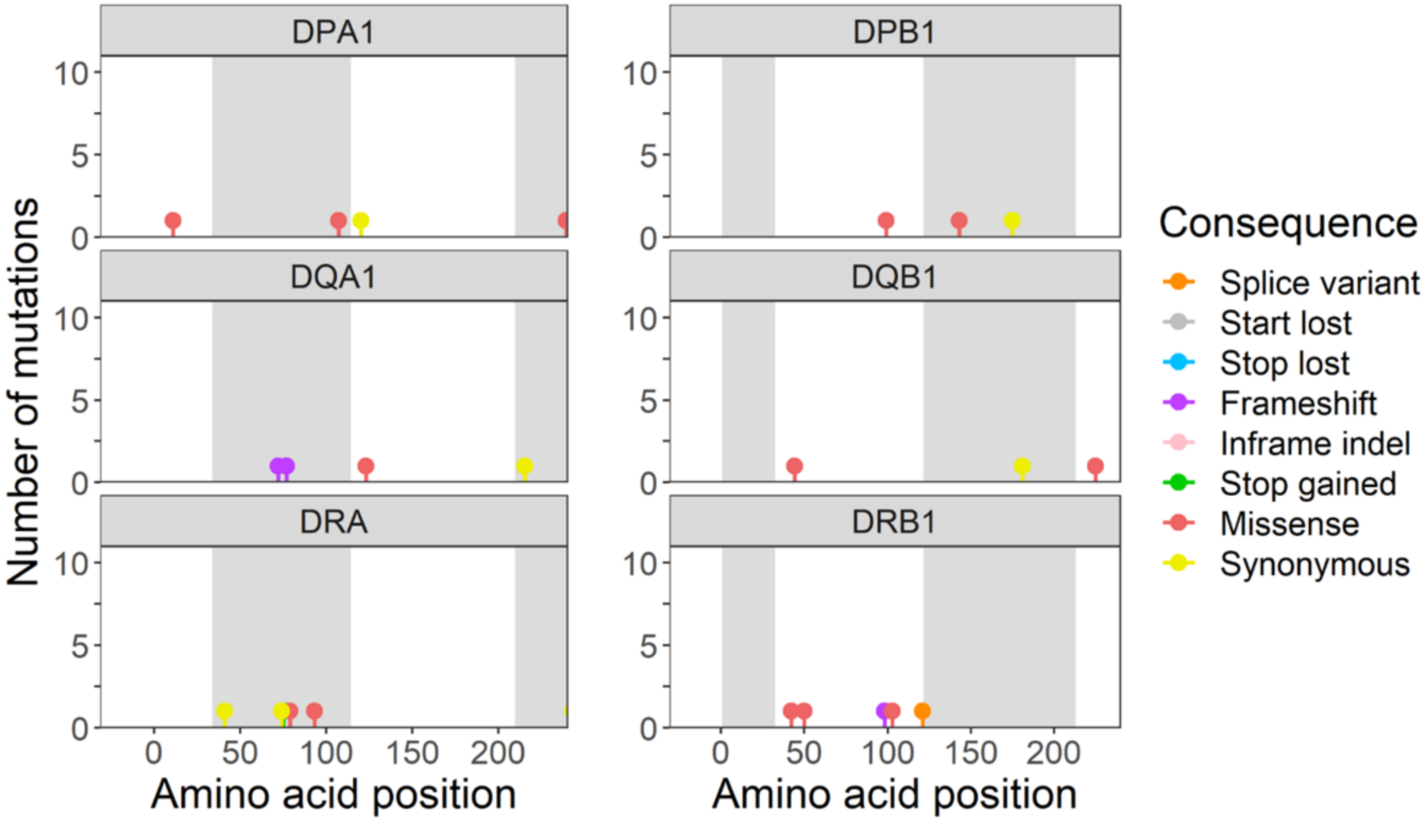

DPA1
DPB1
DQA1
DQB1
DRA
DRB1
Number of mutations
Amino acid position
Amino acid position
Consequence
Splice variant
Start lost
Stop lost
Frameshift
Inframe indel
Stop gained
Missense
Synonymous

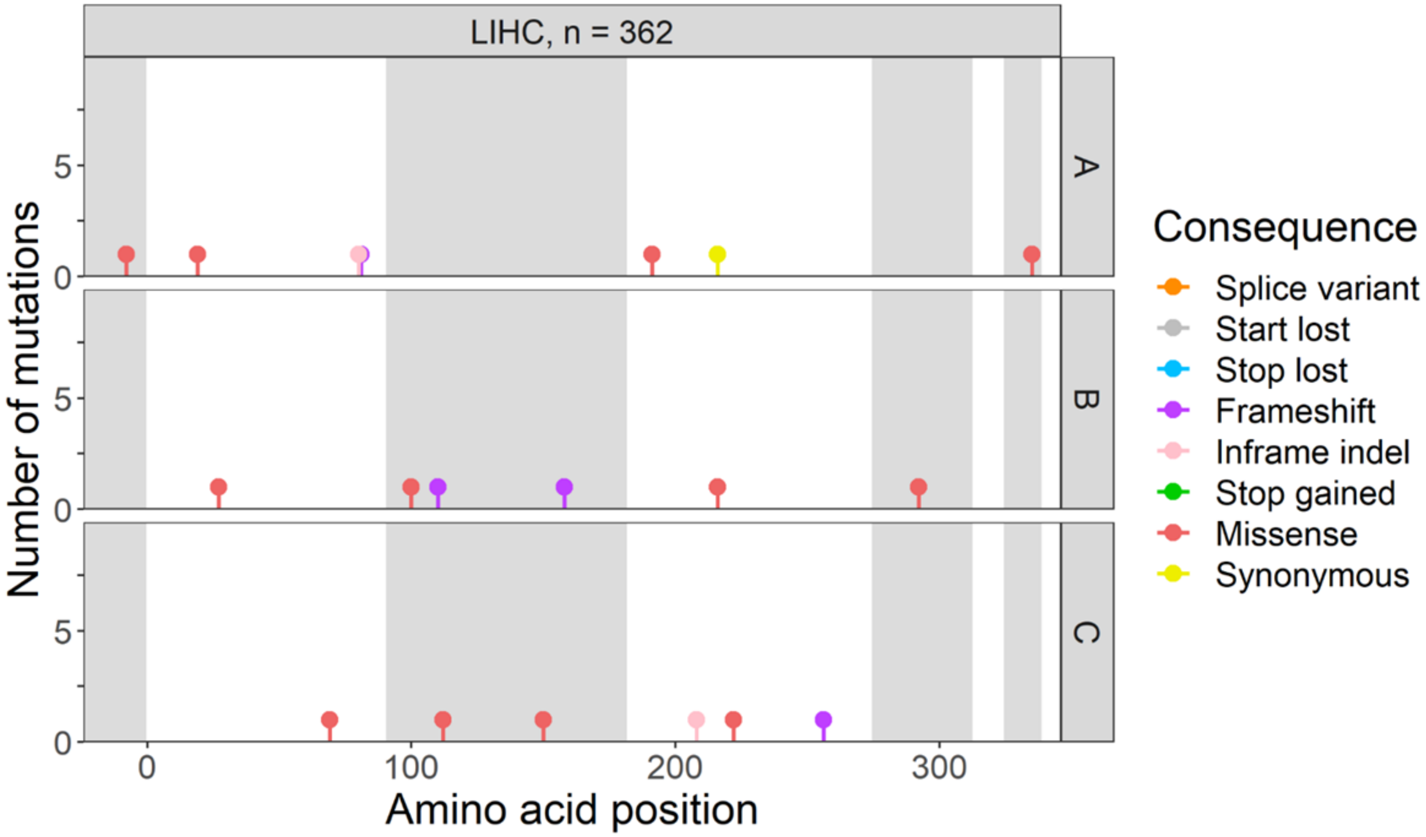

LIHC, n = 362
A
B
C
Number of mutations
Amino acid position
Consequence
Splice variant
Start lost
Stop lost
Frameshift
Inframe indel
Stop gained
Missense
Synonymous


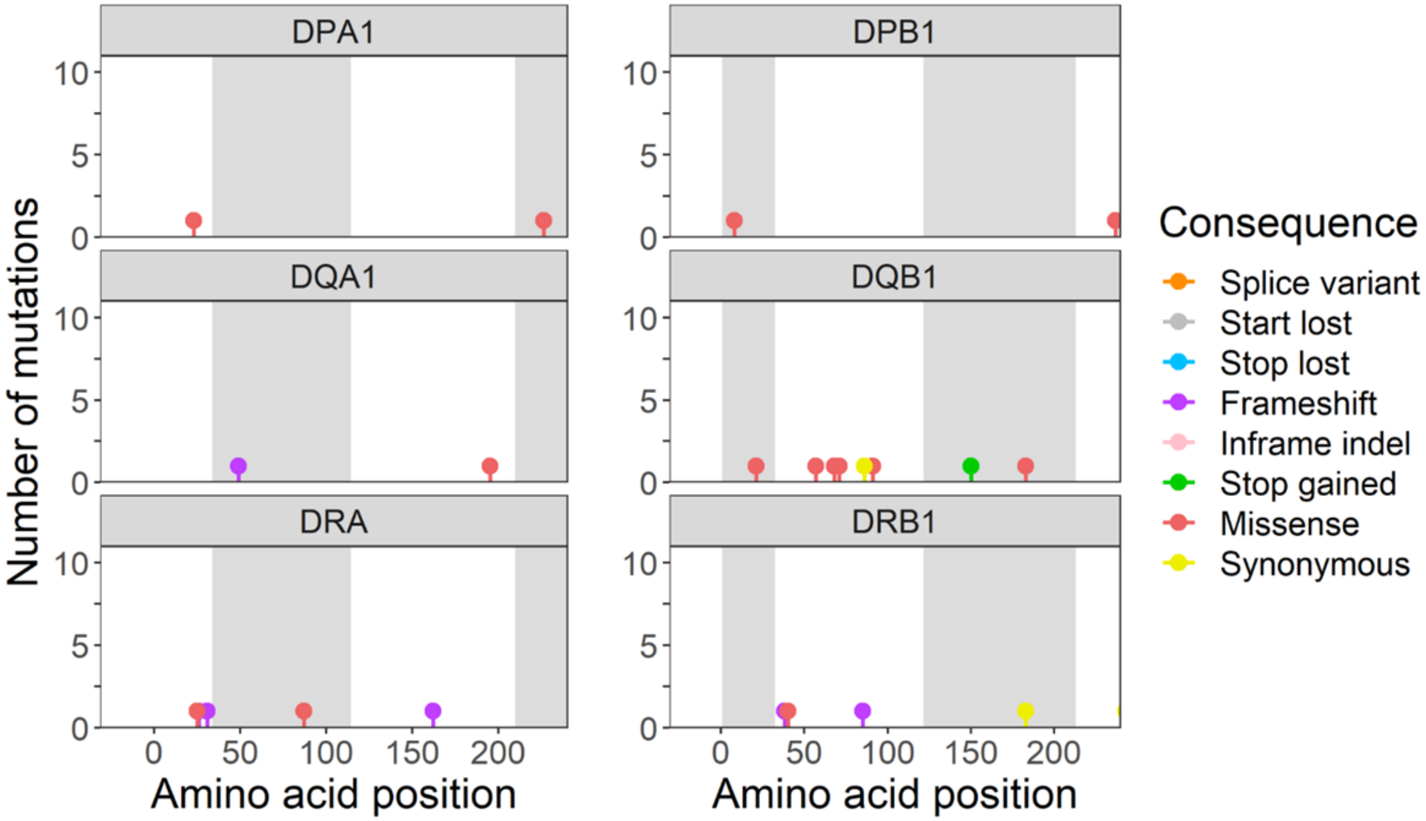

DPA1
DPB1
DQA1
DQB1
DRA
DRB1
Number of mutations
Amino acid position
Amino acid position
Consequence
Splice variant
Start lost
Stop lost
Frameshift
Inframe indel
Stop gained
Missense
Synonymous

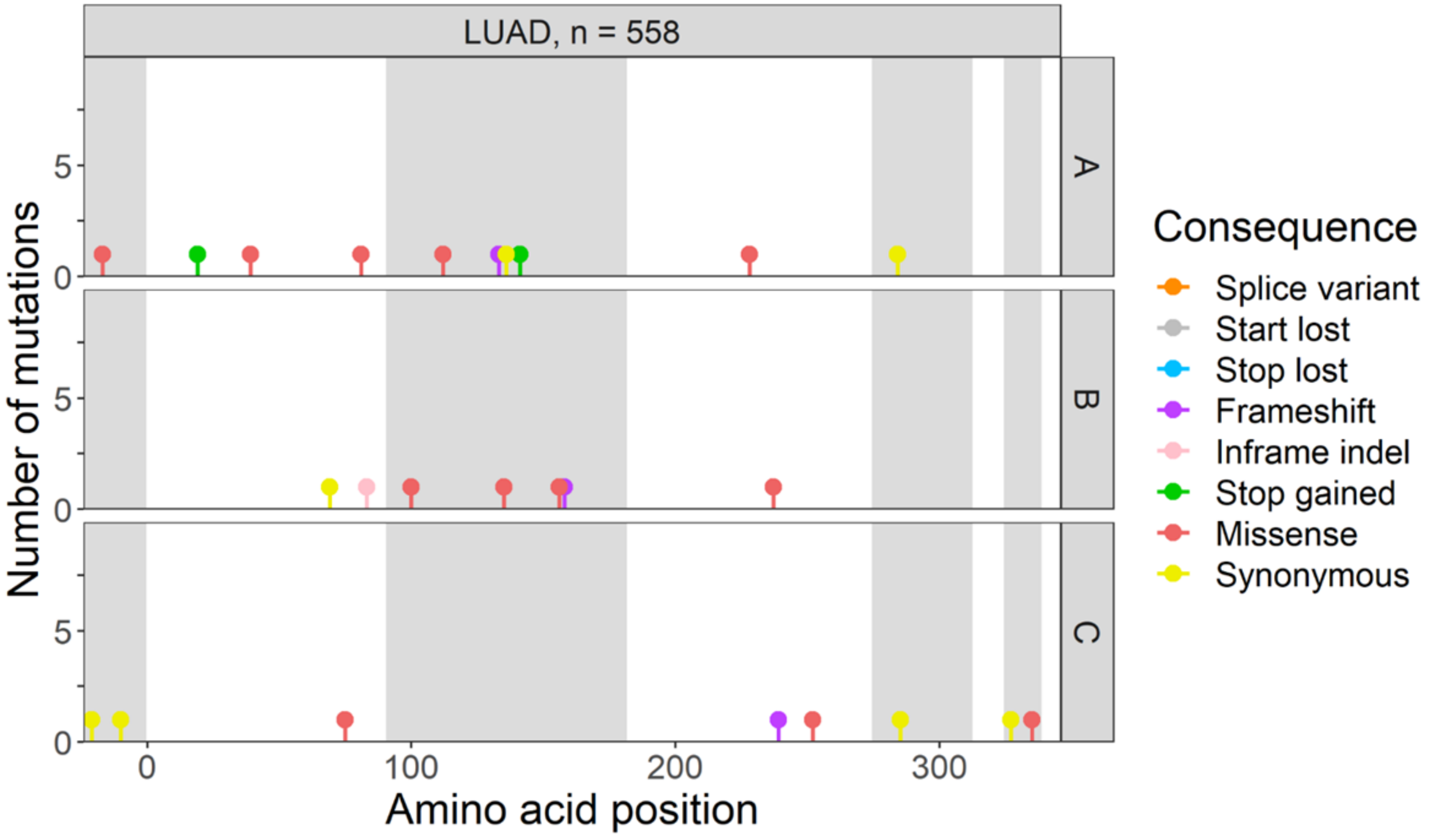

LUAD, n = 558
A
B
C
Number of mutations
Amino acid position
0
5
0
100
200
300
Consequence
Splice variant
Start lost
Stop lost
Frameshift
Inframe indel
Stop gained
Missense
Synonymous


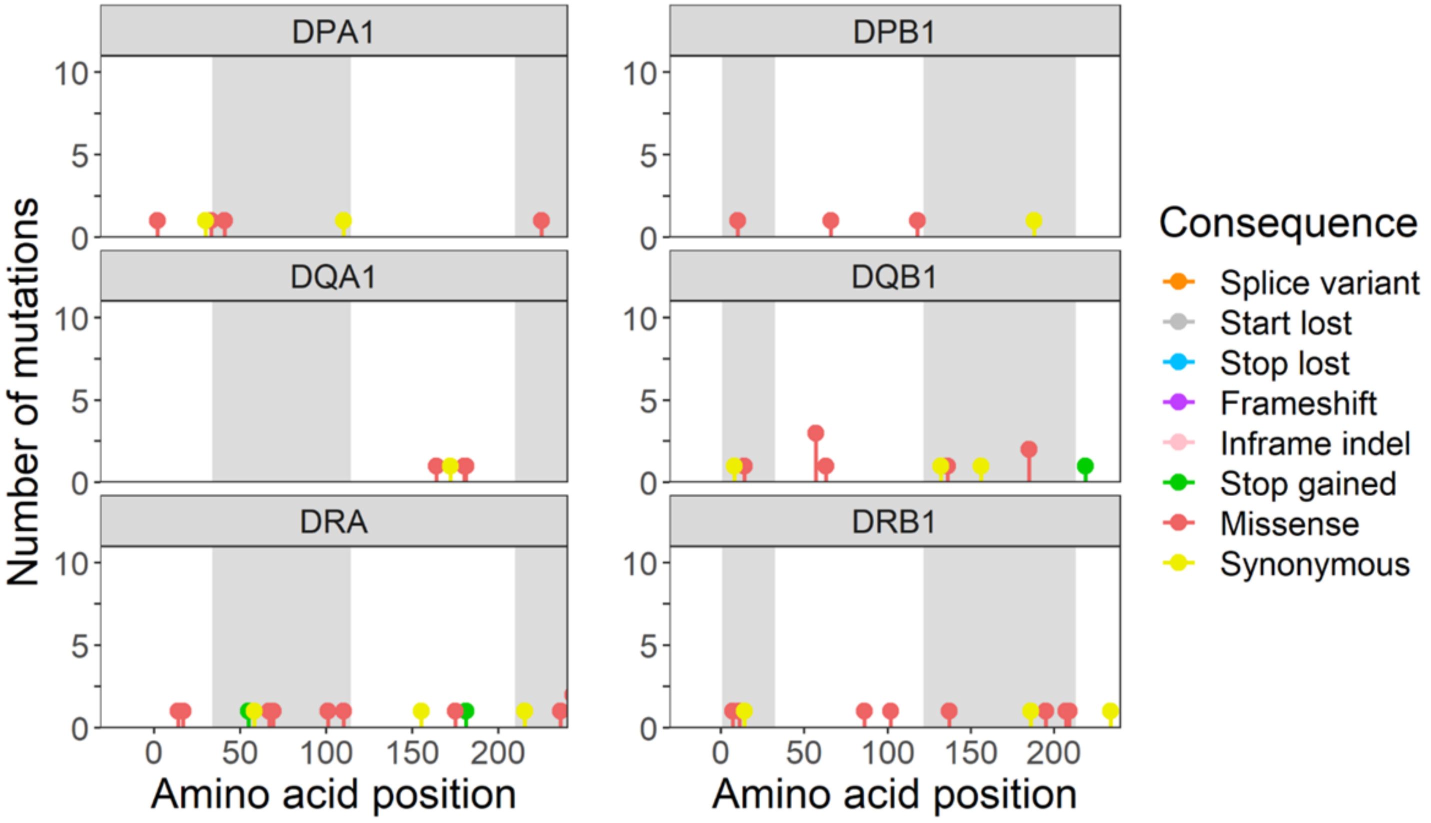

DPA1
DPB1
DQA1
DQB1
DRA
DRB1
Number of mutations
Amino acid position
0
5
10
0
50
100
150
200
Consequence
Splice variant
Start lost
Stop lost
Frameshift
Inframe indel
Stop gained
Missense
Synonymous

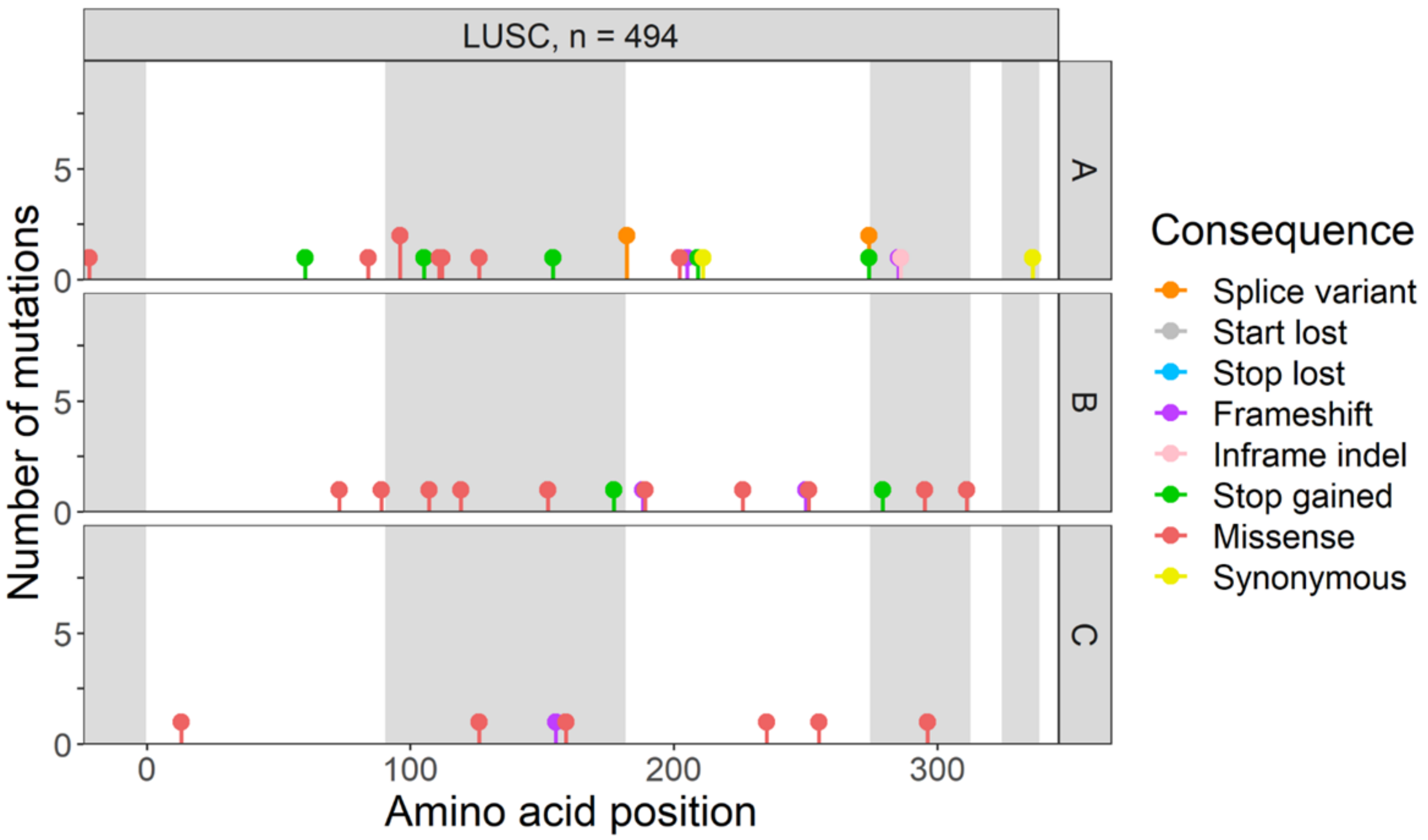

LUSC, n = 494
A
B
C
Number of mutations
Amino acid position
Consequence
Splice variant
Start lost
Stop lost
Frameshift
Inframe indel
Stop gained
Missense
Synonymous


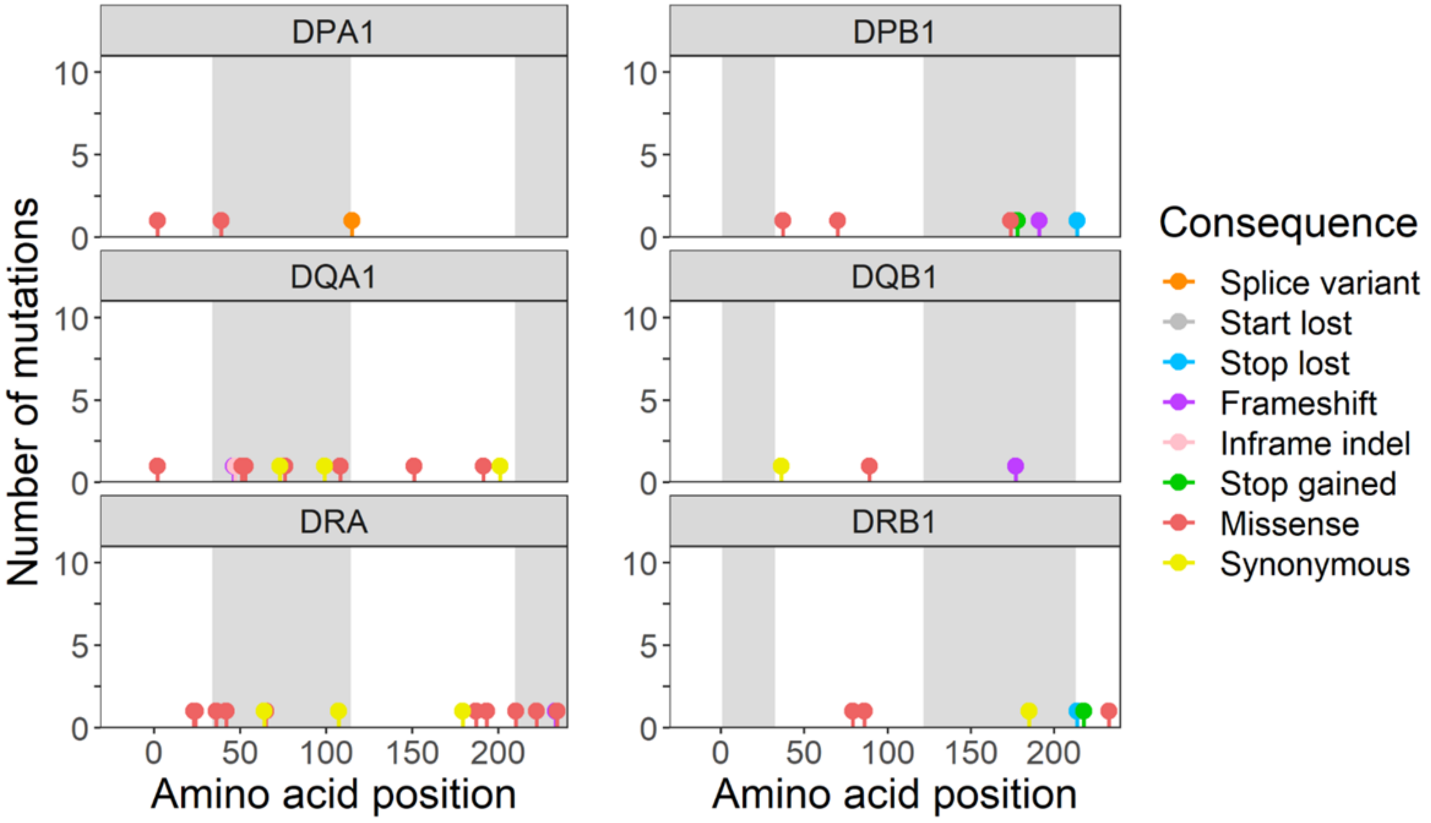

DPA1
DPB1
DQA1
DQB1
DRA
DRB1
Number of mutations
Amino acid position
Amino acid position
Consequence
Splice variant
Start lost
Stop lost
Frameshift
Inframe indel
Stop gained
Missense
Synonymous

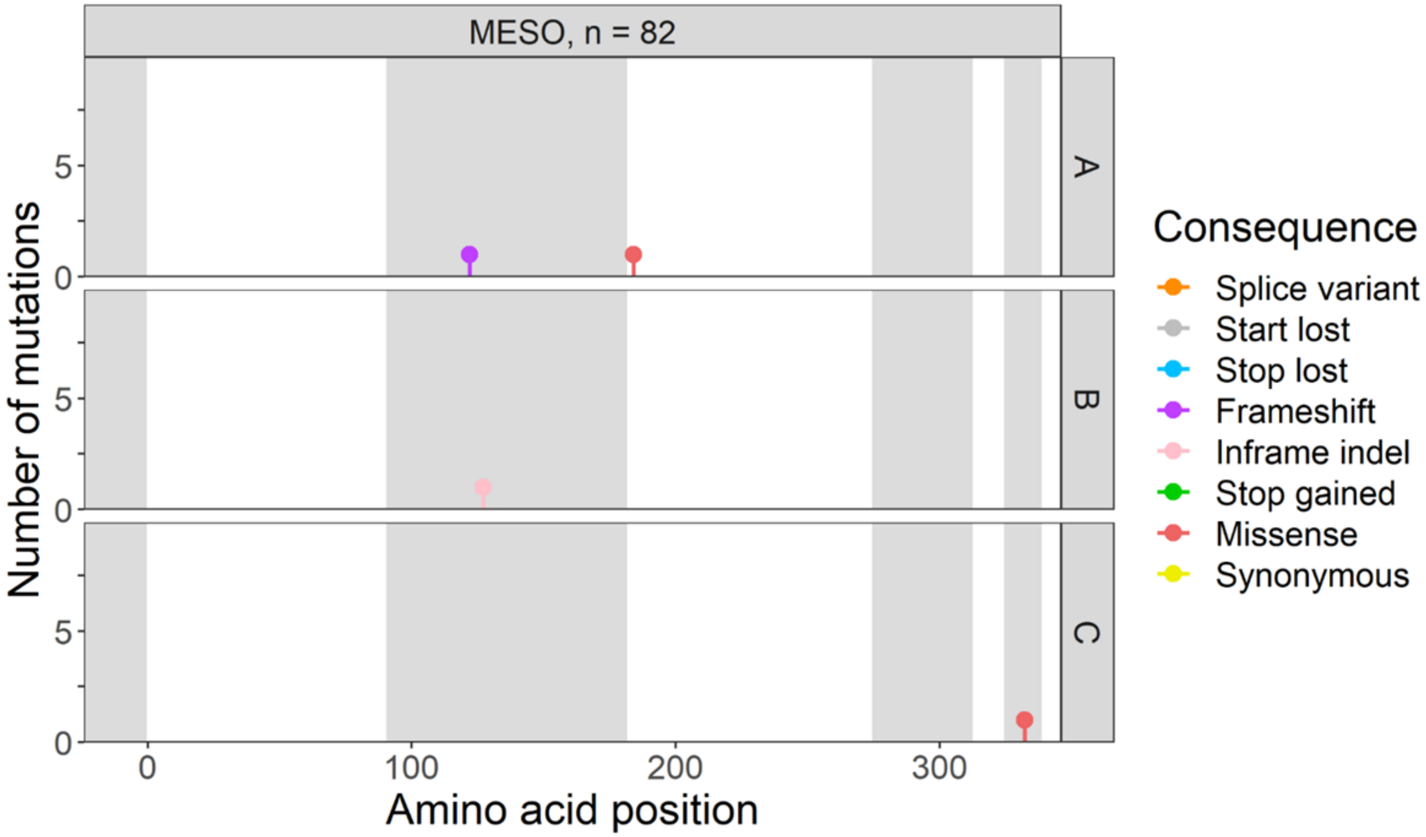
MESO, n = 82
A
B
C
Number of mutations
Amino acid position
Consequence
Splice variant
Start lost
Stop lost
Frameshift
Inframe indel
Stop gained
Missense
Synonymous

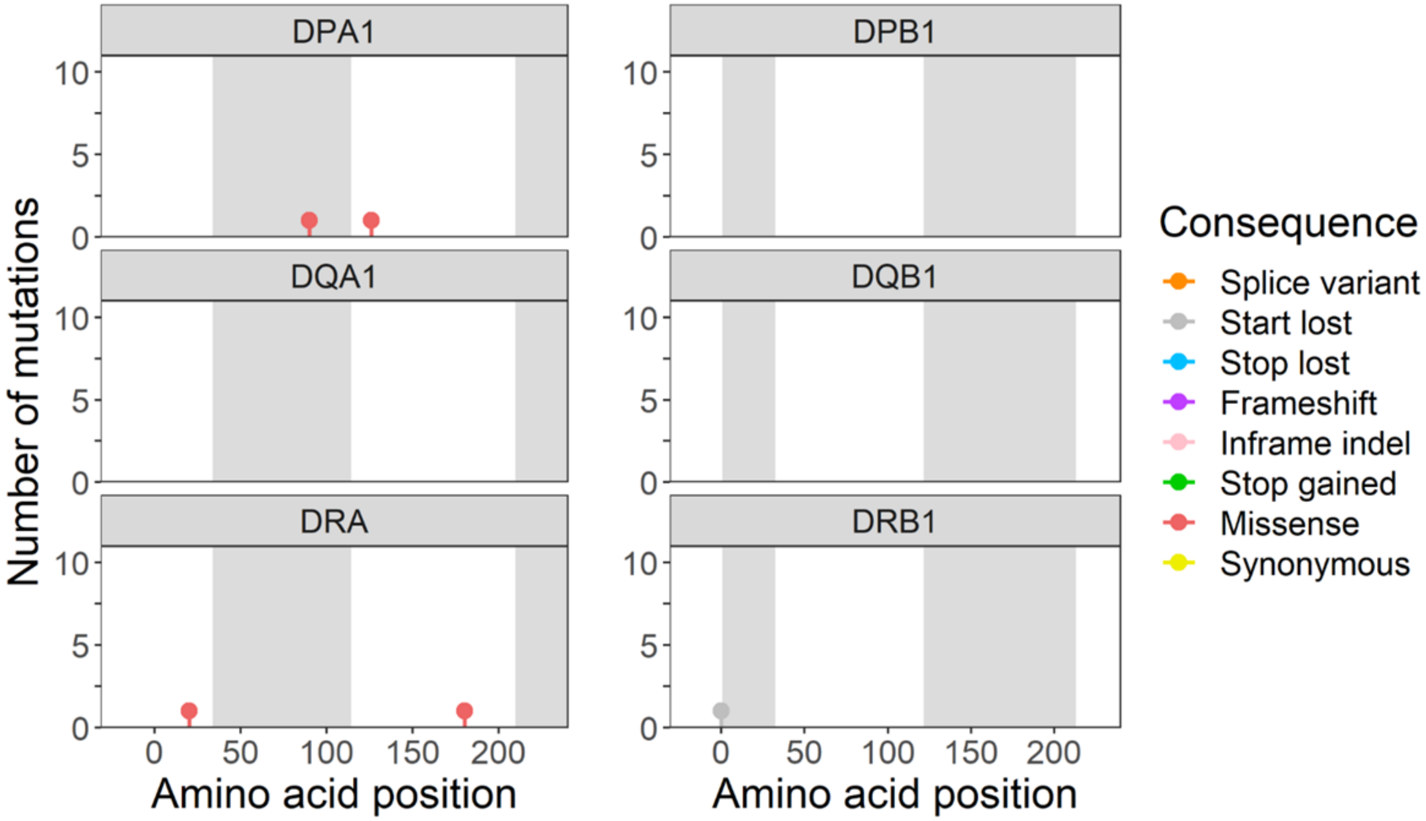
DPA1
DPB1
DQA1
DQB1
DRA
DRB1
Number of mutations
Amino acid position
Amino acid position
Consequence
Splice variant
Start lost
Stop lost
Frameshift
Inframe indel
Stop gained
Missense
Synonymous

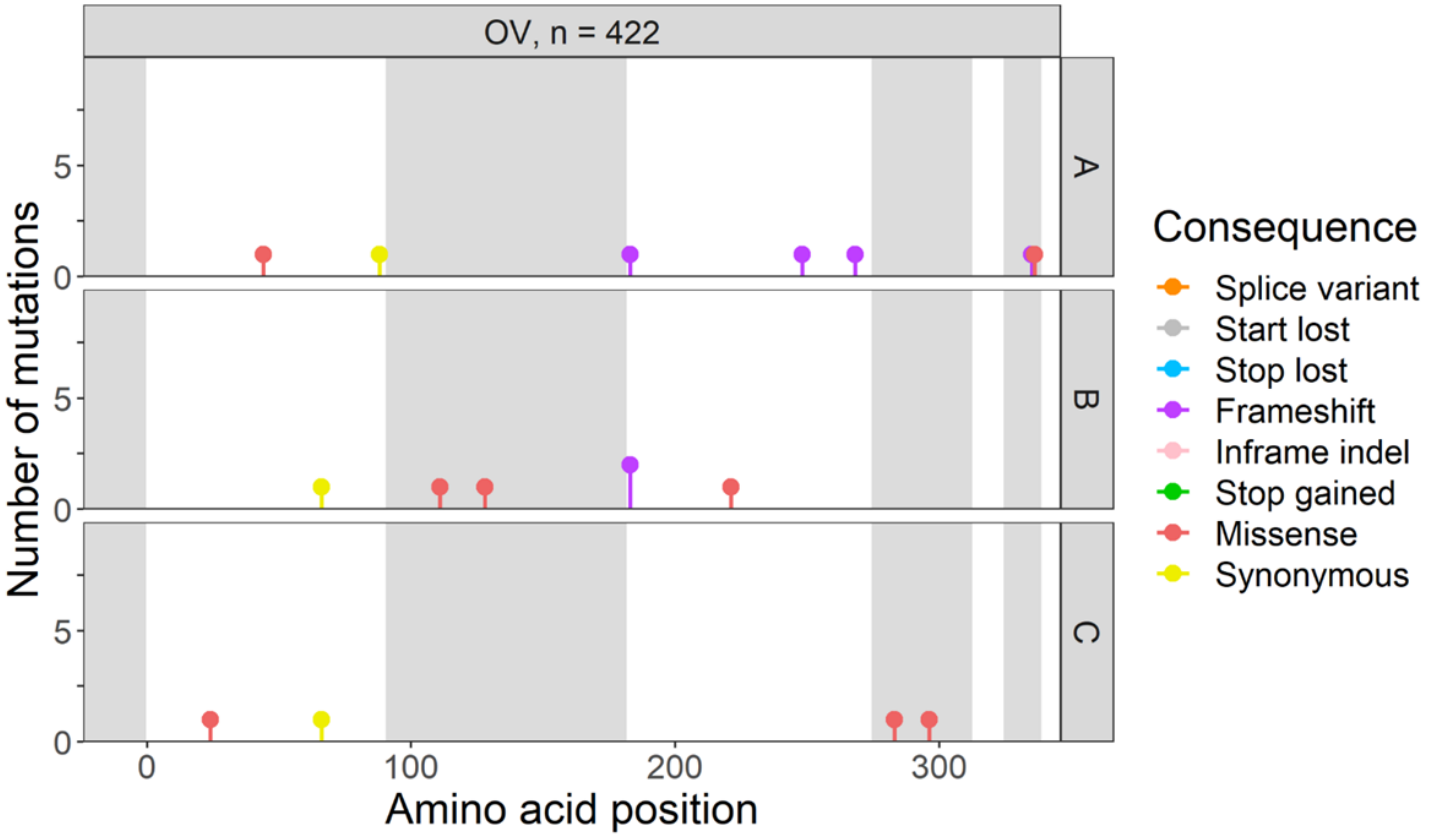

OV, n = 422
A
B
C
Number of mutations
Amino acid position
Consequence
Splice variant
Start lost
Stop lost
Frameshift
Inframe indel
Stop gained
Missense
Synonymous


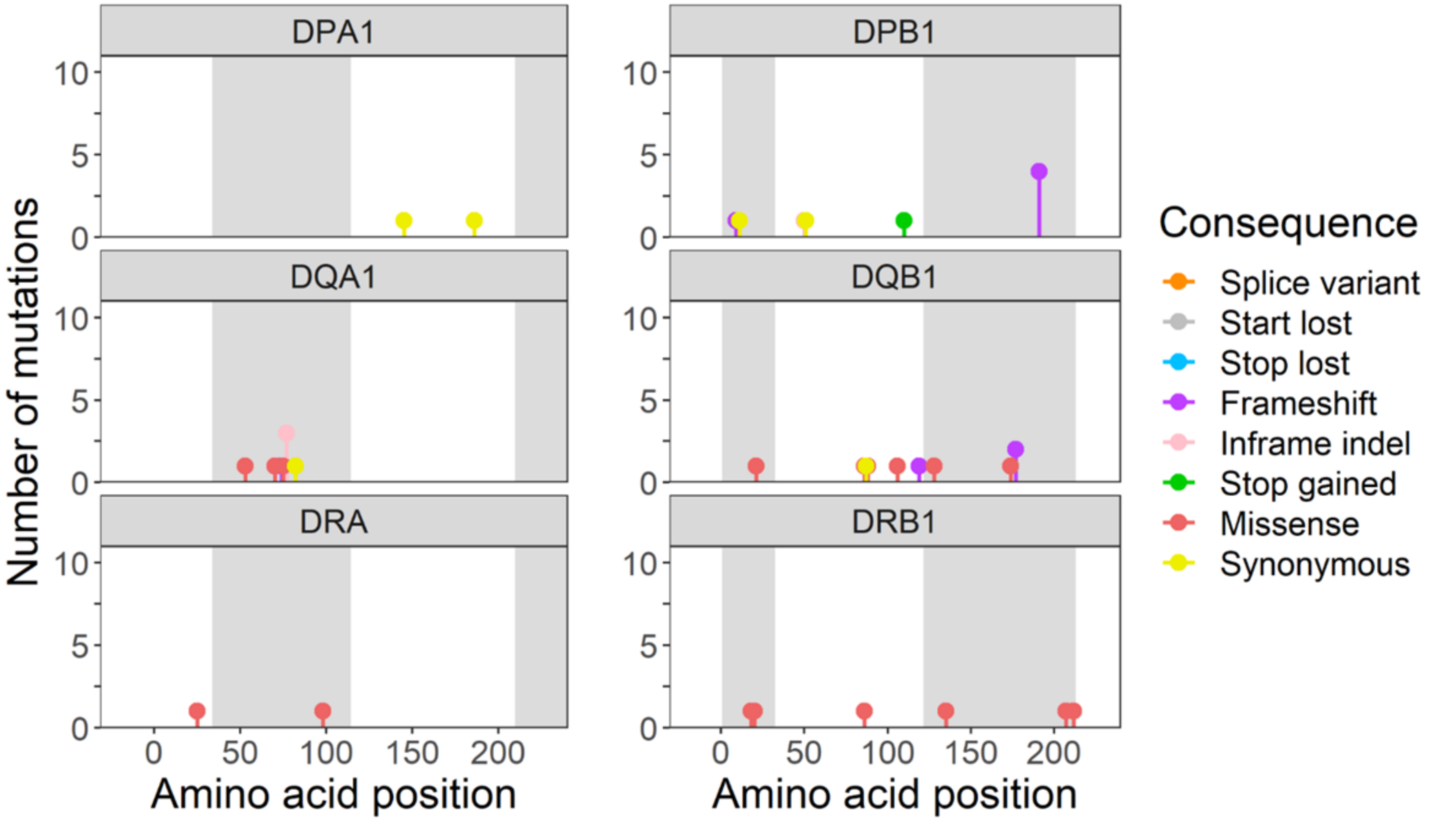

DPA1
DPB1
DQA1
DQB1
DRA
DRB1
Number of mutations
Amino acid position
Amino acid position
Consequence
Splice variant
Start lost
Stop lost
Frameshift
Inframe indel
Stop gained
Missense
Synonymous

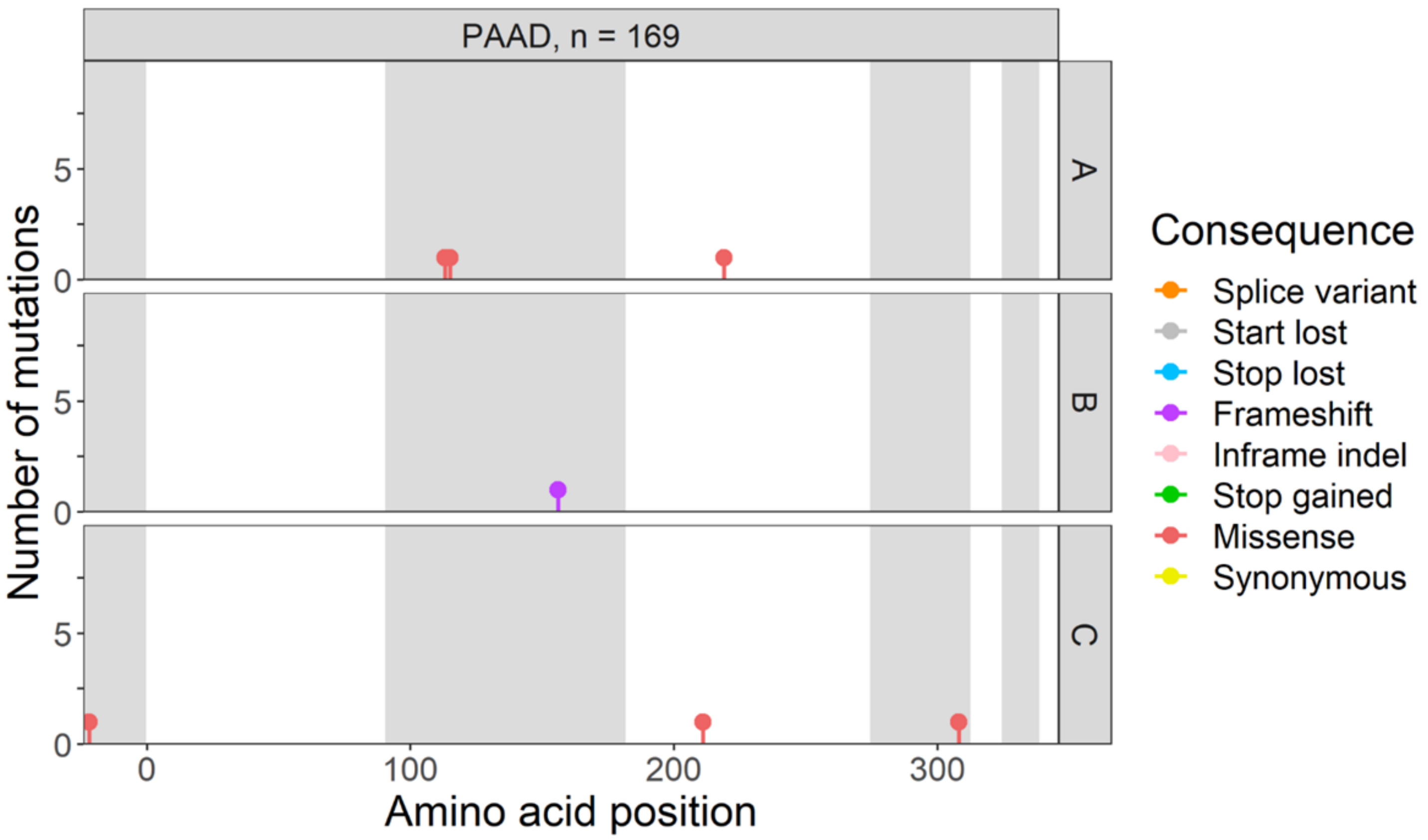

PAAD, n = 169
A
B
C
Number of mutations
Amino acid position
Consequence
Splice variant
Start lost
Stop lost
Frameshift
Inframe indel
Stop gained
Missense
Synonymous


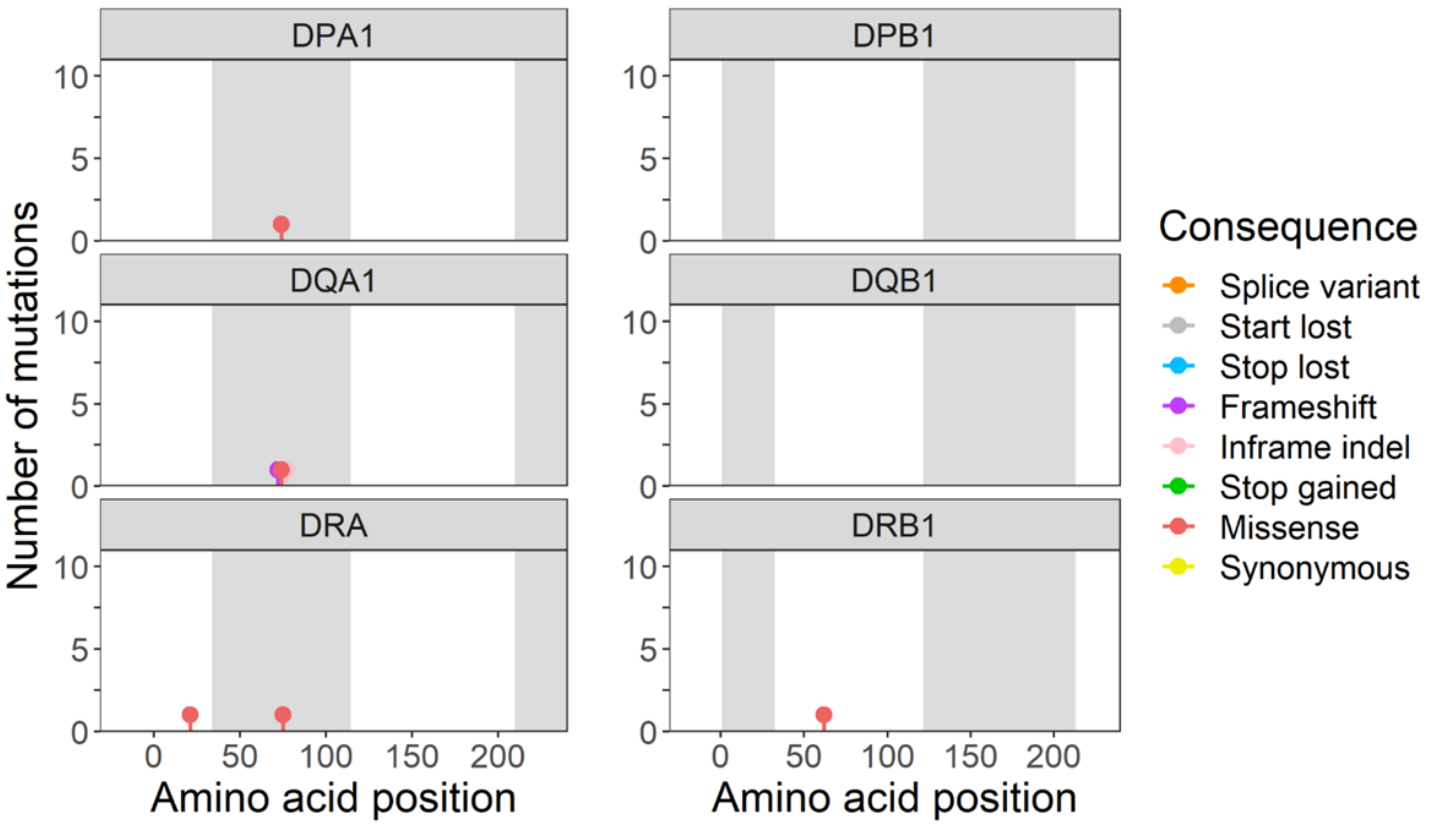

DPA1
DPB1
DQA1
DQB1
DRA
DRB1
Number of mutations
Amino acid position
Amino acid position
Consequence
Splice variant
Start lost
Stop lost
Frameshift
Inframe indel
Stop gained
Missense
Synonymous

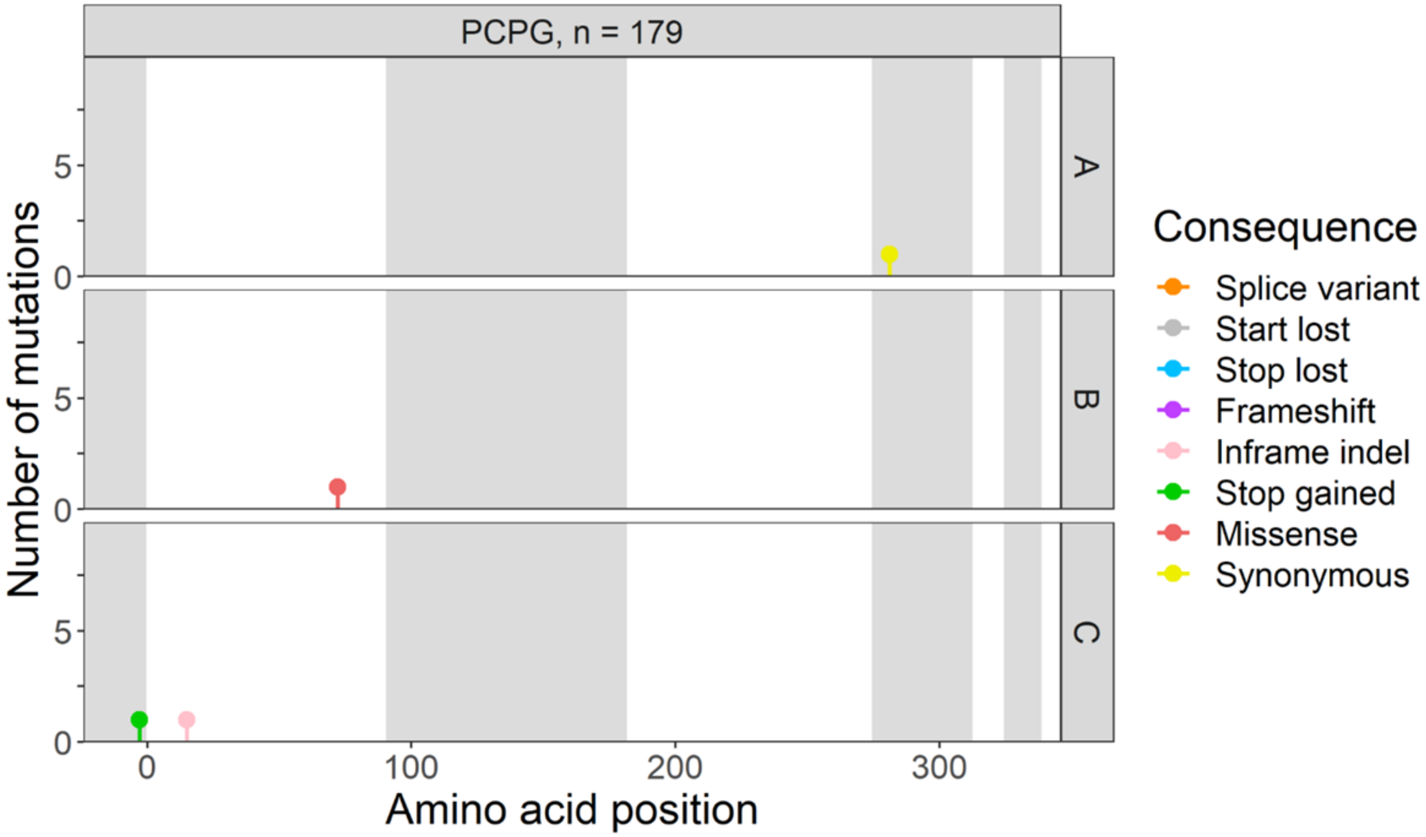

PCPG, n = 179
A
B
C
Number of mutations
Amino acid position
Consequence
Splice variant
Start lost
Stop lost
Frameshift
Inframe indel
Stop gained
Missense
Synonymous


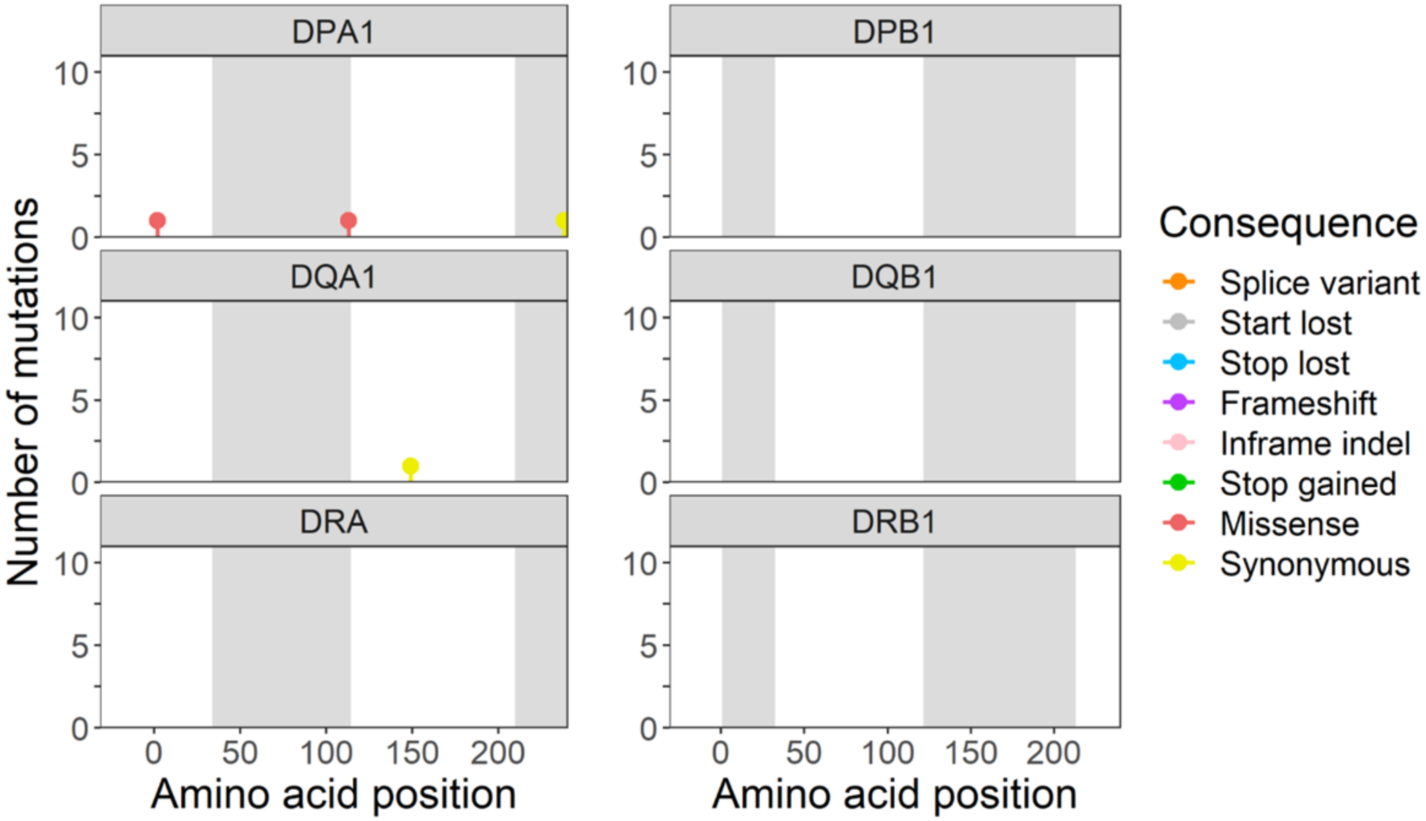

DPA1
DPB1
DQA1
DQB1
DRA
DRB1
Number of mutations
Amino acid position
Amino acid position
Consequence
Splice variant
Start lost
Stop lost
Frameshift
Inframe indel
Stop gained
Missense
Synonymous

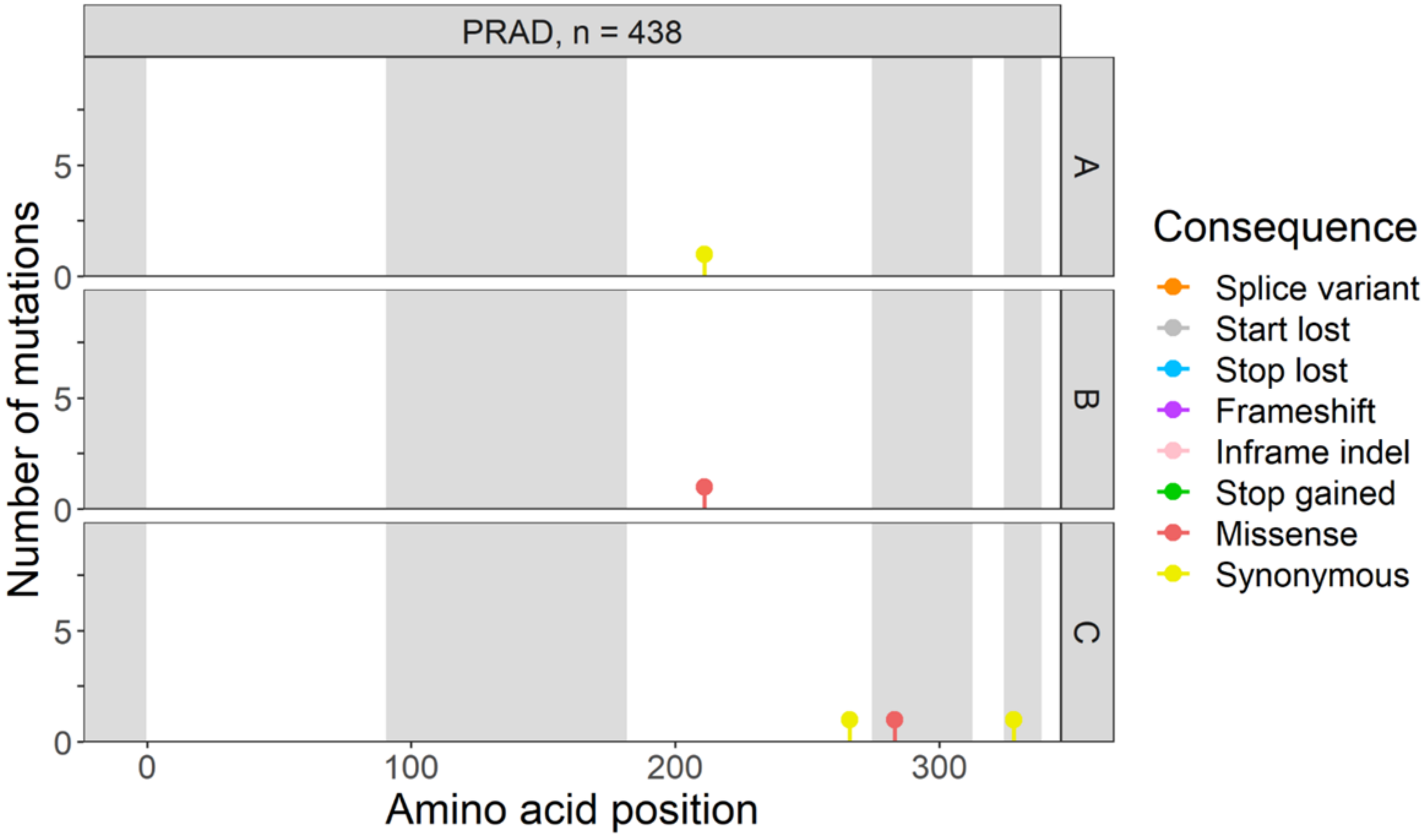

PRAD, n = 438
Number of mutations
A
B
C
Amino acid position
Consequence
Splice variant
Start lost
Stop lost
Frameshift
Inframe indel
Stop gained
Missense
Synonymous


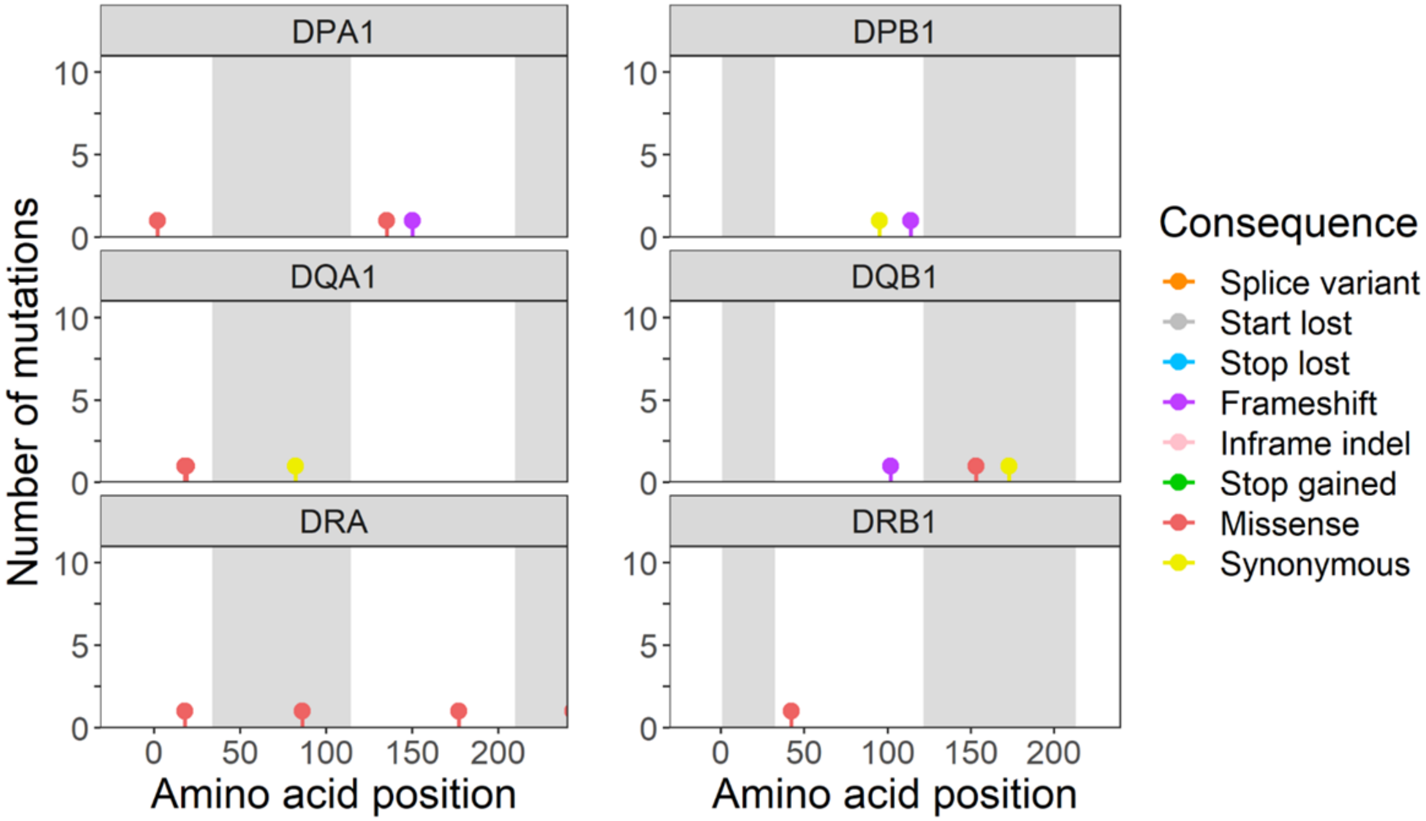

DPA1
DPB1
DQA1
DQB1
DRA
DRB1
Number of mutations
Amino acid position
Amino acid position
Consequence
Splice variant
Start lost
Stop lost
Frameshift
Inframe indel
Stop gained
Missense
Synonymous

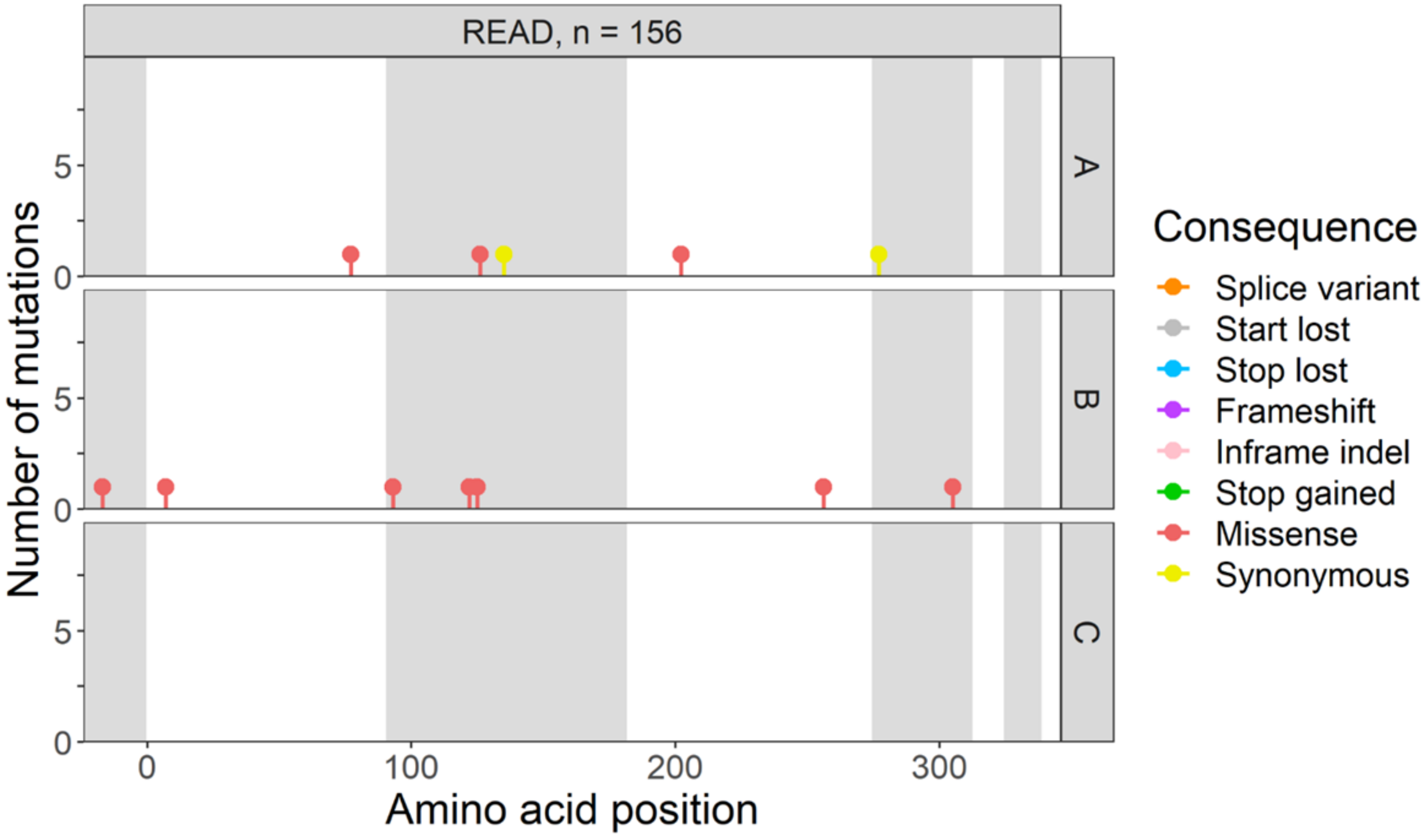

READ, n = 156
A
B
C
Number of mutations
Amino acid position
Consequence
Splice variant
Start lost
Stop lost
Frameshift
Inframe indel
Stop gained
Missense
Synonymous


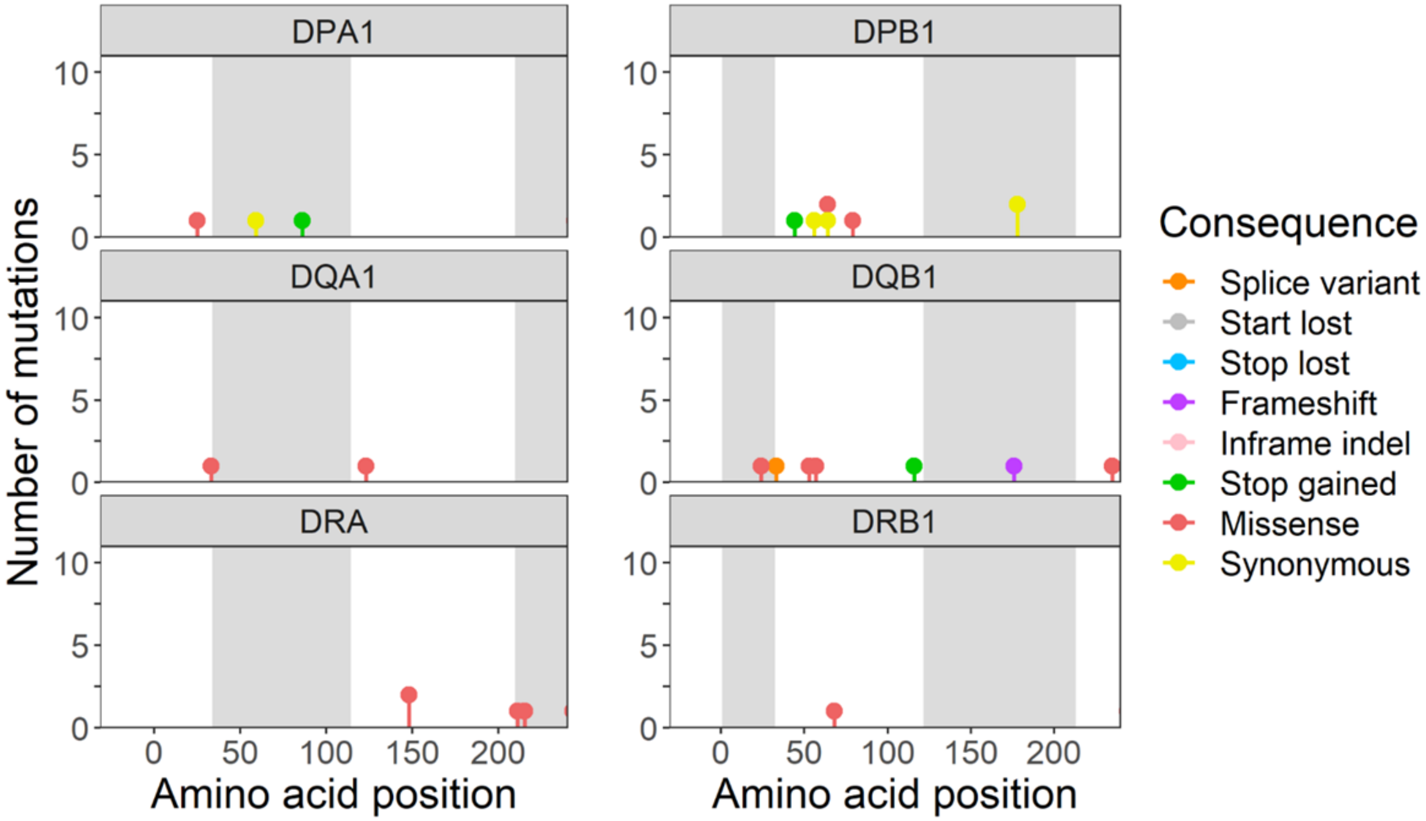

DPA1
DPB1
DQA1
DQB1
DRA
DRB1
Number of mutations
Amino acid position
Amino acid position
Consequence
Splice variant
Start lost
Stop lost
Frameshift
Inframe indel
Stop gained
Missense
Synonymous

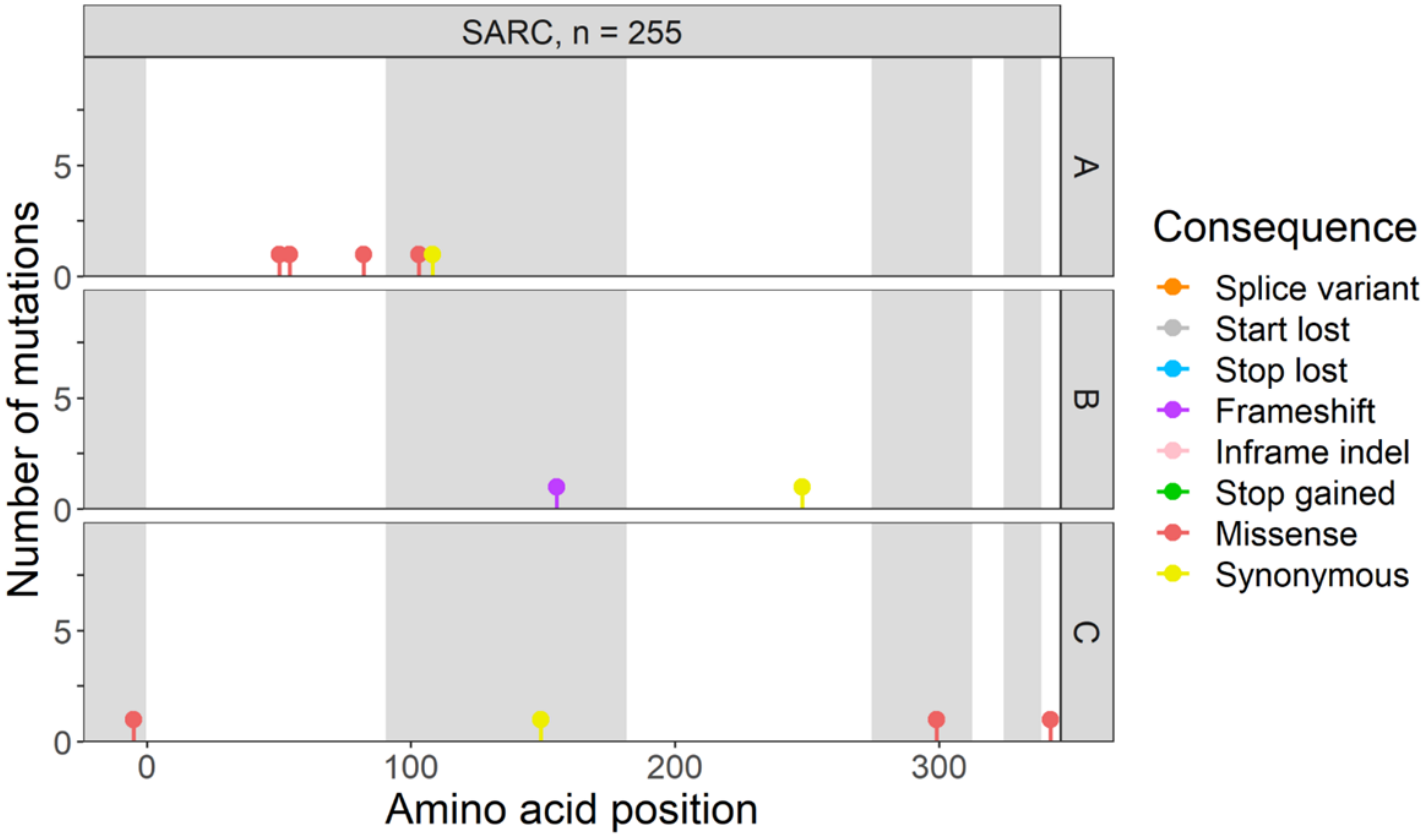
SARC, n = 255
A
B
C
Number of mutations
Amino acid position
0
100
200
300
Consequence
Splice variant
Start lost
Stop lost
Frameshift
Inframe indel
Stop gained
Missense
Synonymous

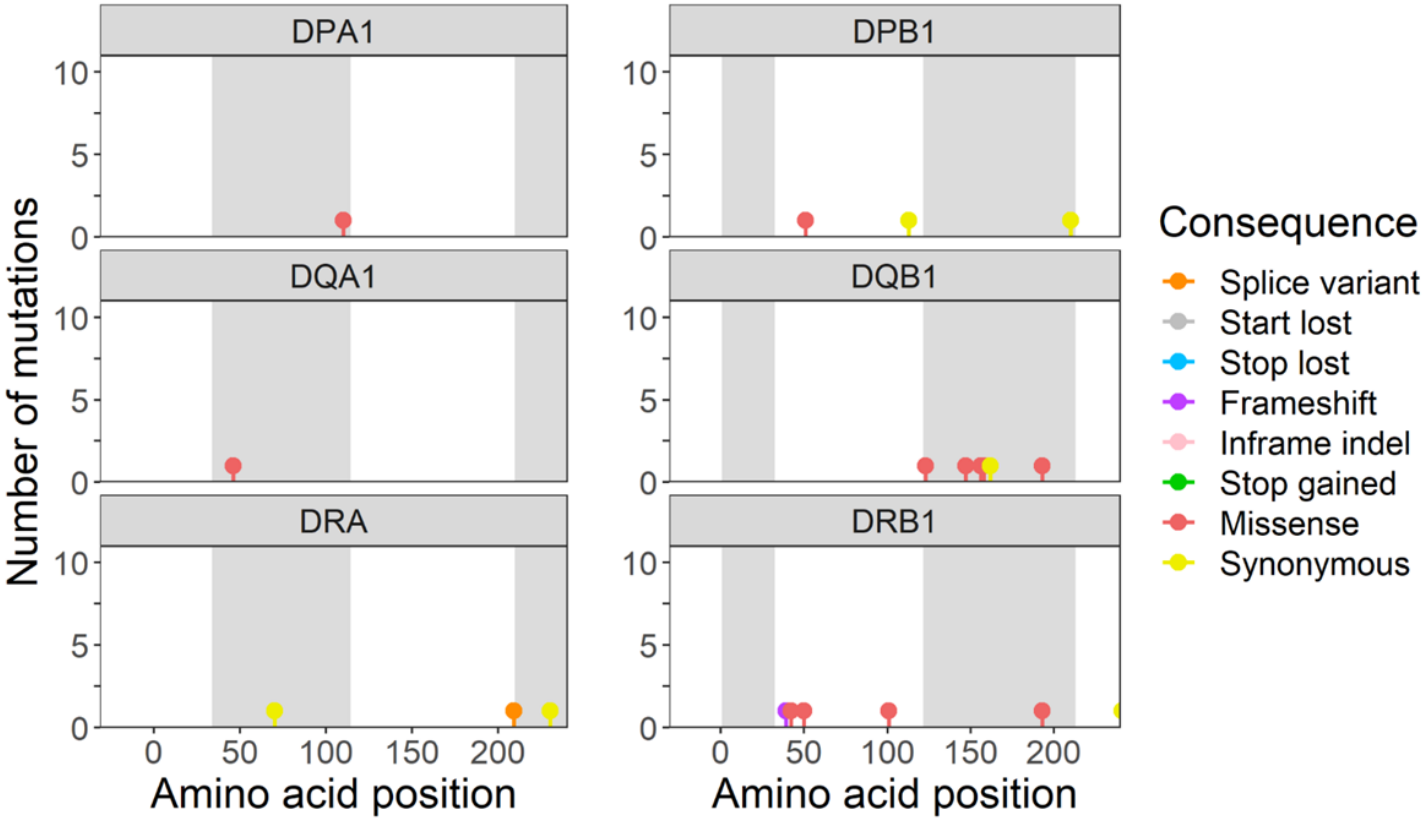
DPA1
DPB1
DQA1
DQB1
DRA
DRB1
Number of mutations
Amino acid position
0
50
100
150
200
Consequence
Splice variant
Start lost
Stop lost
Frameshift
Inframe indel
Stop gained
Missense
Synonymous

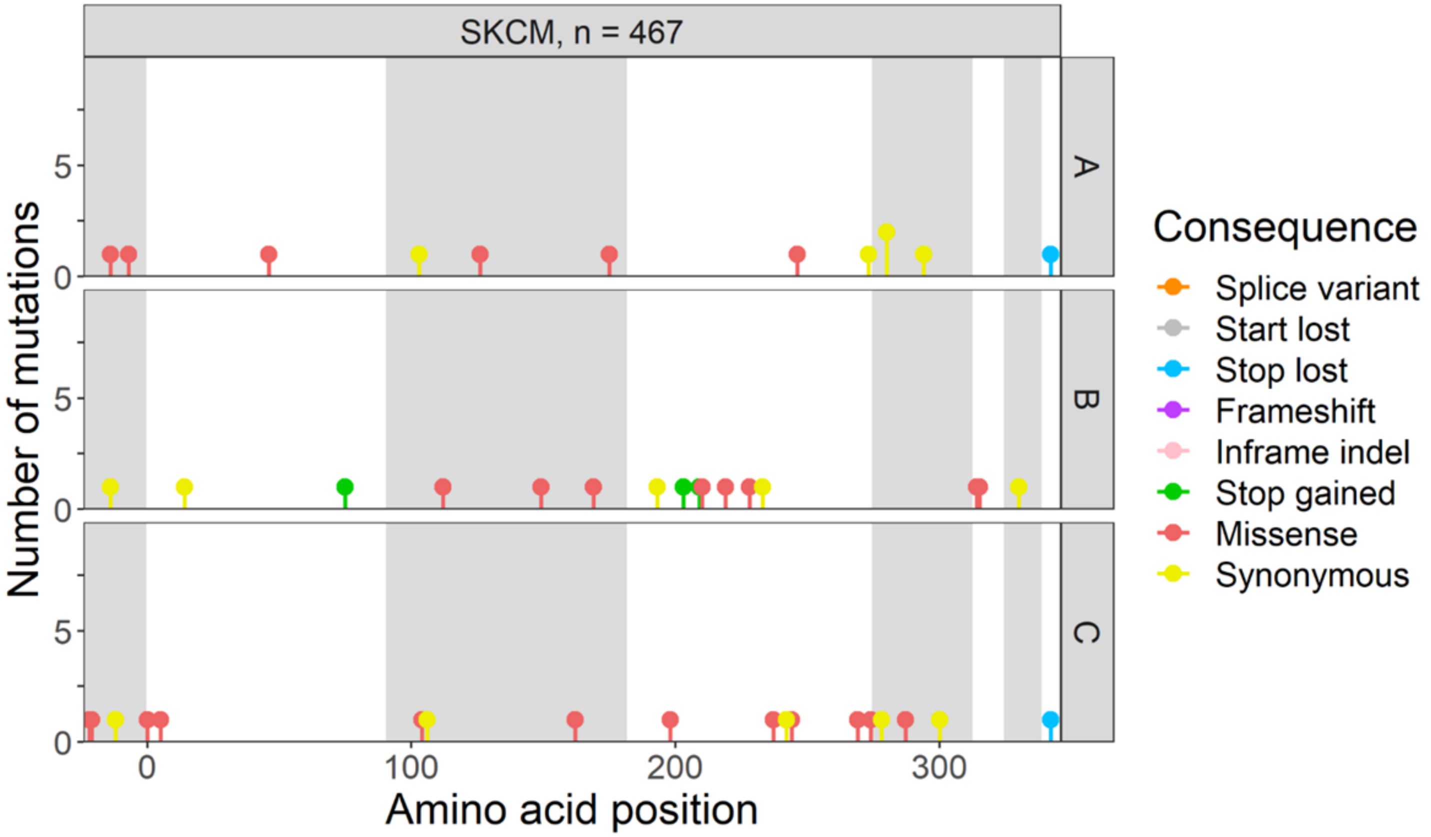
SKCM, n = 467
A
B
C
Number of mutations
Amino acid position
0
5
100
200
300
Consequence
Splice variant
Start lost
Stop lost
Frameshift
Inframe indel
Stop gained
Missense
Synonymous

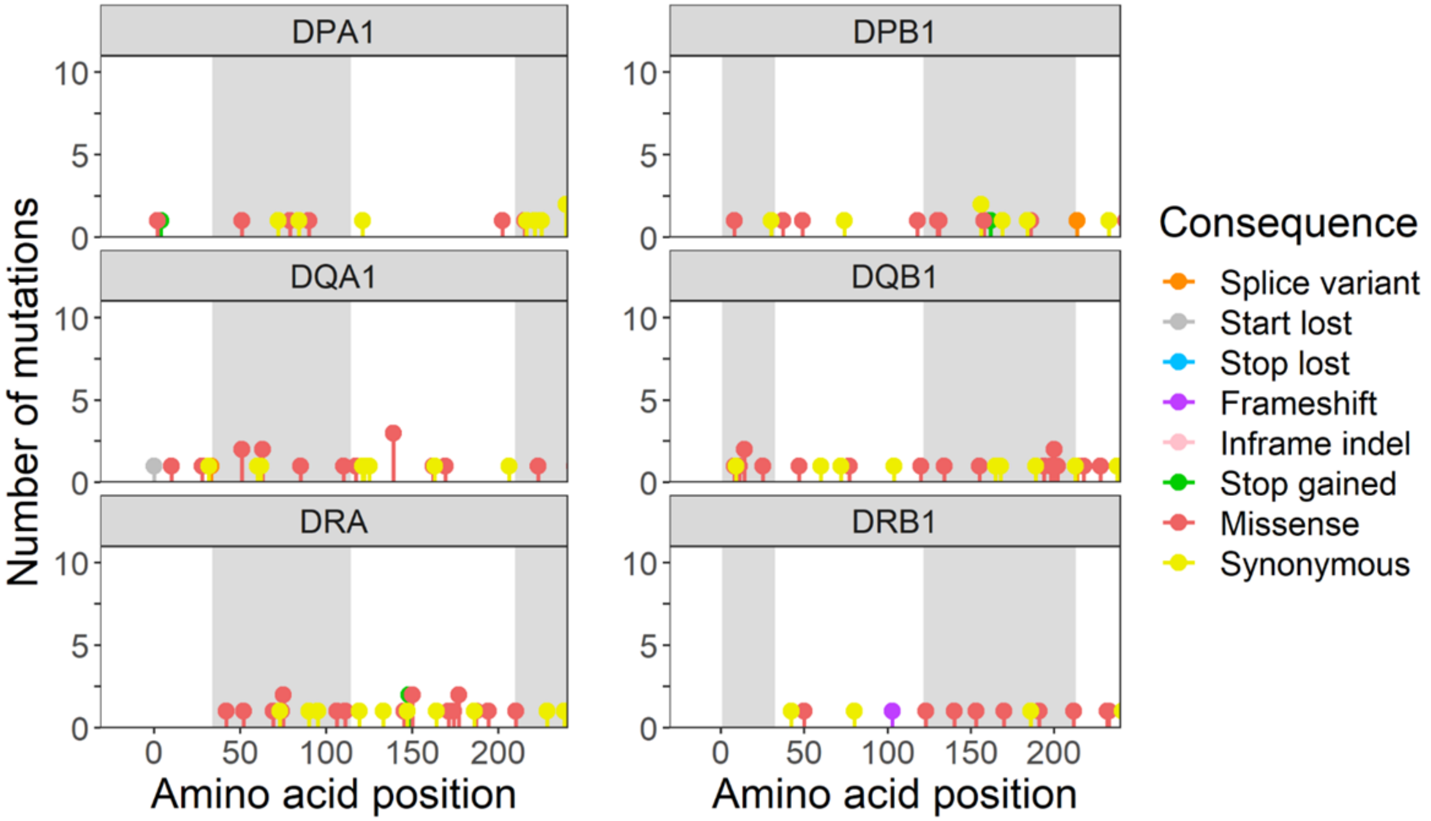
DPA1
DPB1
DQA1
DQB1
DRA
DRB1
Number of mutations
Amino acid position
0
5
10
50
100
150
200
Consequence
Splice variant
Start lost
Stop lost
Frameshift
Inframe indel
Stop gained
Missense
Synonymous

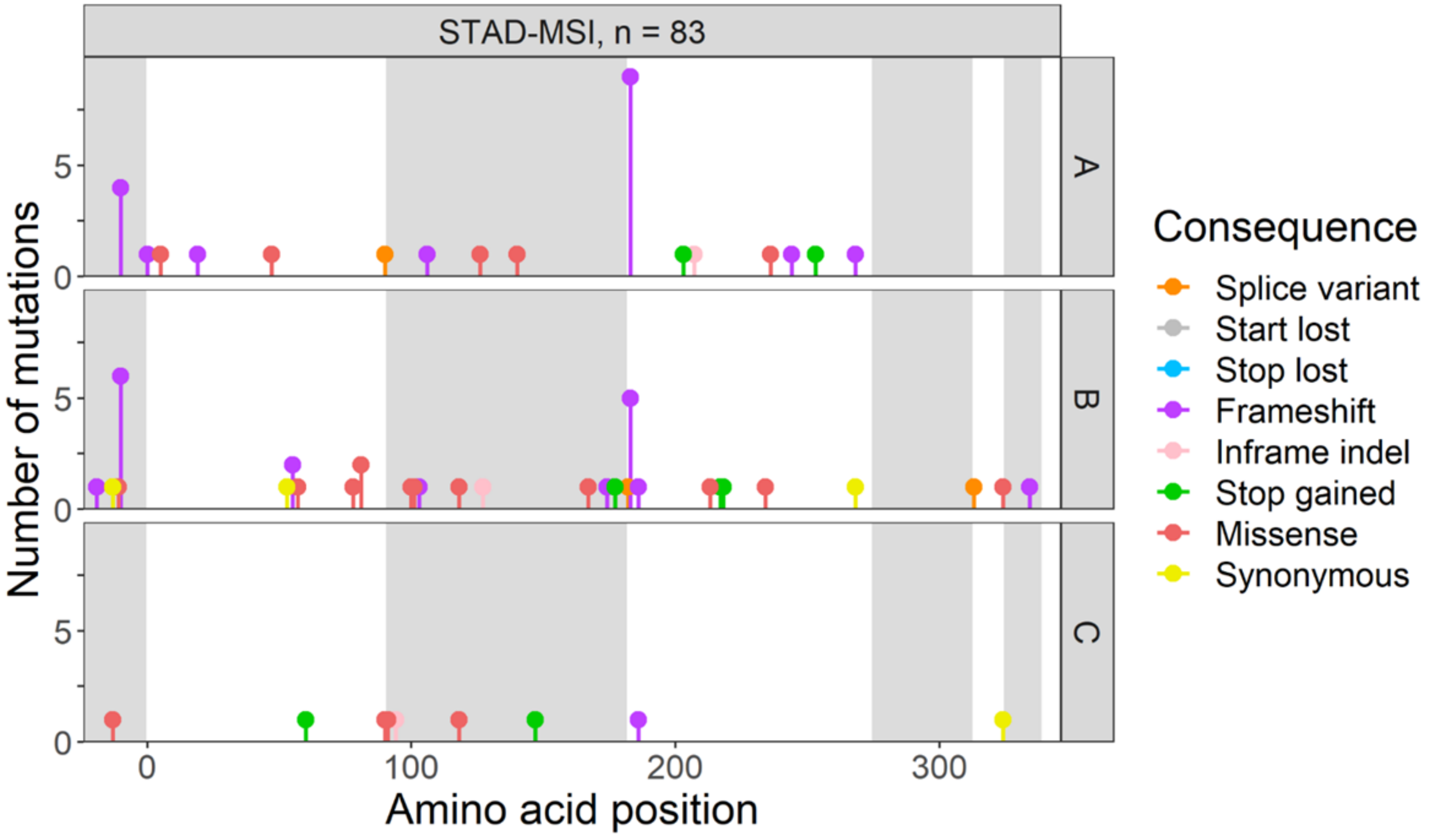
STAD-MSI, n = 83
A
B
C
Number of mutations
Amino acid position
0
5
0
100
200
300
Consequence
Splice variant
Start lost
Stop lost
Frameshift
Inframe indel
Stop gained
Missense
Synonymous

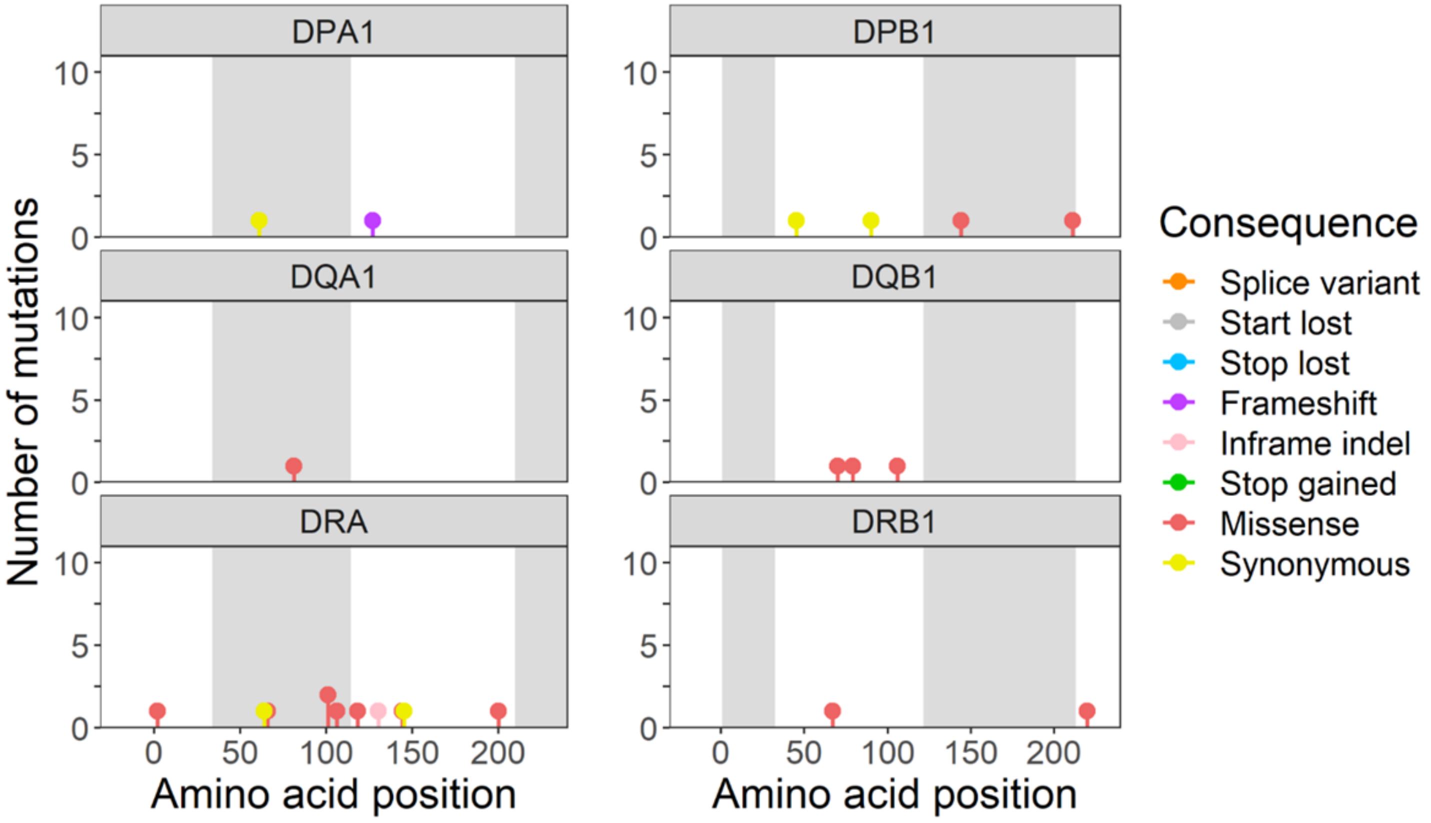
DPA1
DPB1
DQA1
DQB1
DRA
DRB1
Number of mutations
Amino acid position
0
5
10
0
50
100
150
200
Consequence
Splice variant
Start lost
Stop lost
Frameshift
Inframe indel
Stop gained
Missense
Synonymous

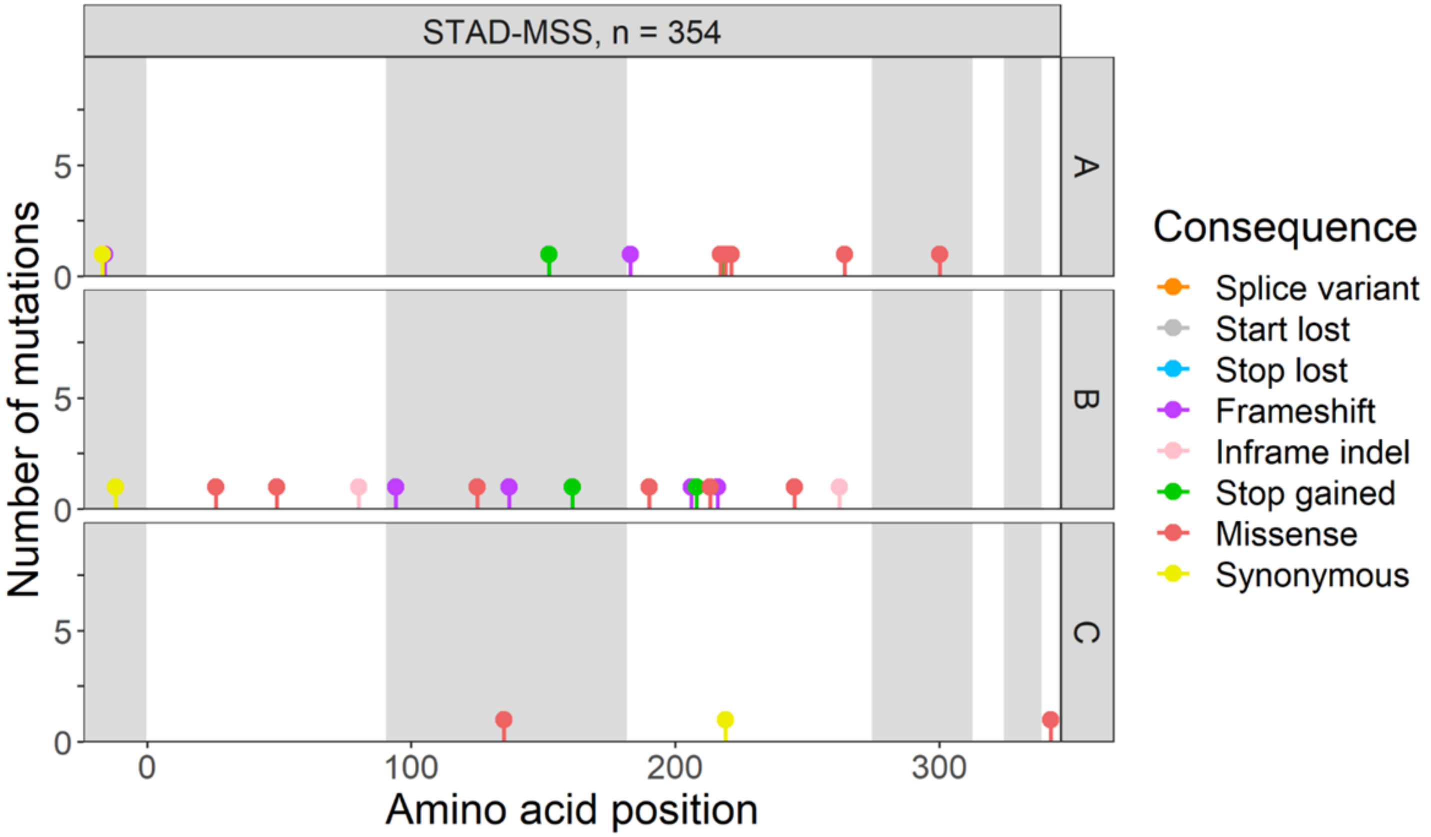
STAD-MSS, n = 354
A
B
C
Number of mutations
0
5
0
100
200
300
Amino acid position
Consequence
Splice variant
Start lost
Stop lost
Frameshift
Inframe indel
Stop gained
Missense
Synonymous

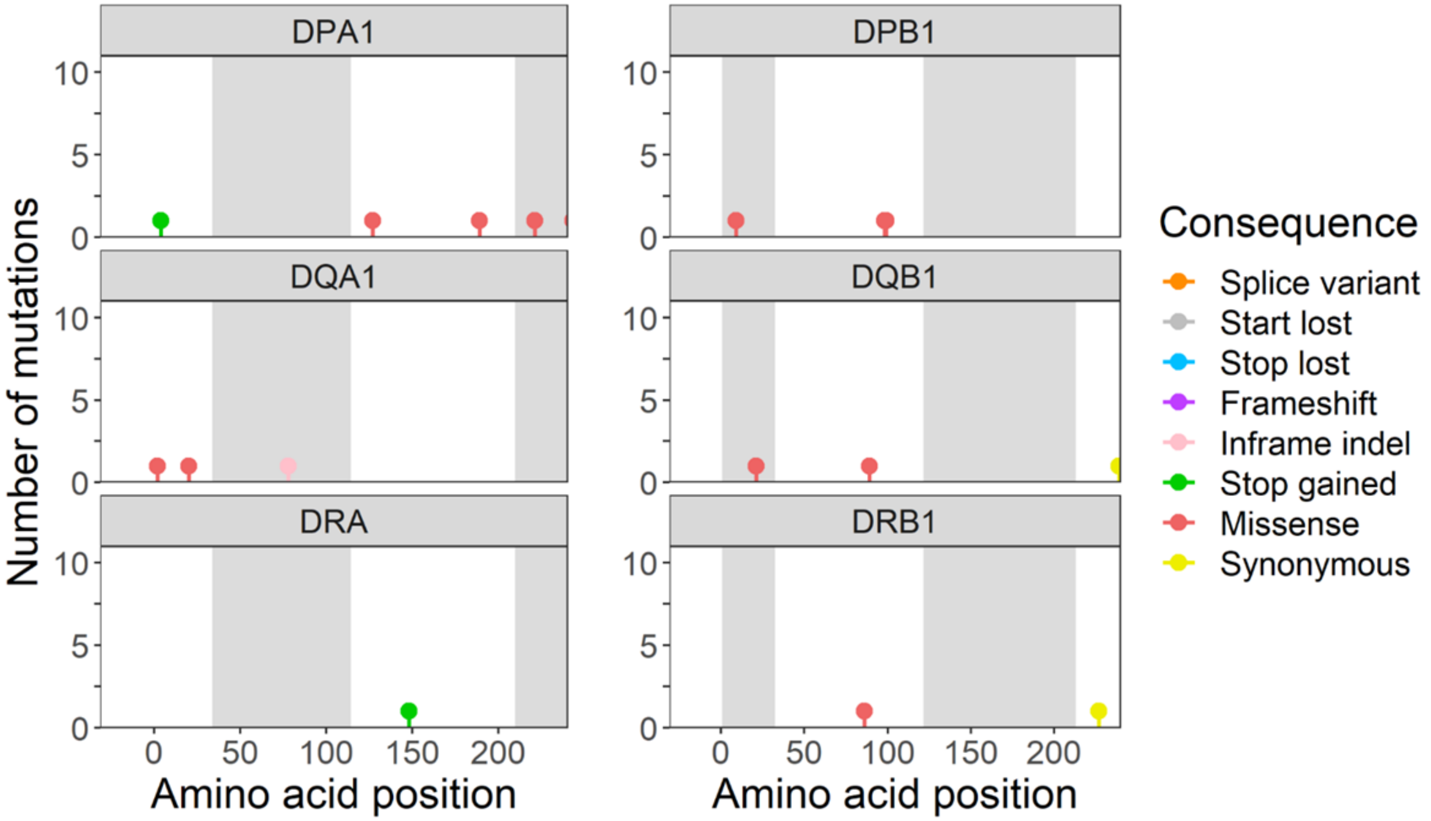
DPA1
DPB1
DQA1
DQB1
DRA
DRB1
Number of mutations
0
5
10
0
50
100
150
200
Amino acid position
Amino acid position
Consequence
Splice variant
Start lost
Stop lost
Frameshift
Inframe indel
Stop gained
Missense
Synonymous

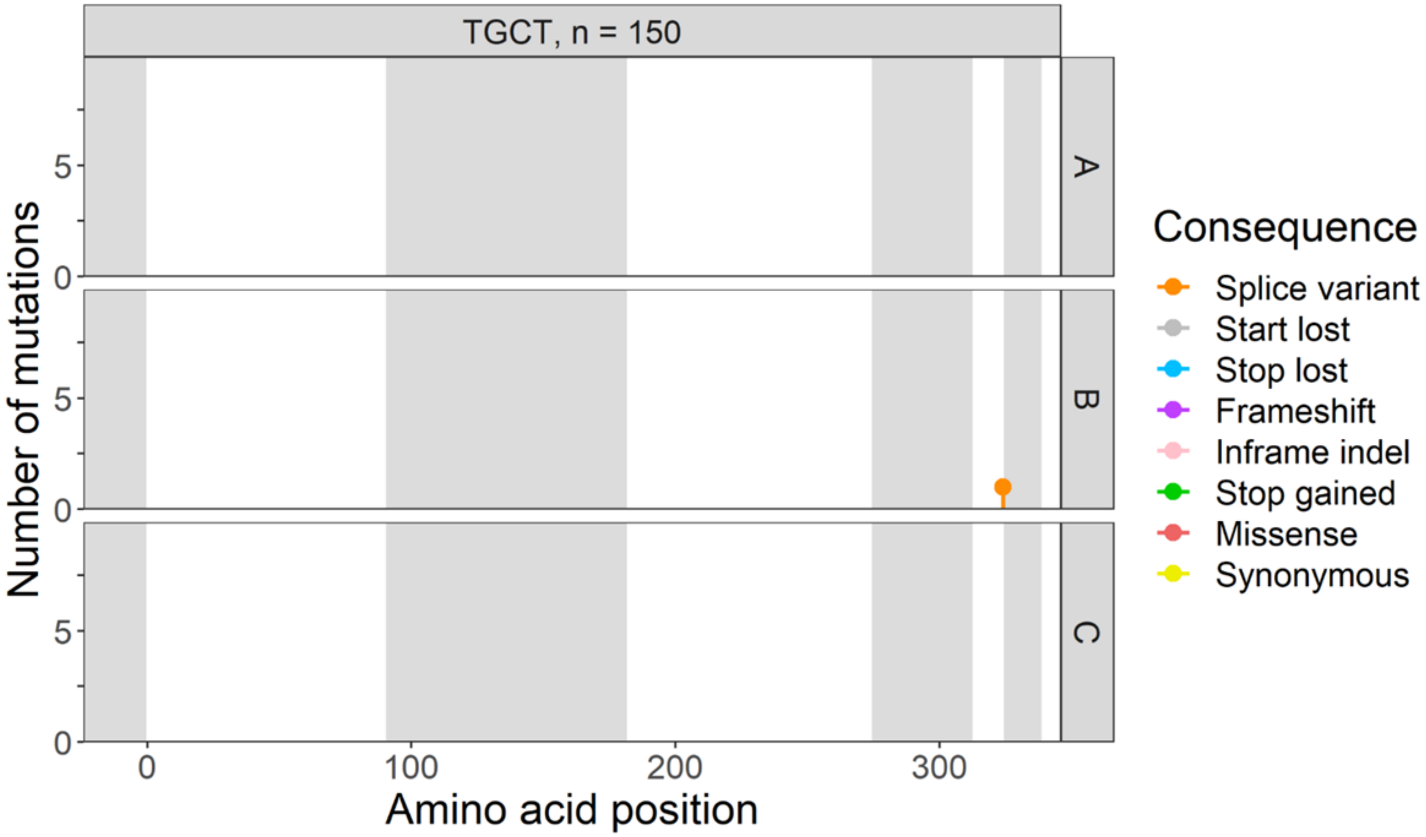
TGCT, n = 150
A
B
C
Number of mutations
0
5
0
100
200
300
Amino acid position
Consequence
Splice variant
Start lost
Stop lost
Frameshift
Inframe indel
Stop gained
Missense
Synonymous

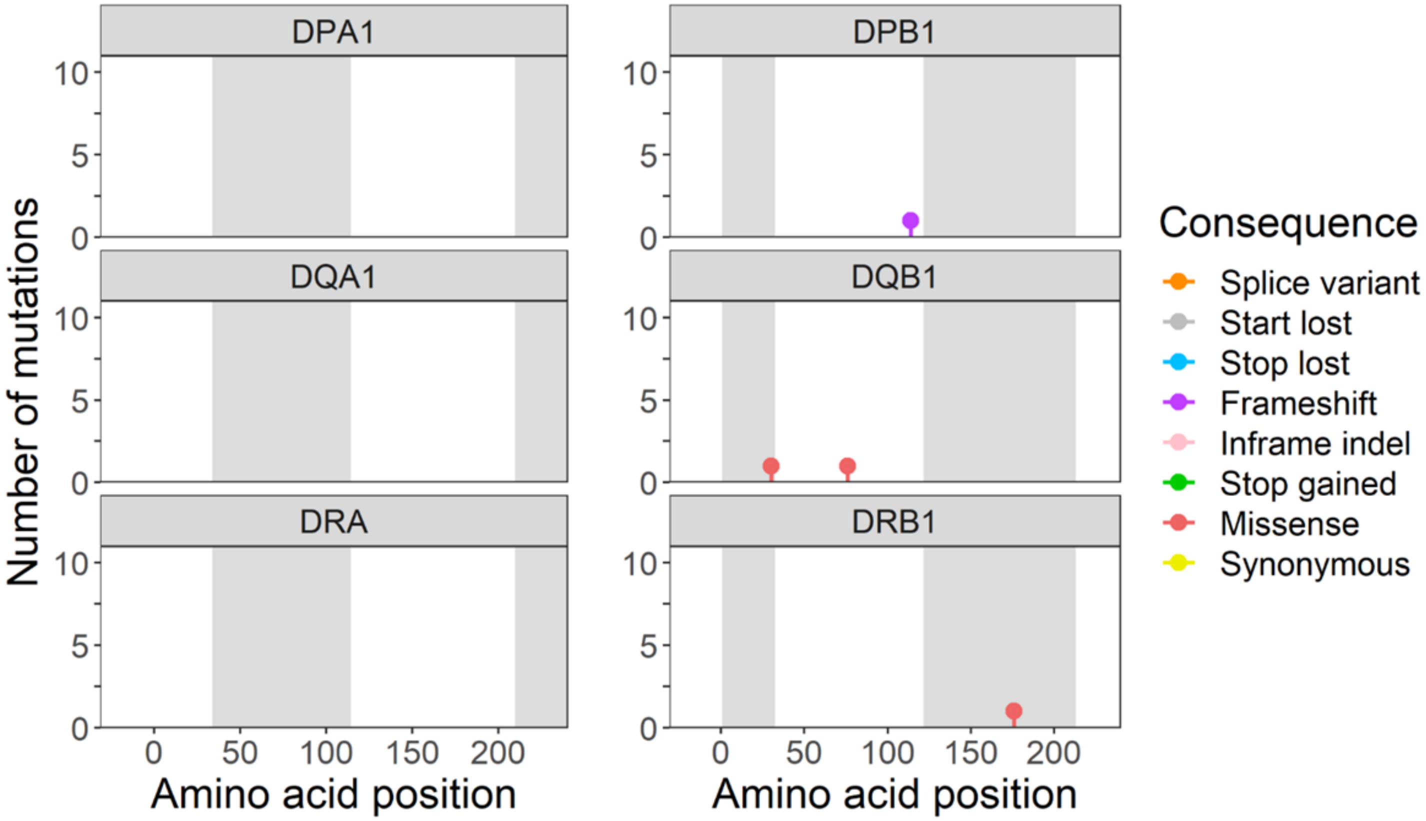
DPA1
DPB1
DQA1
DQB1
DRA
DRB1
Number of mutations
0
5
10
0
50
100
150
200
Amino acid position
Amino acid position
Consequence
Splice variant
Start lost
Stop lost
Frameshift
Inframe indel
Stop gained
Missense
Synonymous

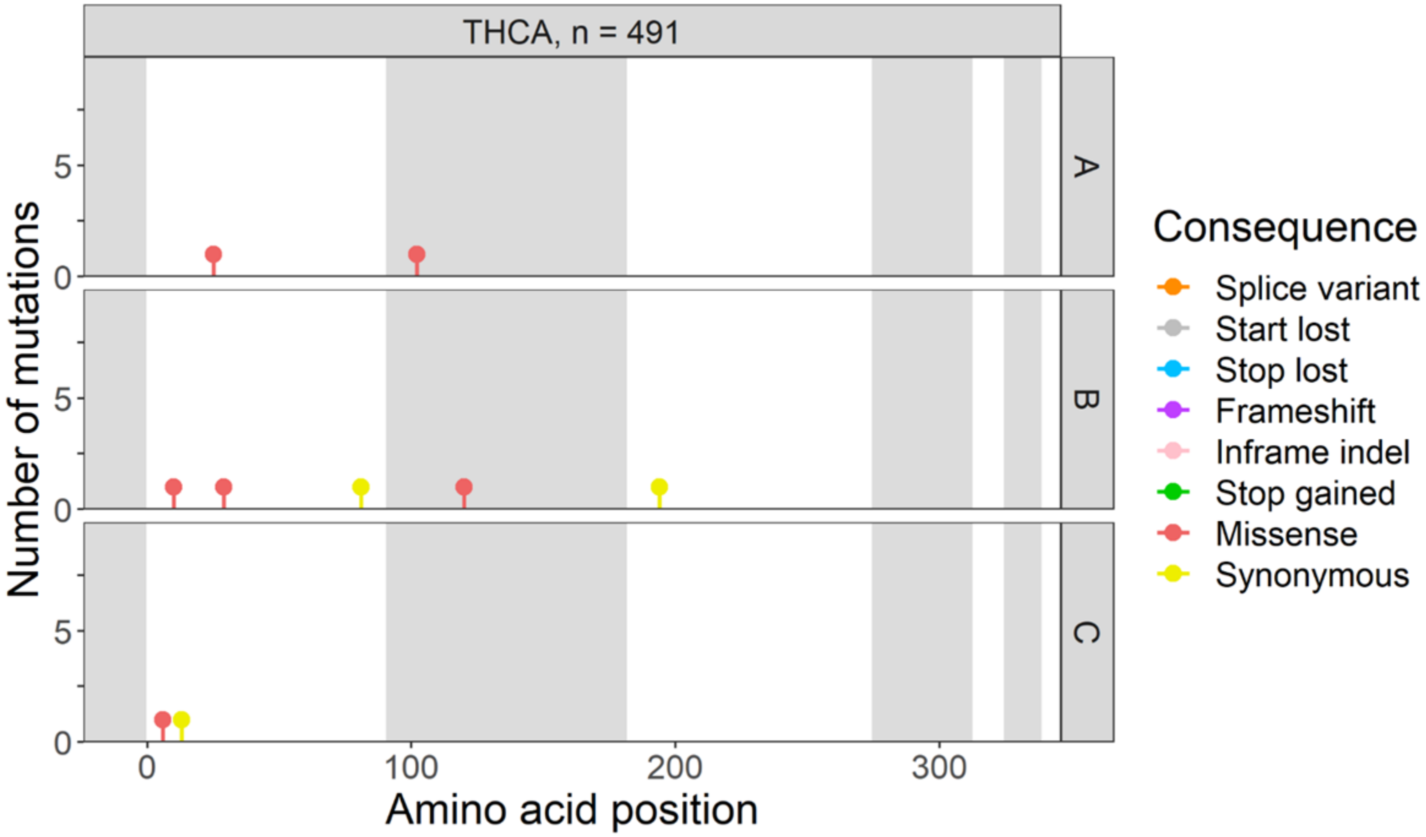
THCA, n = 491
A
B
C
Number of mutations
Amino acid position
Consequence
Splice variant
Start lost
Stop lost
Frameshift
Inframe indel
Stop gained
Missense
Synonymous

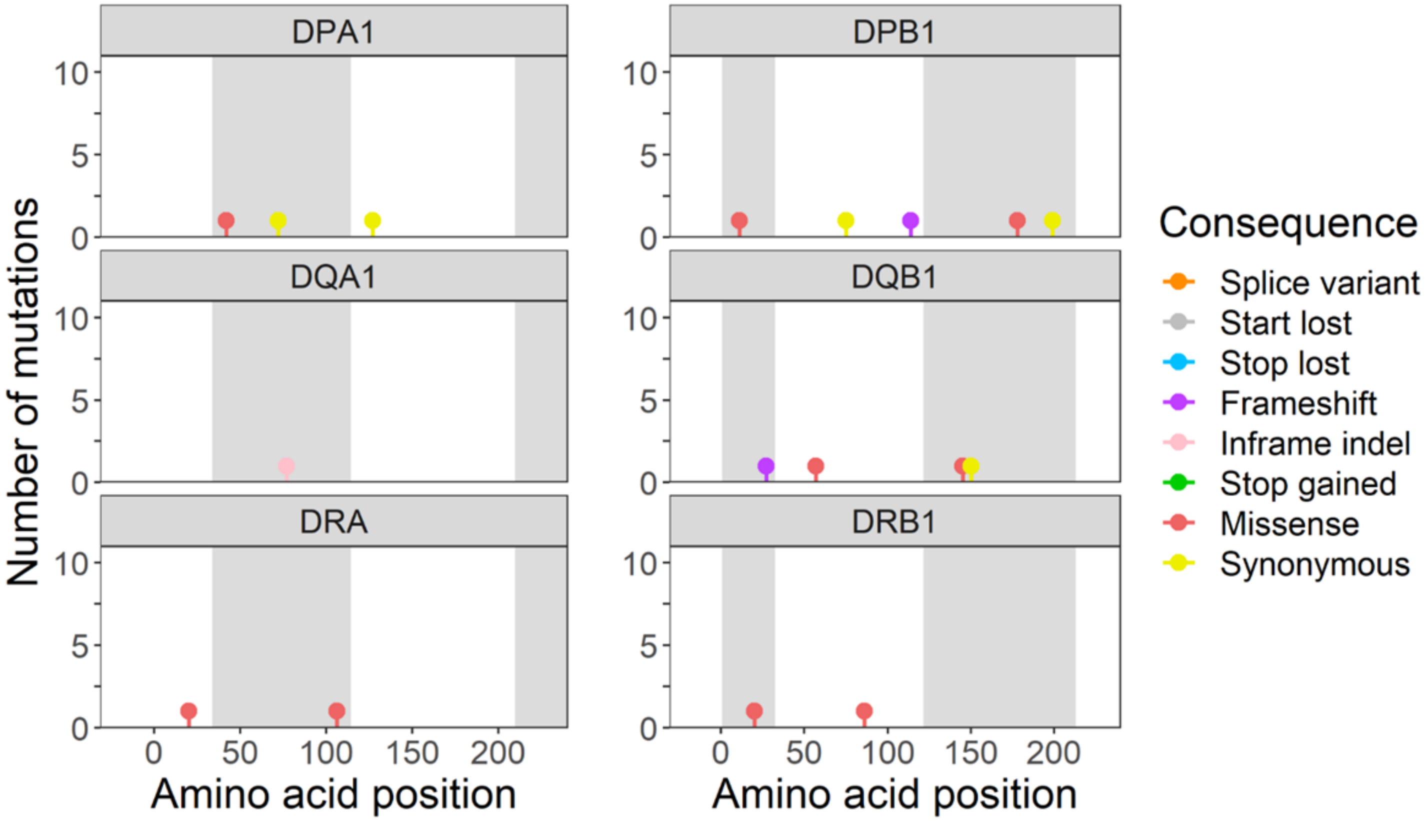
DPA1
DPB1
DQA1
DQB1
DRA
DRB1
Number of mutations
Amino acid position
Amino acid position
Consequence
Splice variant
Start lost
Stop lost
Frameshift
Inframe indel
Stop gained
Missense
Synonymous

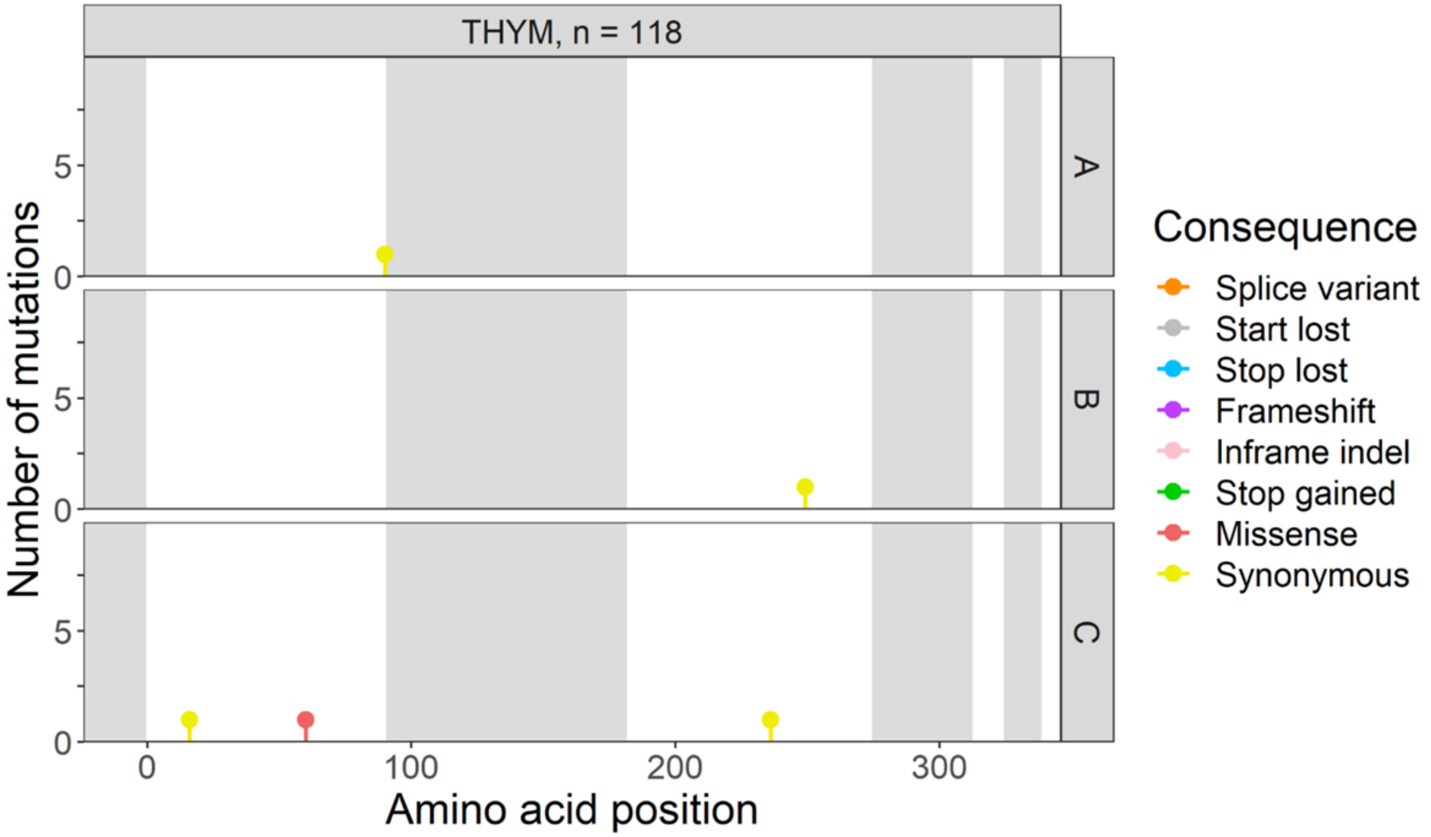
THYM, n = 118
A
B
C
Number of mutations
0
5
0
100
200
300
Amino acid position
Consequence
Splice variant
Start lost
Stop lost
Frameshift
Inframe indel
Stop gained
Missense
Synonymous

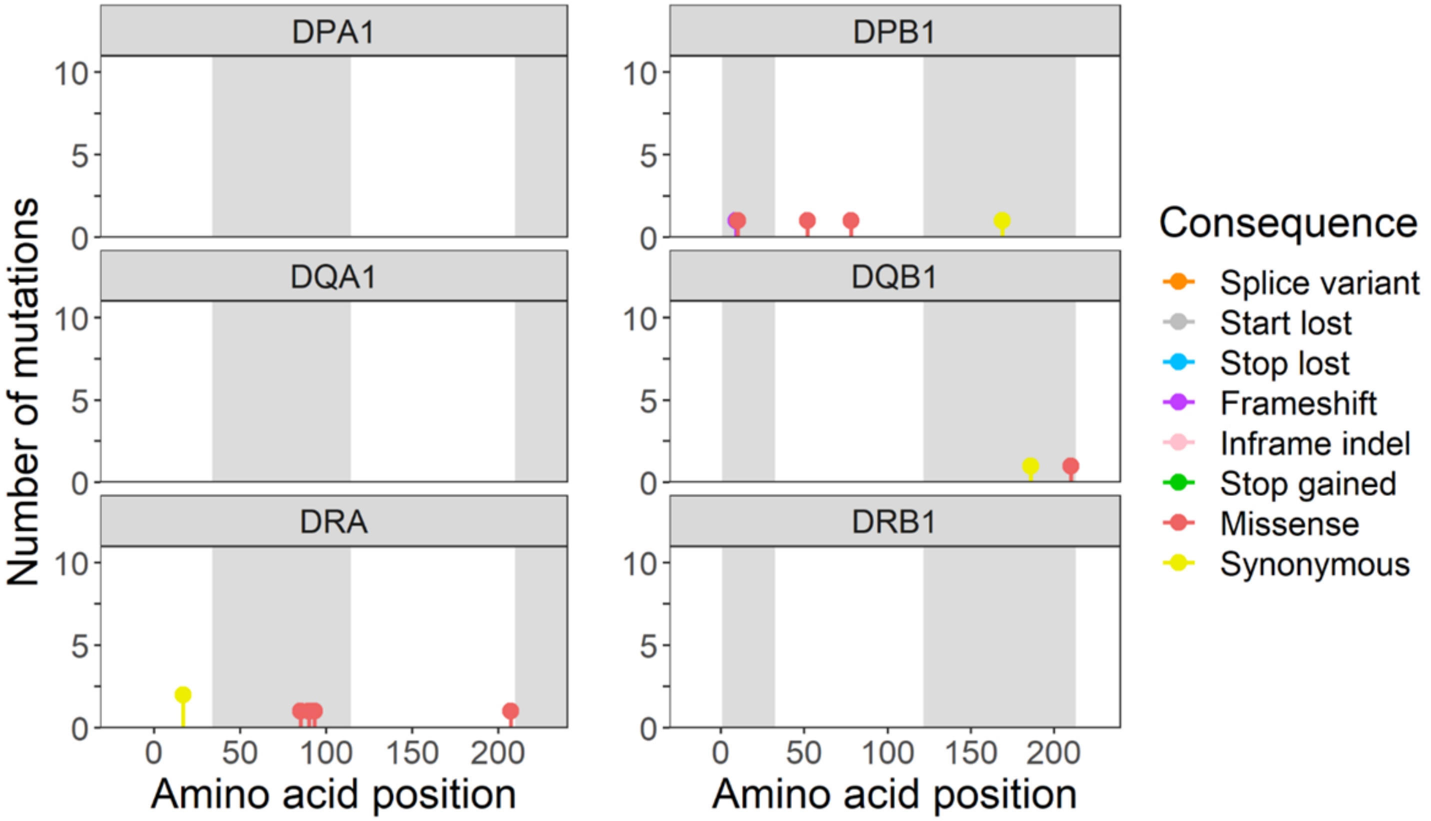
DPA1
DPB1
DQA1
DQB1
DRA
DRB1
Number of mutations
0
5
10
0
50
100
150
200
Amino acid position
Consequence
Splice variant
Start lost
Stop lost
Frameshift
Inframe indel
Stop gained
Missense
Synonymous

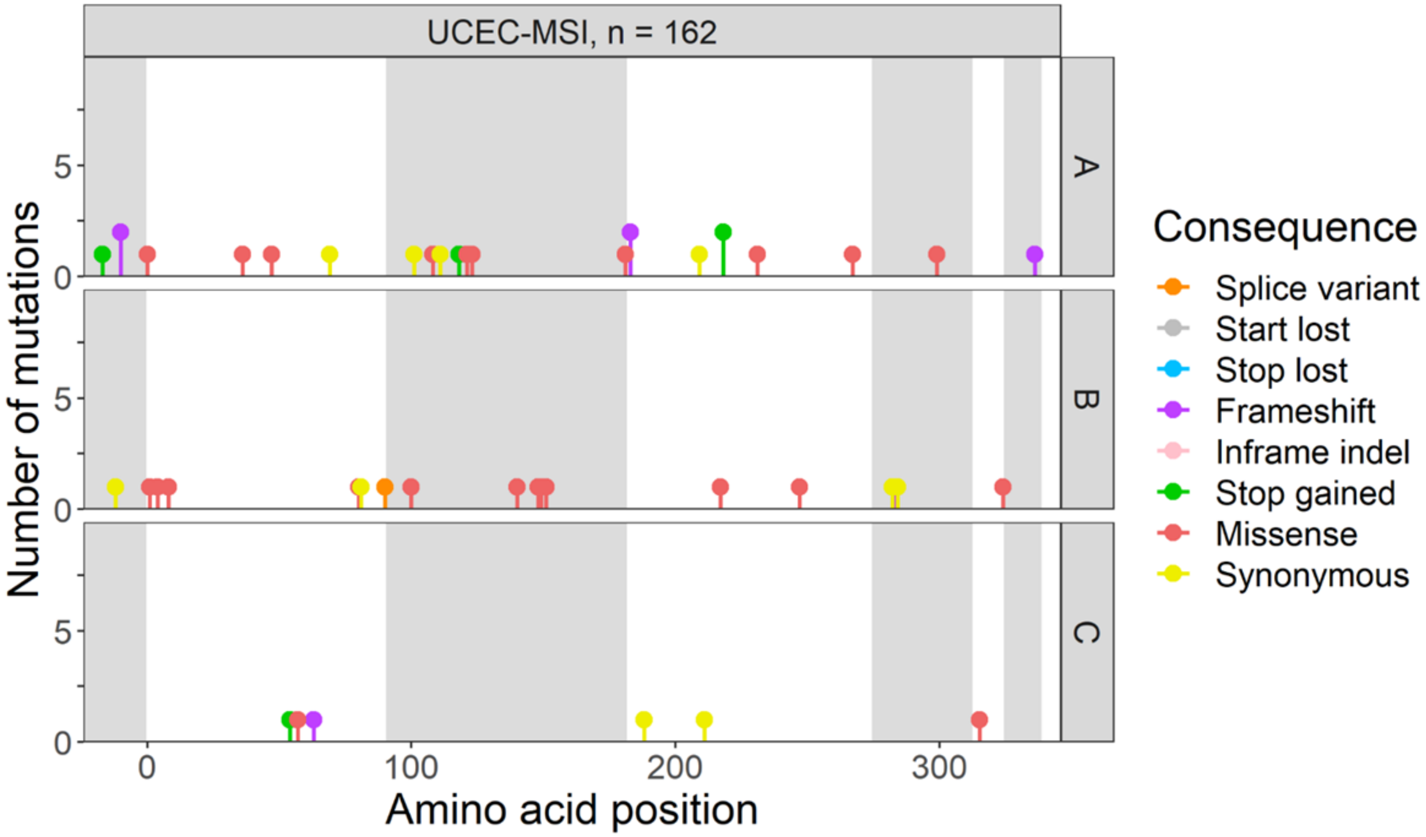
UCEC-MSI, n = 162
A
B
C
Number of mutations
Amino acid position
0
100
200
300
0
5
Consequence
Splice variant
Start lost
Stop lost
Frameshift
Inframe indel
Stop gained
Missense
Synonymous

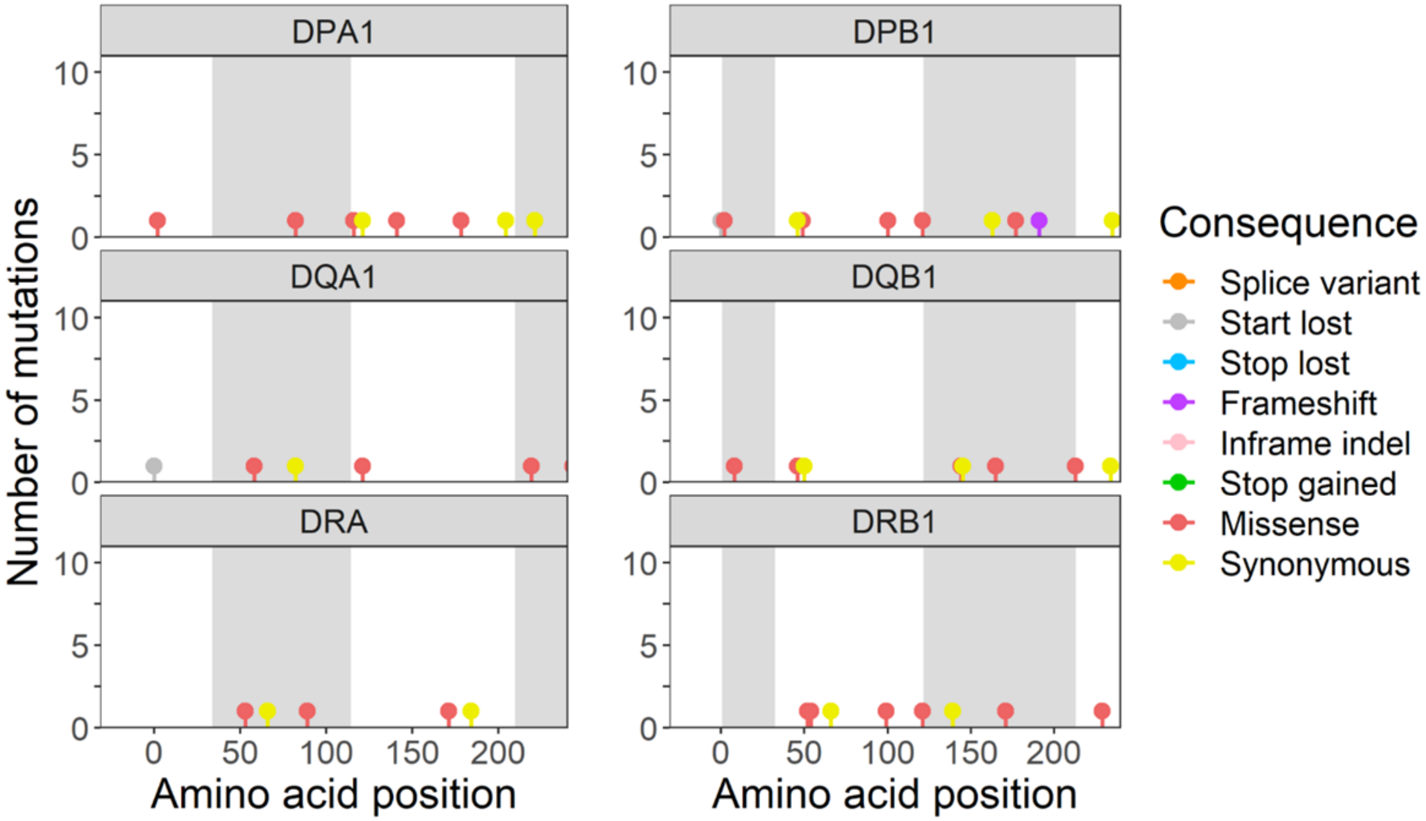
DPA1
DPB1
DQA1
DQB1
DRA
DRB1
Number of mutations
Amino acid position
0
50
100
150
200
0
5
10
Consequence
Splice variant
Start lost
Stop lost
Frameshift
Inframe indel
Stop gained
Missense
Synonymous

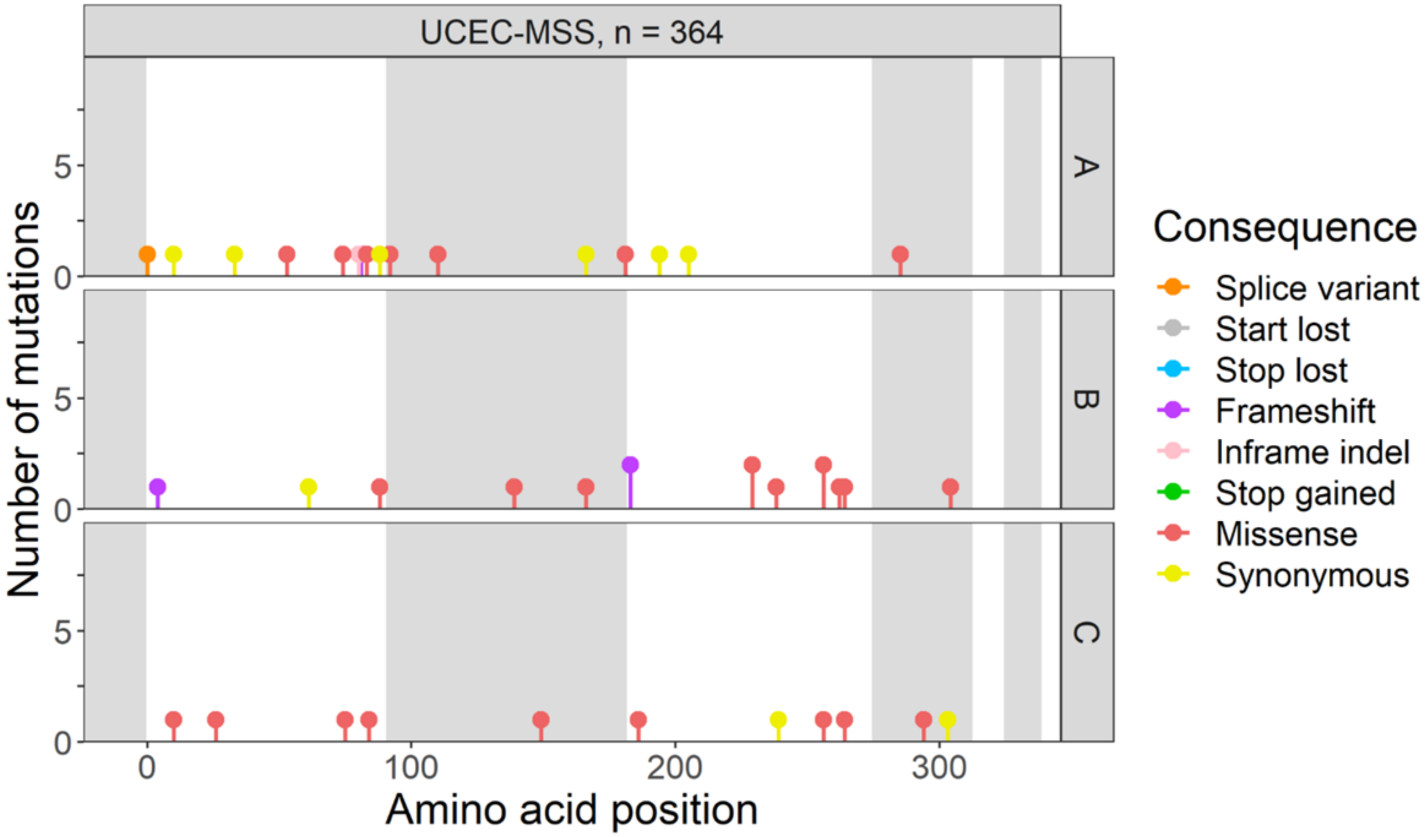
UCEC-MSS, n = 364
A
B
C
Number of mutations
Amino acid position
0
5
0
100
200
300
Consequence
Splice variant
Start lost
Stop lost
Frameshift
Inframe indel
Stop gained
Missense
Synonymous

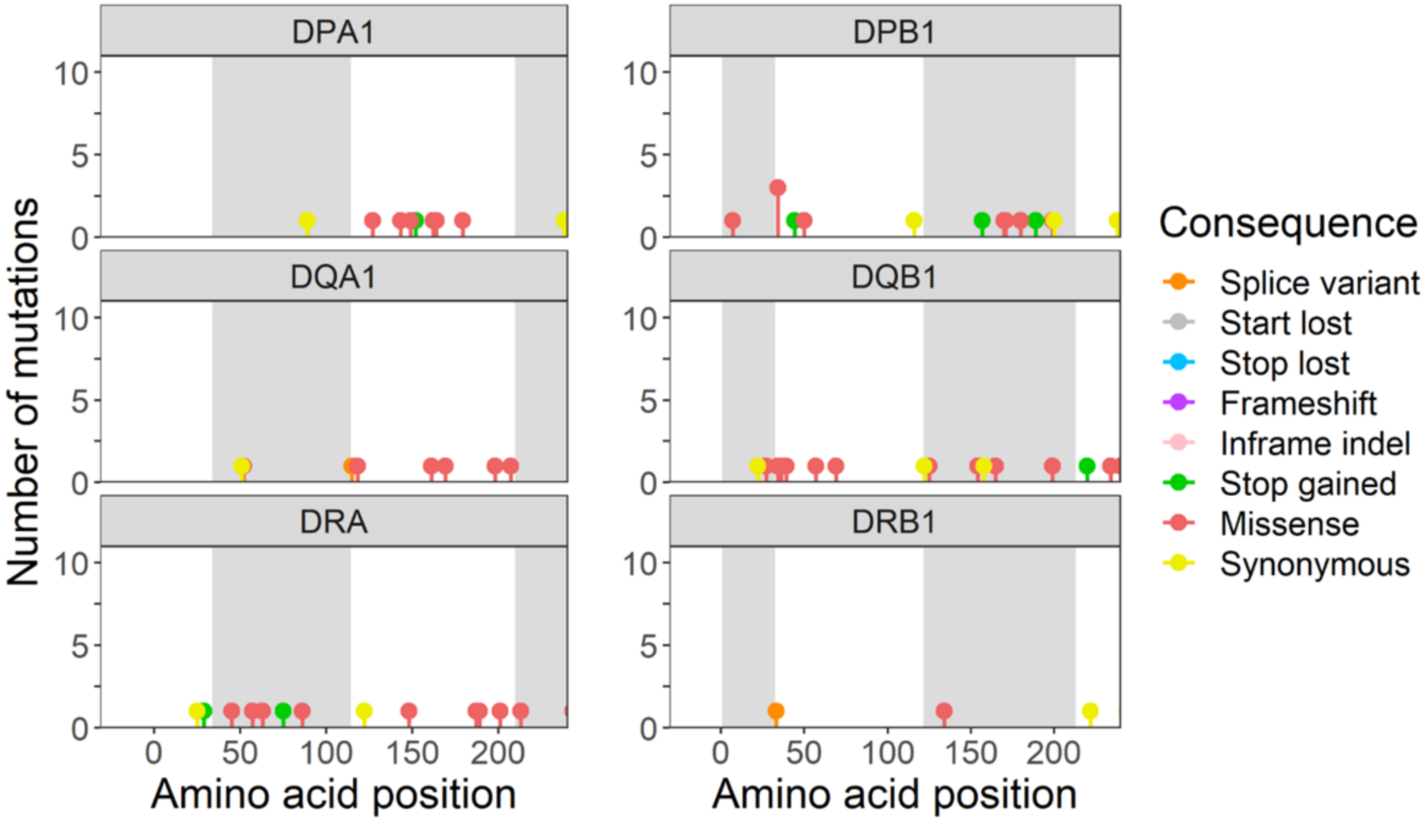
DPA1
DPB1
DQA1
DQB1
DRA
DRB1
Number of mutations
Amino acid position
0
5
10
0
50
100
150
200
Consequence
Splice variant
Start lost
Stop lost
Frameshift
Inframe indel
Stop gained
Missense
Synonymous

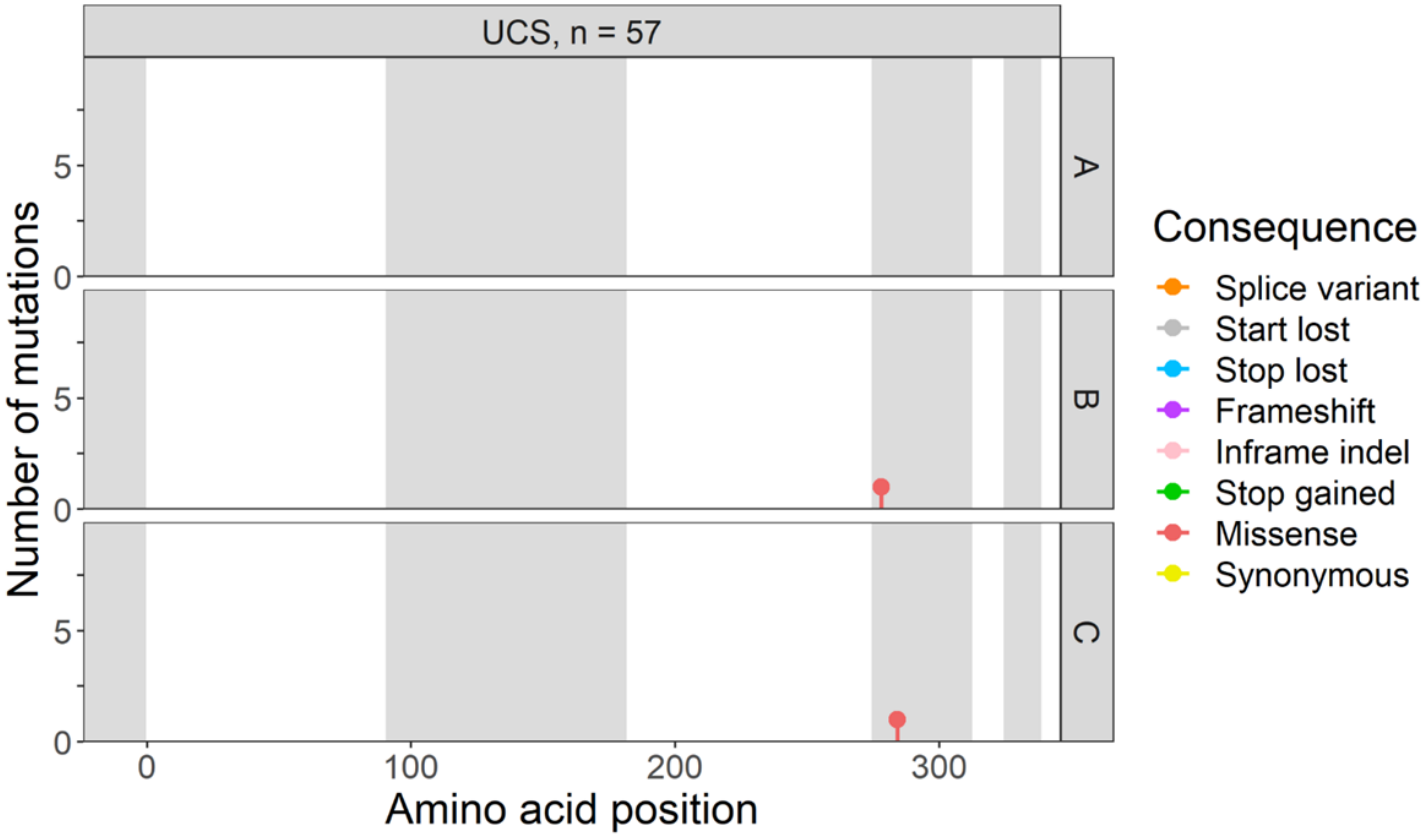
UCS, n = 57
A
B
C
Number of mutations
Amino acid position
0
100
200
300
Consequence
Splice variant
Start lost
Stop lost
Frameshift
Inframe indel
Stop gained
Missense
Synonymous

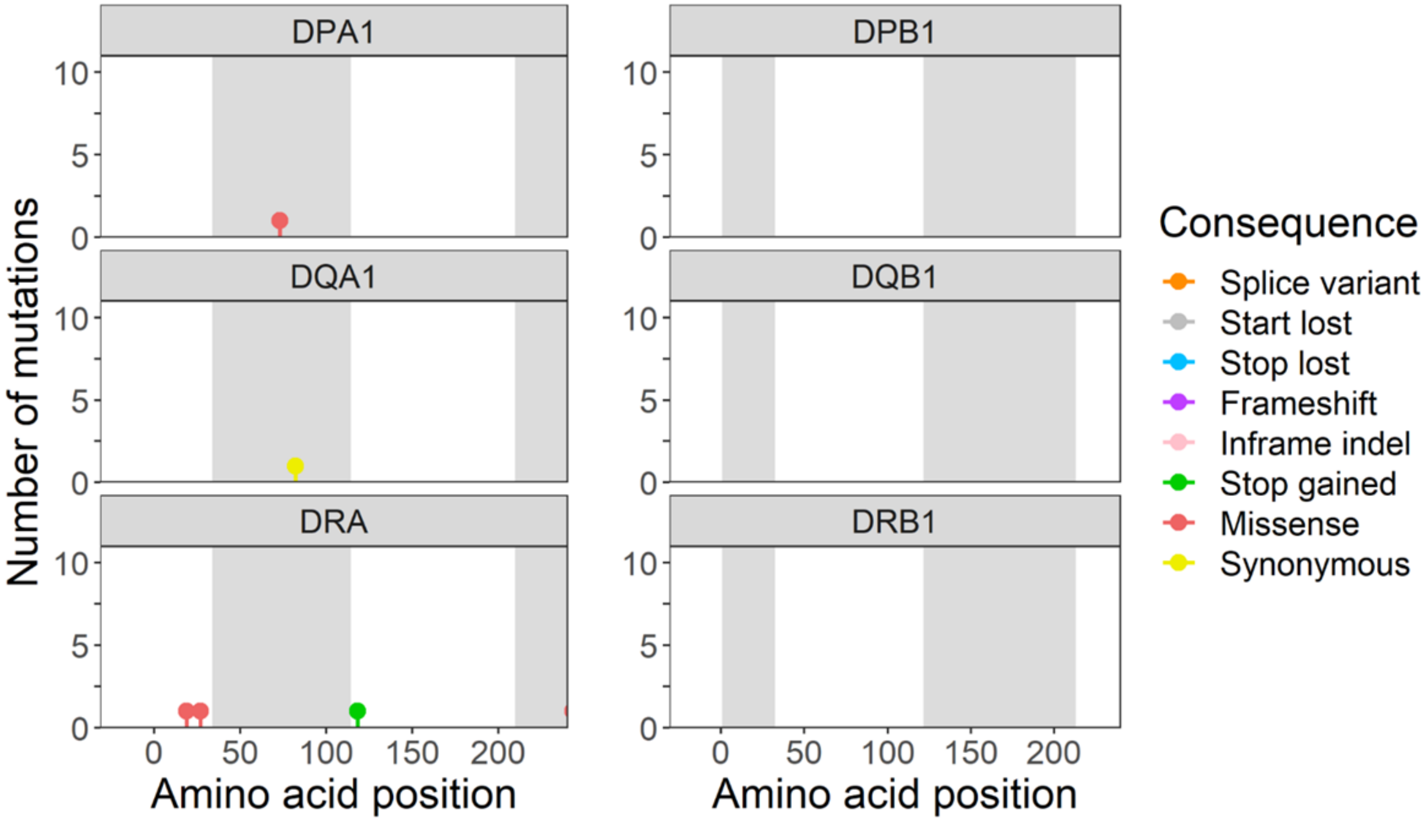
DPA1
DPB1
DQA1
DQB1
DRA
DRB1
Number of mutations
Amino acid position
Amino acid position
Consequence
Splice variant
Start lost
Stop lost
Frameshift
Inframe indel
Stop gained
Missense
Synonymous

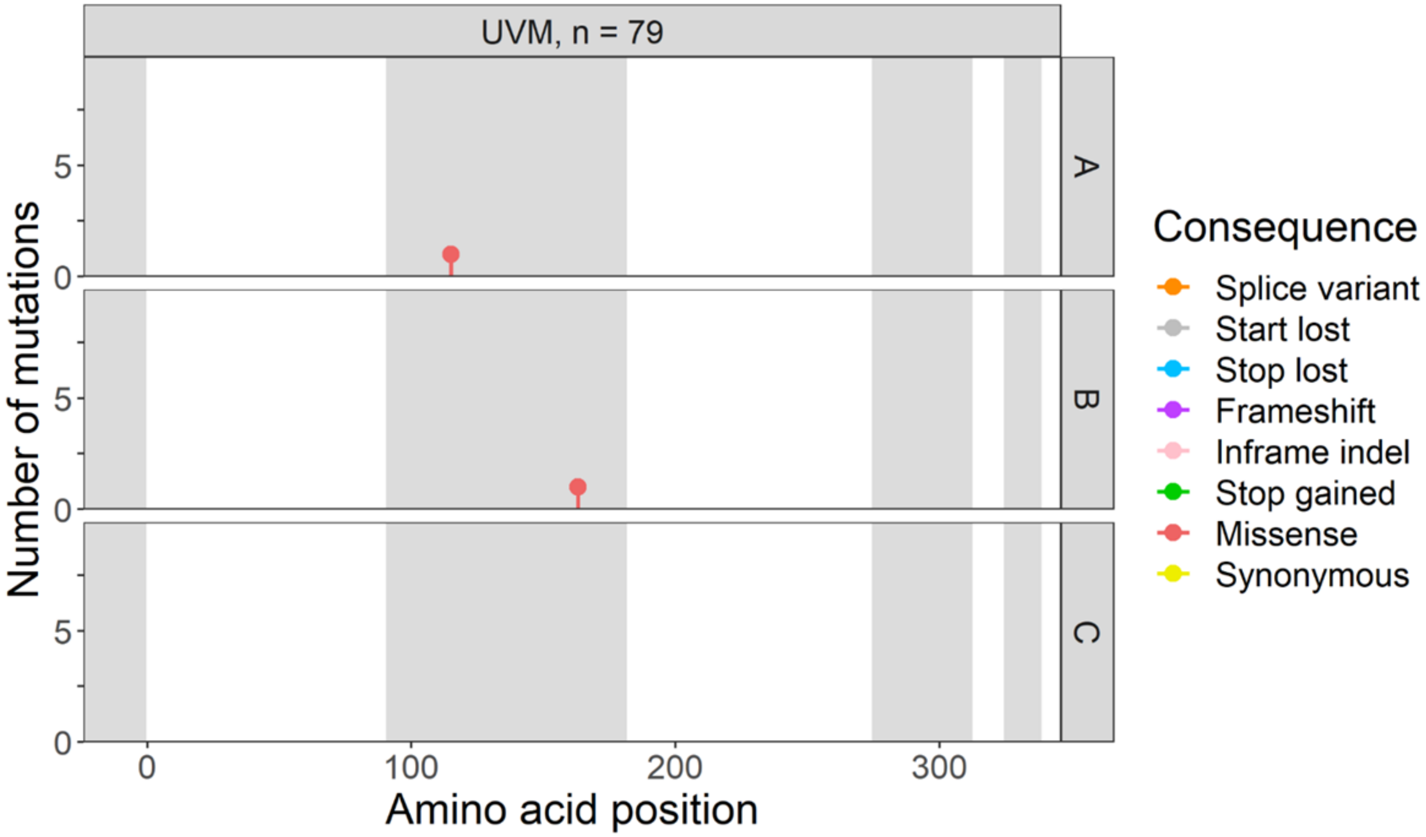
UVM, n = 79
A
B
C
Number of mutations
Amino acid position
0
100
200
300
Consequence
Splice variant
Start lost
Stop lost
Frameshift
Inframe indel
Stop gained
Missense
Synonymous

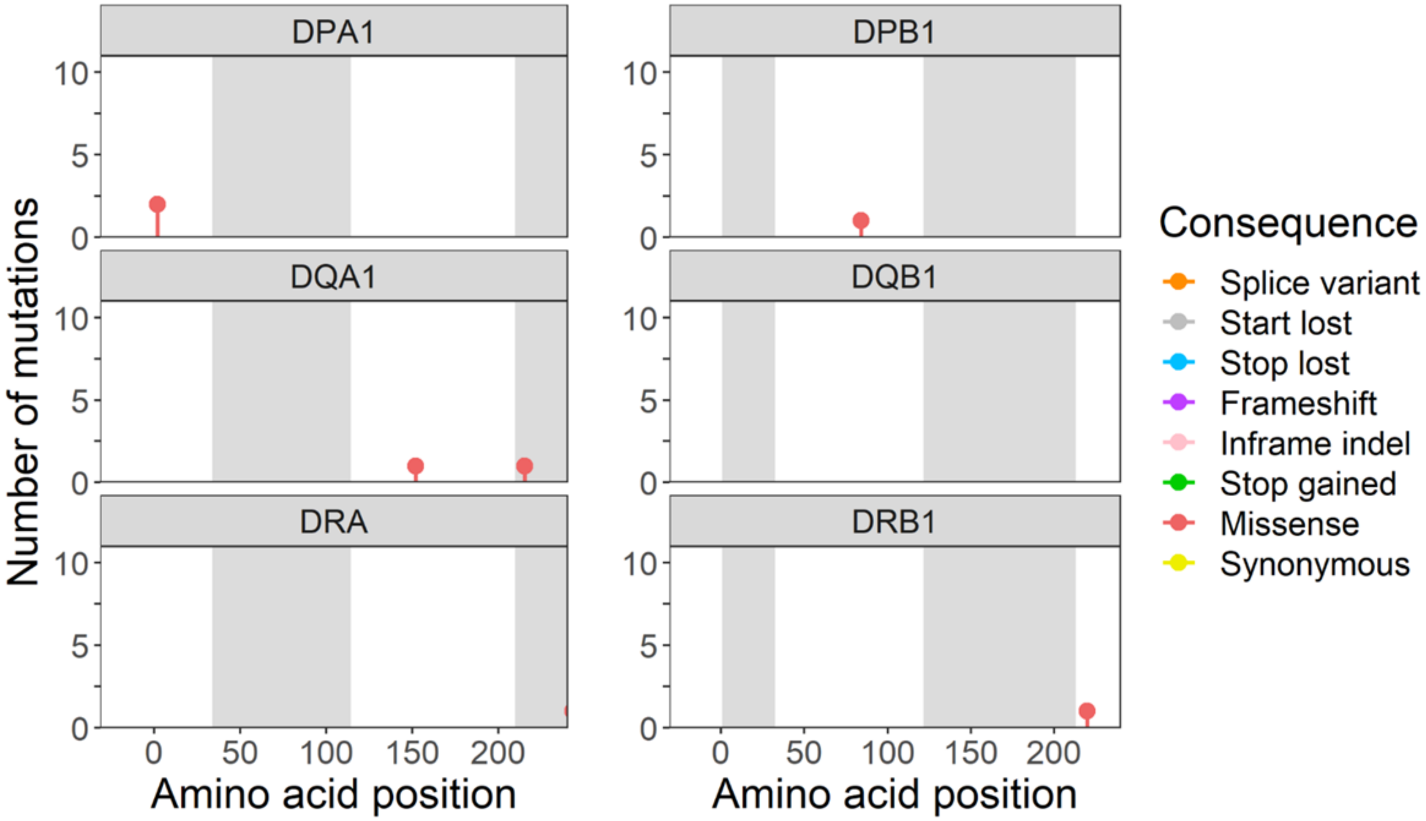
DPA1
DPB1
DQA1
DQB1
DRA
DRB1
Number of mutations
Amino acid position
0
50
100
150
200
Consequence
Splice variant
Start lost
Stop lost
Frameshift
Inframe indel
Stop gained
Missense
Synonymous

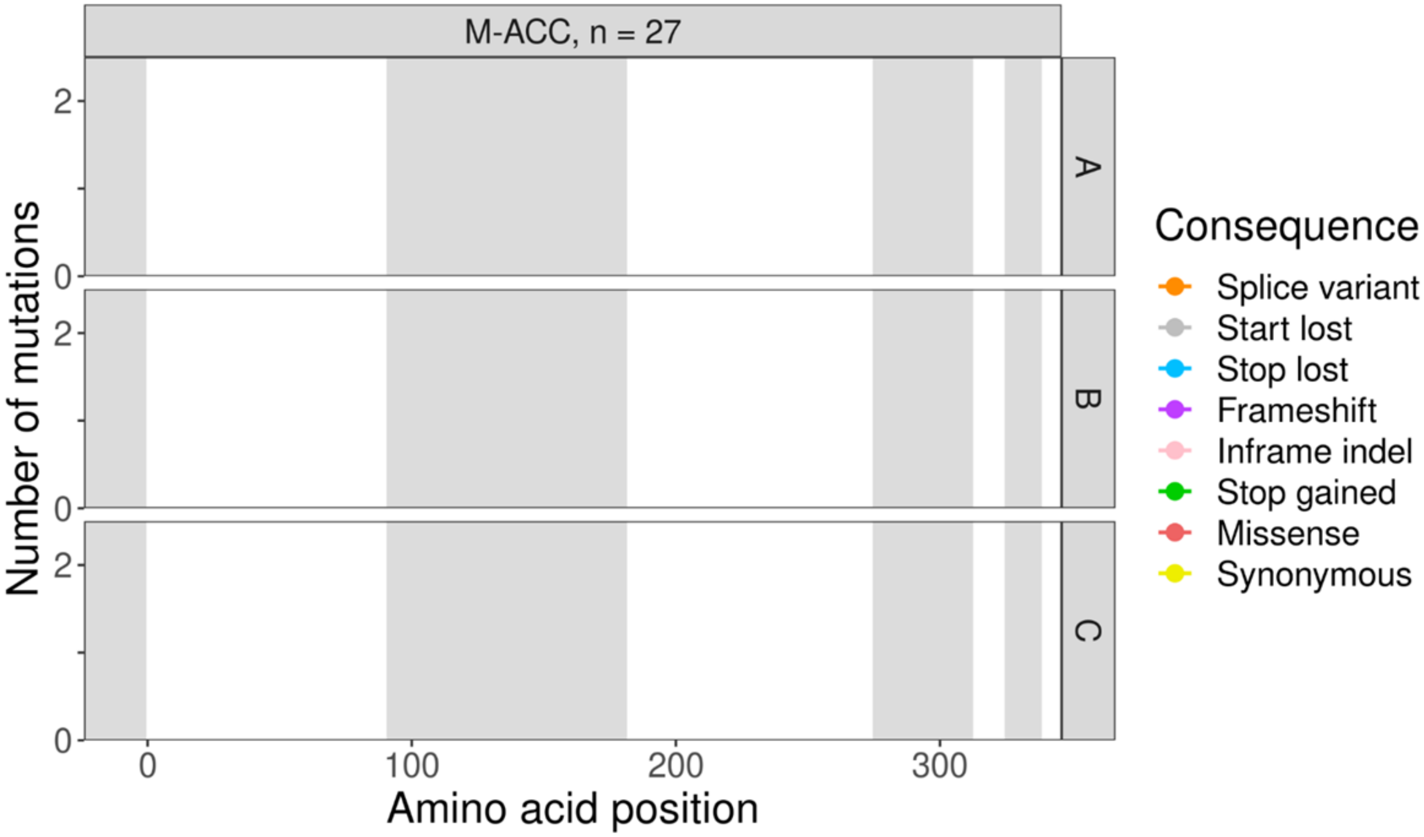
M-ACC, n = 27
A
B
C
Number of mutations
Amino acid position
0
100
200
300
Consequence
Splice variant
Start lost
Stop lost
Frameshift
Inframe indel
Stop gained
Missense
Synonymous

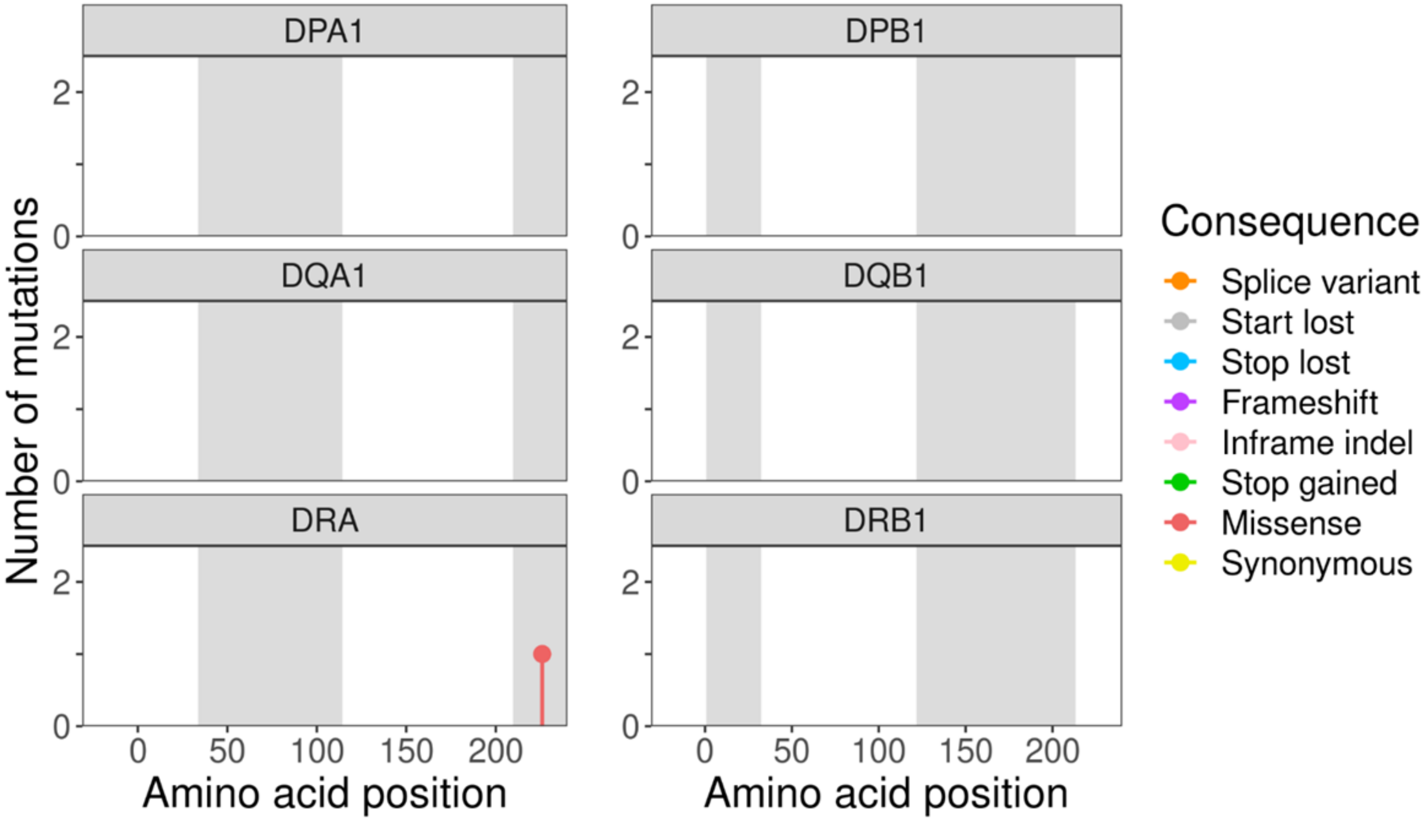
DPA1
DPB1
DQA1
DQB1
DRA
DRB1
Number of mutations
Amino acid position
0
50
100
150
200
Consequence
Splice variant
Start lost
Stop lost
Frameshift
Inframe indel
Stop gained
Missense
Synonymous

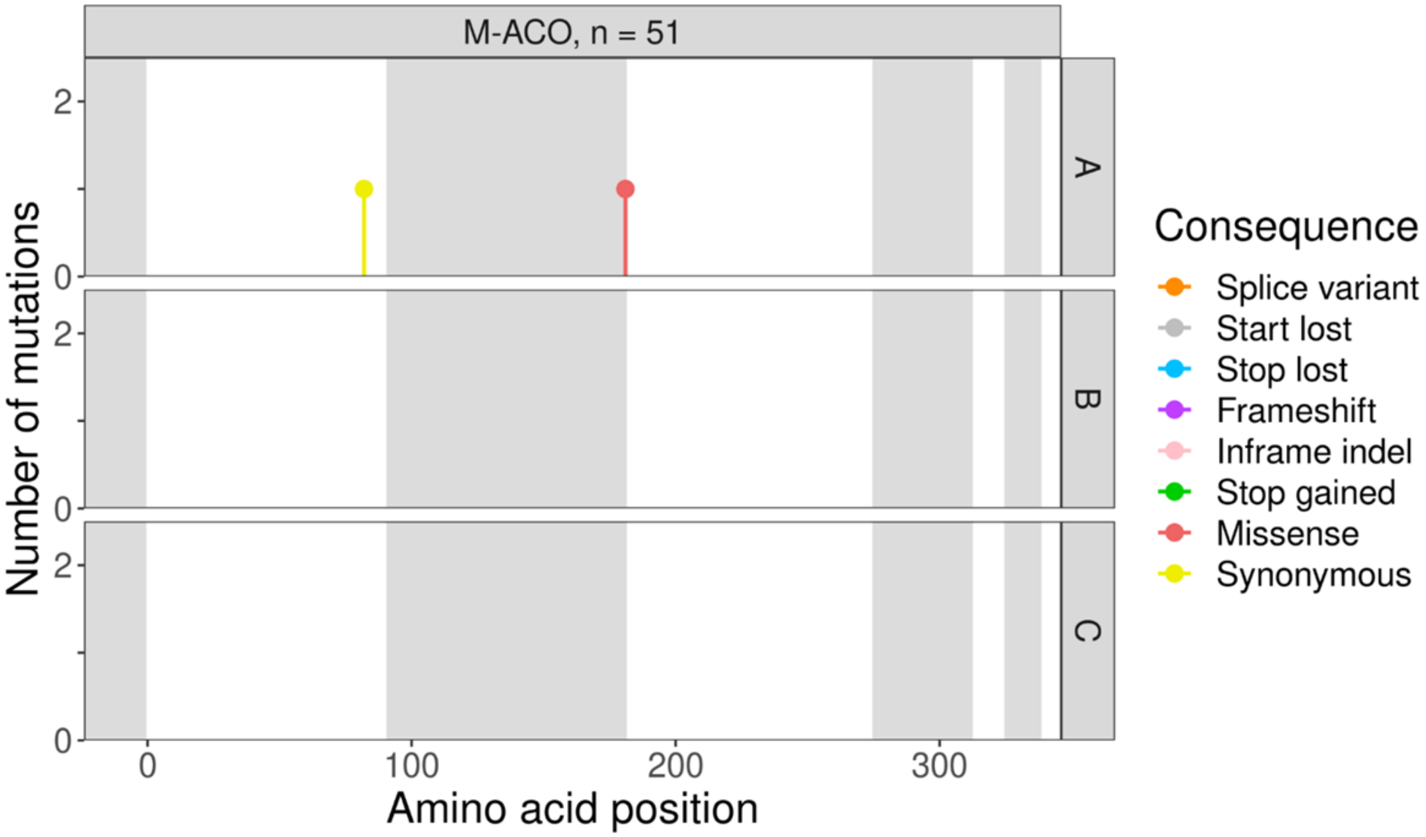
M-ACO, n = 51
A
B
C
Number of mutations
Amino acid position
Consequence
Splice variant
Start lost
Stop lost
Frameshift
Inframe indel
Stop gained
Missense
Synonymous

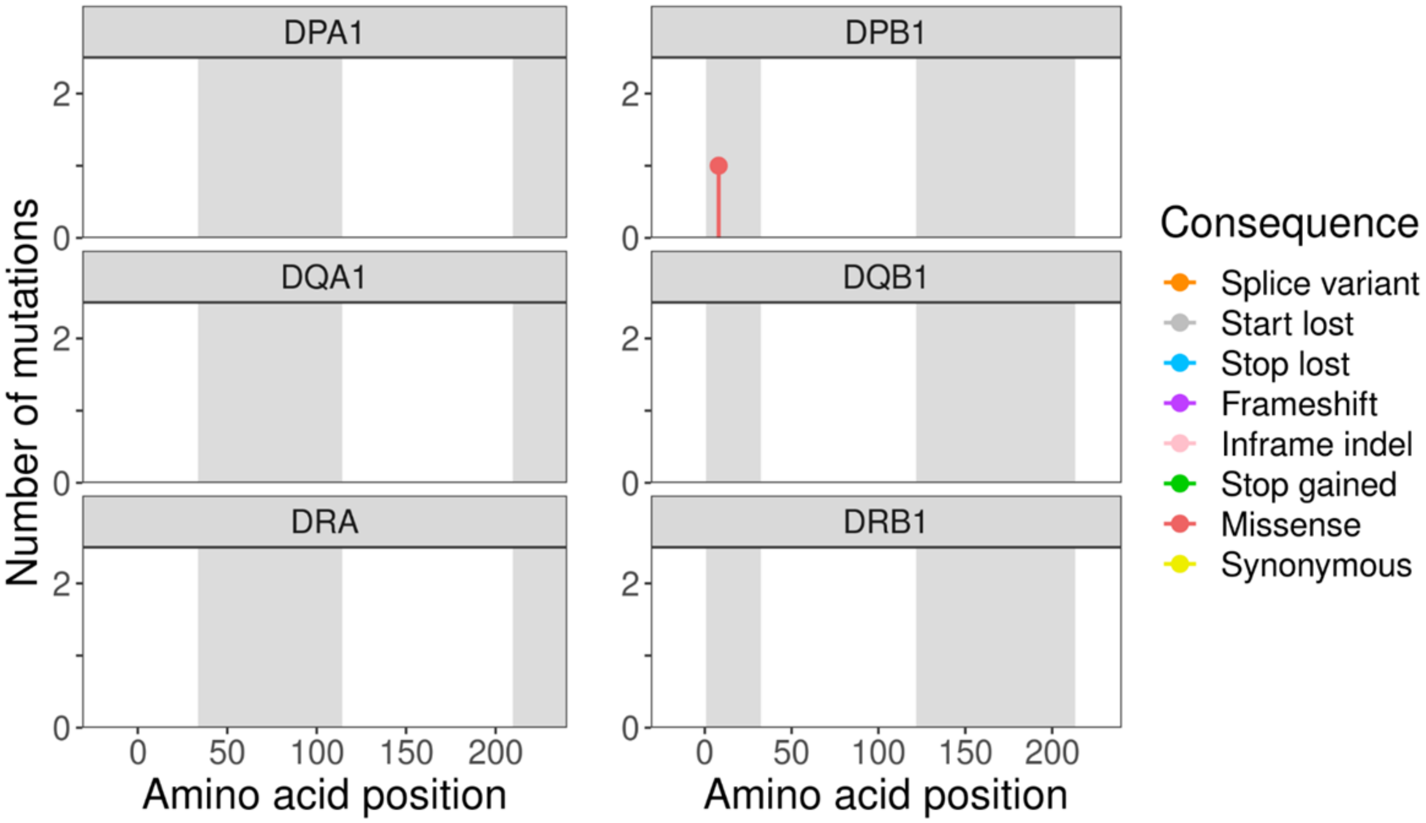
DPA1
DPB1
DQA1
DQB1
DRA
DRB1
Number of mutations
Amino acid position
Amino acid position
Consequence
Splice variant
Start lost
Stop lost
Frameshift
Inframe indel
Stop gained
Missense
Synonymous

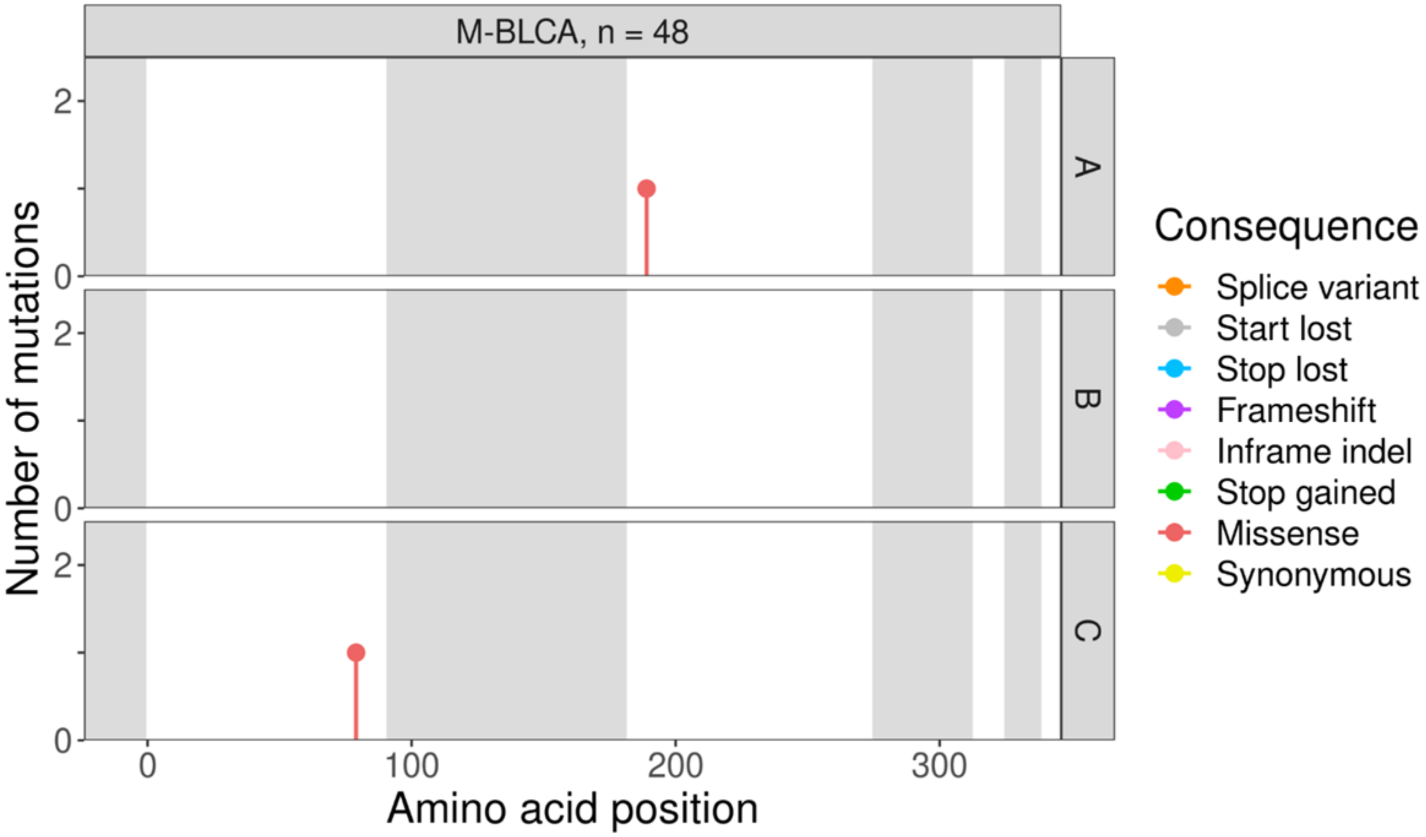
M-BLCA, n = 48
A
B
C
Number of mutations
Amino acid position
Consequence
Splice variant
Start lost
Stop lost
Frameshift
Inframe indel
Stop gained
Missense
Synonymous

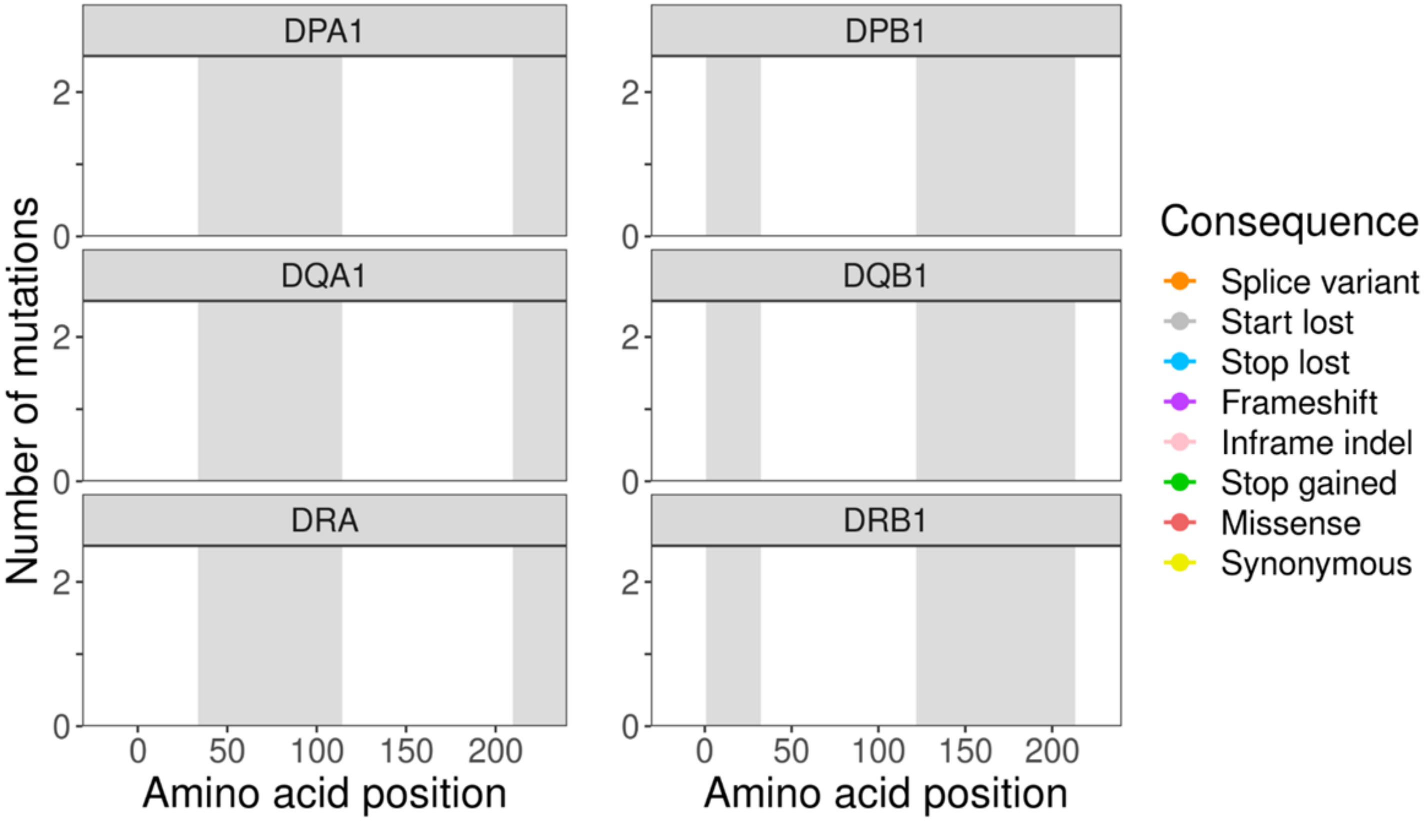
DPA1
DPB1
DQA1
DQB1
DRA
DRB1
Number of mutations
Amino acid position
Amino acid position
Consequence
Splice variant
Start lost
Stop lost
Frameshift
Inframe indel
Stop gained
Missense
Synonymous

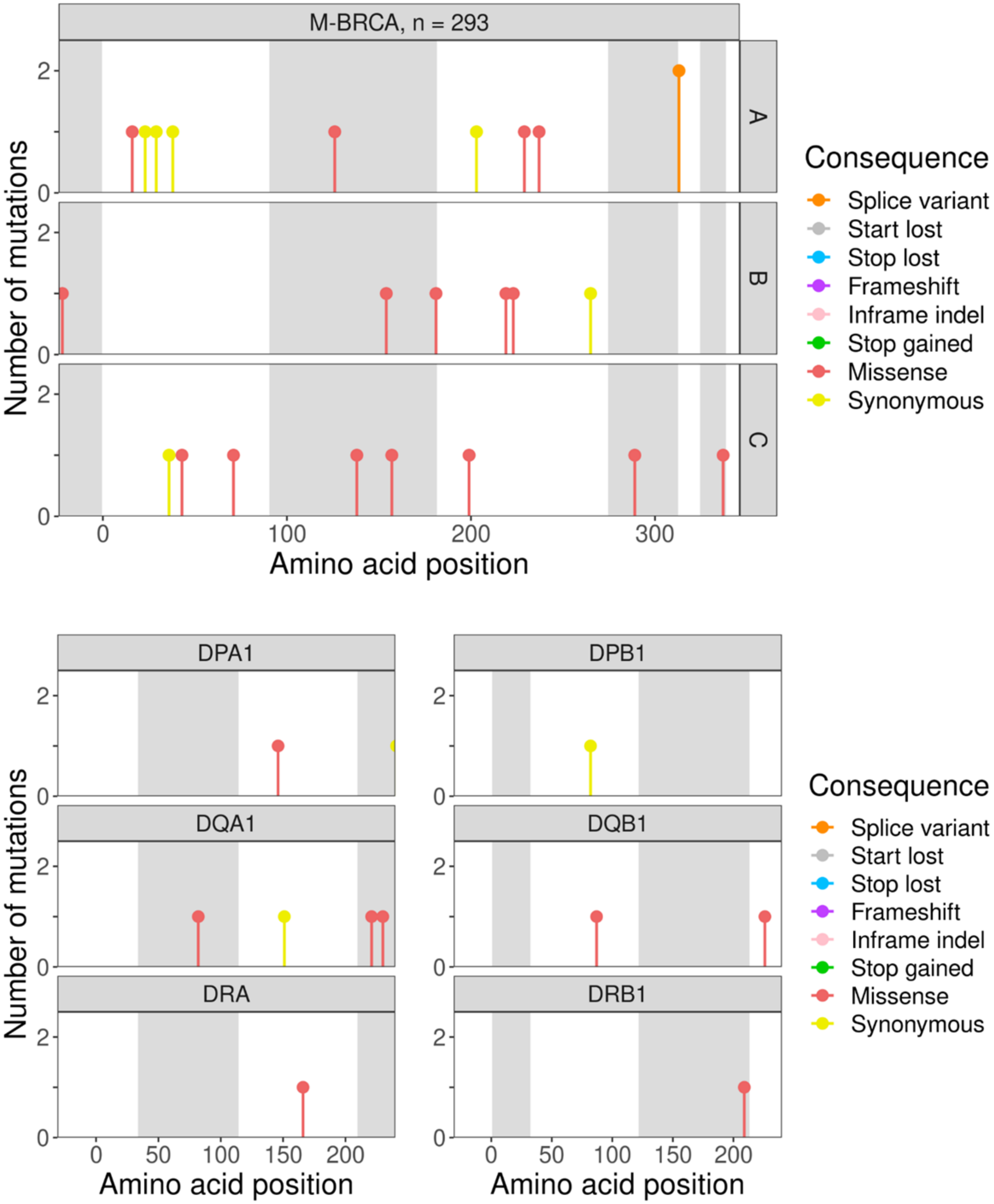

M-BRCA, n = 293
A
B
C
Number of mutations
Amino acid position
Consequence
Splice variant
Start lost
Stop lost
Frameshift
Inframe indel
Stop gained
Missense
Synonymous
DPA1
DPB1
DQA1
DQB1
DRA
DRB1

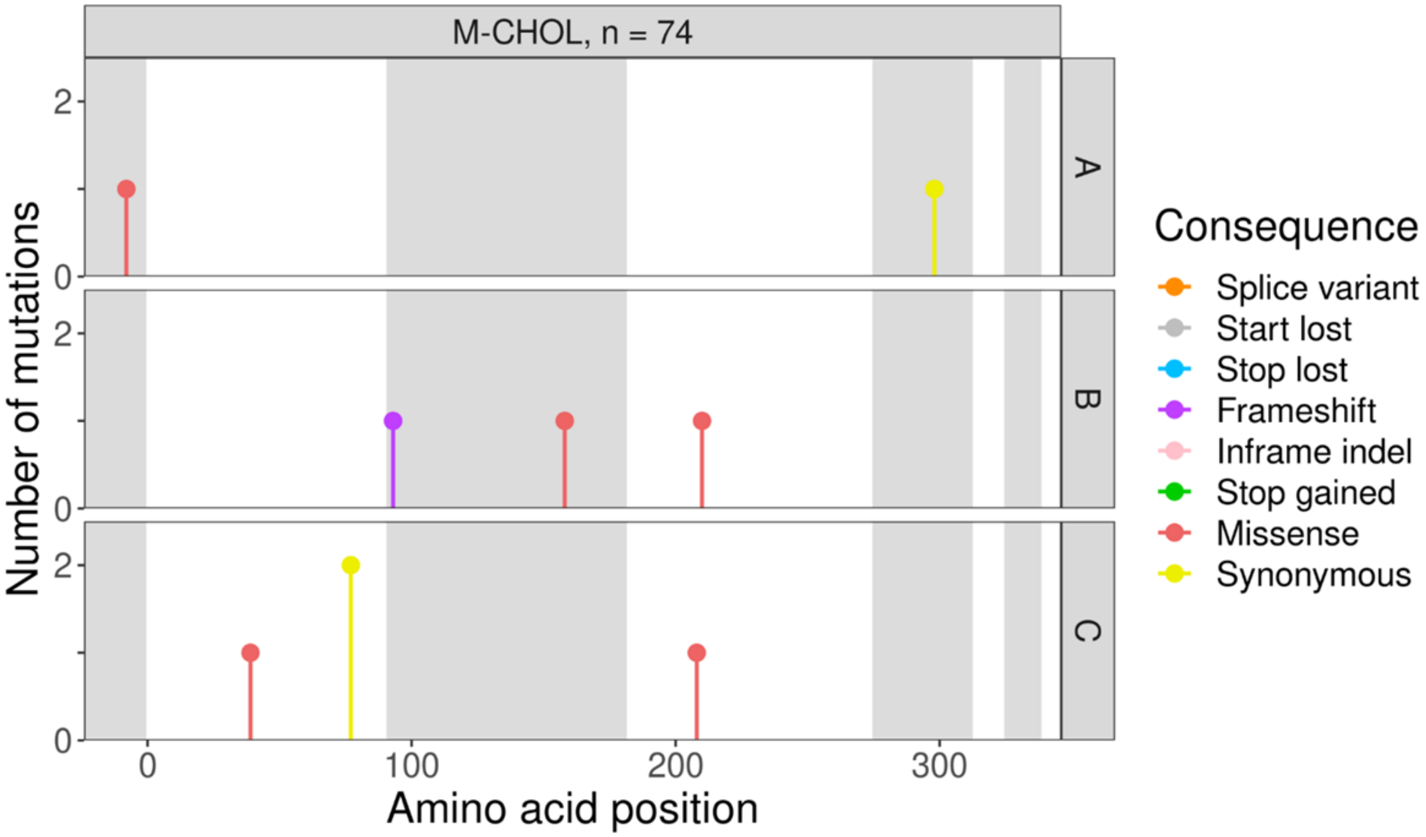

M-CHOL, n = 74
A
B
C
Number of mutations
Amino acid position
0
2
0
100
200
300
Consequence
Splice variant
Start lost
Stop lost
Frameshift
Inframe indel
Stop gained
Missense
Synonymous


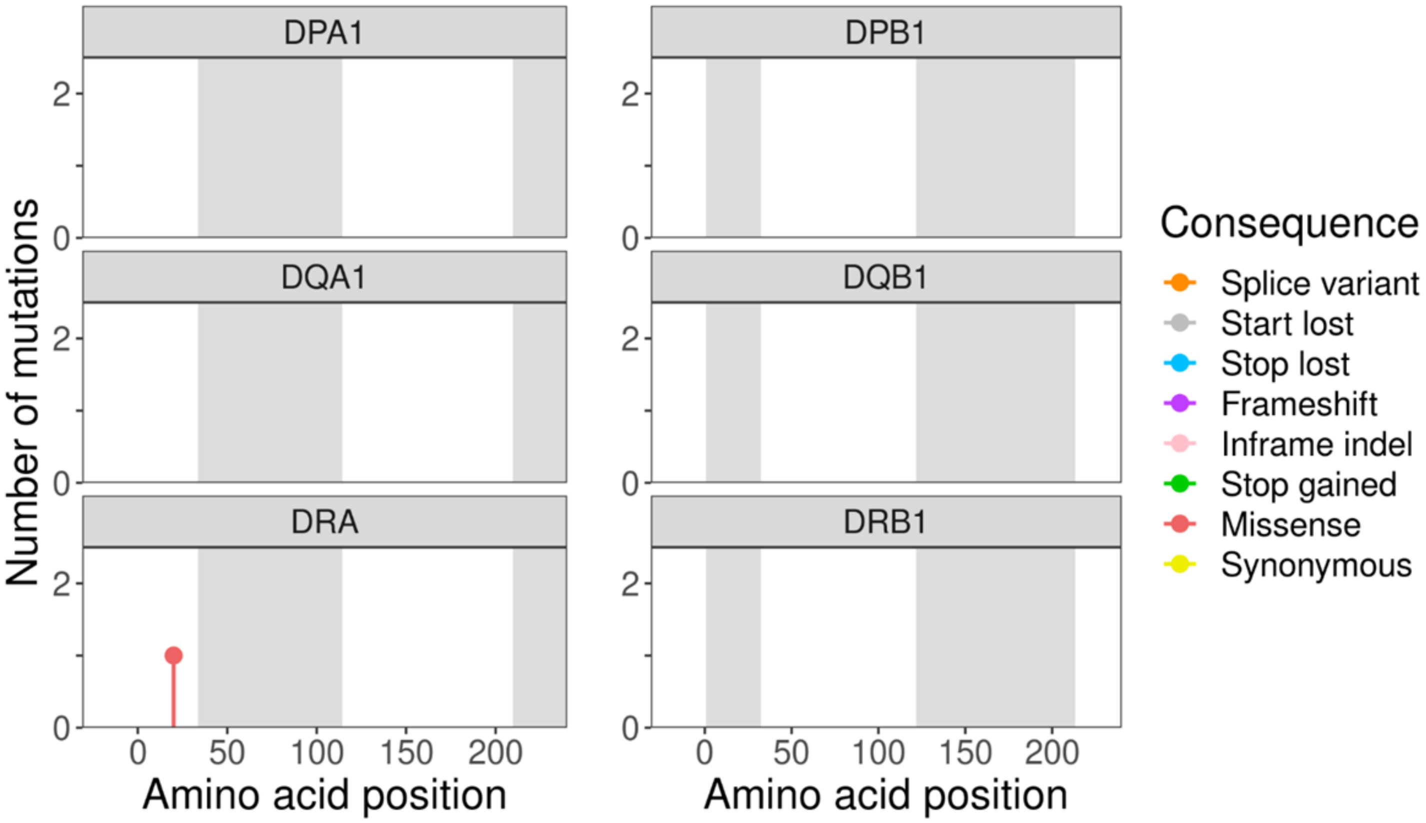

DPA1
DPB1
DQA1
DQB1
DRA
DRB1
Number of mutations
Amino acid position
0
2
0
50
100
150
200
Consequence
Splice variant
Start lost
Stop lost
Frameshift
Inframe indel
Stop gained
Missense
Synonymous

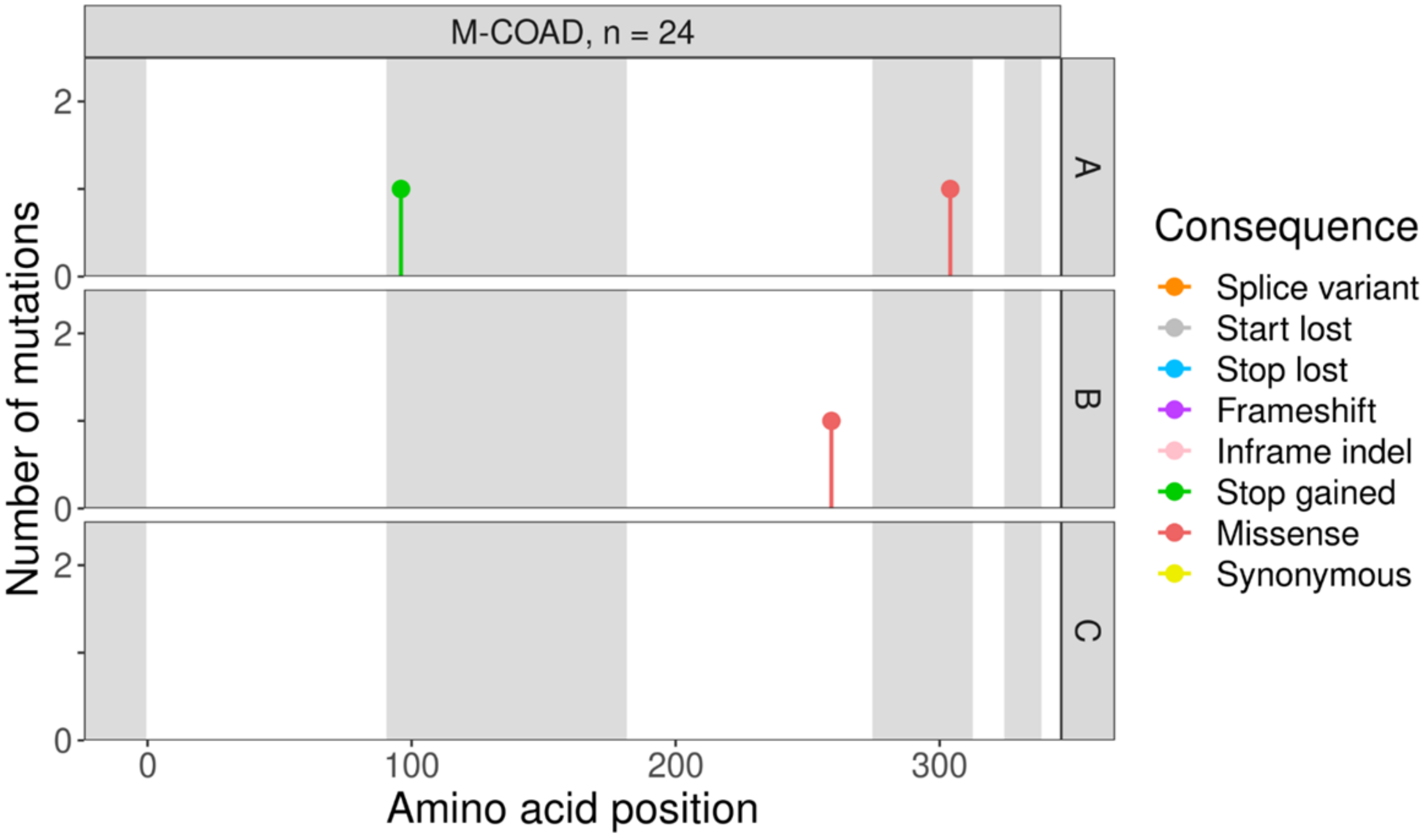
M-COAD, n = 24
A
B
C
Number of mutations
Amino acid position
Consequence
Splice variant
Start lost
Stop lost
Frameshift
Inframe indel
Stop gained
Missense
Synonymous

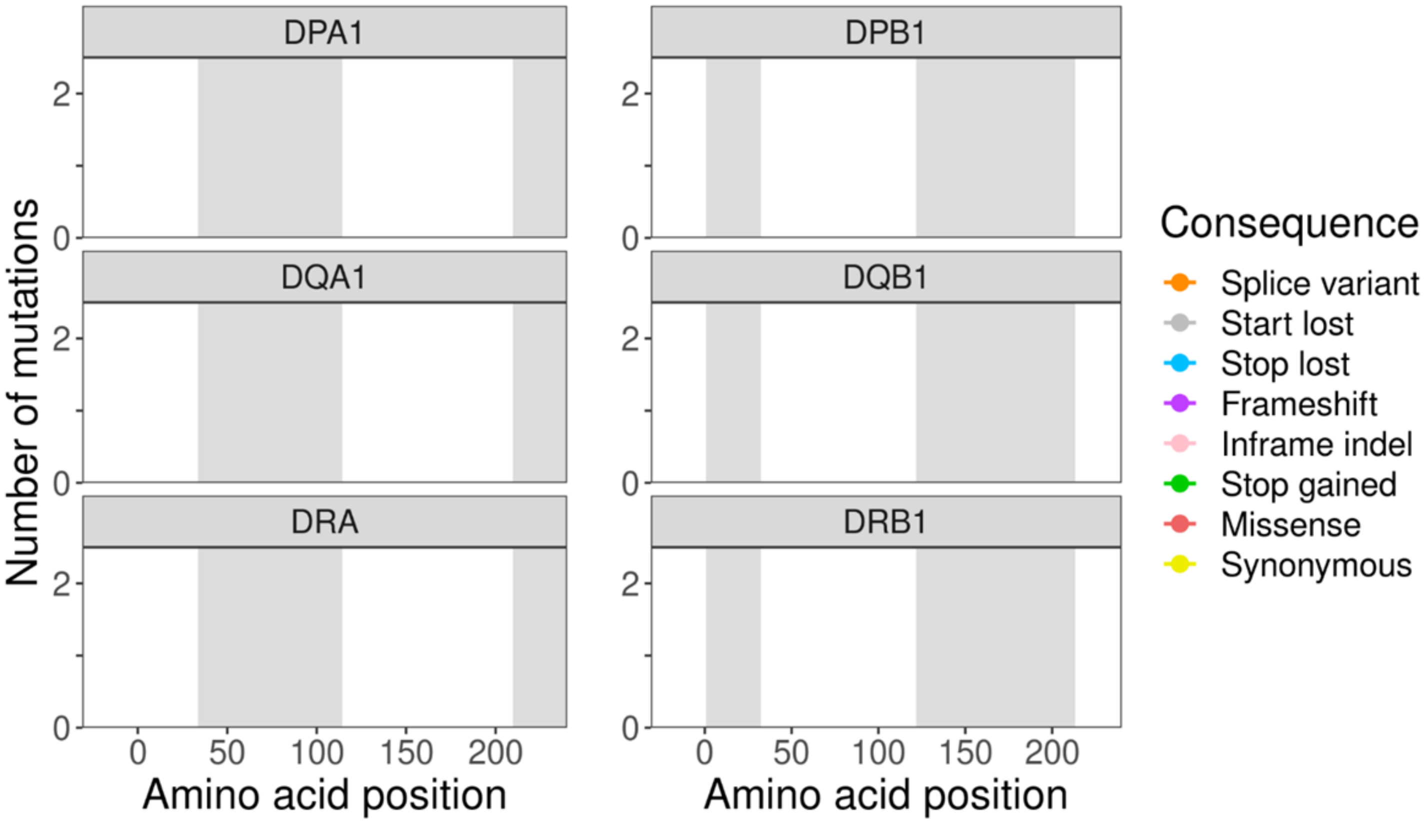
DPA1
DPB1
DQA1
DQB1
DRA
DRB1
Number of mutations
Amino acid position
Amino acid position
Consequence
Splice variant
Start lost
Stop lost
Frameshift
Inframe indel
Stop gained
Missense
Synonymous

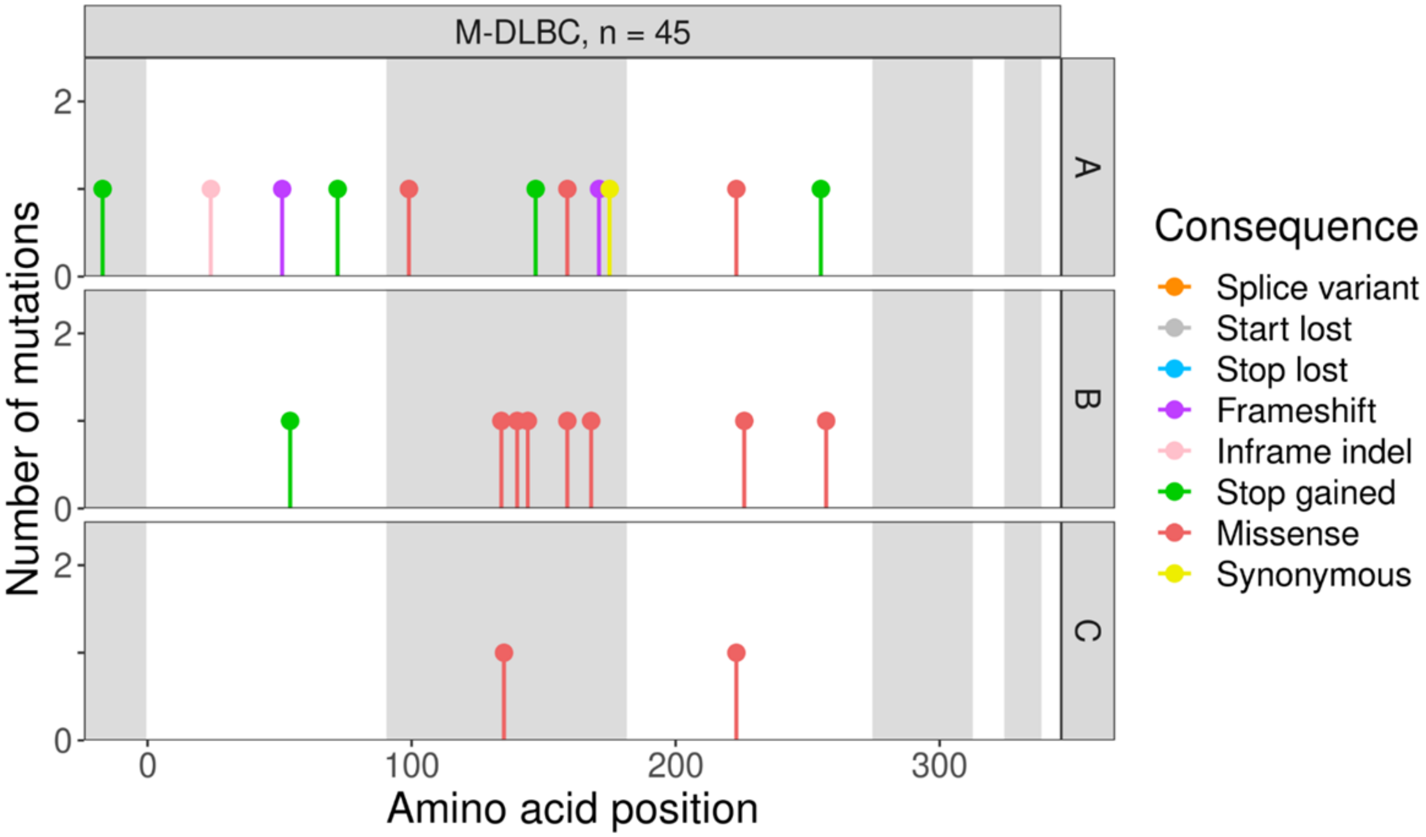
M-DLBC, n = 45
A
B
C
Number of mutations
Amino acid position
Consequence
Splice variant
Start lost
Stop lost
Frameshift
Inframe indel
Stop gained
Missense
Synonymous

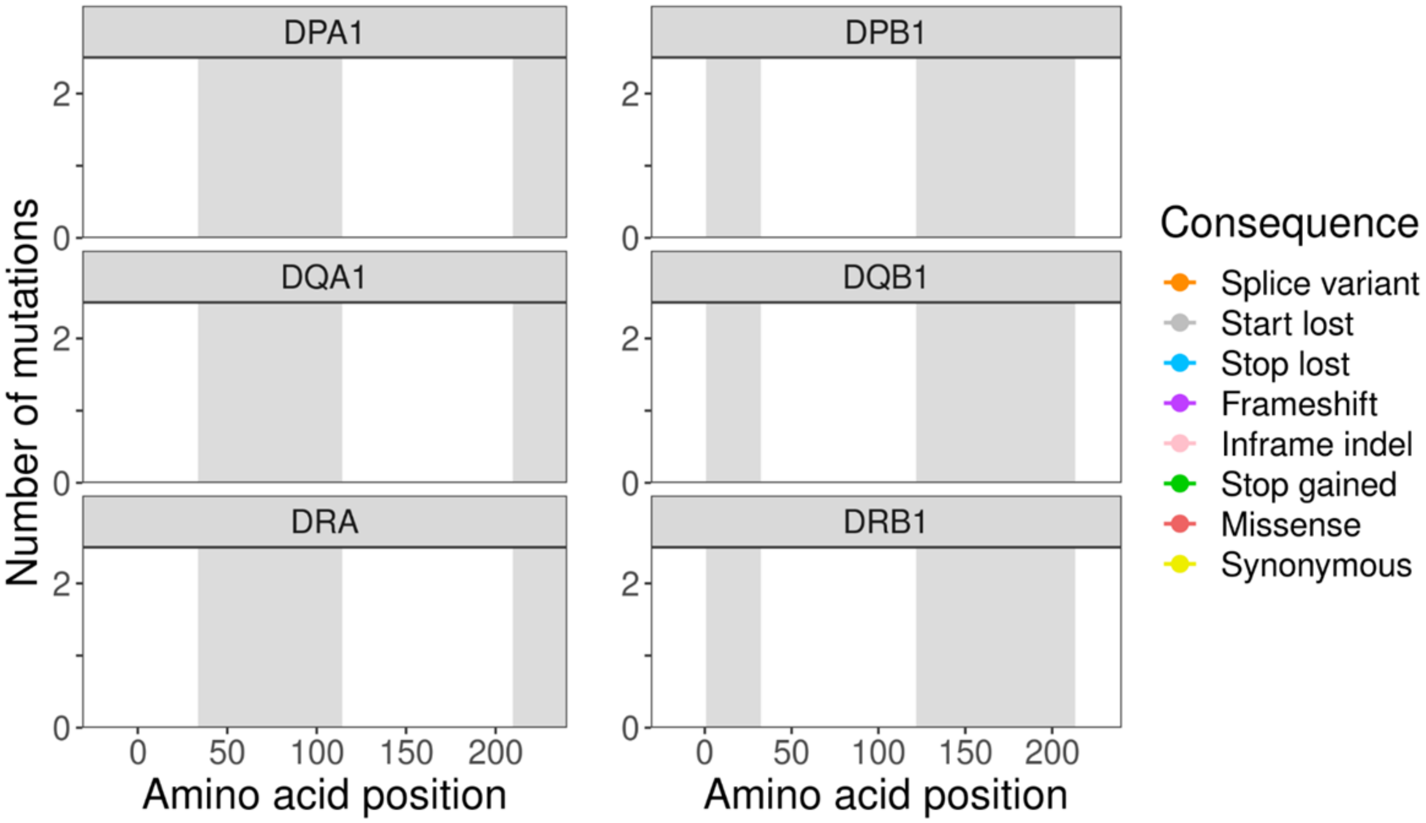
DPA1
DPB1
DQA1
DQB1
DRA
DRB1
Number of mutations
Amino acid position
Amino acid position
Consequence
Splice variant
Start lost
Stop lost
Frameshift
Inframe indel
Stop gained
Missense
Synonymous

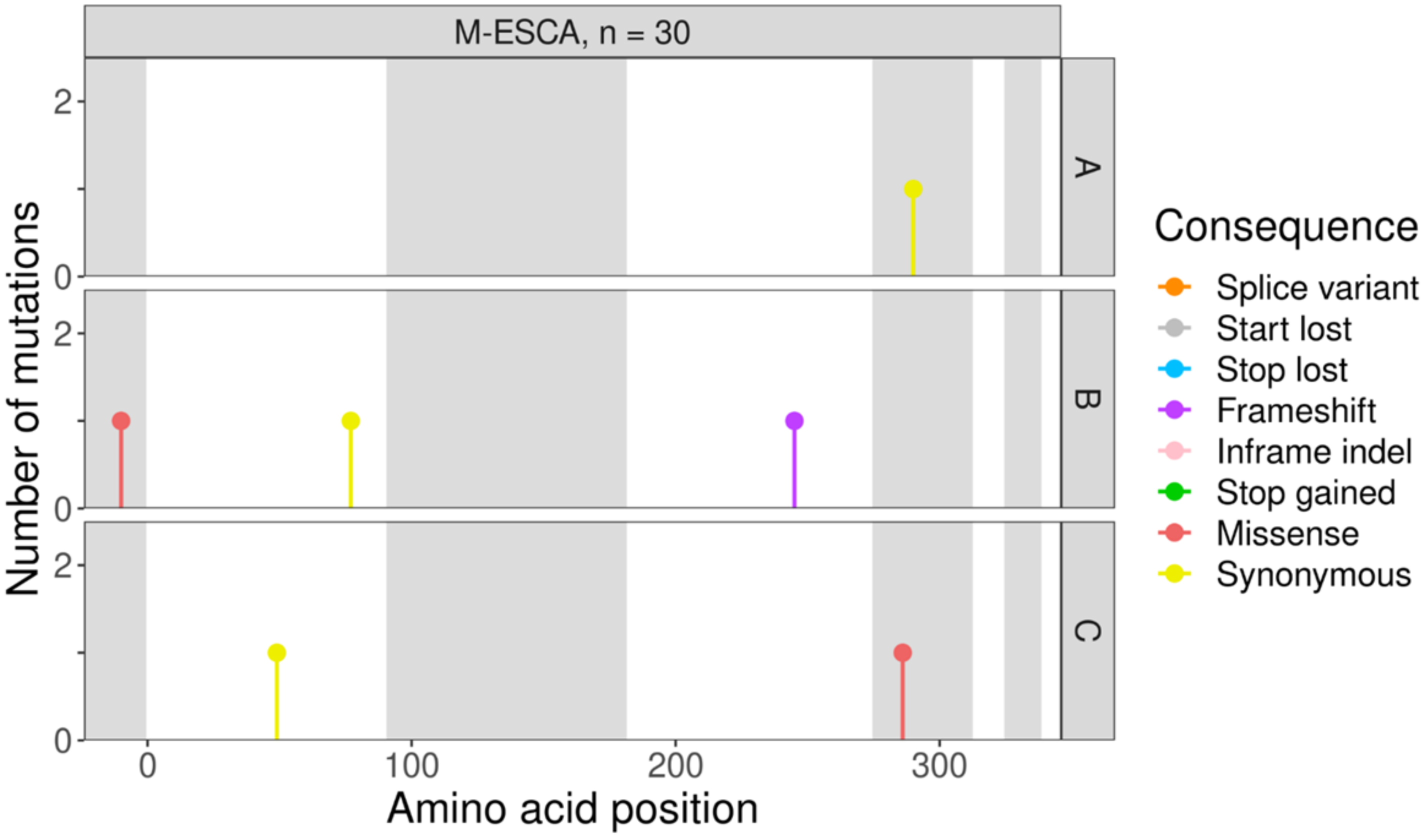
M-ESCA, n = 30
A
B
C
Number of mutations
Amino acid position
Consequence
Splice variant
Start lost
Stop lost
Frameshift
Inframe indel
Stop gained
Missense
Synonymous

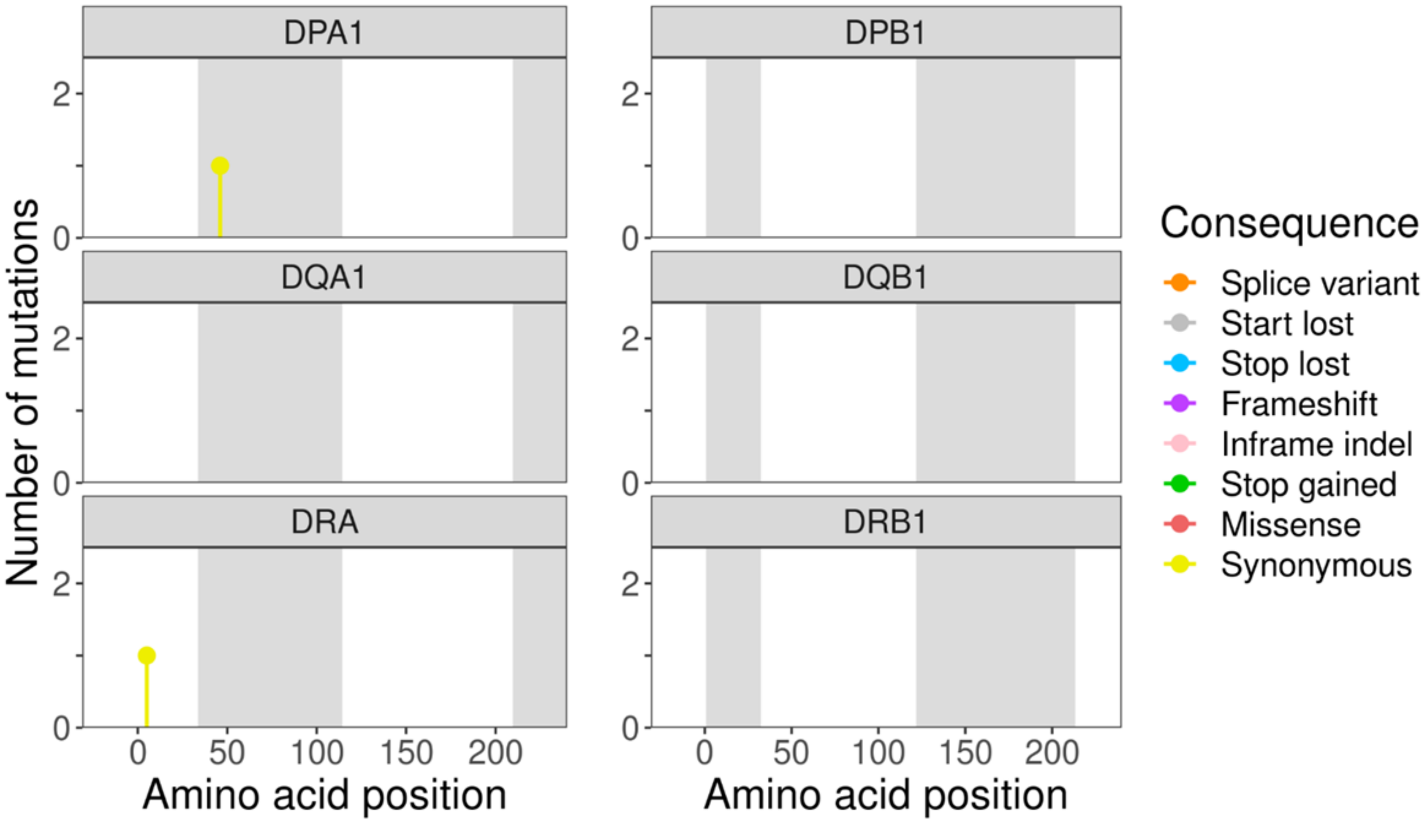
DPA1
DPB1
DQA1
DQB1
DRA
DRB1
Number of mutations
Amino acid position
Amino acid position
Consequence
Splice variant
Start lost
Stop lost
Frameshift
Inframe indel
Stop gained
Missense
Synonymous

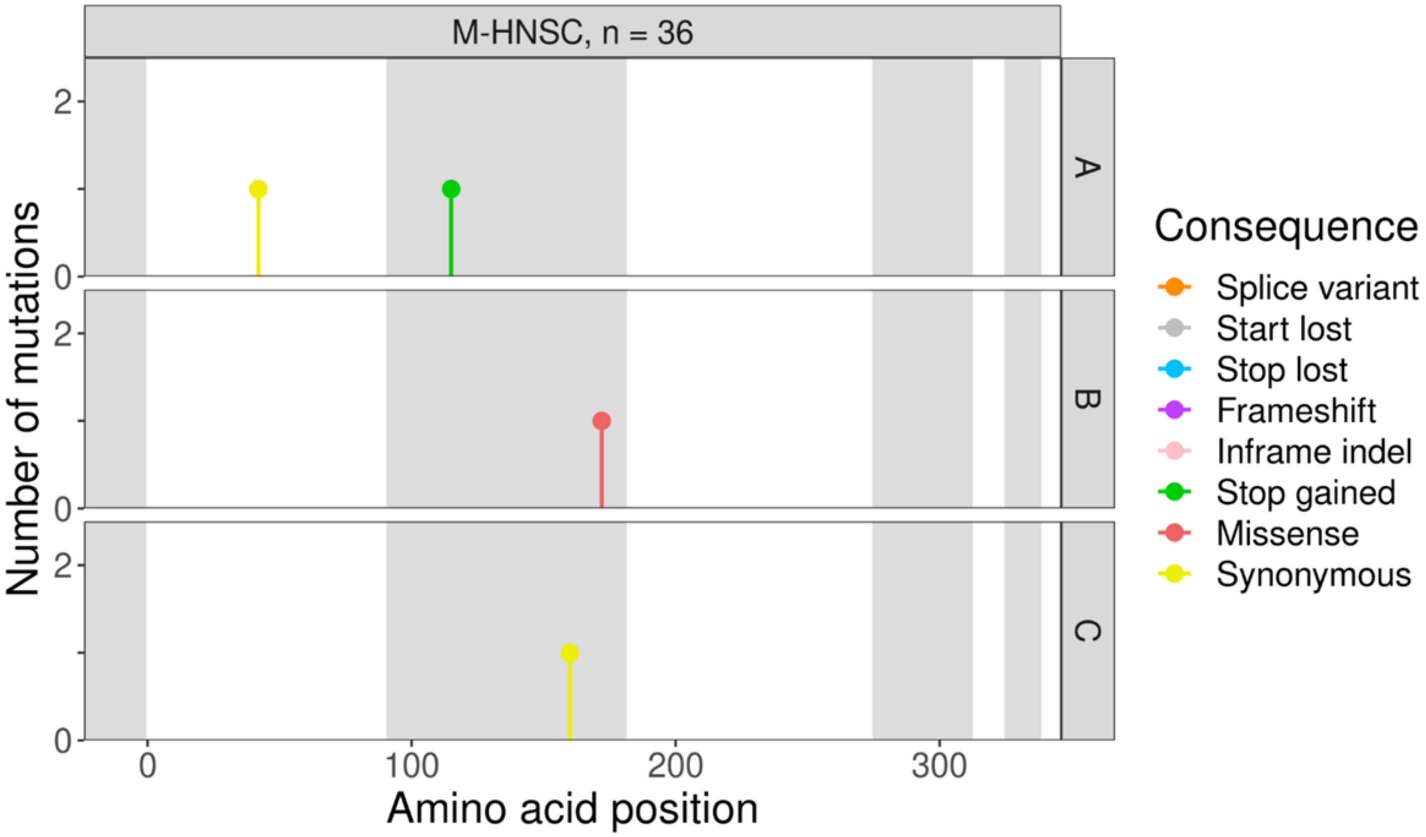

M-HNSC, n = 36
A
B
C
Number of mutations
Amino acid position
Consequence
Splice variant
Start lost
Stop lost
Frameshift
Inframe indel
Stop gained
Missense
Synonymous


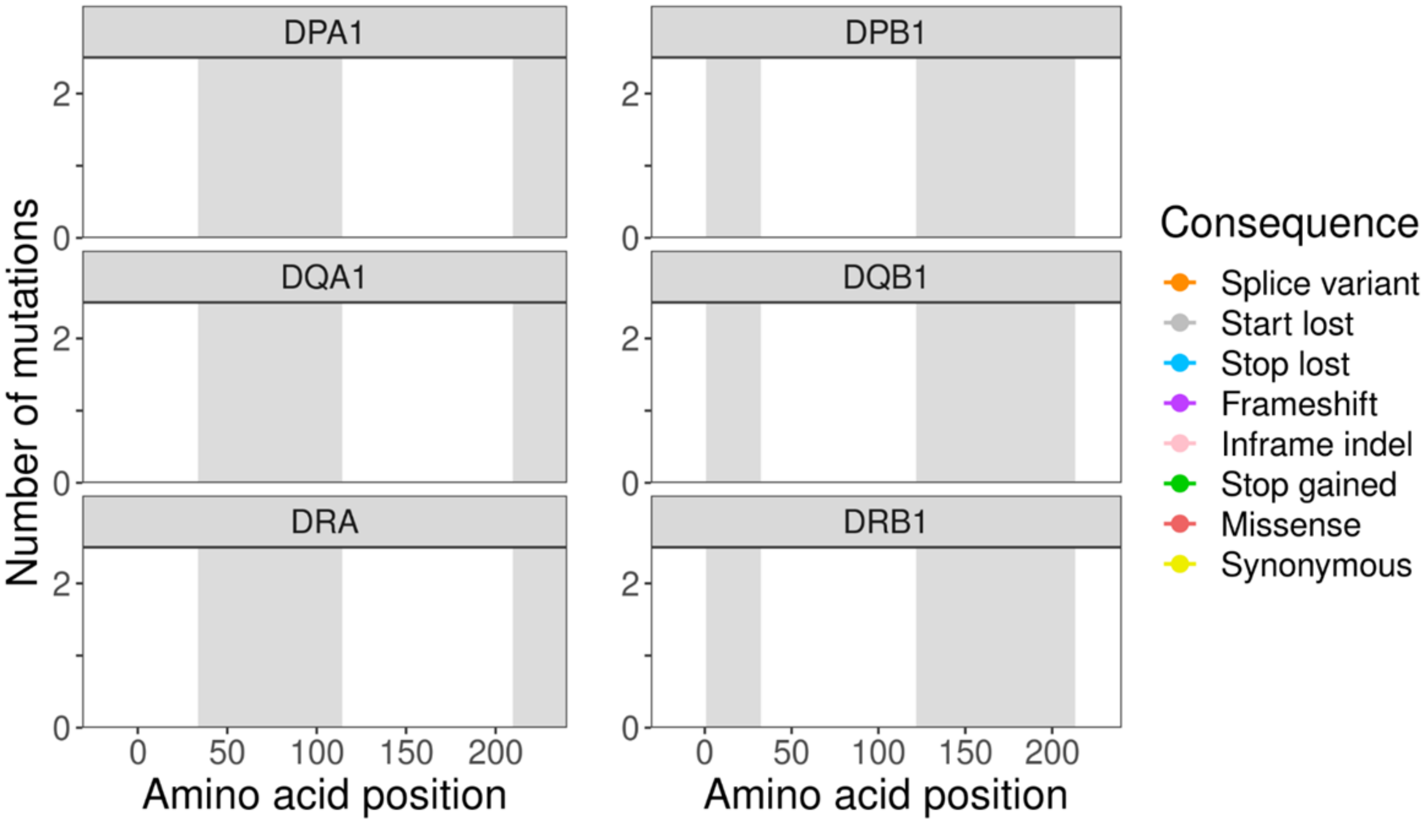

DPA1
DPB1
DQA1
DQB1
DRA
DRB1
Number of mutations
Amino acid position
Amino acid position
Consequence
Splice variant
Start lost
Stop lost
Frameshift
Inframe indel
Stop gained
Missense
Synonymous

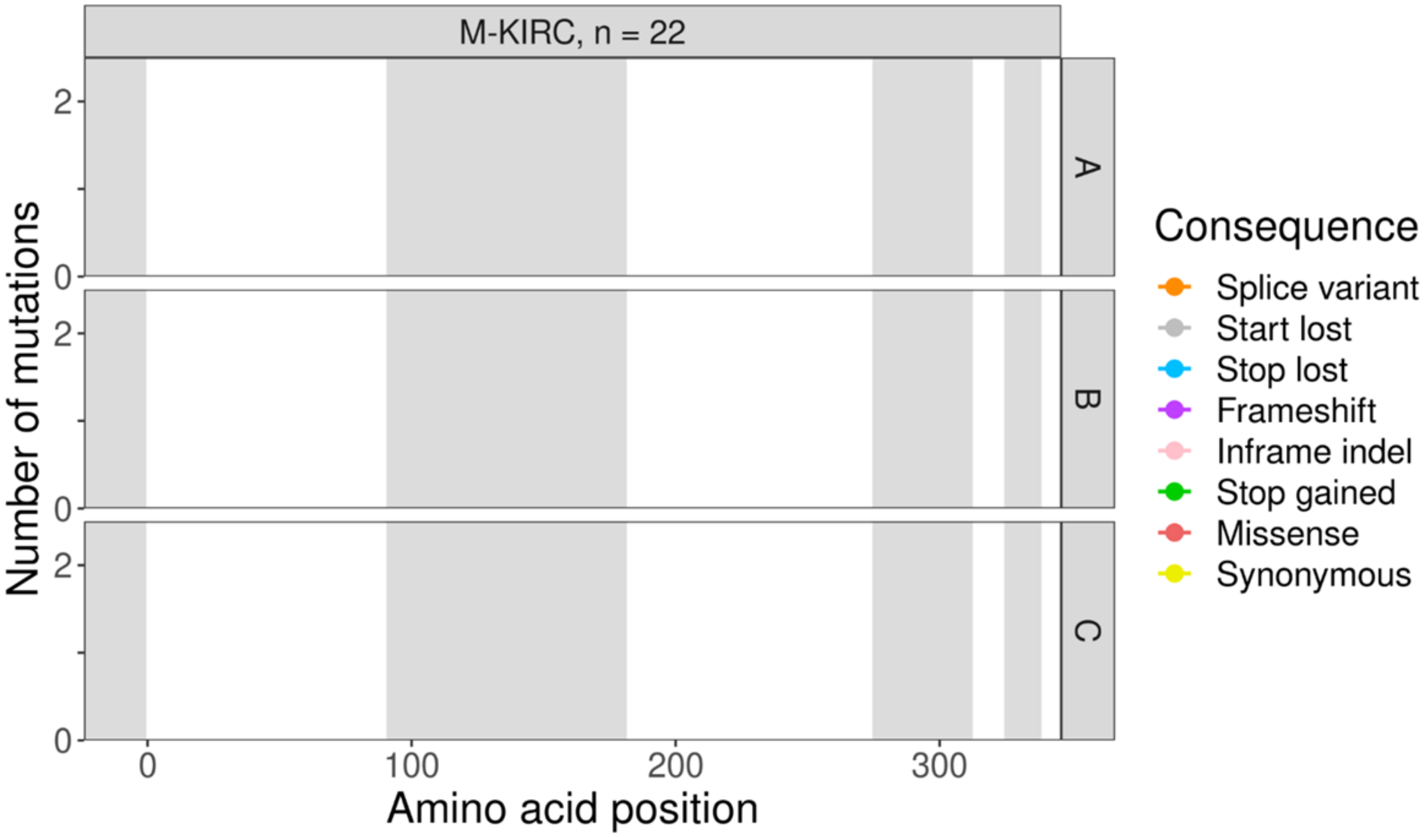
M-KIRC, n = 22
A
B
C
Number of mutations
Amino acid position
Consequence
Splice variant
Start lost
Stop lost
Frameshift
Inframe indel
Stop gained
Missense
Synonymous

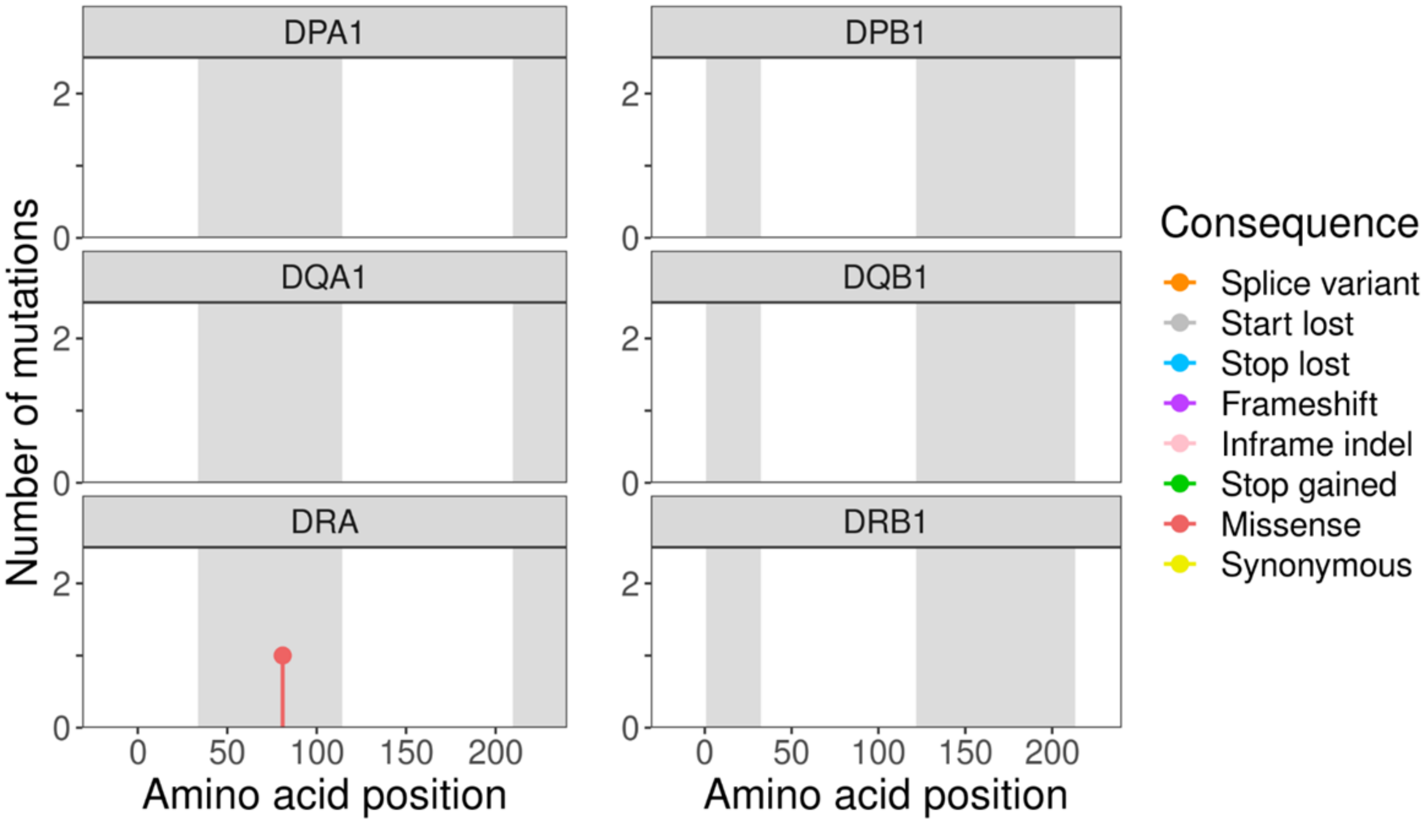
DPA1
DPB1
DQA1
DQB1
DRA
DRB1
Number of mutations
Amino acid position
Consequence
Splice variant
Start lost
Stop lost
Frameshift
Inframe indel
Stop gained
Missense
Synonymous

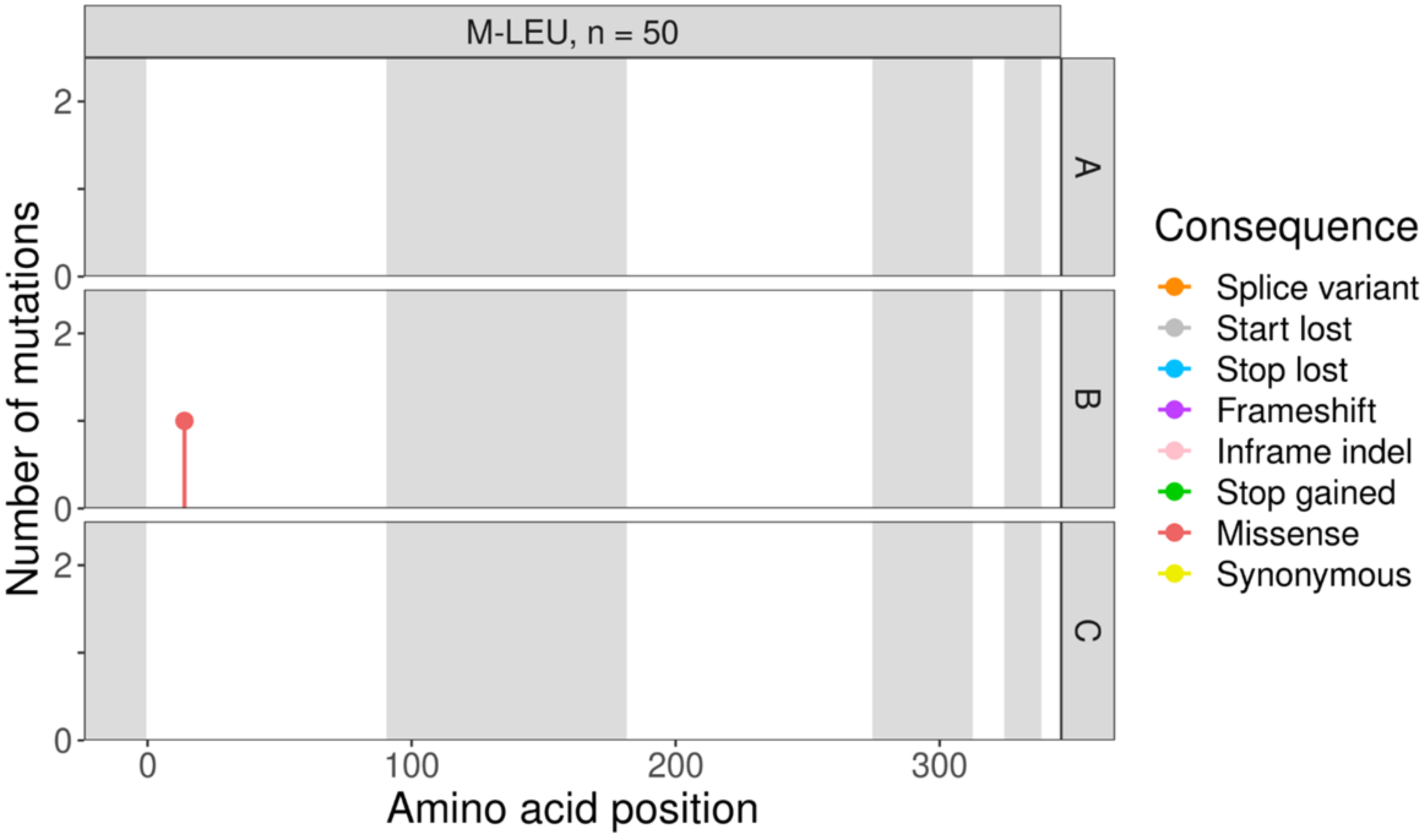
M-LEU, n = 50
A
B
C
Number of mutations
Amino acid position
Consequence
Splice variant
Start lost
Stop lost
Frameshift
Inframe indel
Stop gained
Missense
Synonymous

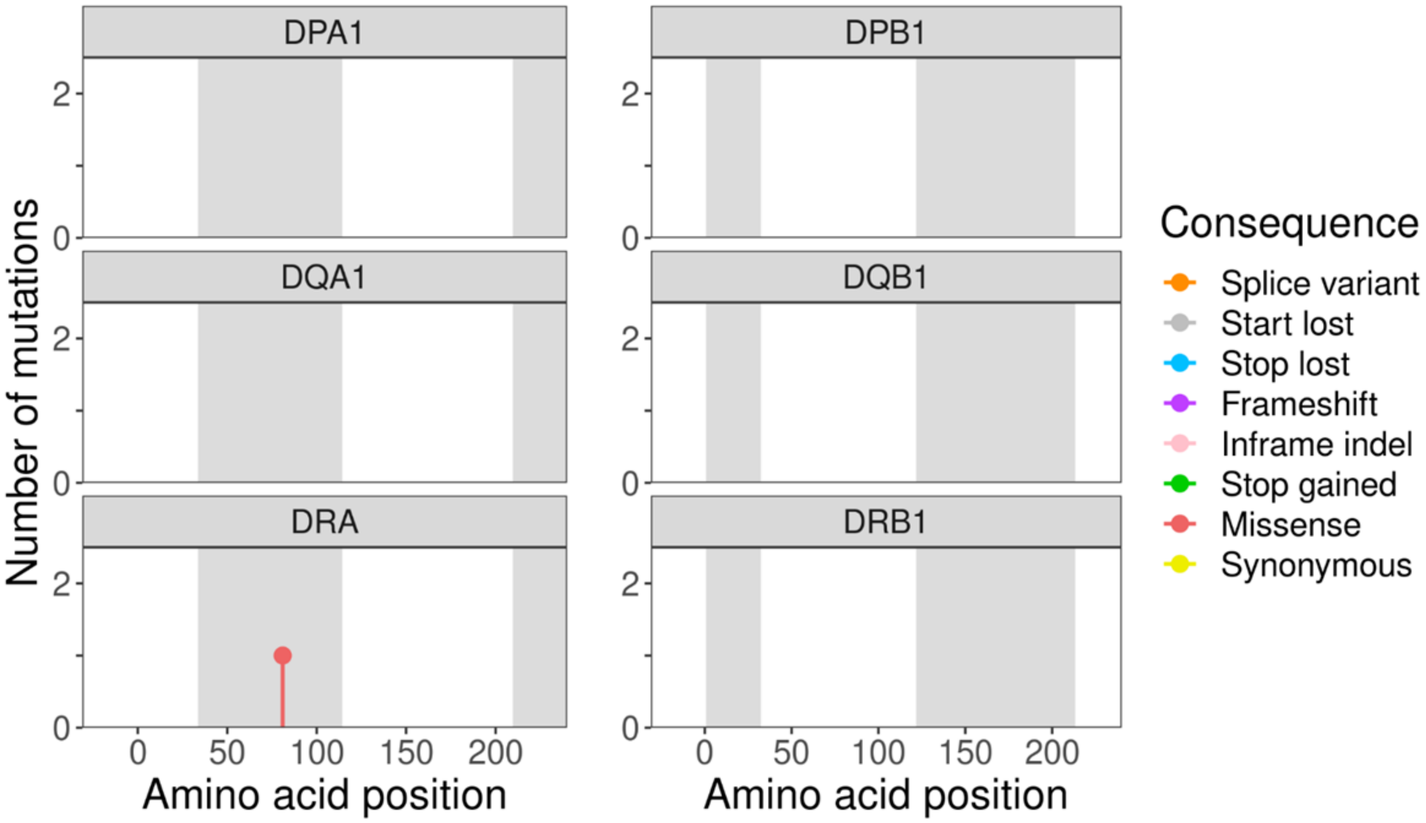
DPA1
DPB1
DQA1
DQB1
DRA
DRB1
Number of mutations
Amino acid position
Consequence
Splice variant
Start lost
Stop lost
Frameshift
Inframe indel
Stop gained
Missense
Synonymous

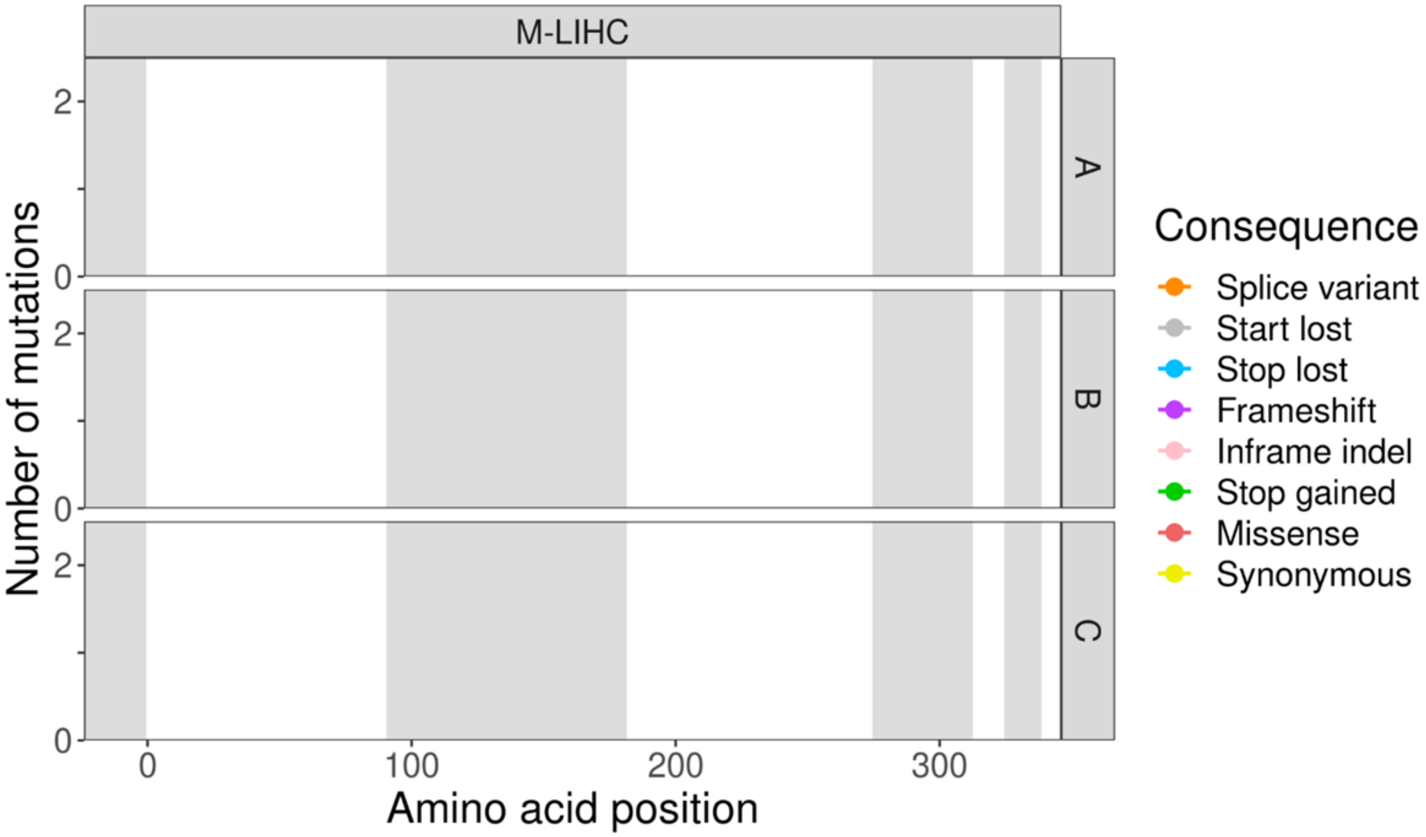
M-LIHC
A
B
C
Number of mutations
Amino acid position
Consequence
Splice variant
Start lost
Stop lost
Frameshift
Inframe indel
Stop gained
Missense
Synonymous

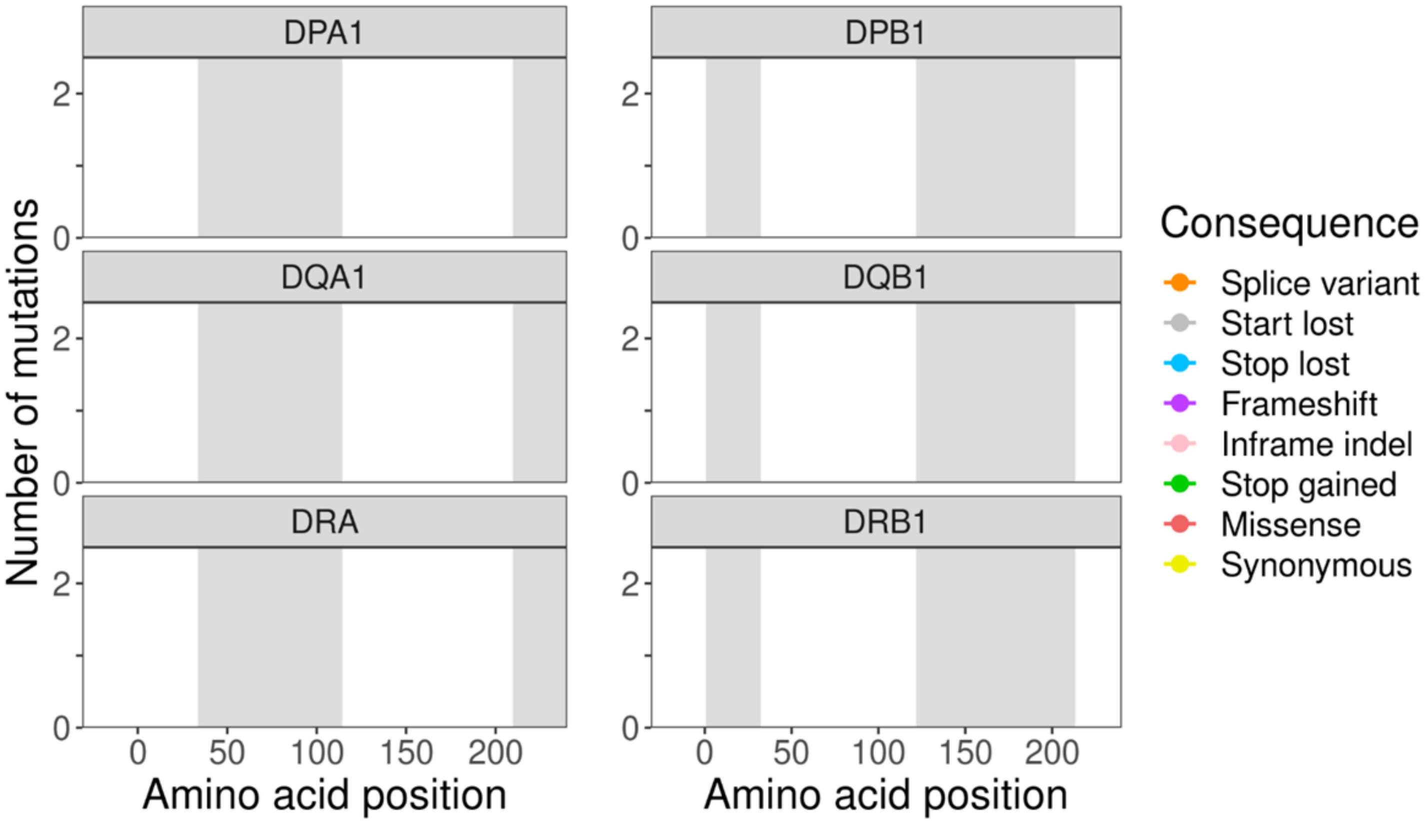
DPA1
DPB1
DQA1
DQB1
DRA
DRB1
Number of mutations
Amino acid position
Consequence
Splice variant
Start lost
Stop lost
Frameshift
Inframe indel
Stop gained
Missense
Synonymous

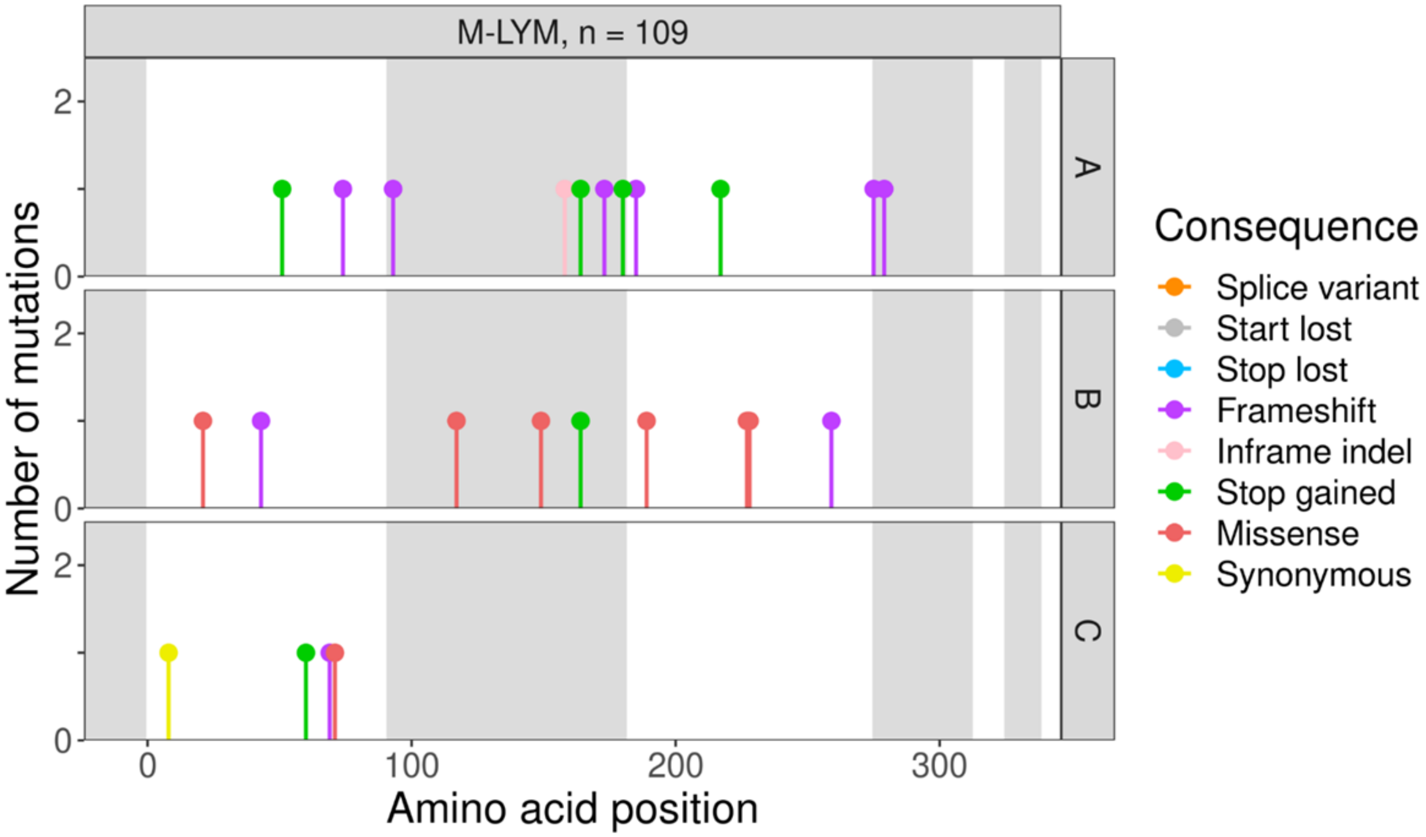
M-LYM, n = 109
A
B
C
Number of mutations
Amino acid position
0
100
200
300
Consequence
Splice variant
Start lost
Stop lost
Frameshift
Inframe indel
Stop gained
Missense
Synonymous

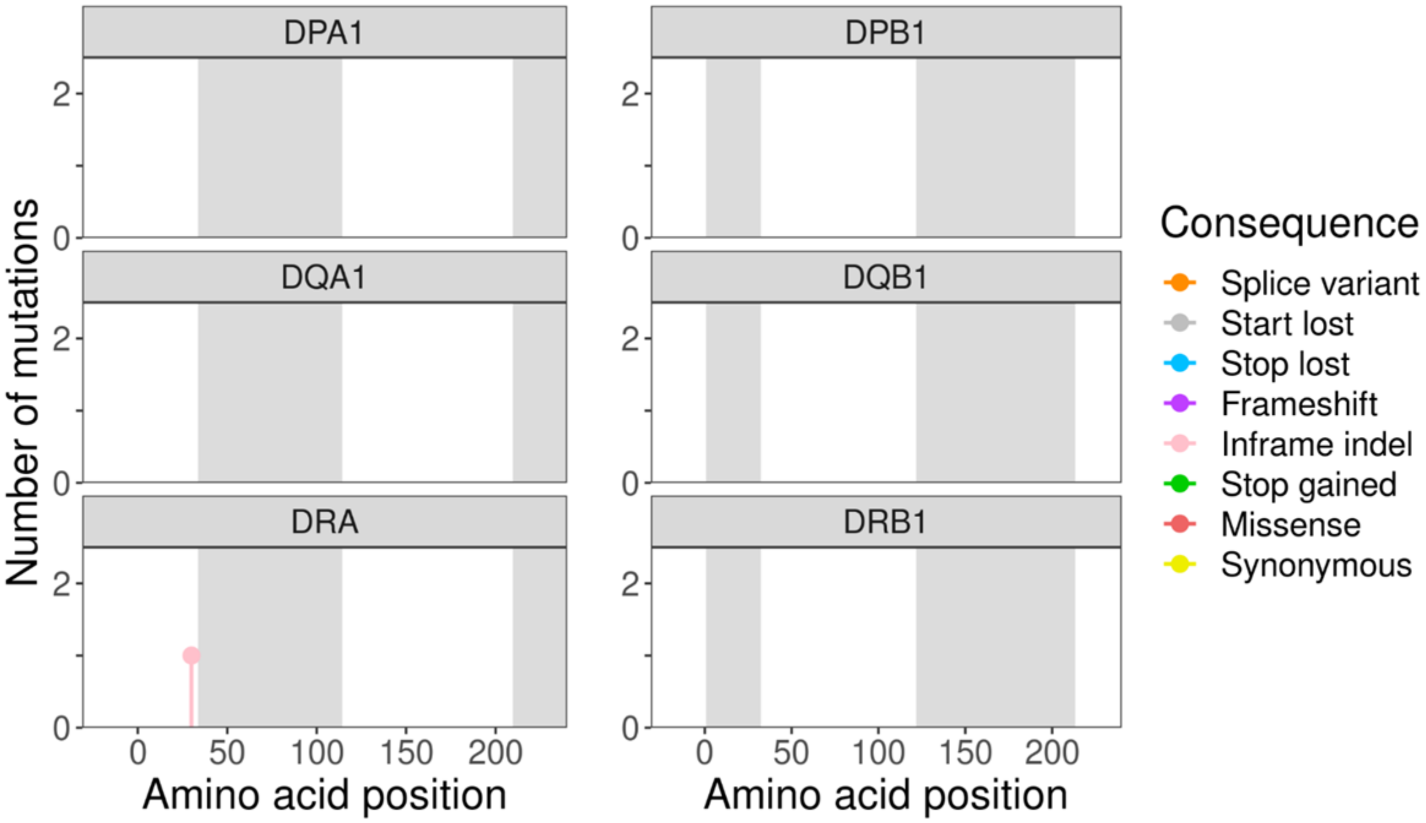
DPA1
DPB1
DQA1
DQB1
DRA
DRB1
Number of mutations
Amino acid position
0
50
100
150
200
Consequence
Splice variant
Start lost
Stop lost
Frameshift
Inframe indel
Stop gained
Missense
Synonymous

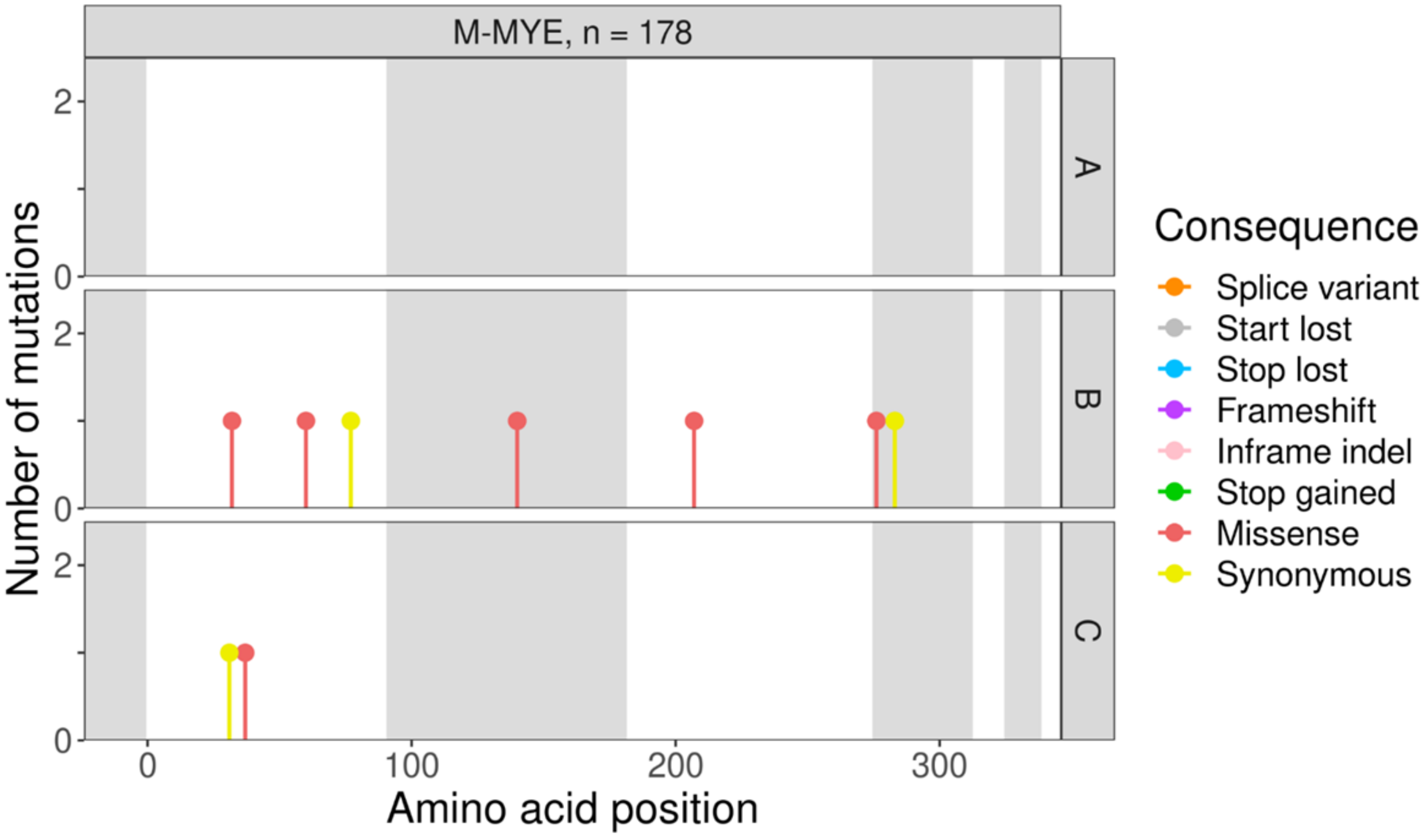

M-MYE, n = 178
A
B
C
Number of mutations
Amino acid position
Consequence
Splice variant
Start lost
Stop lost
Frameshift
Inframe indel
Stop gained
Missense
Synonymous


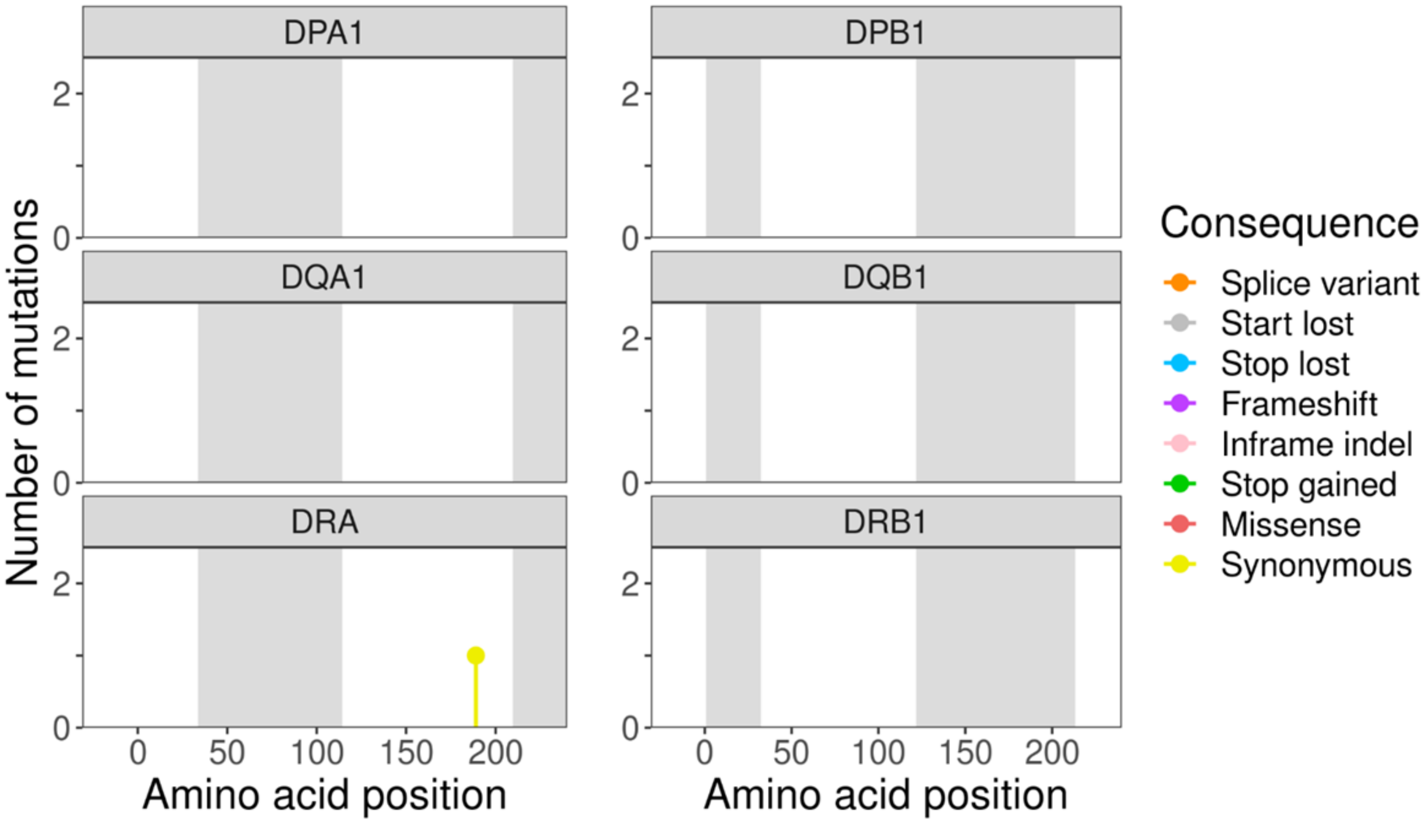

DPA1
DPB1
DQA1
DQB1
DRA
DRB1
Number of mutations
Amino acid position
Consequence
Splice variant
Start lost
Stop lost
Frameshift
Inframe indel
Stop gained
Missense
Synonymous

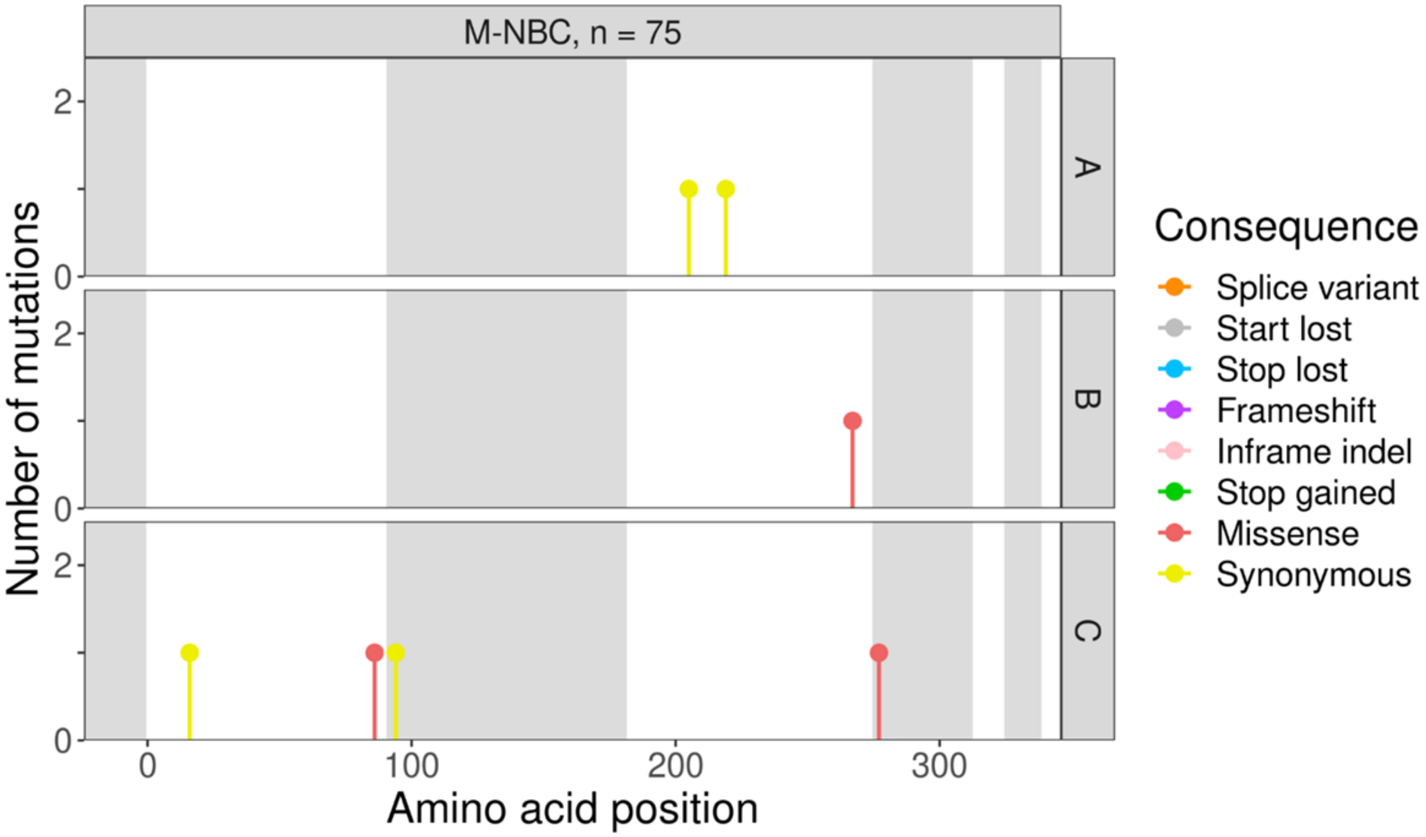
M-NBC, n = 75
A
B
C
Number of mutations
Amino acid position
Consequence
Splice variant
Start lost
Stop lost
Frameshift
Inframe indel
Stop gained
Missense
Synonymous

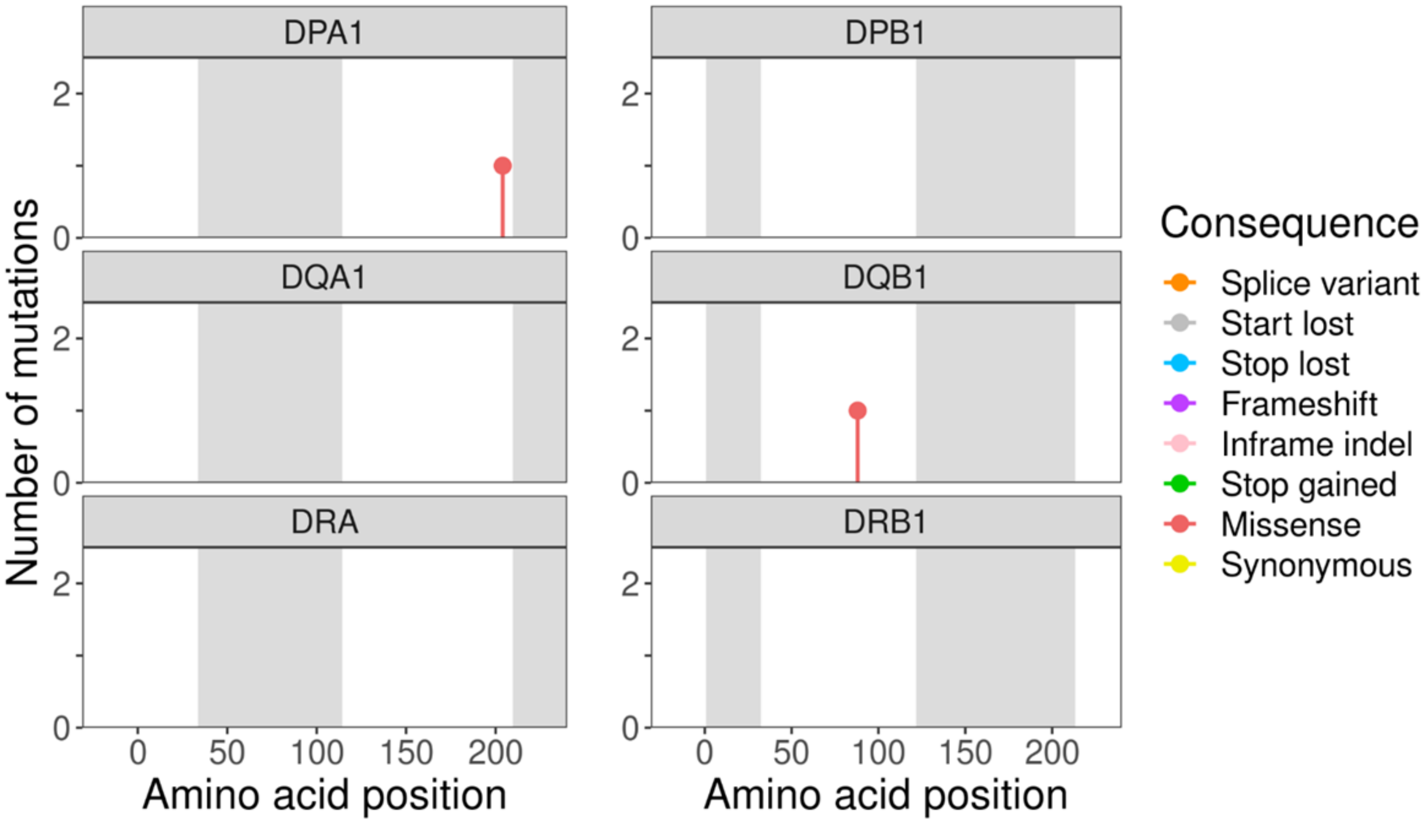
DPA1
DPB1
DQA1
DQB1
DRA
DRB1
Number of mutations
Amino acid position
Amino acid position
Consequence
Splice variant
Start lost
Stop lost
Frameshift
Inframe indel
Stop gained
Missense
Synonymous

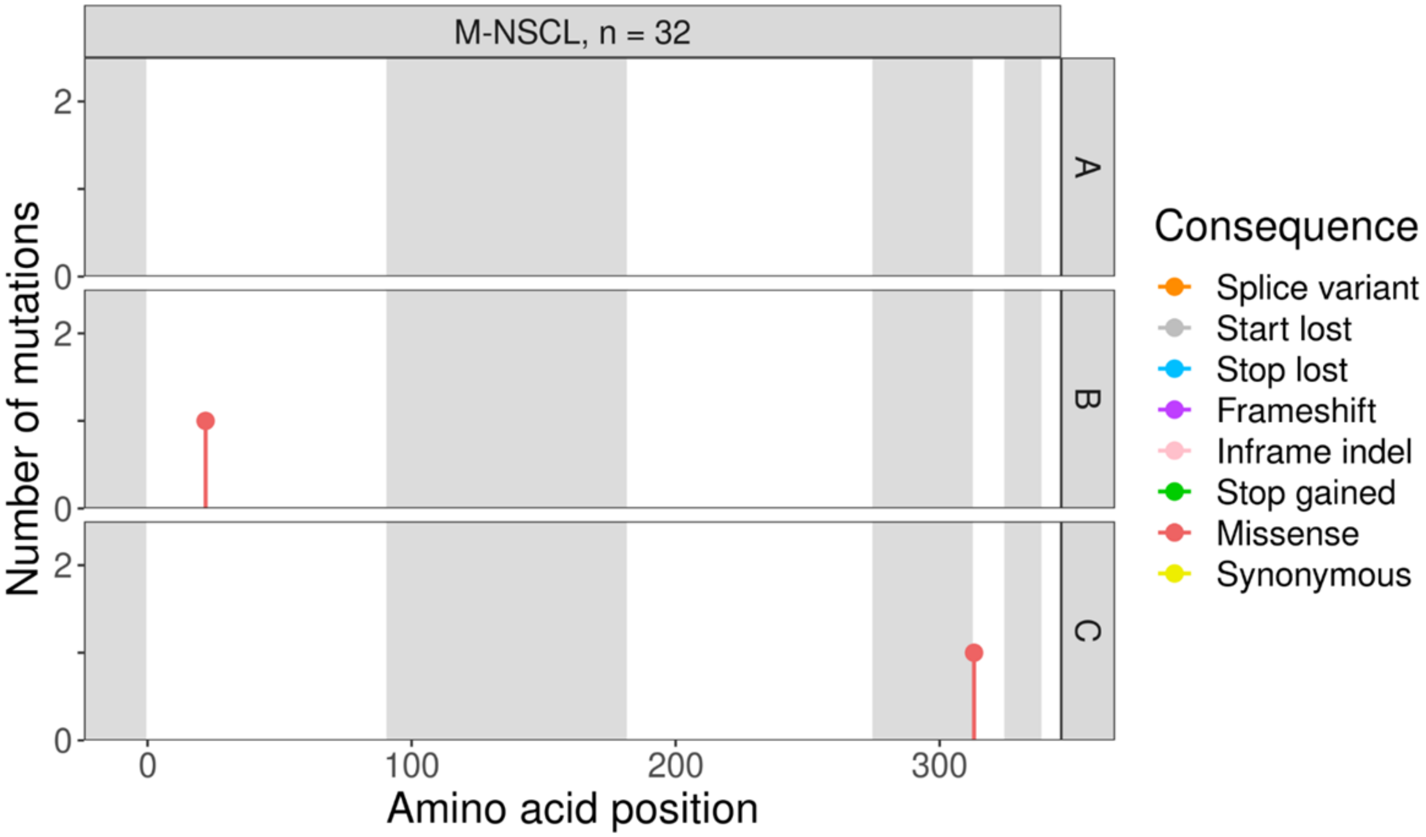
M-NSCL, n = 32
A
B
C
Number of mutations
Amino acid position
Consequence
Splice variant
Start lost
Stop lost
Frameshift
Inframe indel
Stop gained
Missense
Synonymous

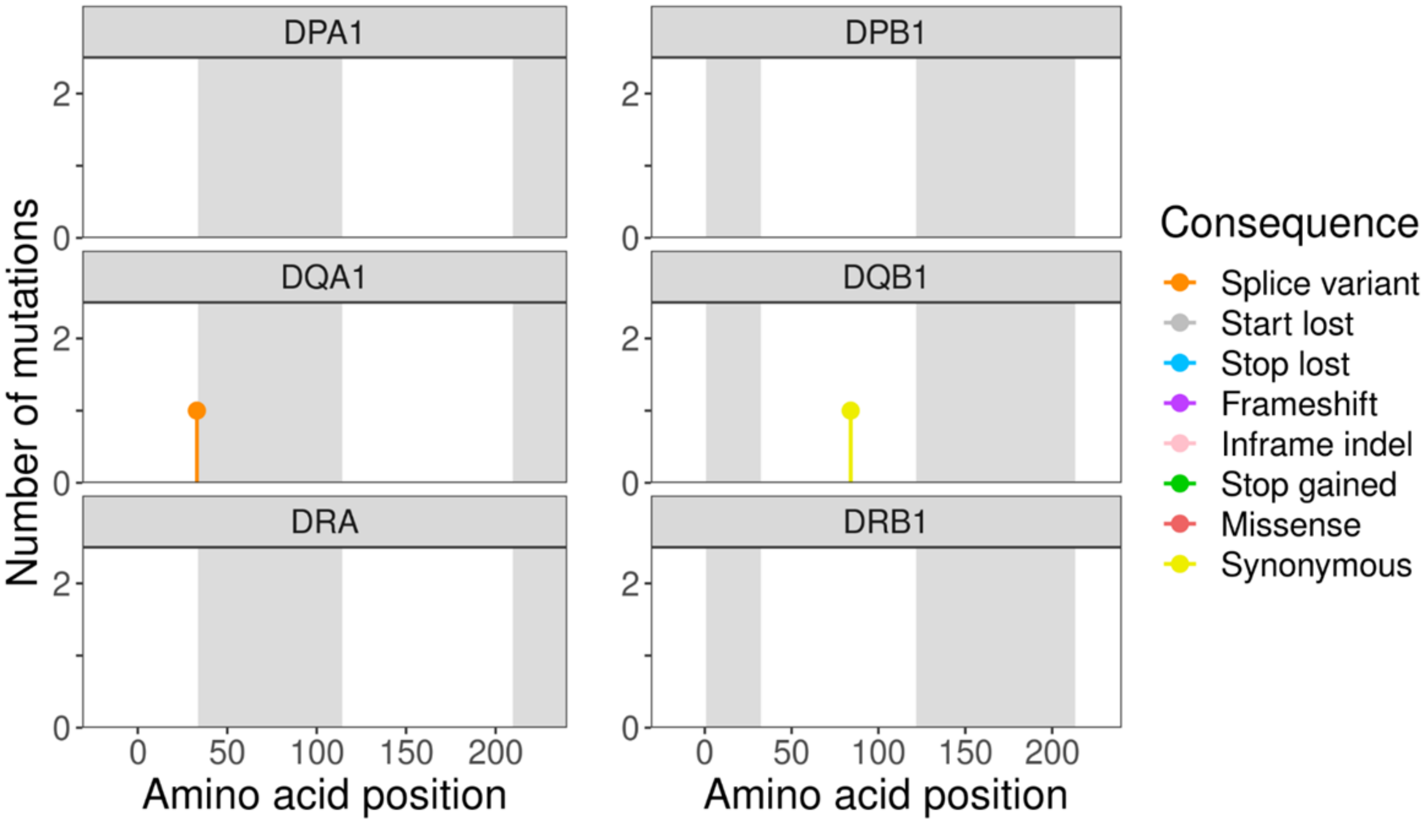
DPA1
DPB1
DQA1
DQB1
DRA
DRB1
Number of mutations
Amino acid position
Amino acid position
Consequence
Splice variant
Start lost
Stop lost
Frameshift
Inframe indel
Stop gained
Missense
Synonymous

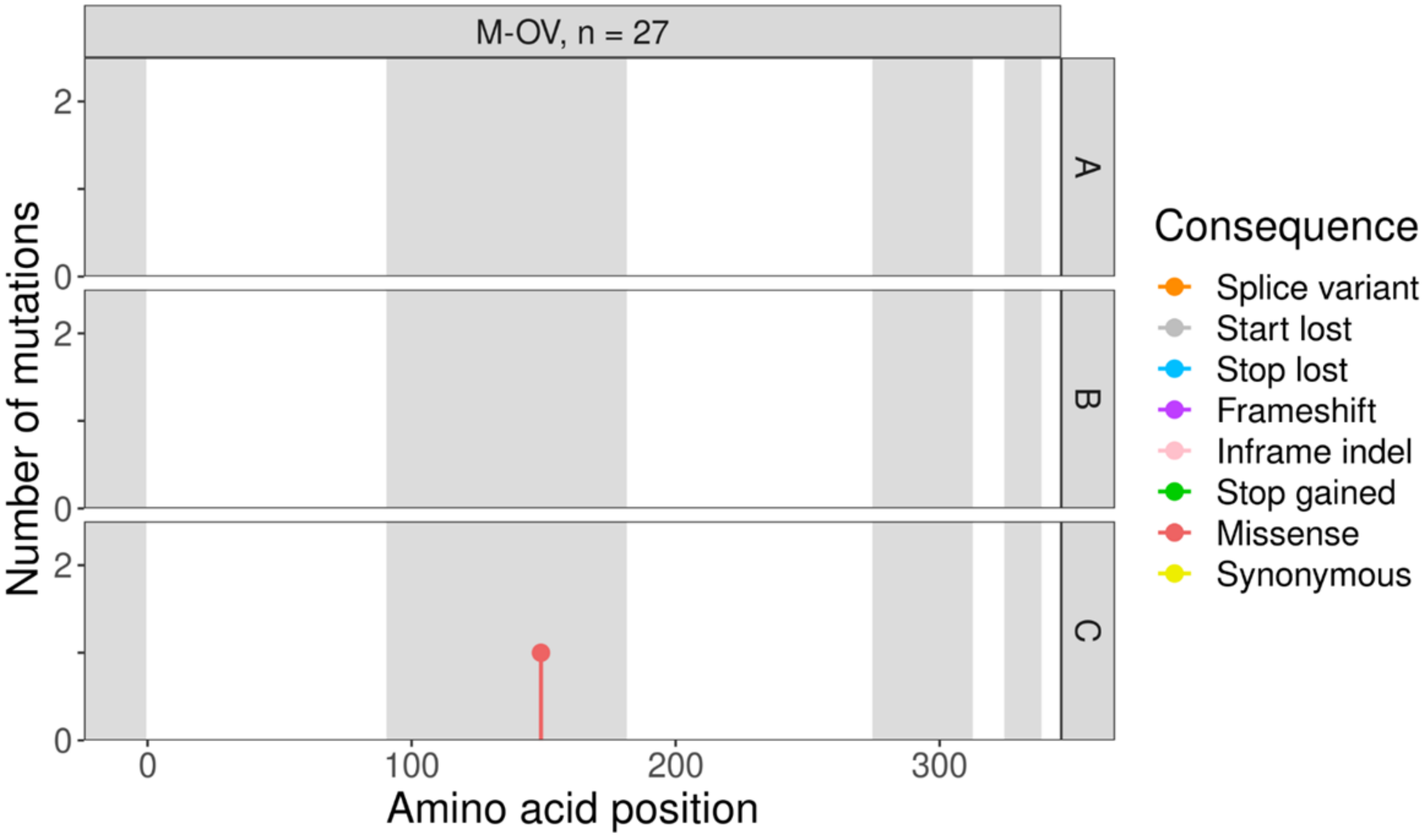
M-OV, n = 27
A
B
C
Number of mutations
Amino acid position
0
100
200
300
Consequence
Splice variant
Start lost
Stop lost
Frameshift
Inframe indel
Stop gained
Missense
Synonymous

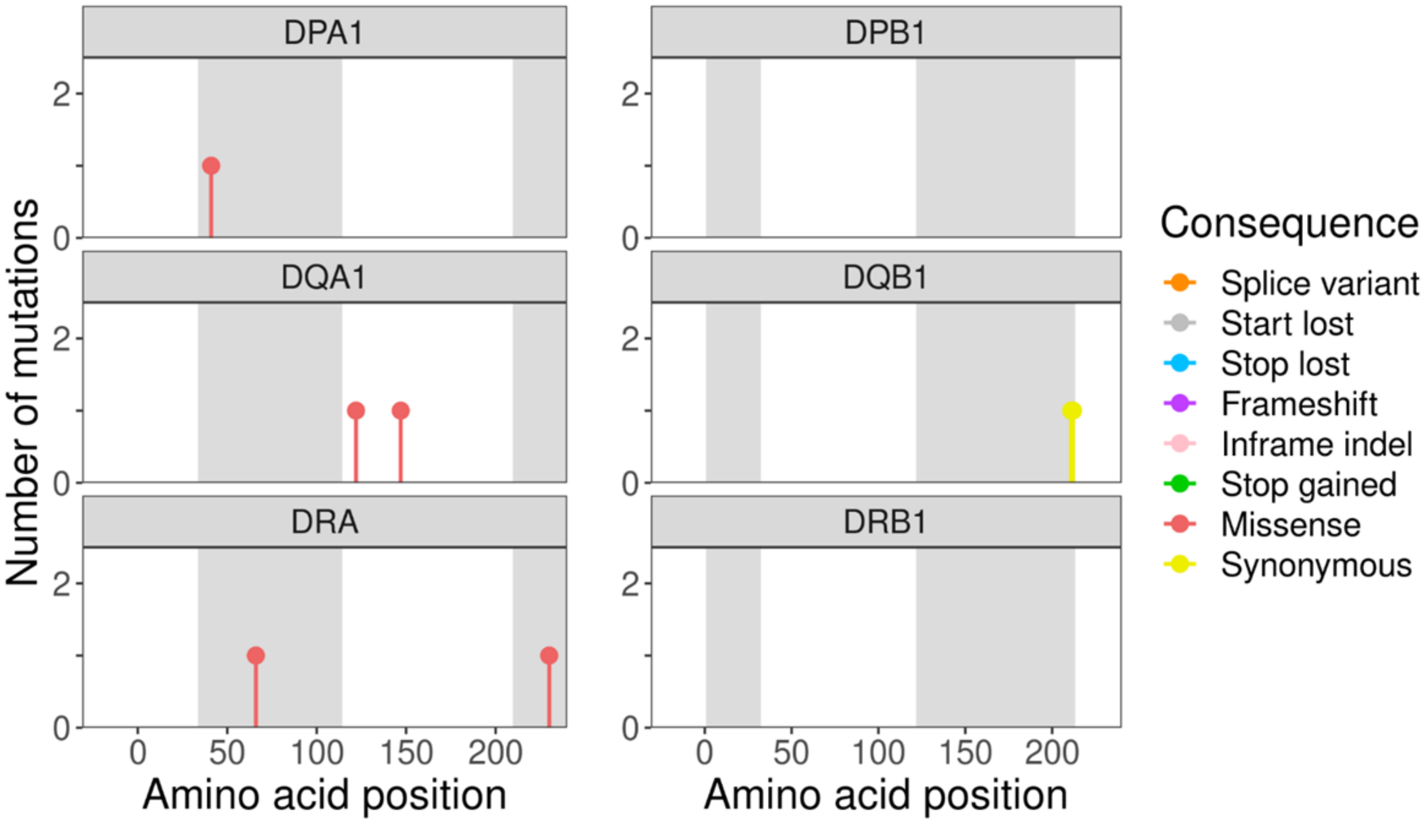
DPA1
DPB1
DQA1
DQB1
DRA
DRB1
Number of mutations
Amino acid position
0
50
100
150
200
Consequence
Splice variant
Start lost
Stop lost
Frameshift
Inframe indel
Stop gained
Missense
Synonymous

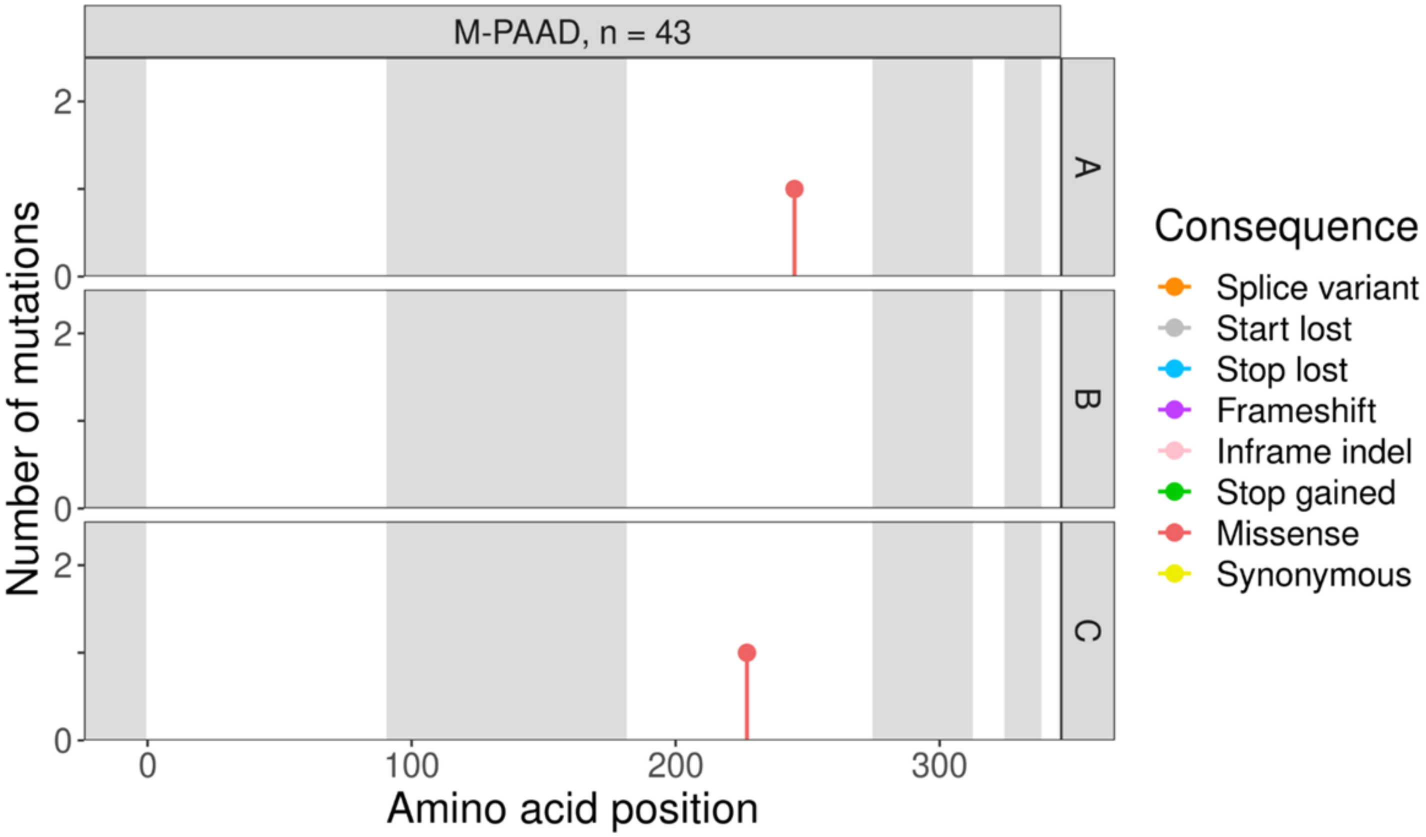
M-PAAD, n = 43
A
B
C
Number of mutations
Amino acid position
0
100
200
300
Consequence
Splice variant
Start lost
Stop lost
Frameshift
Inframe indel
Stop gained
Missense
Synonymous

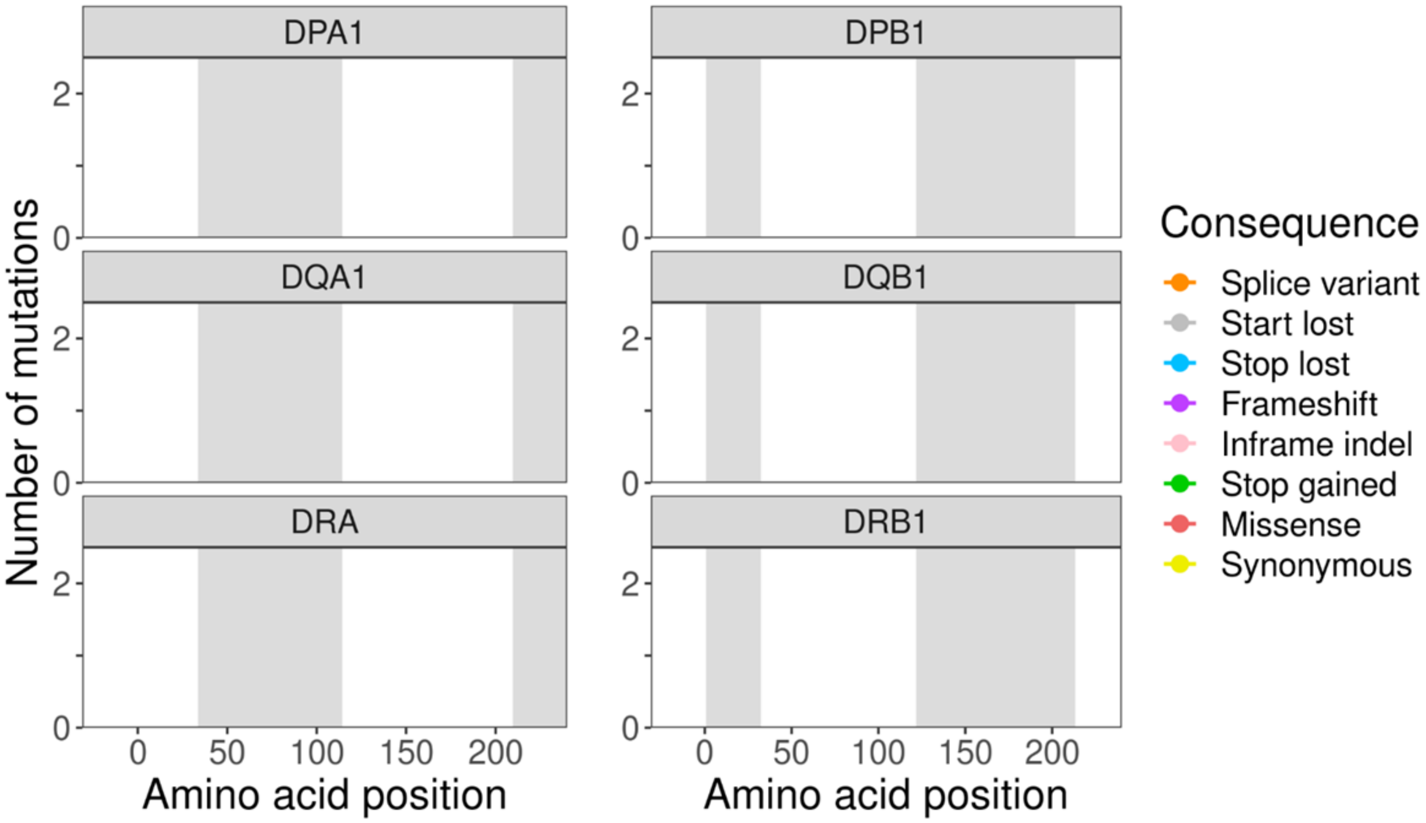
DPA1
DPB1
DQA1
DQB1
DRA
DRB1
Number of mutations
Amino acid position
0
50
100
150
200
Consequence
Splice variant
Start lost
Stop lost
Frameshift
Inframe indel
Stop gained
Missense
Synonymous

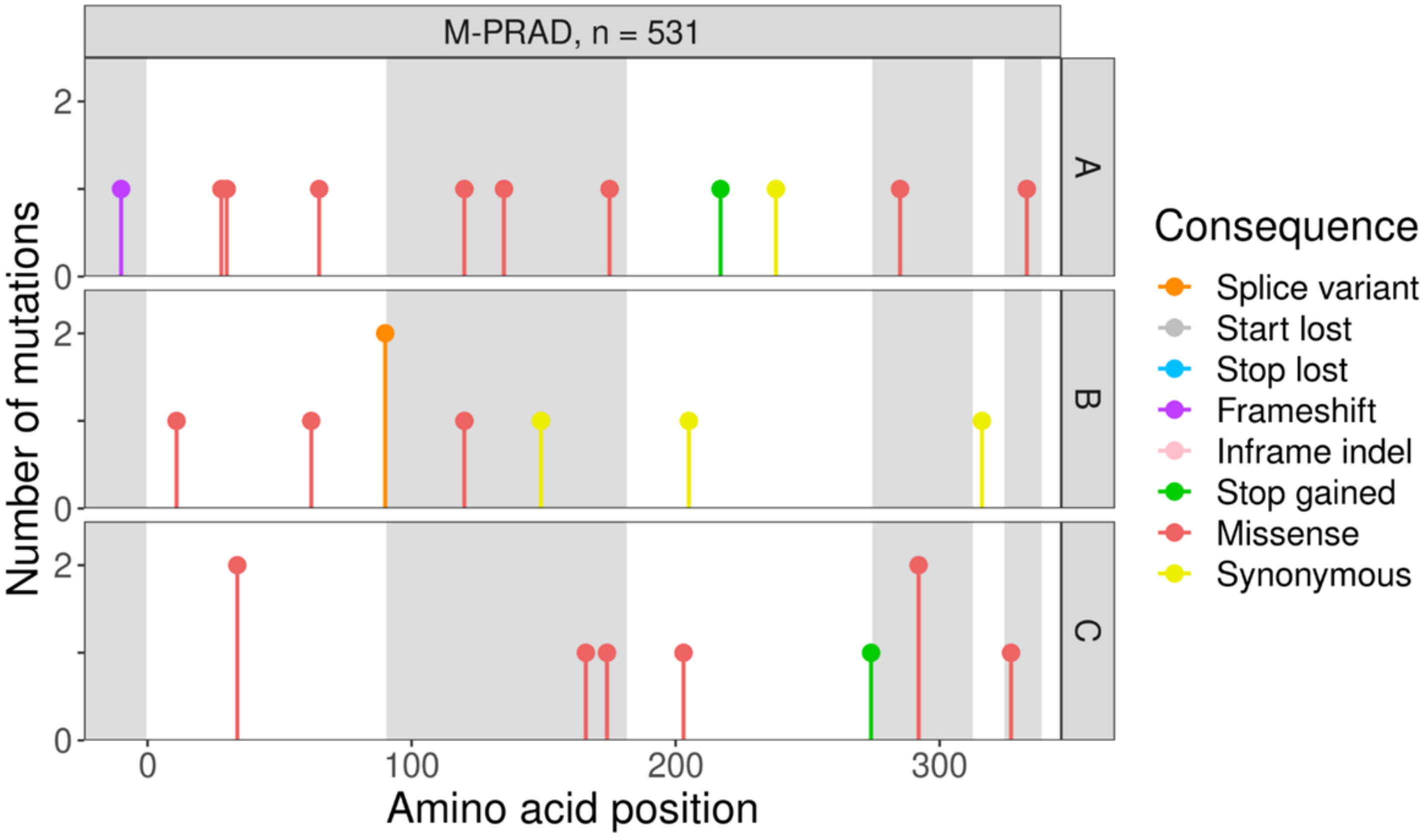
M-PRAD, n = 531
A
B
C
Number of mutations
Amino acid position
Consequence
Splice variant
Start lost
Stop lost
Frameshift
Inframe indel
Stop gained
Missense
Synonymous

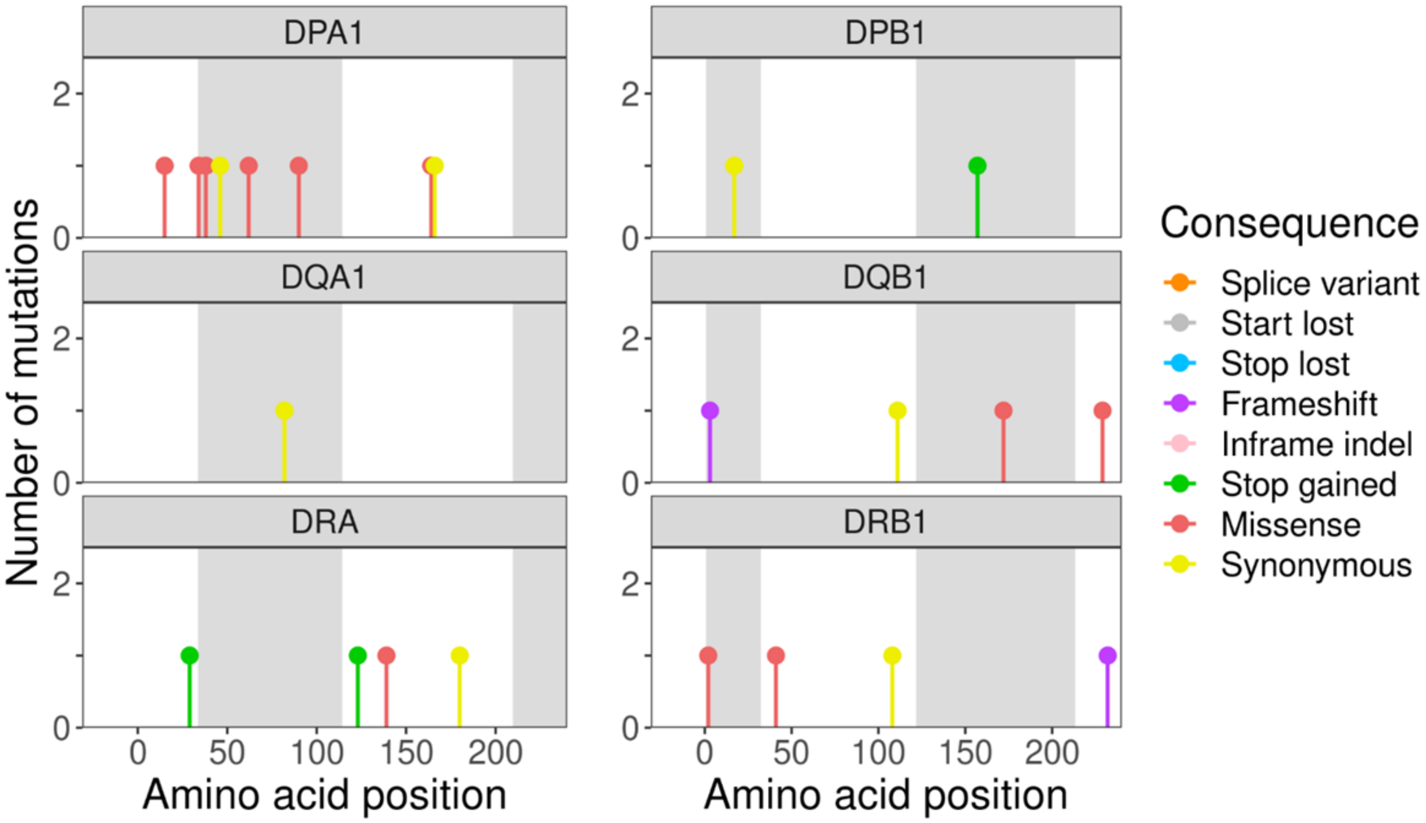
DPA1
DPB1
DQA1
DQB1
DRA
DRB1
Number of mutations
Amino acid position
Amino acid position
Consequence
Splice variant
Start lost
Stop lost
Frameshift
Inframe indel
Stop gained
Missense
Synonymous

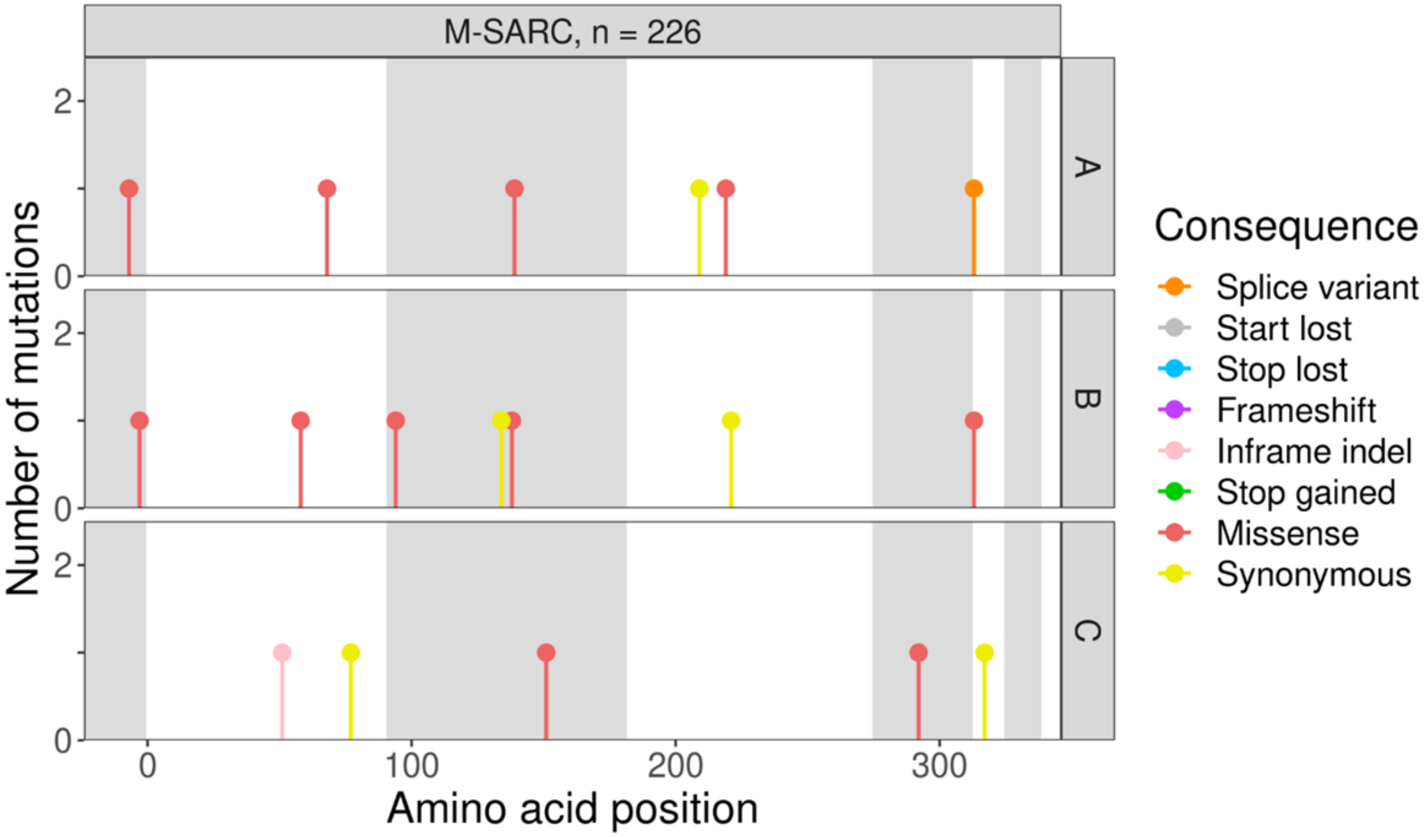

M-SARC, n = 226
A
B
C
Number of mutations
Amino acid position
Consequence
Splice variant
Start lost
Stop lost
Frameshift
Inframe indel
Stop gained
Missense
Synonymous


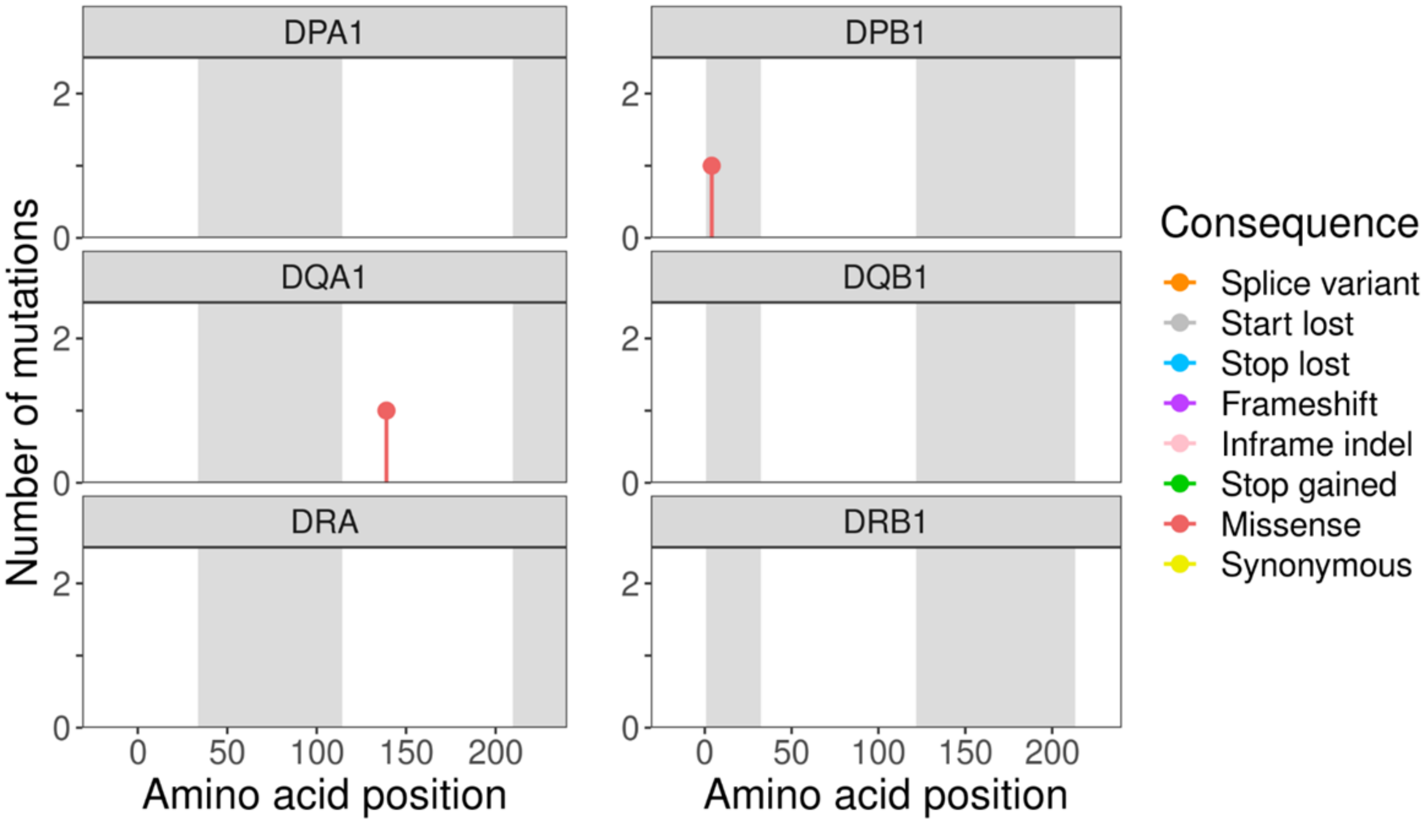

DPA1
DPB1
DQA1
DQB1
DRA
DRB1
Number of mutations
Amino acid position
Amino acid position
Consequence
Splice variant
Start lost
Stop lost
Frameshift
Inframe indel
Stop gained
Missense
Synonymous

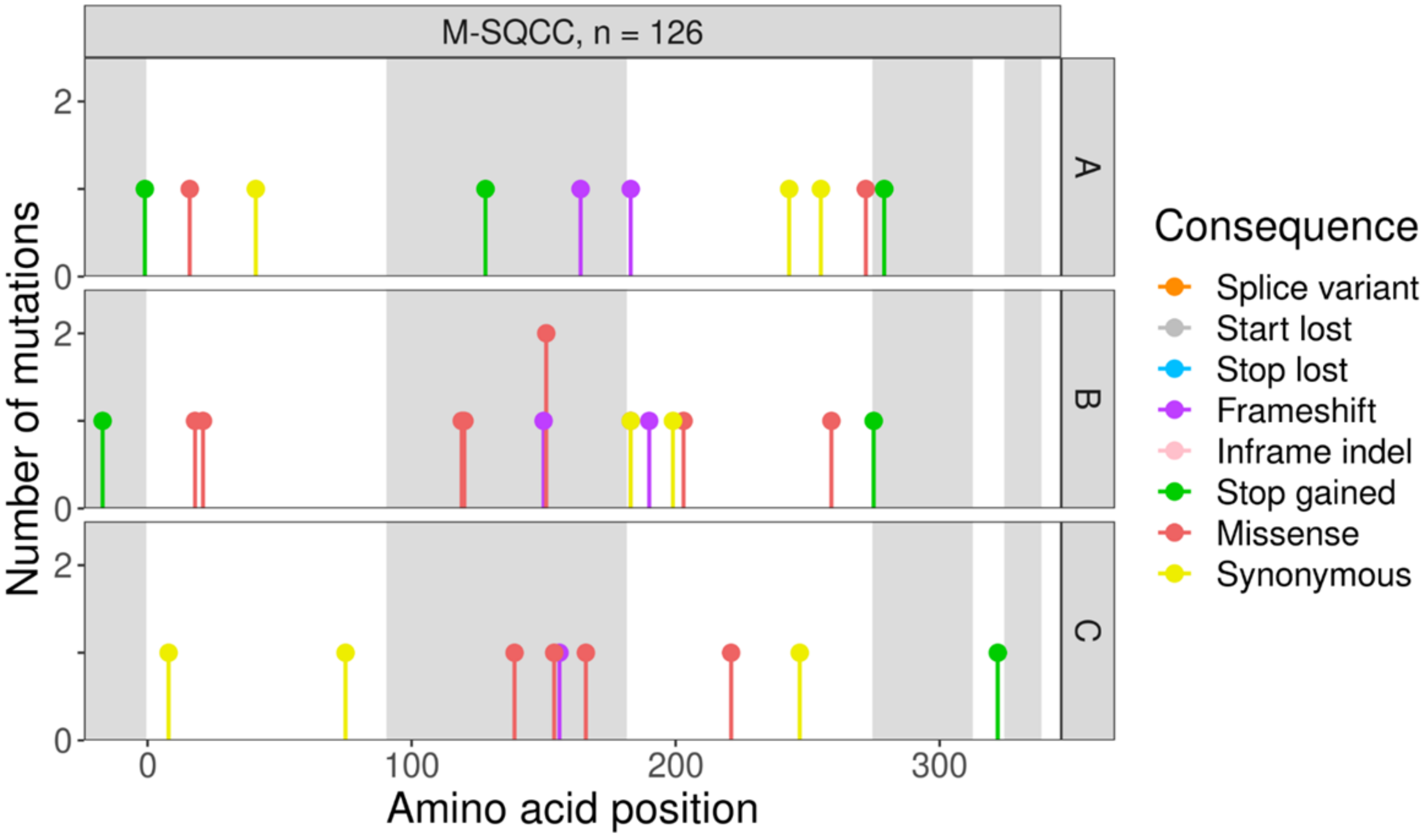
M-SQCC, n = 126
A
B
C
Number of mutations
Amino acid position
0
100
200
300
Consequence
Splice variant
Start lost
Stop lost
Frameshift
Inframe indel
Stop gained
Missense
Synonymous

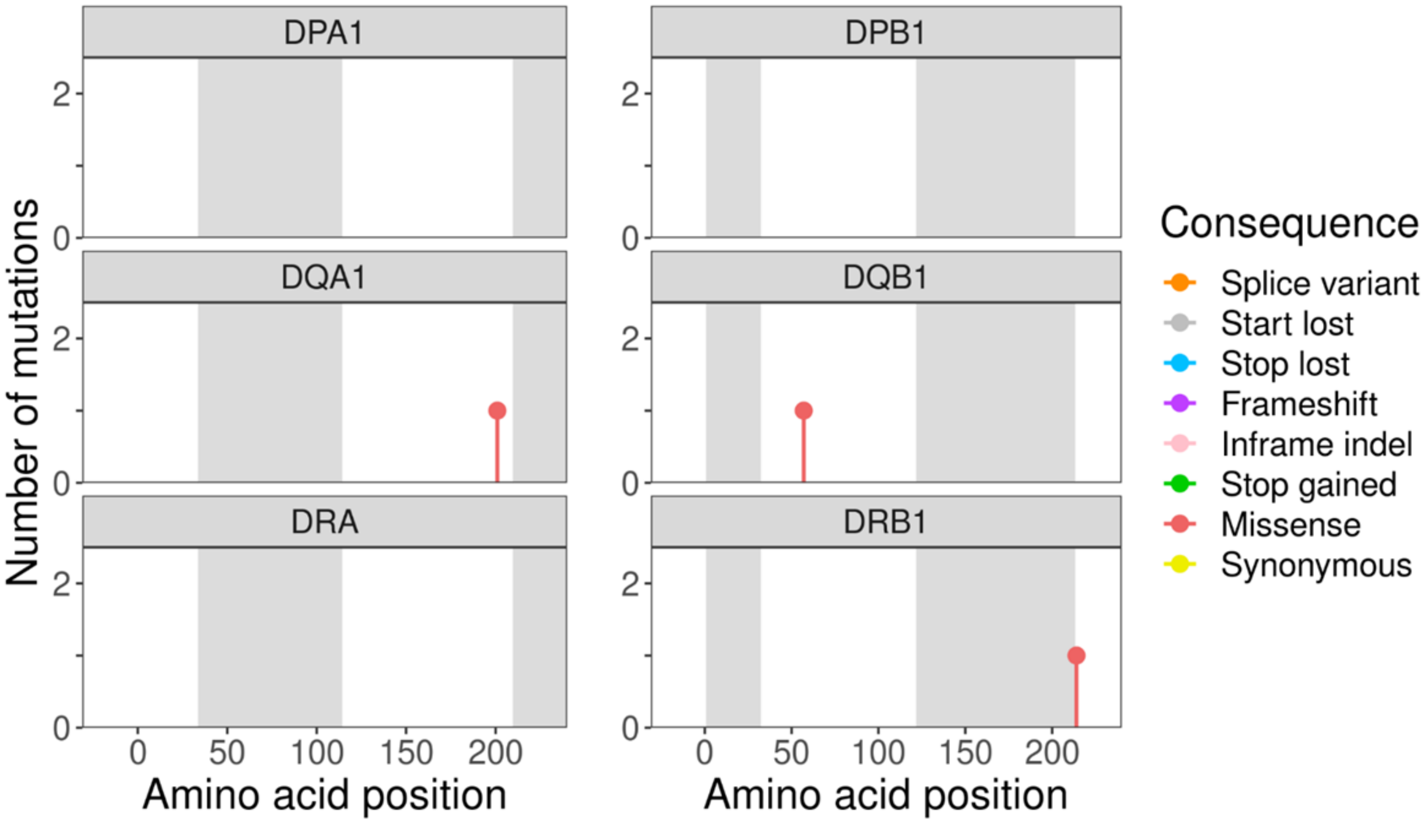
DPA1
DPB1
DQA1
DQB1
DRA
DRB1
Number of mutations
Amino acid position
0
50
100
150
200
Consequence
Splice variant
Start lost
Stop lost
Frameshift
Inframe indel
Stop gained
Missense
Synonymous

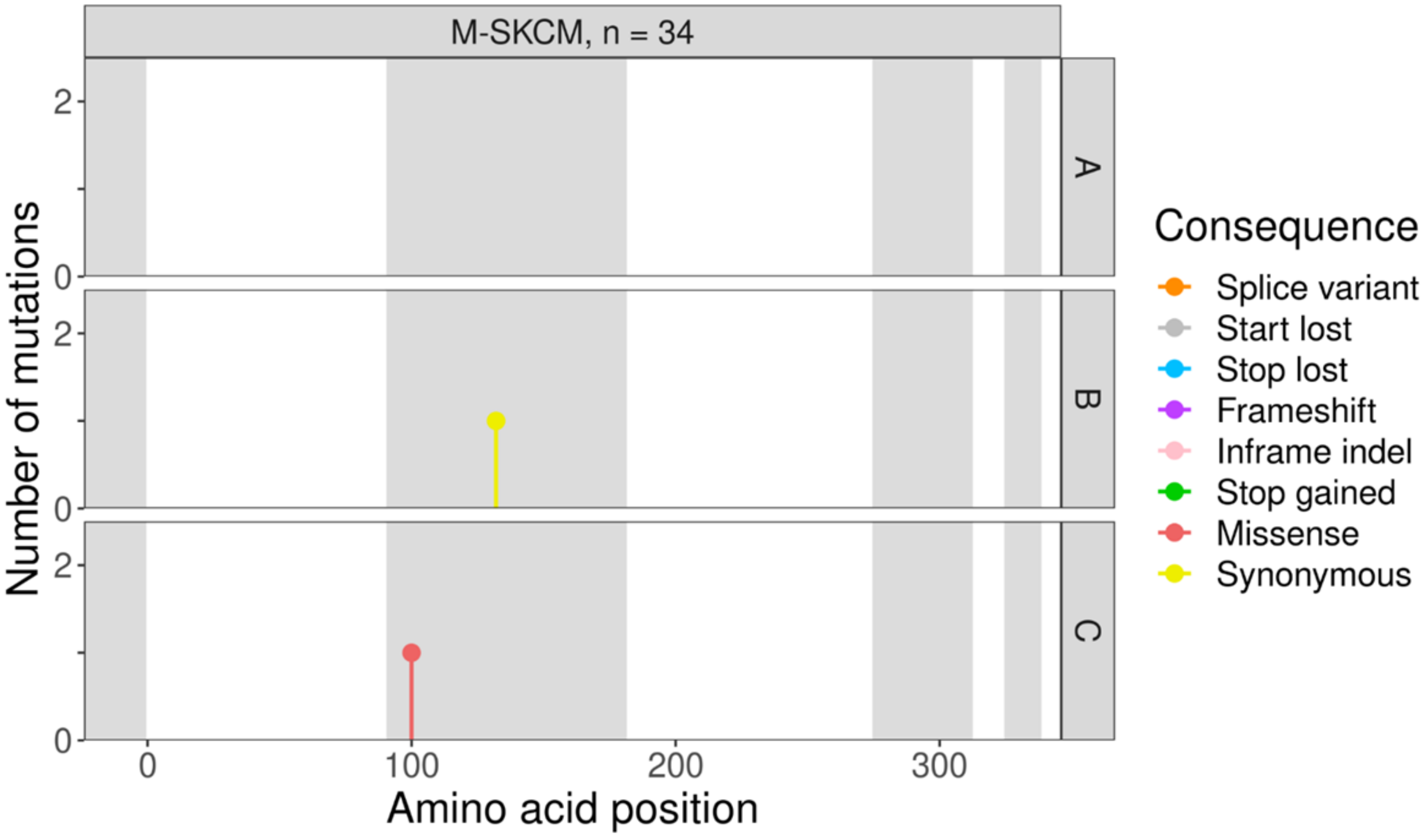
M-SKCM, n = 34
A
B
C
Number of mutations
Amino acid position
0
100
200
300
Consequence
Splice variant
Start lost
Stop lost
Frameshift
Inframe indel
Stop gained
Missense
Synonymous

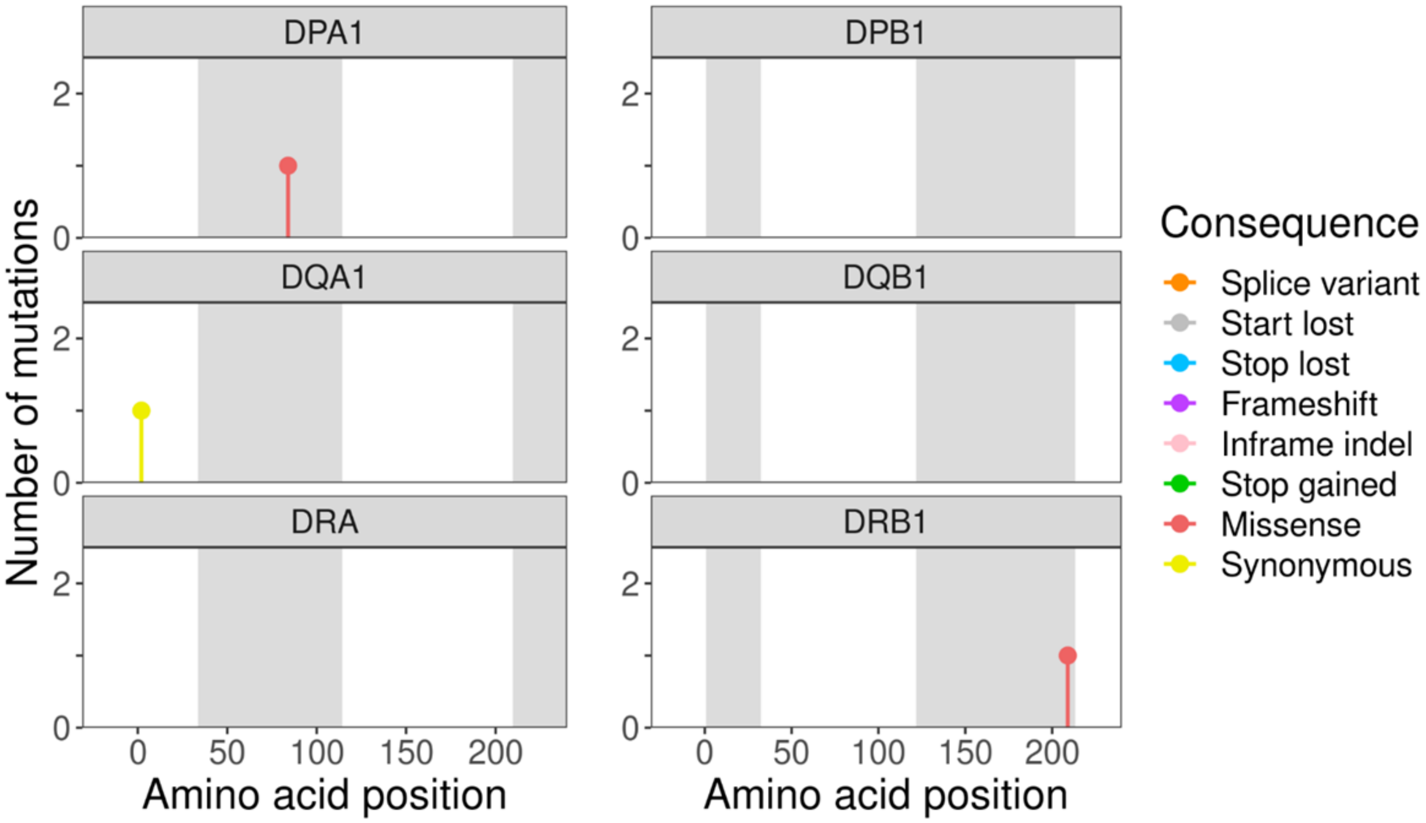
DPA1
DPB1
DQA1
DQB1
DRA
DRB1
Number of mutations
Amino acid position
0
50
100
150
200
Consequence
Splice variant
Start lost
Stop lost
Frameshift
Inframe indel
Stop gained
Missense
Synonymous

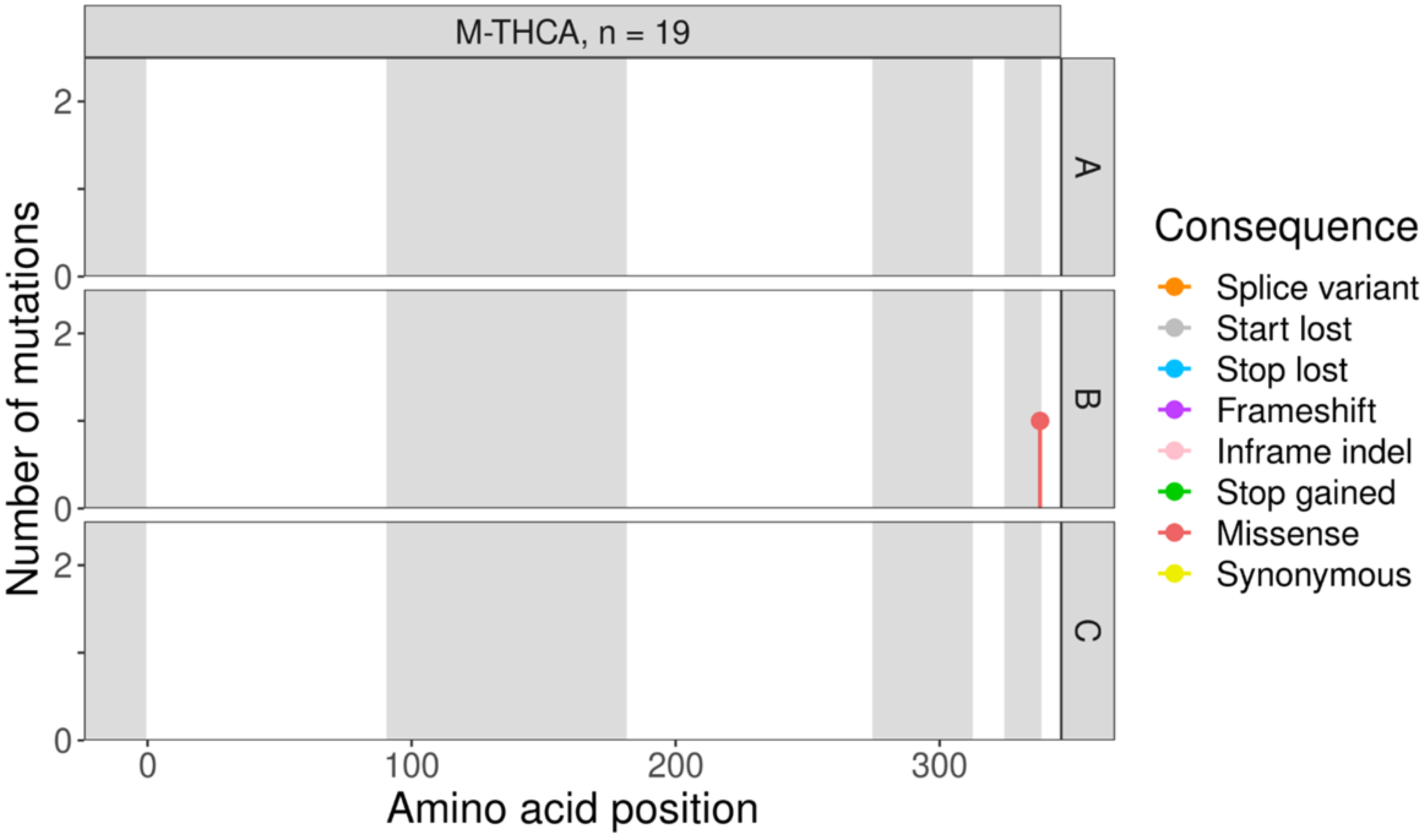
M-THCA, n = 19
A
B
C
Number of mutations
Amino acid position
Consequence
Splice variant
Start lost
Stop lost
Frameshift
Inframe indel
Stop gained
Missense
Synonymous

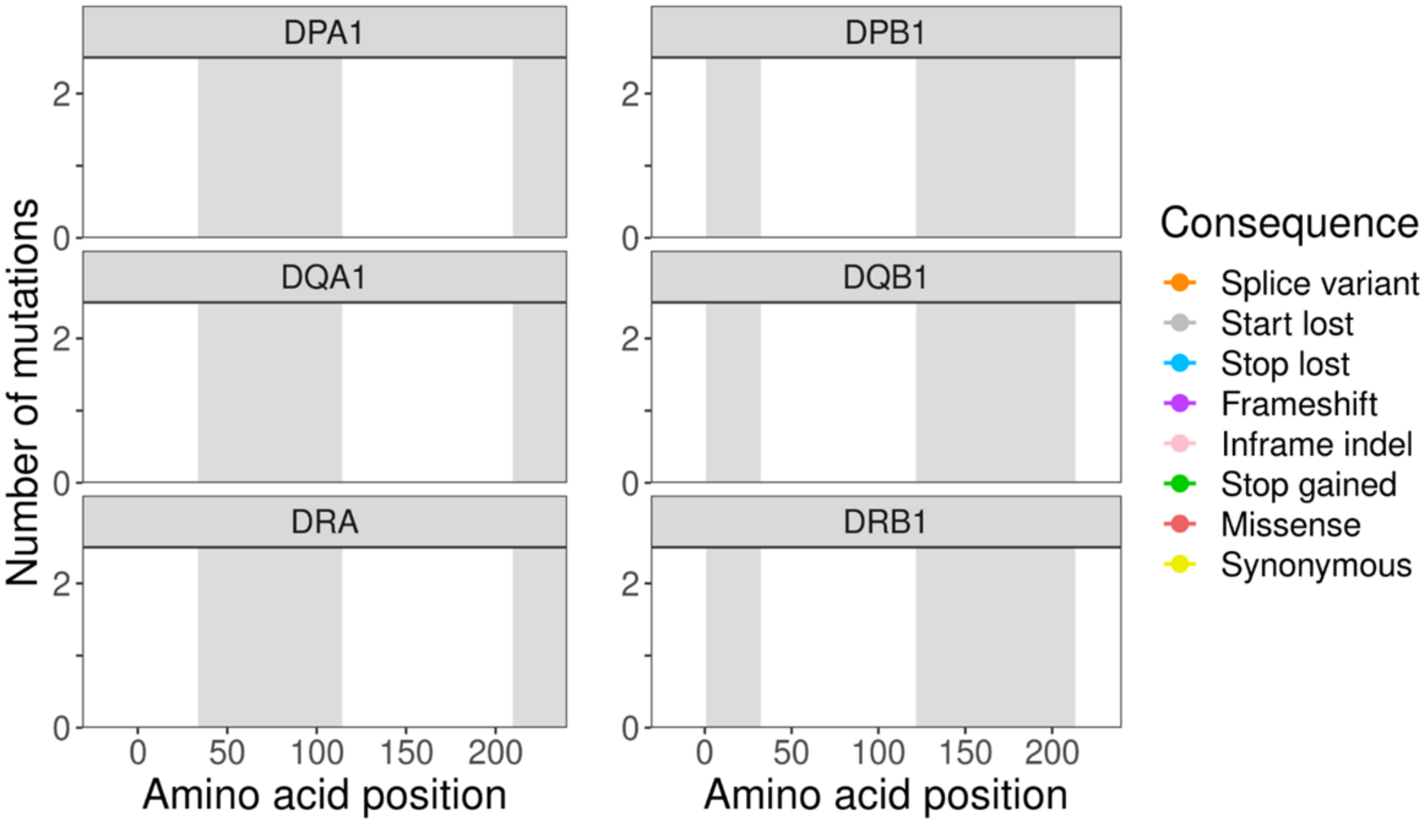
DPA1
DPB1
DQA1
DQB1
DRA
DRB1
Number of mutations
Amino acid position
Consequence
Splice variant
Start lost
Stop lost
Frameshift
Inframe indel
Stop gained
Missense
Synonymous

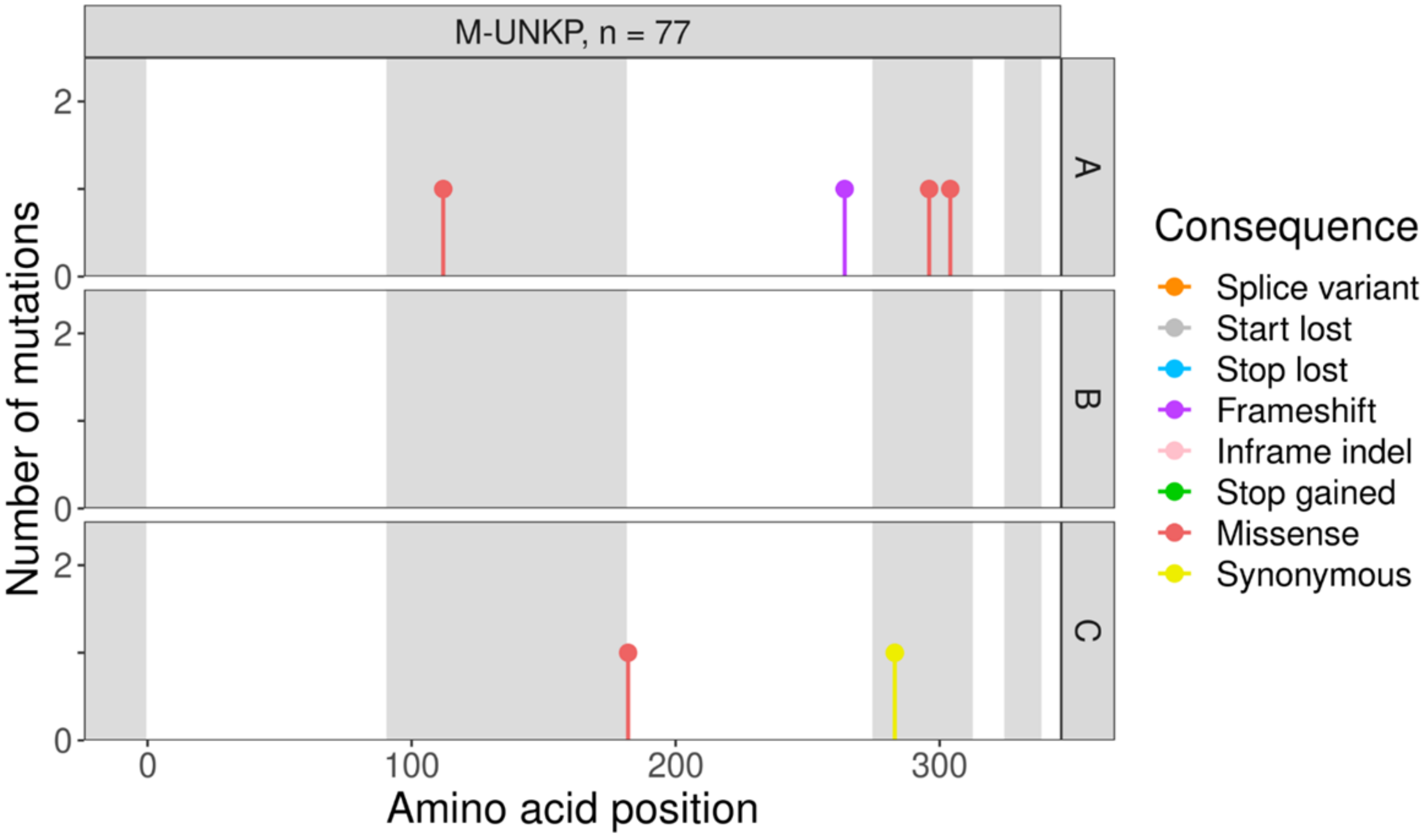
M-UNKP, n = 77
Number of mutations
A
B
C
Amino acid position
Consequence
Splice variant
Start lost
Stop lost
Frameshift
Inframe indel
Stop gained
Missense
Synonymous

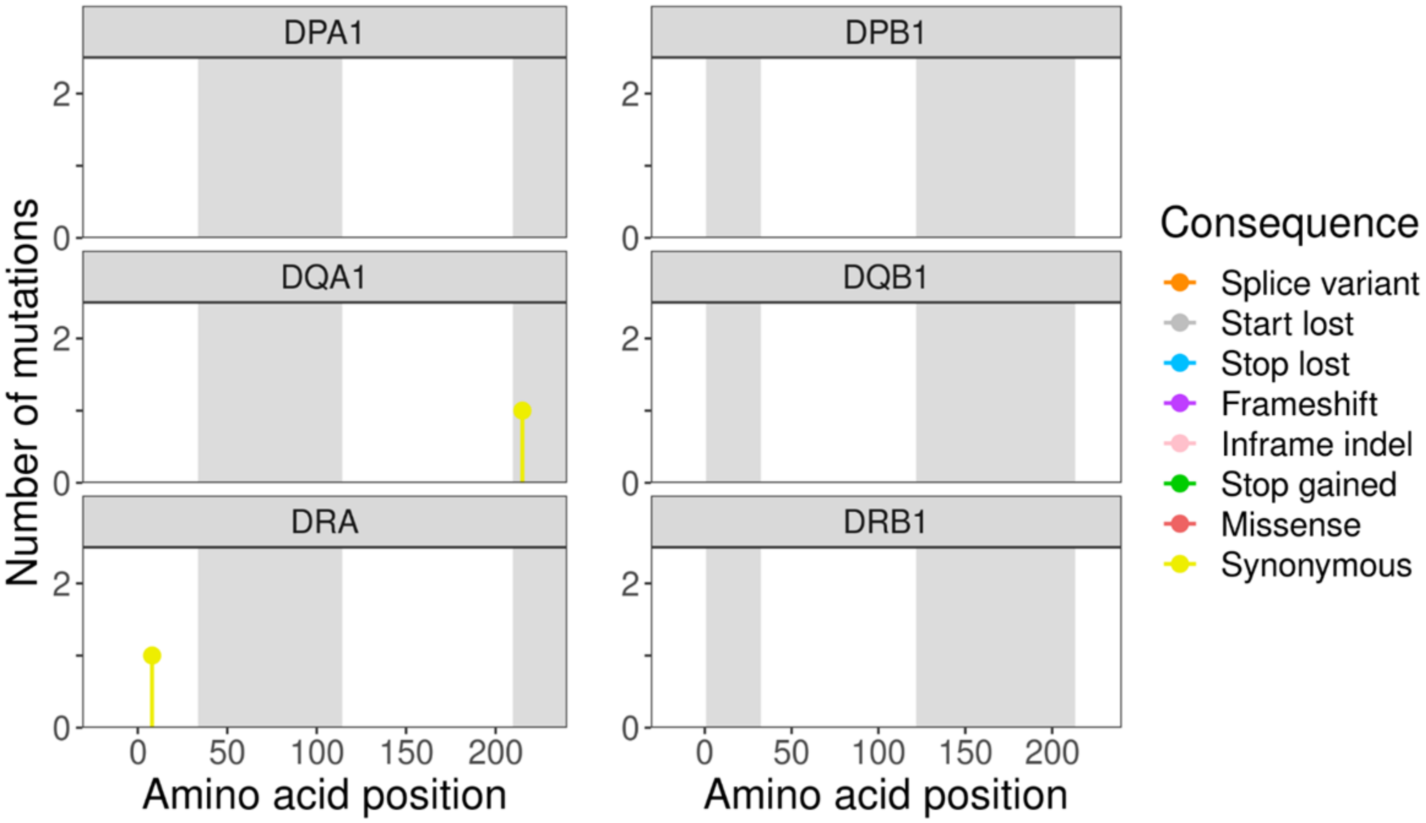
DPA1
DPB1
DQA1
DQB1
DRA
DRB1
Number of mutations
Amino acid position
Amino acid position
Consequence
Splice variant
Start lost
Stop lost
Frameshift
Inframe indel
Stop gained
Missense
Synonymous
